# Enzymatic epimerization of monoterpene indole alkaloids in Kratom

**DOI:** 10.1101/2024.12.13.628308

**Authors:** Allwin McDonald, Yoko Nakamura, Carsten Schotte, Kin Lau, Ryan Alam, Adriana A. Lopes, C. Robin Buell, Sarah O’Connor

## Abstract

Monoterpene indole alkaloids (MIAs) are a large, structurally diverse class of bioactive natural products. These compounds are biosynthetically derived from a stereoselective Pictet-Spengler condensation that generates a tetrahydro-β-carboline scaffold characterized by a 3*S* stereocenter. However, a subset of MIAs contain a non-canonical 3*R* stereocenter. Herein, we report the basis for 3*R*-MIA biosynthesis in *Mitragyna speciosa* (Kratom). We discover the presence of the iminium species, 20*S*-3-dehydrocorynantheidine, which led us to hypothesize that isomerization of 3*S* to 3*R* occurs by oxidation and stereoselective reduction downstream of the initial Pictet-Spengler condensation. Isotopologue feeding experiments implicated young leaves and stems as the sites for pathway biosynthesis, facilitating the identification of an oxidase/reductase pair that catalyzes this epimerization. This enzyme pair has broad substrate specificity, suggesting that the oxidase and reductase may be responsible for the formation of many 3*R*-MIAs and downstream spirooxindole alkaloids in Kratom. These enzymes allow biocatalytic access to a range of previously inaccessible pharmacologically active compounds.

## Introduction

Monoterpene indole alkaloids (MIAs) are a class of structurally diverse and pharmacologically important plant-derived natural products. This natural product family includes vinblastine (anti-cancer), ajmalicine (antihypertensive),^1–3^ strychnine (infamous toxin and pesticide),^4^ mitragynine (pain relief/psychedelic),^5^ and ibogaine (candidate for opioid withdrawal treatment) (Figure 1a).^6,7^ A highly conserved feature of MIA biosynthesis^4,8–10^ is the stereoselective Pictet-Spengler condensation between the aldehyde secologanin and tryptamine, catalyzed by the enzyme strictosidine synthase (STR). Strictosidine, the resulting tetrahydro-β-carboline product, is characterized by a 3*S* stereocenter, the stereochemical configuration observed in the vast majority of MIAs (Figure 1a). More rarely, however, MIAs possess a 3*R* stereocenter (Figure 1b), a subset which includes reserpine (an adergenic blocking agent previously approved to treat high blood pressure),^11^ hirsutine (a potent antiarrhythmic and vasodilator),^12,13^ and speciociliatine (a more potent µ-opioid receptor agonist than the corresponding 3*S* epimer, mitragynine).^14^ 3*R*-MIAs are concentrated within the *Naucleeae* tribe of the *Rubiaceae* plant family,^15^ in species such as *Mitragyna speciosa*,^16^ *Uncaria sp*.,^17–19^ *Pausinystalia yohimbe*,^20^ and *Cephalanthus occidentalis*.^21^ The mechanism by which this 3*R* stereocenter is formed is unknown.

**Figure 1.**
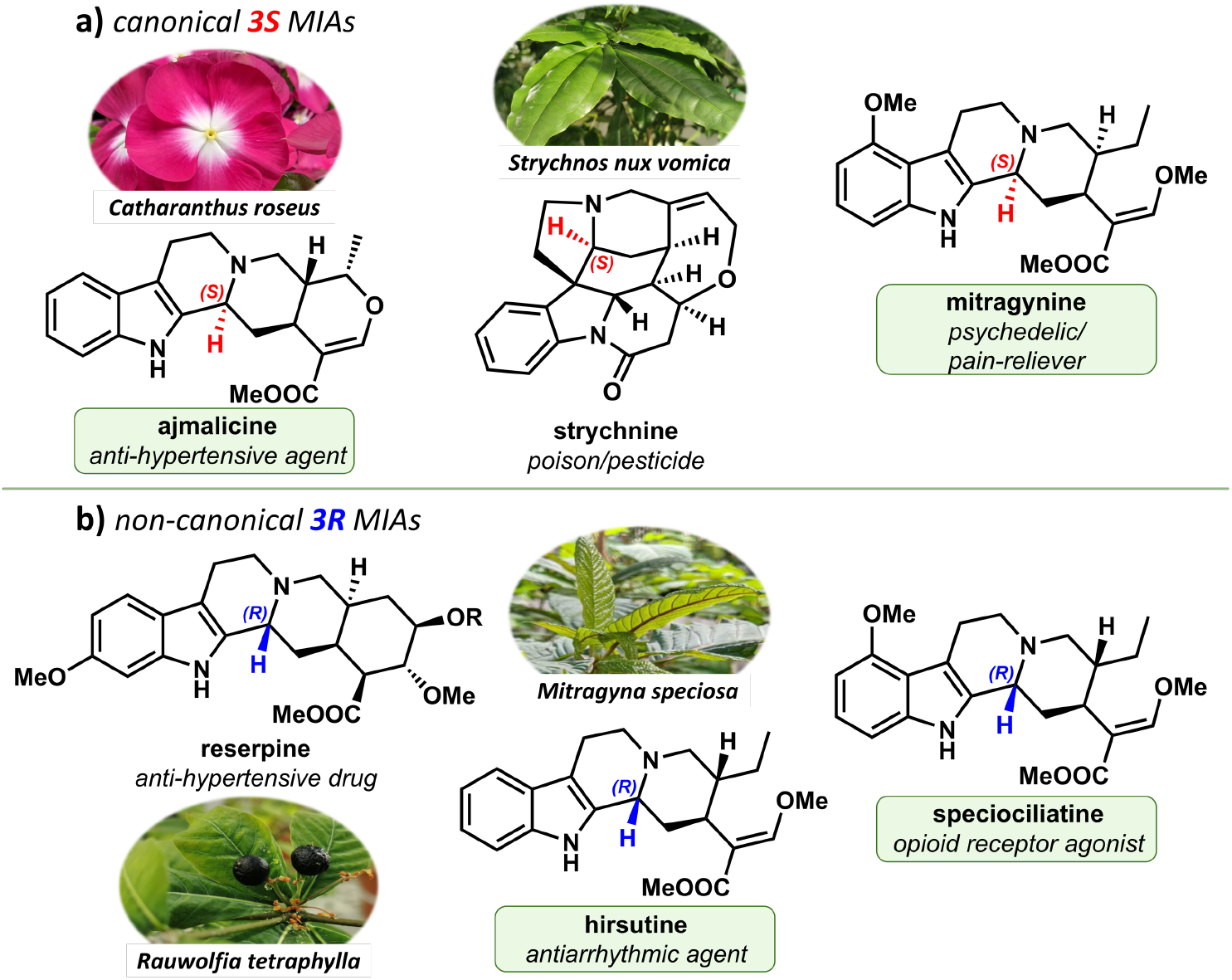
Examples of monoterpene indole alkaloids (MIAs) with the corresponding plant producers. Alkaloids observed in *Mitragyna speciosa* (Kratom) are highlighted in green. **a**) MIAs possessing the canonical 3*S* stereocenter (3-position highlighted in red); **b**) MIAs possessing the non-canonical 3*R* stereocenter (3-position highlighted in blue). Reserpine, *R = - (O)C-Ph(OMe)*_*3*_

Kratom (*M. speciosa*) has attracted wide interest for its use in managing chronic pain and its potential in treating opiate withdrawal. Its pharmacological properties are derived from both the 3*S*-(*e*.*g*. mitragynine (**5a**)) and 3*R*-(*e*.*g* speciociliatine (**7a**)) MIAs found in this plant (Figure S1, S2).^5,14^ The biosynthetic pathway of one Kratom MIA, 20*S*-corynantheidine (**3a**, 3*S*), has been elucidated (Figure 2).^9,22^ Briefly, strictosidine (**2a**) is formed via Pictet-Spengler condensation of tryptamine (**1**) and secologanin by *Ms*STR, deglucosylation occurs via action of SGD, and the resulting aglycone is reduced via *Ms*DCS isoforms to produce dihydrocorynantheine (both 20*S* and 20*R* isomers). *Ms*EnolMT catalyzes an unusual enol methylation, resulting in corynantheidine (**3a/3b**), which is hypothesized to then be derivatized to form a variety of downstream alkaloids (Figure 2). Kratom additionally accumulates a range of corynanthe- and heteroyohimbine-type 3*R*-MIAs, which, in addition to having important pharmacological properties, are postulated to serve as precursors for a plethora of biologically active spirooxindole alkaloids, such as mitraphylline (**12b** (3*S*, 7*R*), a potential treatment for Parkinson’s disease).^18,21,23–25^ A cytochrome P450 enzyme, spirooxindole alkaloid synthase (*Ms*SAS, also known as *Ms*3eCIS), is reported to produce spirooxindole alkaloids from 3*R*-MIAs **6b** and **6c**.^26^

**Figure 2.**
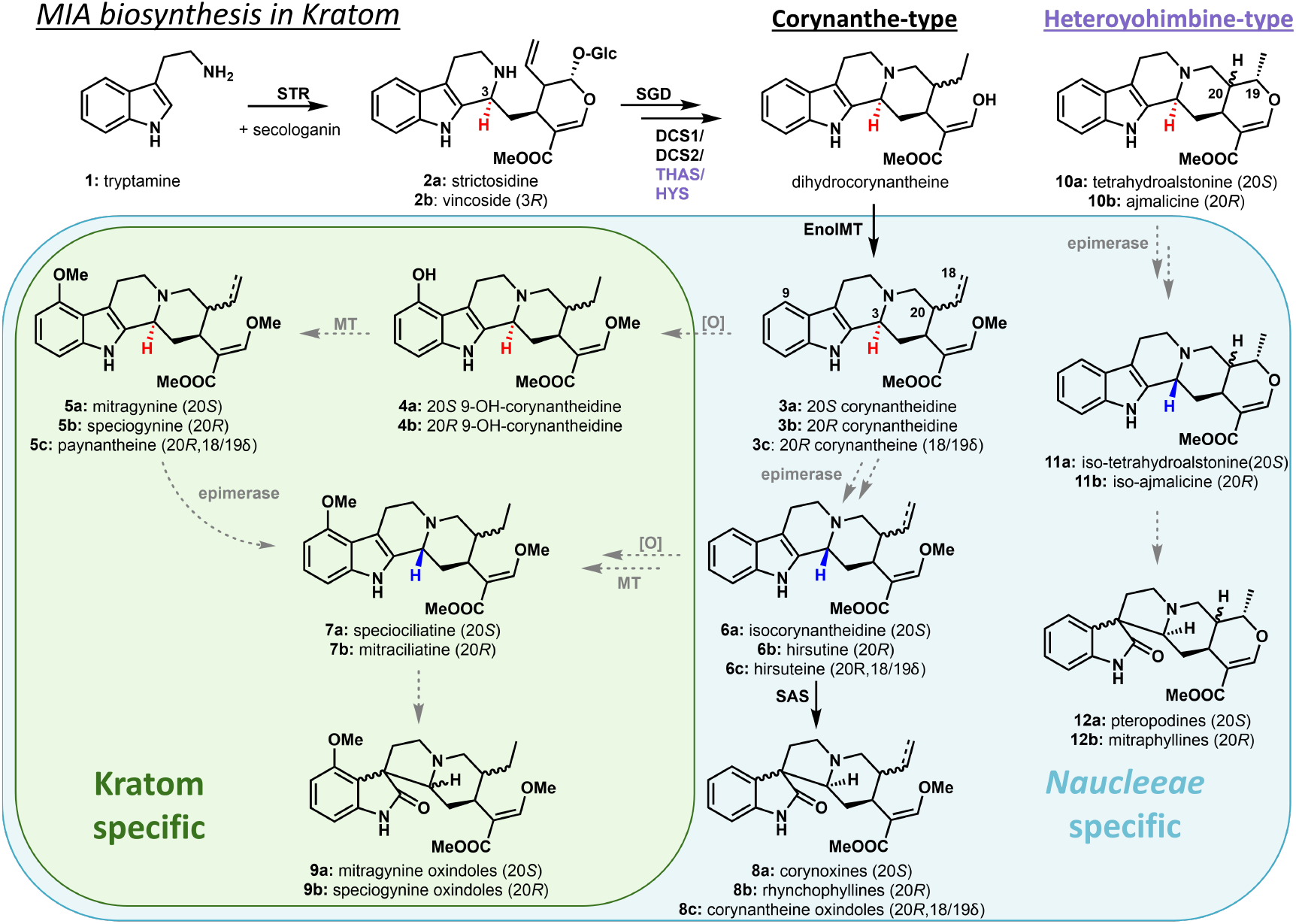
Kratom MIA biosynthetic network. Known transformations depicted with black arrows, unknown or unverified transformations depicted with gray dashed arrows, 3-*S* stereocenters are shown in red, and 3-*R* stereocenters are shown in blue. Enzyme abbreviations: STR = strictosidine synthase, SGD = strictosidine β-D-glucosidase, DCS = dihydrocorynantheine synthase, THAS = tetrahydroalstonine synthase, HYS = heteroyohimbine synthase, EnolMT = enol-methyltransferase, SAS = spirooxindole alkaloid synthase, MT = unknown methyltransferase, [O] = unknown oxidase.

Here, we report the elucidation of the enzymatic steps responsible for biosynthesis of 3*R*-MIAs in Kratom. Isolation of a previously unreported Kratom biosynthetic intermediate led us to the hypothesis that epimerization of the 3*S* center occurs on corynantheidine (**3a/3b**). Feeding of isotopically labelled substrates to Kratom tissues indicated that epimerization occurs in only specific tissues, an observation that facilitated the identification of an oxidase (*Ms*CO)/reductase (*Ms*DCR) pair responsible for epimerization of corynantheidine (**3a/3b**) to 3*R*-corynantheidine (**6a/6b**). Additionally, we demonstrate that the resulting 3*R*-MIAs can be enzymatically converted into spirooxindole alkaloids by the cytochrome P450 *Ms*SAS. Furthermore, we established that *Ms*CO, *Ms*DCR and *Ms*SAS each have broad substrate specificity, suggesting that these three enzymes are collectively responsible for biosynthesis of many *3R*-MIAs and spirooxindole alkaloids in Kratom.

## Results

### Discovery of new Kratom MIA iminium species

Kratom tissues were subjected to metabolic profiling. A diversity of alkaloids, including 3*S-*MIAs (*e*.*g*. 20*S*-corynantheidine (**3a**) and mitragynine (**5a**)) as well as 3*R*-MIAs (*e*.*g*. speciociliatine (**7a**)), were found in young leaves (Figure S3, S4). In contrast, mature leaves contained primarily 3*S*-MIAs (Figure S5). Stems and roots contained greater amounts of 3*R*-MIAs and spirooxindole alkaloids (Figure S6, S7). In young leaf extracts, we noticed two especially abundant peaks (*m/z* [M+H]^+^ = 367 and *m/z* [M+H]^+^ = 397) that did not match any available MIA standard (Figures 3A). Since these masses were consistent with an oxidized MIA derivative, we incubated the young leaf extract with the reducing agent NaBH_4_, which resulted in efficient conversion to either corynantheidine (**3a**) or mitragynine (**5a**). Compounds purified from young leaf extracts were shown to be the iminium species 20*S*-3-dehydrocorynantheidine (DHC, **13a**) and 3-dehydromitragynine (DHM, **14a**) by NMR analysis (Figure 3a, Figure S8). Both iminium species showed minimal decomposition during purification and were stereoselectively reduced by NaBH_4_ to the respective 3*S* epimers (Figure S8). DHM (**14a**) had been previously reported to be present in Kratom leaves,^27^ but DHC (**13a**) has not been previously reported. The presence of these compounds led us to hypothesize that 3*R*-MIAs could be formed via stereoselective reduction of one or both of these iminium species.

**Figure 3.**
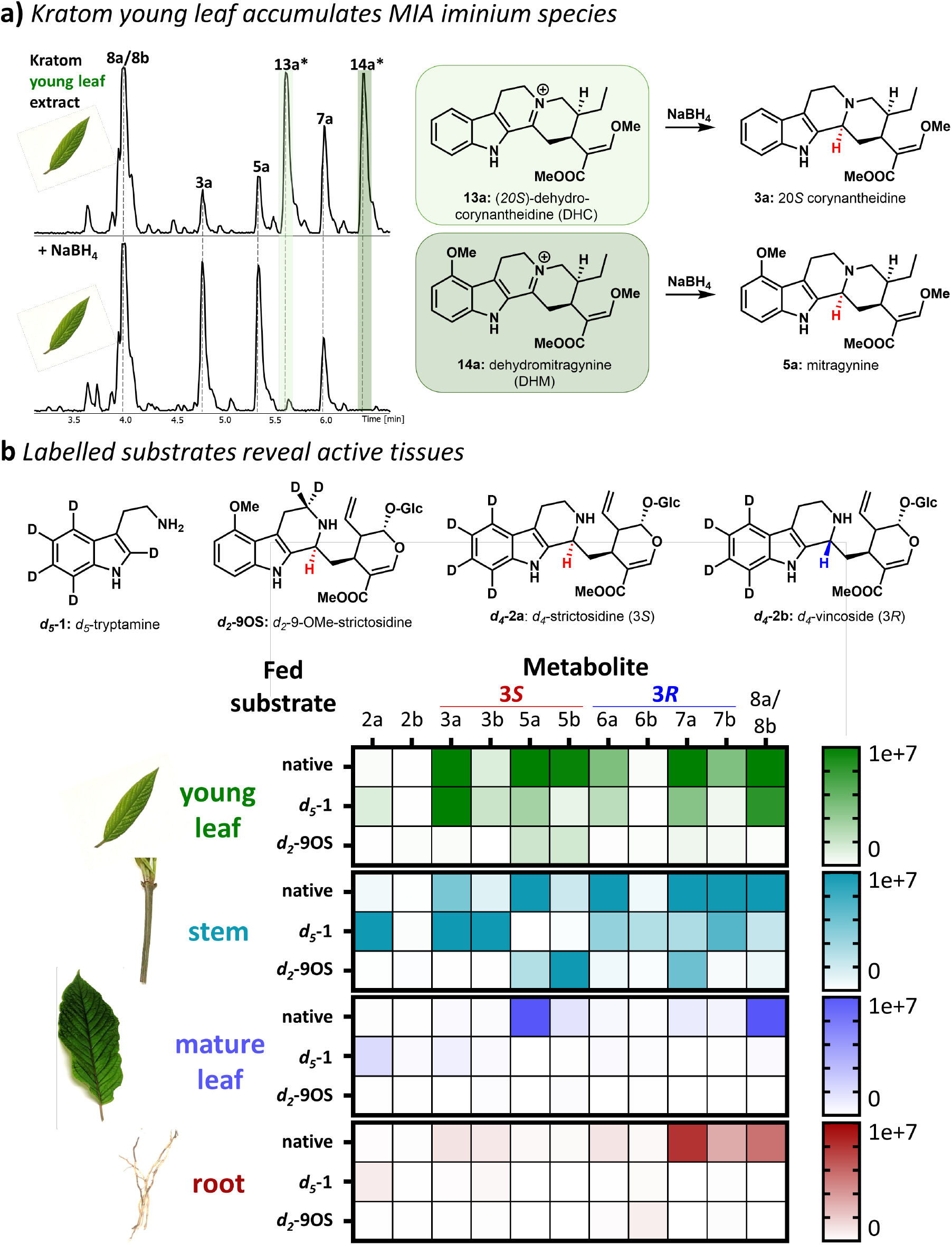
**A)** Comparison of Kratom leaf extract with and without added NaBH_4_. Peaks corresponding to 20*S*-3-dehydrocorynantheidine (**13a**, DHC) and dehydromitragynine (**14a**, DHM) are highlighted. These compounds were isolated and characterized by NMR (Figures S56-S61). **B)** Native metabolite distribution and metabolite incorporation from fed labeled precursors. Note that values for fed tissues are scaled 2x (***d***_***5***_**-1**) or 5x (***d***_***2***_**-9OS**) in the above heat map for better visualization (Table S1). *d*_*4*_-Strictosidine (***d***_***4***_**-2a**) and *d*_*4*_-vincoside (***d***_***4***_**-2b**) feeding results can be found in Figure S11.

### Isotopologue feeding to identify metabolically active Kratom tissue

Kratom MIA biosynthesis appears to involve transport of MIA intermediates between tissues,^23^ a scenario that complicates biosynthetic gene discovery. Known pathway enzymes are preferentially expressed in roots,^9^ although mitragynine (**5a**) and related isomers accumulate predominantly in leaves and stems. To deconvolute metabolite and gene location, we fed labeled MIA isotopologues to cuttings of Kratom tissues: young/mature leaf disks, green stem cuttings, and cut roots (Figure 3b, Table S1). Young leaves and stems produced substantial levels of labeled MIAs from *d*_*5*_-tryptamine (***d***_***5***_**-1**, Figure 3b, S9-S10), most notably labelled mitragynine (**5a**, 3*S*) and speciociliatine (**7a**, 3*R*), demonstrating that these tissues harbor the genes responsible for establishing the 3*R* stereochemistry. Interestingly, some metabolites, such as hirsutine (**6b**, 3*R*), were not detected in untreated stem tissue, but were observed after isotopologue feeding, highlighting potential discrepancies between gene expression and native metabolite profiles. Mature leaves contained large amounts of mitragynine (**5a**, 3*S*) and corynantheidine oxindoles (**8a/8b**, 3*R* derived), but labelled substrates were not incorporated to significant levels in this tissue. Roots primarily contained speciociliatine (**7a**, 3*R*) and likewise showed minimal isotopologue incorporation. Feeding of *d*_*4*_-strictosidine (***d***_***4***_**-2a**, 3*S*) resulted in similar incorporation as observed for *d*_*5*_-tryptamine (***d***_***5***_**-1**), but with lowered efficiency (Figure S11). The 3*R* epimer of strictosidine, *d*_*4*_-vincoside (***d***_***4***_**-2b**, 3*R*), was not incorporated in any tissue, suggesting that epimerization happens at a later biosynthetic stage. Feeding of *d*_*2*_-9-OMe-strictosidine (***d***_***2***_**-9OS**, 3*S*) produced labelled mitragynine (**5a**, 3*S*) and speciogynine (**5b**, 3*S*) in both stem and young leaf (Figure 3b). However, only small amounts of labelled speciociliatine (**7a**, 3*R*) were observed (stems), suggesting that epimerization does not occur efficiently on C9 methoxylated intermediates. Additionally, feeding of ***d***_***5***_**-1** into stems resulted in significant **7a**/**7b** labeling, but little **5a**/**5b** labeling, which may indicate that **5a**/**5b** are not precursors to **7a**/**7b**. These data suggest that epimerization occurs in young leaves after formation of strictosidine (**2a**), but prior to methoxylation at C9. Along with the identification of the iminium moiety DHC (**13a**), these observations led us to hypothesize that corynantheidine (**3a**) could be a primary substrate for epimerization.

### Identification and characterization of epimerase genes

Using a coupled feeding/RNA extraction procedure, we extracted RNA from young leaf and green stem tissue that showed robust conversion of *d*_*5*_-tryptamine (***d***_***5***_**-1**) to speciociliatine (**7a**, 3*R*). For comparison, we also isolated RNA from mature leaf tissue that did not incorporate *d*_*5*_-tryptamine (***d***_***5***_**-1**). From the resultant RNA-sequencing data, we searched for gene candidates that could be responsible for the oxidation of the proposed substrate, 20*S*-corynantheidine (**3a**), to the iminium species DHC (**13a**). We noted that in the biosynthetically unrelated alkaloid morphine, isomerization of a key stereocenter of the tetrahydroisoquinoline scaffold also occurs via formation and reduction of an iminium intermediate (from *S*-reticuline via 3-dehydroreticuline to *R*-reticuline). In this case, a fusion protein consisting of a cytochrome P450 linked to an aldo-keto reductase performs both the oxidation and reduction.^28,29^ Taking inspiration from this transformation, we targeted genes encoding cytochromes P450 that were co-expressed with the pathway gene *Ms*EnolMT in young leaves, though gene candidates annotated as polyphenol oxidases and berberine bridge-like enzymes (BBE-like) were also considered. Oxidase gene candidates were agroinfiltrated into *Nicotiana benthamiana* along with 20*S*-corynantheidine (**3a**). This screening approach resulted in identification of two BBE-like enzymes that catalyzed the oxidation of 20*S*-corynantheidine (**3a**) to DHC (**13a**) (Figure 4a, Figure S12a). No activity was observed when mitragynine was used as a substrate (Figure S13), suggesting an alternate route to DHM (**14a**) production in leaves (Figure 4b). Thus, these enzymes were termed corynantheidine oxidases 1-2 (CO1, CO2). *Ms*CO1 and *Ms*CO2 share 94% sequence identity and were coexpressed with *Ms*EnolMT in young leaf tissue (Figure 4b, Figure S12b). No differences in substrate specificity were noted among these isoforms from *in planta* assays; therefore, *Ms*CO1 was used for all subsequent assays. Negative (empty vector (EV)) controls indicated that native *N. benthamiana* enzymes could oxidize 20*S*-corynantheidine (**3a**) to DHC (**13a**), although with low efficiency (Figure 4a).

**Figure 4.**
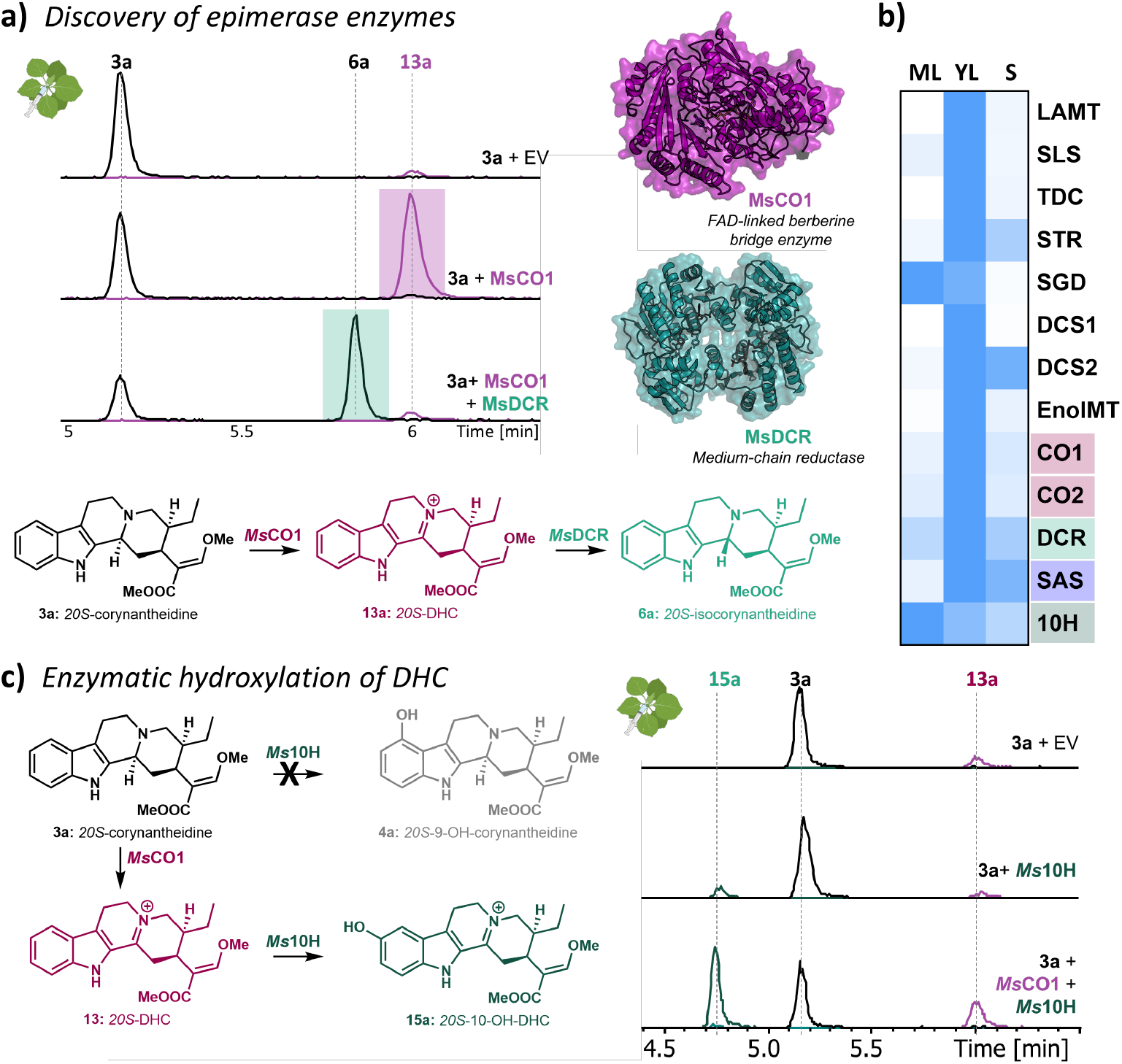
**a)** *Ms*CO1 and *Ms*DCR epimerize 20*S*-corynantheidine (**3a**, 3*S*) to 20*S*-isocorynantheidine (**6a**, 3*R*) via iminium intermediate **13a**. Enzyme activities assessed via agroinfiltration in *N. benthamiana*. Protein structures generated using AlphaFold3. **b)** Coexpression matrix of Kratom pathway enzymes in mature leaves (ML), young leaves (YL), and stems (S). **c)** Activity of a newly discovered CYP81, *Ms*10H, hydroxylates 20*S-*DHC to form **15a**.

Reductase candidates were initially chosen based on co-expression with *Ms*CO1 and/or homology to known iminium reductases (alcohol dehydrogenases: *Ms*DCS1, *Cr*THAS1, *Cr*DPAS^30^), but no enzyme that reduced **13a** to 20*S*-isocorynantheidine (**6a**) could be identified. We then compared the reductase genes that are conserved among the known 3*R*-MIA producing species of the *Naucleeae* tribe of *Rubiaceae* (Kratom, *Uncaria guianensis*, and *Uncaria rhynchophylla*) with the reductases that are found in non-producers outside of the *Naucleeae* (*Rubiaceae: Cinchona pubescens* and *Coffea arabica*; *Apocynaceae*: *Catharanthus roseus*). Sequence similarity networks (SSNs) are a useful visualization tool for analysis of related sequences, and we hypothesized that *Naucleeae*-specific clusters could contain the reductase responsible for stereoselectively reducing DHC (**13a**). To this aim, we built an SSN with every annotated Kratom reductase/dehydrogenase/methyltransferase and the closest homolog found in each of the aforementioned plant transcriptomes. Through increasing the stringency of the node alignment threshold, we observed the emergence of *Naucleeae*-specific clusters containing Kratom gene candidates. As a control, we tested this approach using the known *Naucleeae* pathway-specific methyltransferase gene, *Ms*EnolMT, which appeared in a *Naucleeae*-specific cluster (Figure S14).

Encouraged, we identified nine reductase candidates that appeared in *Naucleeae*-specific clusters (Figure S15). One of these candidates, annotated as an isoflavone reductase, reduced DHC (**13a**) to 20*S*-isocorynantheidine (**6a**) (Figure 4a). This enzyme, here named dehydrocorynantheidine reductase (DCR), belongs to the class of lignan aromatic alcohol dehydrogenases^31^ and shares high sequence identity (73%) to isoflavone reductase in *C. arabica* (Figure S12c). Notably, this enzyme is not related to the other known Kratom MIA iminium reductase, *Ms*DCS1 (8% amino acid sequence identity). Although *Ms*DCR was only identified after SSN analysis, this gene does in fact have a similar expression pattern with the upstream pathway (Figure 4b). Both *Ms*CO1 and *Ms*DCR could be heterologously expressed and purified, and we verified their activities using *in vitro* assays (Figure S16).

### Iminium intermediate DHC is a substrate for 10-hydroxylation

We envisioned that SSN analysis for all annotated Kratom oxidases/P450s and the closest homologs in related *Rubiaceae* and *Apocynaceae* species would be a viable strategy to uncover the hydroxylases hypothesized to be involved in downstream MIA biosynthesis. Since 9- and 10-hydroxylation has been reported in both Kratom and *Uncaria* species^32^, we targeted candidates in *Naucleeae*-specific clusters along with Kratom-specific clusters (Figure S17). Although no candidate gene hydroxylated 20*S*-corynantheidine (**3a**), one CYP81, identified from a *Naucleeae*-specific cluster, catalyzed an unexpected 10-hydroxylation on DHC to form **15a**. This enzyme showed no activity on other Kratom metabolites (Figure 4D, Figure S18-S19). The gene encoding this enzyme, here termed *Ms*10H, was expressed primarily in mature leaves. No 10-hydroxylated MIAs have been observed in Kratom, and the corresponding 10-hydroxy-DHC (**15a**) and reduced 10-hydroxycorynantheidine (**16a**) peaks do not correspond to any observed compounds from Kratom extracts (Figure S18). The *in planta* role of *Ms*10H in Kratom MIA biosynthesis, therefore, remains unclear. Nevertheless, this discovery highlights that the iminium intermediate DHC can serve as a substrate for downstream pathway steps.

### Substrate activity assays for *Ms*CO1, *Ms*DCR, and *Ms*SAS

In Kratom and related plants, it is hypothesized that only MIAs with 3*R* stereochemistry can be converted to spirooxindole alkaloids. A recently discovered P450, *Ms*SAS (also known as *Ms*3eCIS), was shown to convert hirsutine (**6b**, 3*R*, 20*R*) and hirsuteine (**6c**, 3*R*, 20*R*) to spirooxindole alkaloids **8b** and **8c**,^26^ though additional substrates were not tested. We sought to determine the substrate scope of 3*S*-MIA to 3*R*-MIA epimerization, as well as spirocyclization of formed 3*R*-MIAs. To this aim, we assayed a variety of substrates with *Ms*CO1, *Ms*DCR, and *Ms*SAS in *N. benthamiana* (Figures 5, S20-S31, S34) to assess how these enzymes functioned *in planta*.

**Figure 5.**
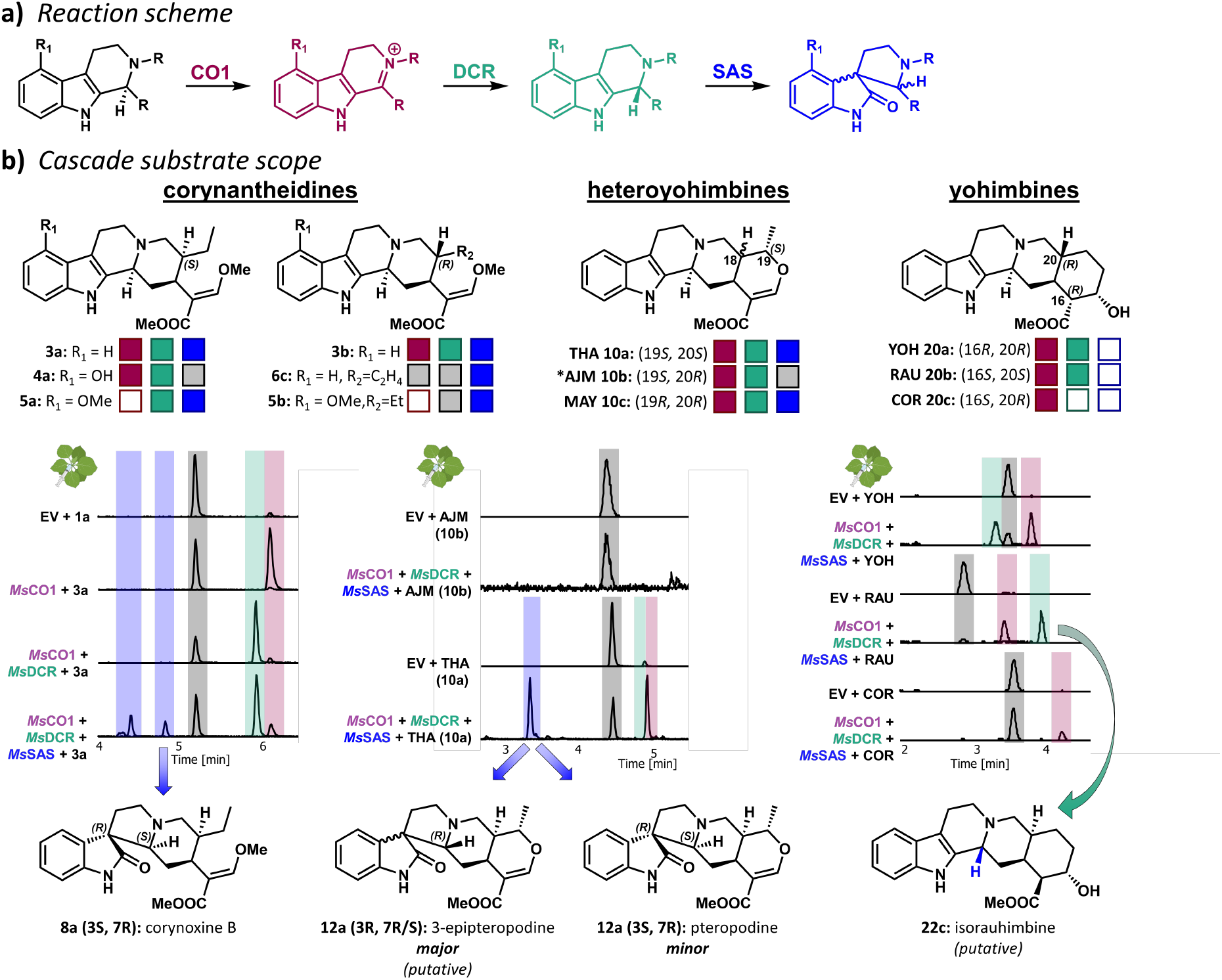
**a)** Investigation of substrate scope of *Ms*CO1, *Ms*DCR and *Ms*SAS. **b)** Tested substrates for epimerization: THA = tetrahydroalstonine, AJM = ajmalicine, MAY = mayumbine, YOH = yohimbine, RAU = rauwolscine, COR = corynanthine. A colored square indicates an observed transformation for a given substrate, an empty square indicates that no reaction was observed, and a gray square indicates that a given transformation could not be tested. *For ajmalicine (**10b**), activity was observed *in vitro*, but not *in planta*. (Figure S27). For HPLC traces, gray bars indicate starting material [*m/z*] extracted ion chromatogram (EIC), red bars indicate [*m/z* -2] EIC, green bars indicate [*m/z*] EIC, and blue bars indicate [*m/z* + 16]. See Figures S19-S31, S34 for detailed analyses of all tested reactions.

Hirsuteine (**6c**) has previously been reported to be a substrate for *Ms*SAS,^26^ and reaction *in planta* with this substrate resulted in two 3*S* spirocyclic products: isocorynoxeine (**8c** (3*S*, 7*S*)) and corynoxeine (**8c** (3*S*, 7*R*)) (Figures 5b, S21). We then tested the activity of *Ms*SAS in the context of the newly discovered upstream enzymes *Ms*CO1 and *Ms*DCR on 20*S*-corynantheidine (**3a**), and found that this substrate was converted to a mixture of three corynoxine isomers (**8a**), including corynoxine A (**8a** (3*S*, 7*S*)) and corynoxine B (**8a** (3*S*, 7*R*)) (Figure S22). A compound putatively assigned as 3-epicorynoxine (**8a** (3*R*, 7*R/S*)) was also observed as a major product. Consistent with previous reports of spirooxindole alkaloid interconversion, we noted that produced corynantheidine oxindoles (and standards) spontaneously isomerize.^33^ Lack of availability of 20*R*-corynantheidine (**3b**) precluded direct assay of this substrate.

20*S*-9-Hydroxycorynantheidine (**4a**), a presumed intermediate *en route* to mitragynine (**5a**), appeared to be readily oxidized by *Ms*CO1, but only traces of reduced product were observed with the subsequent reaction with *Ms*DCR (Figure S23). Neither mitragynine (**5a**, 20*S*) nor speciogynine (**5b**, 20*R*) were oxidized by *Ms*CO1, but DHM (**14a**) was reduced via *Ms*DCR to speciociliatine (**7a** (3*R*, 20*S*) Figure S24). Both speciociliatine (**7a**) and mitraciliatine (**7b**) act as substrates for *Ms*SAS, forming compounds with mass fragmentation patterns consistent with mitragynine oxindoles (**9a**) and speciogynine oxindoles (**9b**), respectively (Figures S24-S25). Peaks coeluting with these observed methoxylated spirooxindole alkaloid products were highly abundant in stem and root tissue.

Heteroyohimbine-type MIAs such as tetrahydroalstonine (THA) (**10a**) and ajmalicine (**10b**) are also observed in Kratom (Figure 2). When *Ms*CO1, *Ms*DCR, and *Ms*SAS were assayed with THA (**10a**), we observed formation of minor amounts of isopteropodine (**12a** (3*S*, 7*S*)) and pteropodine (**12a** (3*S*, 7*S*)), with the major peak likely corresponding to one of the two possible 3*R* **12a** isomers (3-epipteropodine) (Figure S26). Additional compounds assigned as dehydro-THA (**19a**) and iso-THA (**11a**) were also observed. In contrast, *Ms*CO1 incubation with ajmalicine (**10b**) did not result in observable product formation *in planta* (Figure 5b), although *Ms*CO1 and *Ms*DCR were shown to be active on ajmalicine (**10b**) *in vitro*, forming 3-epi-ajmalicine (Figure S27). The related heteroyohimbine mayumbine (**10c**) was also carried through the cascade, resulting in the formation of compounds that could be putatively assigned as mayumbine oxindoles (**12c**), alongside intermediates dehydro-mayumbine (**19c**) and isomayumbine (**11c**) (Figure S28).

Finally, we tested the activity of *Ms*CO1, *Ms*DCR, and *Ms*SAS on yohimbine-type alkaloids (Figure 5b), even though yohimbanes are not present in Kratom. Yohimbine (**20a**) was readily converted by both *Ms*CO1 and *Ms*DCR to form pseudoyohimbine (**22a**) (Figure S29). Rauwolscine (**20b**) was similarly isomerized to isorauhimbine (**22b**) (Figure S30), but corynanthine (**20c**) could only be converted to 3-dehydro-corynanthine (**21c**), and no subsequent reduction to isocorynanthine (**22c**) was observed (Figure S31). Interestingly, *Ms*SAS showed no activity on any of these yohimbine-type substrates, highlighting the greater specificity of this P450 for conversion to the spirooxindole alkaloids.

### Pathway construction in *N. benthamiana* from tryptamine

Finally, we reconstituted the full pathway from tryptamine (**1**) to spirooxindole alkaloids (**8a/8b**) by coexpressing Kratom pathway enzymes in *N. benthamiana* (*Ms*STR, *Cr*SGD, *Ms*DCS1/*Cp*DCS, *Ms*EnolMT, *Ms*CO1, *Ms*DCR, and *Ms*SAS), along with infiltration of the starting substrates tryptamine (**1**) and secologanin. Successful stepwise formation of all 20*S* and 20*R* isomers leading to the respective corynantheidine oxindoles (**8a/8b**) was observed though product levels were low (Figure S32). When *Ms*DCS1, which is selective for 20*S* products, was used, corynoxine isomers (**8a**) were observed (Figure S33). To obtain the 20*R* series, we used the DCS reductase from *C. pubescens, Cp*DCS, which selectively produces this epimer.^34^ In this cascade, only two 20*R* products were noted, both with 3*S* stereochemistry corresponding to standards of isorhynchophylline (**8b** (3*S*, 7*S*)) and rhynchophylline (**8b** (3*S*, 7*R*)), Figure S34). Therefore, 20*R*-corynantheidine (**3b**) also acts as a substrate for *Ms*CO1, *Ms*DCR, and *Ms*SAS (Figure 5b).

## Discussion

The mechanism(s) by which the 3*R* stereocenter is generated in MIA biosynthesis was unknown at the outset of this study. Here, we show that Kratom (*Rubiaceae* family, *Naucleeae* tribe) evolved an oxidase/reductase pair to epimerize a variety of corynanthe-, heteroyohimbine-, and yohimbine-type *3S*-MIAs to the corresponding *3R-*MIAs. An analogous strategy of epimerization following a stereoselective Pictet-Spengler reaction has been previously characterized for biosynthesis of the tetrahydroisoquinoline alkaloid morphine. This morphine biosynthetic enzyme, reticuline epimerase, a fused P450-AKR enzyme, carries out both oxidation and reduction. Fusing the epimerase in this way is proposed to limit the reactivity of the dehydro-reticuline iminium intermediate and increases pathway flux.^28,29^ In Kratom, we show that the intermediate iminium species are surprisingly long-lived and decoupled from subsequent reduction, accumulating to a high degree in growing leaf tissue. *Ms*DCR and *Ms*SAS are expressed to high levels in stem, suggesting that iminium species are synthesized in young leaves and then transported to the stems/roots. In support of this hypothesis, both roots and stems primarily accumulate 3*R*-MIAs and downstream 3*R*-derived spirooxindole MIAs, while leaves contain primarily 3*S*-MIAs and iminium MIAs.

Iminium formation from BBE-like enzymes has been previously observed. In a related MIA biosynthetic pathway in *C. roseus*, the related BBE-like enzyme *Cr*PAS (54% identity) produces the conjugated iminium precondylocarpine acetate.^35^ The newly discovered hydroxylase *Ms*10H has activity specific to DHC (**13a**), acting only on this iminium species (**13a**) and not 20*S*-corynantheidine (**3a**). Alongside the high abundance of DHC (**13a**) and DHM (**14a**) in leaf tissue (and presence of **17a** (Figure S35)), this data raises the possibility that DHC (**13a**) could act as a central Kratom metabolite and branchpoint of Kratom MIA biosynthesis (Figure 6). *Ms*DCR catalyzes a well-precedented 1,2-iminium reduction but is not related to other known MIA-associated iminium reductases, *Cr*THAS1, *Ms*DCS1, *Cp*DCS or *Cr*DPAS (< 15% sequence identity). The most closely related sequences to *Ms*DCR are isoflavone reductase-like enzymes, known to catalyze the reduction of alkenes in isoflavonoid biosynthesis (∼70% sequence identity).^31^ Thus, *Ms*DCR represents a new class of iminium reductase. The relatively broad substrate scope of this oxidase/reductase pair will be useful for the formation of intermediates of other known 3*R*-MIA biosynthetic pathways. Notably, yohimbine-type alkaloids could also be epimerized by these enzymes, even though these pharmacologically important alkaloids are not produced by Kratom. Along with the recently reported yohimbine synthase from *Rauwolfia tetraphylla*,^36^ biocatalytic production of 3*R* yohimbanes, such as pseudoyohimbine (**22a**) and isorauhimbine (**22b**) is now possible.

**Figure 6.**
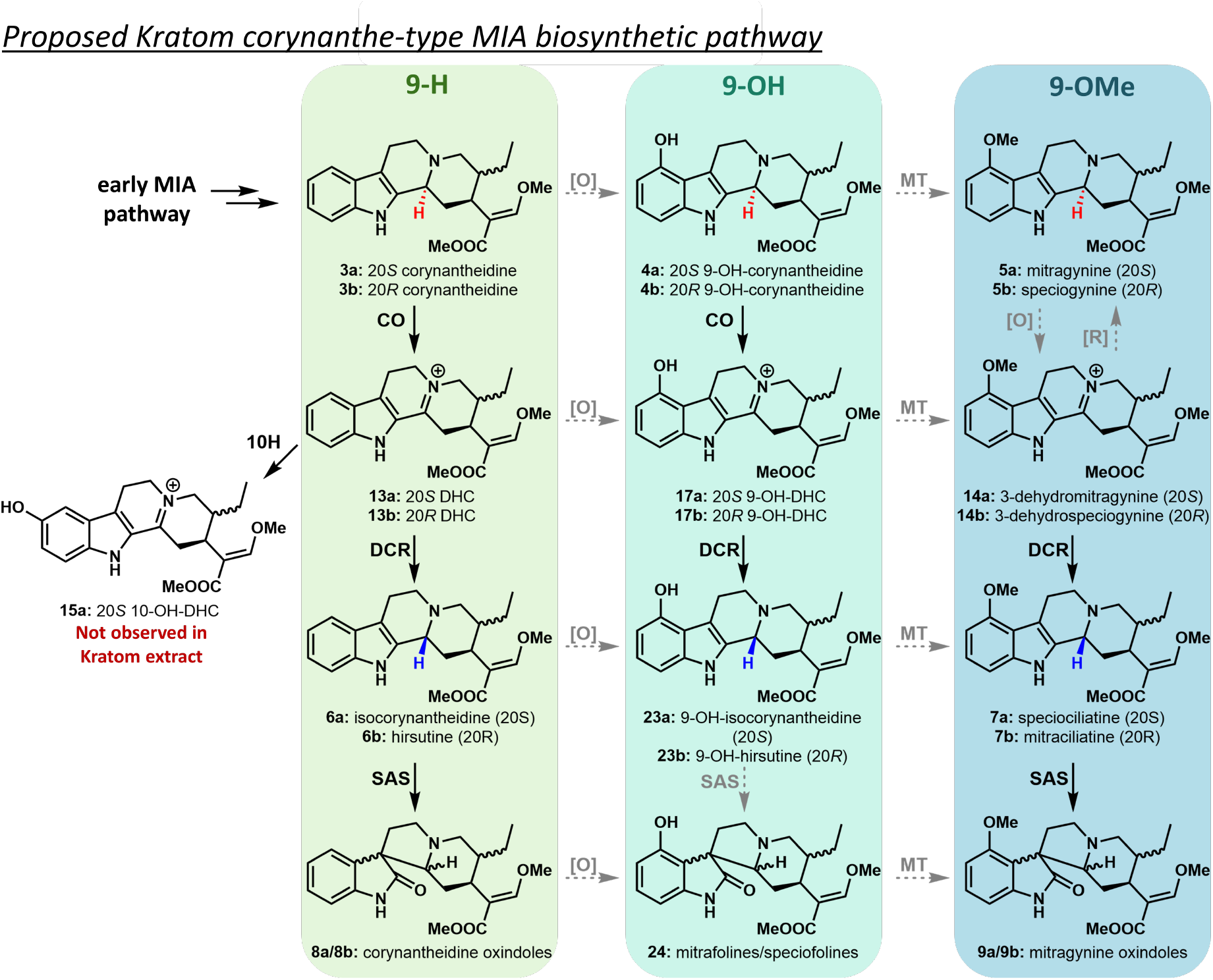
Proposed and updated Kratom corynanthe-type MIA biosynthetic pathway from corynantheidine (**3**). 9-H MIAs are shown highlighted in green, 9-OH MIAs in teal, and 9-OMe MIAs in blue. Unknown transformations are depicted in gray.

Finally, establishment of enzymatic production of numerous 3*R*-MIAs allowed assessment of the activity of the spirocyclase CYP71, *Ms*SAS.^26^ We showed that *Ms*SAS turns over all tested 3*R* corynanthe- and heteroyohimbine-type MIAs but is inactive on yohimbane scaffolds. We reconstituted the biosyntheses of spirooxindoles corynoxine (**8a**) and rhynchophylline (**8b**) from tryptamine. Additionally, both speciociliatine (**7a**) and mitraciliatine (**7b**) are spirocyclized via this enzyme, indicating that presence of the 9-methoxy group does not inhibit catalytic activity. When 3*R*-MIAs with 20*R* stereochemistry (hirsutine (**6b**), hirsuteine (**6c**), and isomayumbine (**11c**)) are used as substrates, two 3*S* spirooxindole alkaloid products are formed (Figure S36). Conversely, substrates with 20*S* stereochemistry (20S-corynantheidine (**3a**) or THA (**10a**)) are converted into a more diverse product mixture comprising both 3*R* and 3*S* geometries (Figure S37). This is consistent with previous studies that have confirmed the effect of ring stereochemistry on product distribution resulting from isomerization.^33,37^ Notably, for both 20*S* and 20*R* geometries, the thermodynamic equilibrium favors 3*S* spirocyclic products, meaning the formation of (3*R*, 20*S*) spirooxindole alkaloids is due to the stereochemical control of *Ms*SAS. Our data, therefore, shows that the *Ms*SAS active site is able to promote formation of otherwise thermodynamically-unfavorable spirocyclic products with 3*R* geometries.

## Conclusion

3*R*-MIAs such as hirsutine (**6b**) and speciociliatine (**7a**) have long been recognized as having valuable bioactivites, but the biosynthetic pathways for these compounds have remained cryptic. Labelled substrate feeding, isolation of intermediates, and bioinformatic analysis of RNAseq data led to the discovery of two enzymes, a BBE-like FAD-linked oxidase, *Ms*CO, and an iminium reductase, *Ms*DCR, that act together to epimerize 3*S*-MIAs to 3*R*-MIAs in *M. speciosa* (Kratom). The resulting 3R-MIAs can also be combined with the previously discovered *Ms*SAS to form spiroxindole alkaloids. The wide substrate specificity of this two-enzyme cascade expands the available stereochemical space of MIAs, thereby substantially improving access to these medicinally relevant compounds.

## Acknowledgments

We acknowledge Dr. Gabriel Titchiner, Dr. Ling Chuang, and Dr. Maite Colinas for insightful feedback/advice. We acknowledge the greenhouse team with special thanks to Eva Rothe. We acknowledge Dr. Maritta Kunert and Sarah Heinicke for their support and assistance with mass spectrometry data collection. We acknowledge Satya Swathi Nadakuduti for procurement of Kratom plants. We acknowledge funding from the Max Planck Society, DfG Leibniz Award (505457618) and the National Institutes of Health (AT012783-02).We acknowledge funding from Michigan State University to C.R.B.

## References

(1) Allain, H.; Bentue-Ferrer, D. Clinical Efficacy of Almitrine-Raubasine. Eur. Neurol. 1998, 39, 39–44.

(2) Roquebert, J.; Demichel, P. INHIBITION OF THE a I-AND A2-ADRENOCEPTOR-MEDIATED PRESSOR RESPONSE IN PITHED RATS BY RAUBASINE, TETRAHYDROALSTONINE AND AKUAMMIGINE. Eur. J. Pharmacol. 1985, 106, 203–205.

(3) Dhyani, P.; Quispe, C.; Sharma, E.; Bahukhandi, A.; Sati, P.; Attri, D. C.; Szopa, A.; Rad, J. S.; Docea, A. O.; Mardare, I.; Calina, D.; Cho, W. C. Anticancer Potential of Alkaloids : A Key Emphasis to Colchicine , Vinblastine , Vincristine , Vindesine , Vinorelbine and Vincamine. Cancer Cell Int. 2022, 1–20. 10.1186/s12935-022-02624-9.

(4) Hong, B.; Grzech, D.; Caputi, L.; Sonawane, P.; López, C. E. R.; Kamileen, M. O.; Hernández Lozada, N. J.; Grabe, V.; O’Connor, S. E. Biosynthesis of Strychnine. Nature 2022, 607 (7919), 617–622. 10.1038/s41586-022-04950-4.

(5) Harun, N.; Kamaruzaman, N. A.; Sofian, Z. M.; Hassan, Z. Potential Therapeutic Values of Mitragynine as an Opioid Substitution Therapy. Neurosci. Lett. 2022, 773 (December 2021), 136500. 10.1016/j.neulet.2022.136500.

(6) Koenig, X.; Hilber, K. The Anti-Addiction Drug Ibogaine and the Heart: A Delicate Relation. Molecules 2015, 20, 2208–2228. 10.3390/molecules20022208.

(7) Mash, D. C. IUPHAR – Invited Review - Ibogaine – A Legacy within the Current Renaissance of Psychedelic Therapy. Pharmacol. Res. 2023, 190 (November 2022), 106620. 10.1016/j.phrs.2022.106620.

(8) Boccia, M.; Grzech, D.; Lopes, A. A.; O’Connor, S. E.; Caputi, L. Directed Biosynthesis of New to Nature Alkaloids in a Heterologous Nicotiana Benthamiana Expression Host. Front. Plant Sci. 2022, 13 (June), 1–8. 10.3389/fpls.2022.919443.

(9) Schotte, C.; Jiang, Y.; Grzech, D.; Dang, T.-T. T.; Laforest, L.; León, F.; Mottinelli, M.; Nadakuduti, S. S.; McCurdy, C. R.; O’Connor, S. E. Directed Biosynthesis of Mitragynine Stereoisomers. JACS 2022, 2022.12.22.521574. 10.1021/jacs.2c13644.

(10) Loris, E. A.; Panjikar, S.; Ruppert, M.; Barleben, L.; Unger, M.; Schübel, H.; Stöckigt, J. Structure-Based Engineering of Strictosidine Synthase: Auxiliary for Alkaloid Libraries. Chem. Biol. 2007, 14 (9), 979–985. 10.1016/j.chembiol.2007.08.009.

(11) Shore, P. A.; Giachetti, A. RESERPINE : BASIC AND CLINICAL PHARMACOLOGY. Handb. Psychopharmacol. 1978, 197–219.

(12) Li, J.; Li, J.; Jiang, H.; Li, M.; Chen, L.; Wang, Y.; Wang, L.; Zhang, N.; Guo, H.; Ma, K. Phytochemistry and Biological Activities of Corynanthe Alkaloids. Phytochemistry 2023, 213 (February), 113786. 10.1016/j.phytochem.2023.113786.

(13) Zhu, K.; Yang, S.; Ma, F.; Gu, X.; Zhu, Y.; Zhu, Y. The Novel Analogue of Hirsutine as an Anti-Hypertension and Vasodilatary Agent Both In Vitro and In Vivo. 2015, 1–14. 10.1371/journal.pone.0119477.

(14) Behnke, M.; Chear, N. J.; Ramanathan, S.; Sharma, A.; Leo, F.; Hiranita, T.; Avery, B. A.; Mcmahon, L. R.; Mccurdy, C. R. Investigation of the Adrenergic and Opioid Binding A Ffi Nities, Metabolic Stability, Plasma Protein Binding Properties, and Functional E Ff Ects of Selected Indole-Based Kratom Alkaloids. J. Med. Chem. 2020, 63, 433–439. 10.1021/acs.jmedchem.9b01465.

(15) Bremer, B.; Eriksson, T. TIME TREE OF RUBIACEAE : PHYLOGENY AND DATING THE FAMILY , SUBFAMILIES , AND TRIBES. Int. J. Plant Sci. 2009, 170 (6), 766–793. 10.1086/599077.

(16) Smith, K. E.; Sharma, A.; Grundmann, O.; Mccurdy, C. R. Kratom Alkaloids: A Blueprint? ACS Chem. Neurosci. 2023, 2022–2024. 10.1021/acschemneuro.2c00704.

(17) Zhou, J.; Zhou, S. Isorhynchophylline: A Plant Alkaloid with Therapeutic Potential for Cardiovascular and Central Nervous System Diseases. Fitoterapia 2012, 83 (4), 617–626. 10.1016/j.fitote.2012.02.010.

(18) Zhang, J.; Geng, C.; Huang, X.; Chen, X.; Ma, Y.; Zhang, X.; Chen, J. Chemical and Biological Comparison of Different Sections of Uncaria Rhynchophylla ( Gou-Teng ). 2017, No. 132. 10.1177/1469066717694044.

(19) Qi, W.; Zhou, C.; Bai, X.; Kano, Y.; Chen, Y.; Yuan, D. Metabolites Identification and Pharmacokinetic Profile of Hirsuteine, a Bioactive Component in Uncaria in Rats by Ultra-High Performance Liquid Chromatography Coupled with Quadrupole Time-of-Flight Mass Spectrometry. J. Sep. Sci. 2022, 45, 4145–4157. 10.1002/jssc.202200452.

(20) van der Meulen, T. H.; van der Kerk, G. J. M. Alkaloids in Pausinystalia Yohimbe (K. Schum.) Ex Pierre Part II. The Isolation of a New Alkaloid. RECUEIL 1964, 83, 148–153.

(21) Phillipson, J. D.; Hemingway, S. R. Indole and Oxindole Alkaloids from Cephalanthus Occidentalis. Phytochem. Reports 1974, 13 (1972), 2621–2622.

(22) Kim, K.; Shahsavarani, M.; Johnathan Oswaldo Garza-García, J.; Carlisle, J. E.; Guo, J.; De Luca, V.; Qu, Y. Biosynthesis of Kratom Opioids. New Phytol. 2023. 10.1111/nph.19162.

(23) Laforest, L. C.; Kuntz, M. A.; Rama, S.; Kanumuri, R.; Mukhopadhyay, S.; Sharma, A.; Connor, S. E. O.; Mccurdy, C. R.; Nadakuduti, S. S. Metabolite and Molecular Characterization of Mitragyna Speciosa Identifies Developmental and Genotypic Effects on Monoterpene Indole and Oxindole Alkaloid Composition. J. Nat. Prod. 2023. 10.1021/acs.jnatprod.3c00092.

(24) Flores-bocanegra, L.; Raja, H. A.; Graf, T. N.; Augustinovic, M.; Wallace, E. D.; Hematian, S.; Kellogg, J. J.; Todd, D. A.; Cech, N. B.; Oberlies, N. H. The Chemistry of Kratom [Mitragyna Speciosa]: Updated Characterization Data and Methods to Elucidate Indole and Oxindole Alkaloids. J. Nat. Prod. 2020, 83, 2165–2177. 10.1021/acs.jnatprod.0c00257.

(25) Panda, S. S.; Girgis, A. S.; Aziz, M. N.; Bekheit, M. S. Spirooxindole: A Versatile Biologically Active Heterocyclic Scaffold. Molecules 2023, 28 (2). 10.3390/molecules28020618.

(26) Nguyen, T. M.; Grzech, D.; Chung, K.; Xia, Z.; Nguyen, T.; Dang, T. T. Discovery of a Cytochrome P450 Enzyme Catalyzing the Formation of Spirooxindole Alkaloid Scaffold. Front. Plant Sci. 2023, No. February, 1–7. 10.3389/fpls.2023.1125158.

(27) Houghton, P. J.; Said, I. M. 3-Dehydromitragynine: An Alkaloid from Mitragyna Speciosa. Phytochemistry 1986, 25, 2910–2912.

(28) Farrow, S. C.; Hagel, J. M.; Beaudoin, G. A. W.; Burns, D. C.; Facchini, P. J. Stereochemical Inversion of (S)-Reticuline by a Cytochrome P450 Fusion in Opium Poppy. Nat. Chem. Biol. 2015, 11 (September), 728–732. 10.1038/nchembio.1879.

(29) Winzer, T.; Kern, M.; King, A. J.; Larson, T. R.; Teodor, R. I.; Donninger, S. L.; Li, Y.; Dowle, A. A.; Cartwright, J.; Bates, R.; Ashford, D.; Thomas, J.; Walker, C.; Bowser, T. A.; Graham, I. A. Morphinan Biosynthesis in Opium Poppy Requires a P450-Oxidoreductase Fusion Protein. Science (80-. ). 2015, 349 (6245), 309–312.

(30) Langley, C.; Tatsis, E.; Hong, B.; Paetz, C.; Stevenson, C. E. M.; Lawson, D. M.; Caputi, L.; O’Connor, S. E. Expansion of the Catalytic Repertoire of Alcohol Dehydrogenases in Plant Metabolism. Angew. Chemie 2022. 10.1002/anie.202210934.

(31) Min, T.; Kasahara, H.; Bedgar, D. L.; Youn, B.; Lawrence, P. K.; Gang, D. R.; Halls, S. C.; Park, H. J.; Hilsenbeck, J. L.; Davin, L. B.; Lewis, N. G.; Kang, C. H. Crystal Structures of Pinoresinol-Lariciresinol and Phenylcoumaran Benzylic Ether Reductases and Their Relationship to Isoflavone Reductases. J. Biol. Chem. 2003, 278 (50), 50714–50723. 10.1074/jbc.M308493200.

(32) Merlini, L.; Mendelli, R.; Masini, G. Gambirine, a New Indole Alkaloid from Uncaria Gambier. Tetrahedron Asymmetry 1967, 16, 1571–1574.

(33) Laus, G.; Brossner, D.; Senn, G.; Wurst, K. Analysis of the Kinetics of Isomerization of Spiro Oxindole Alkaloids. J. Chem. Soc. 1996, 1931–1936.

(34) Trenti, F.; Yamamoto, K.; Hong, B.; Paetz, C.; Nakamura, Y.; Connor, S. E. O. Early and Late Steps of Quinine Biosynthesis. 2021, 1–5. 10.1021/acs.orglett.1c00206.

(35) Caputi, L.; Franke, J.; Farrow, S. C.; Chung, K.; Payne, R. M. E.; Nguyen, T. D.; Dang, T. T. T.; Teto Carqueijeiro, I. S.; Koudounas, K.; De Bernonville, T. D.; Ameyaw, B.; Jones, D. M.; Curcino Vieira, I. J.; Courdavault, V.; O’Connor, S. E. Missing Enzymes in the Biosynthesis of the Anticancer Drug Vinblastine in Madagascar Periwinkle. Science (80-. ). 2018, 360 (6394), 1235–1239. 10.1126/science.aat4100.

(36) Stander, E. A.; Lehka, B.; Carqueijeiro, I.; Cuello, C.; Hansson, F. G.; Jansen, H. J.; Bernonville, T. D. De; Williams, C. B.; Vergès, V.; Lezin, E.; Daniel, M.; Bohn, B.; Dang, T.; Oudin, A.; Lanoue, A.; Durand, M.; Giglioli-guivarc, N.; Janfelt, C.; Papon, N.; Dirks, R. P. O S. E.; Jensen, M. K.; Besseau, S.; Courdavault, V. The Rauvolfia Tetraphylla Genome Suggests Multiple Distinct Biosynthetic Routes for Yohimbane Monoterpene Indole Alkaloids. Commun. Biol. 2023, 6, 1997. 10.1038/s42003-023-05574-8.

(37) Wang, X.; Zhang, M.; Liu, X.; Lou, M.; Li, G.; Qi, X. Total Synthesis of Tetracyclic Spirooxindole Alkaloids via a Double Oxidative Rearrangement/Cyclization Cascade. Org. L 2024, 26, 824–828. 10.1021/acs.orglett.3c03938.

