## Supplementary material for "Enzymatic epimerization of monoterpene indole alkaloids in Kratom": SI Data

**Contents:**

**Supplementary Figures** S2– S36

**Supplementary Tables** S36– S38

**Supplementary Materials and Methods** S39– S48

**Coding Sequences** S49– S56

**Synthetic Methods** S57– S61

**Structural Data** S62– S94

**Supplementary References** S95– S96

### Supplementary Figures

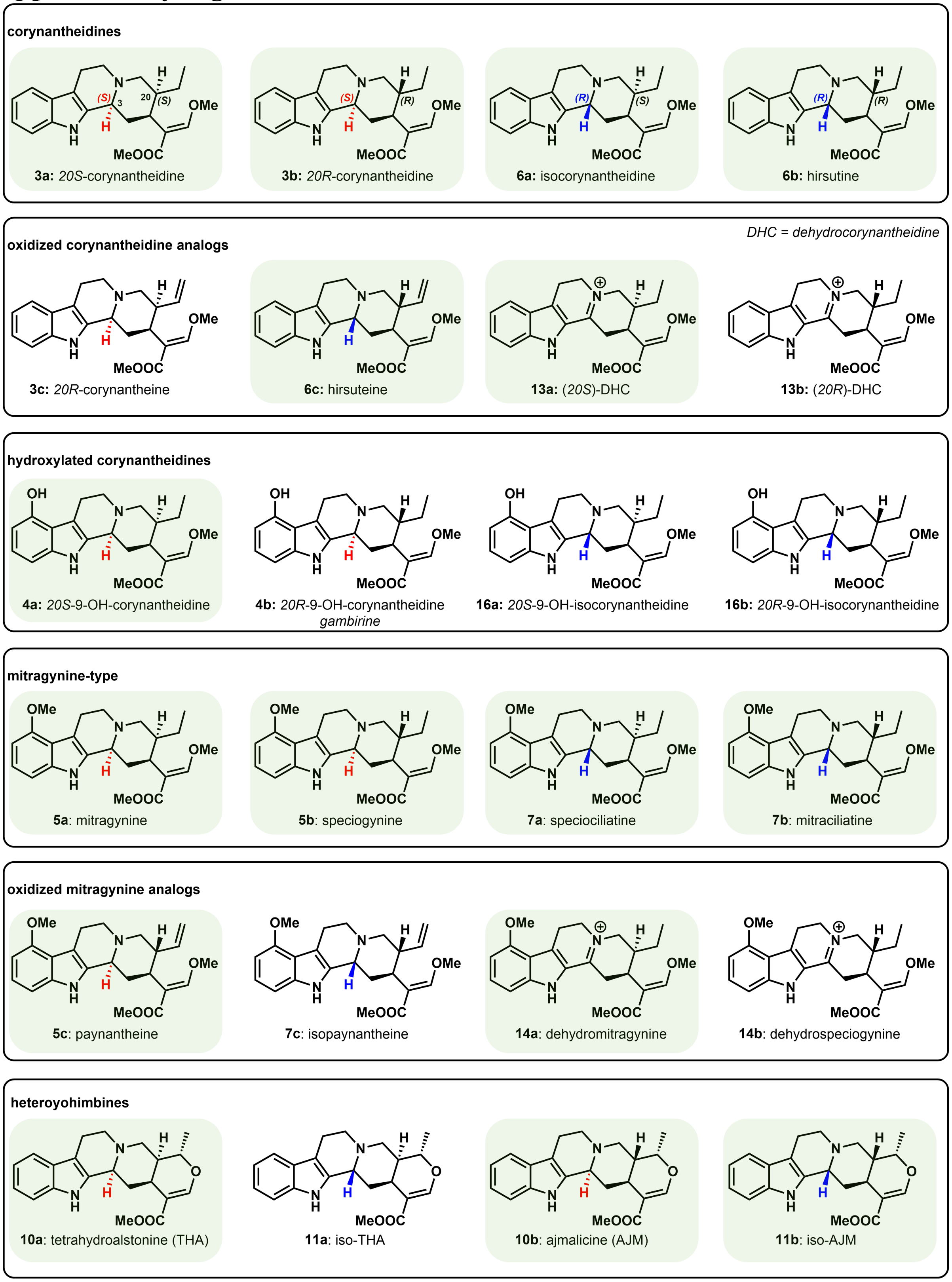


**Figure S1:** Structures and names of select Kratom alkaloids. Compounds with verified standards are highlighted in green.

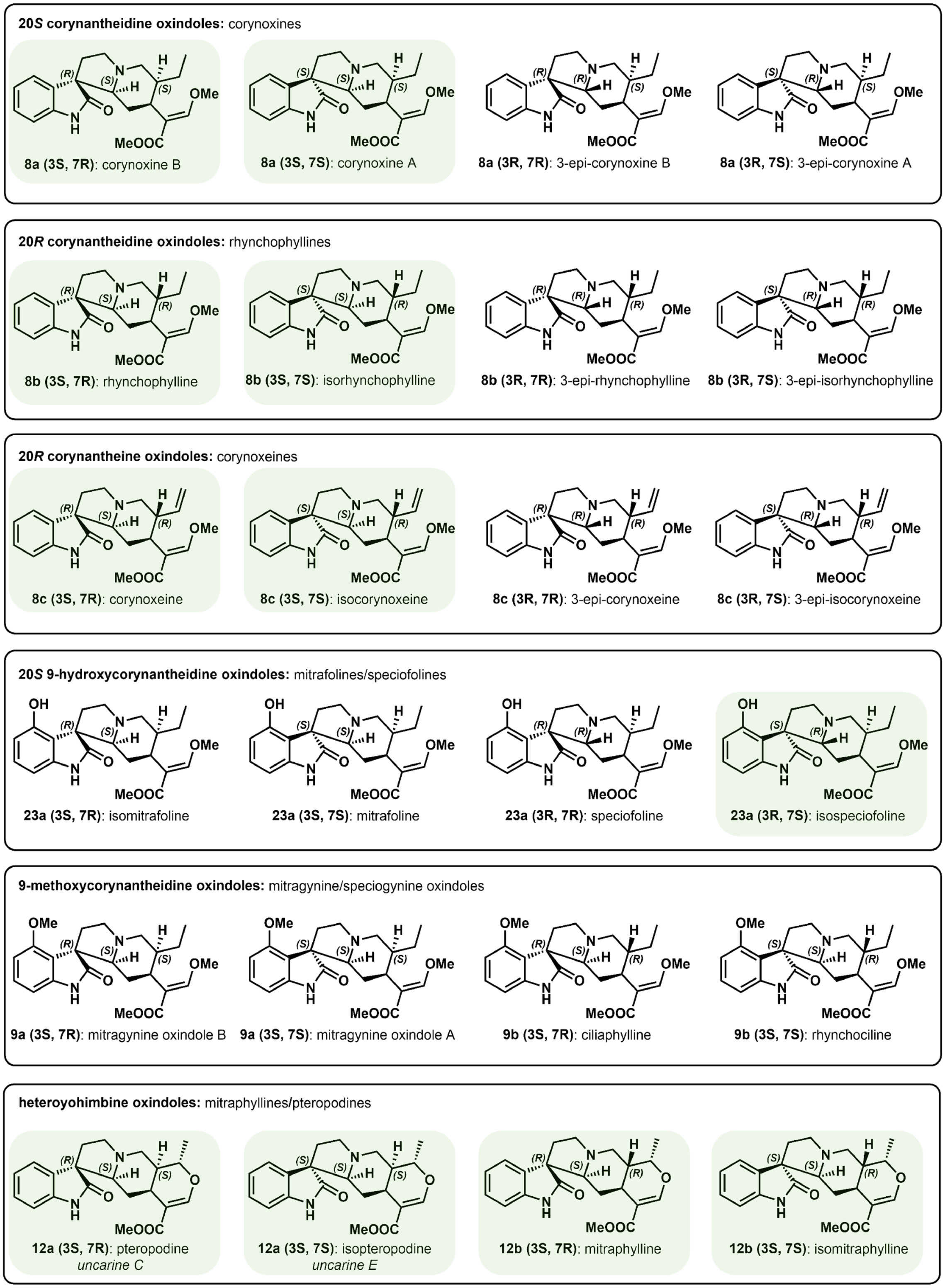


**Figure S2:** Structures and names of select Kratom spirooxindole alkaloids. Compounds with verified standards are highlighted in green. **Note:** different spirocylized isomers (*i.e.* **8a**) retain the same compound numbering but are differentiated by stereochemistry at C-3 and C-7. We adopted this numbering scheme since these isomers interconvert readily.

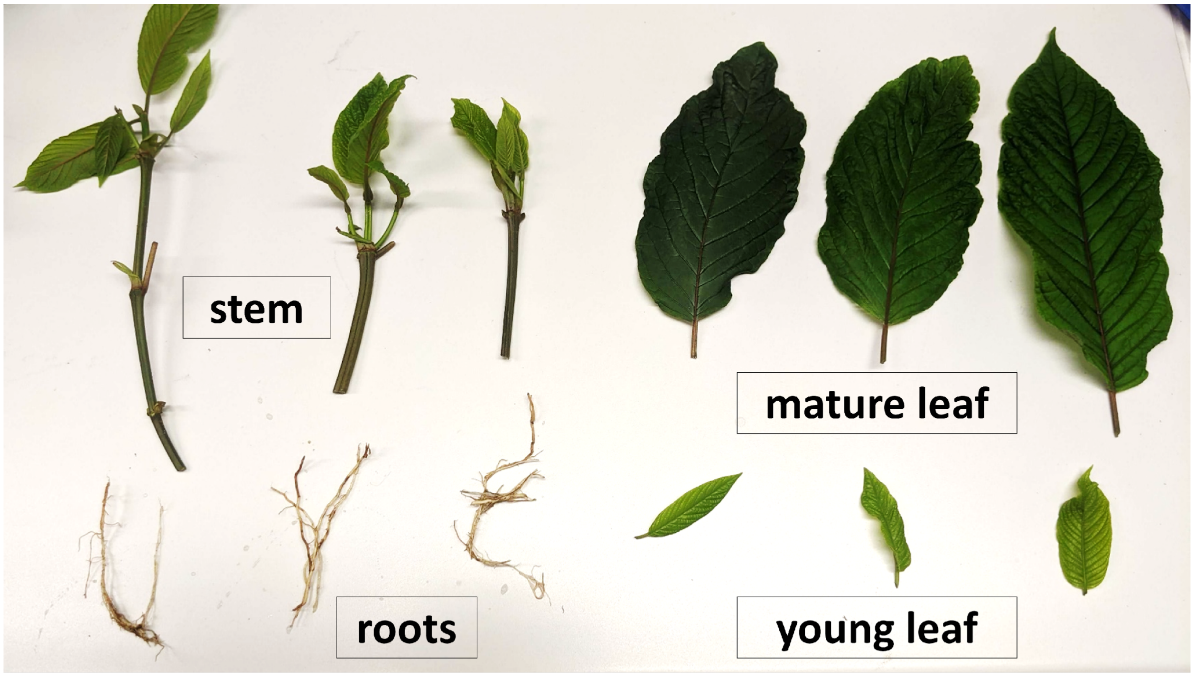


**Figure S3:** Kratom tissues used for alkaloid analysis and feeding studies. Note that for stems, cut stem disks were acquired from the stem internode preceding young leaf petioles. Roots were cut into small 2 mm fragments, while 5 mm leaf disks were cut out from leaf tissue.

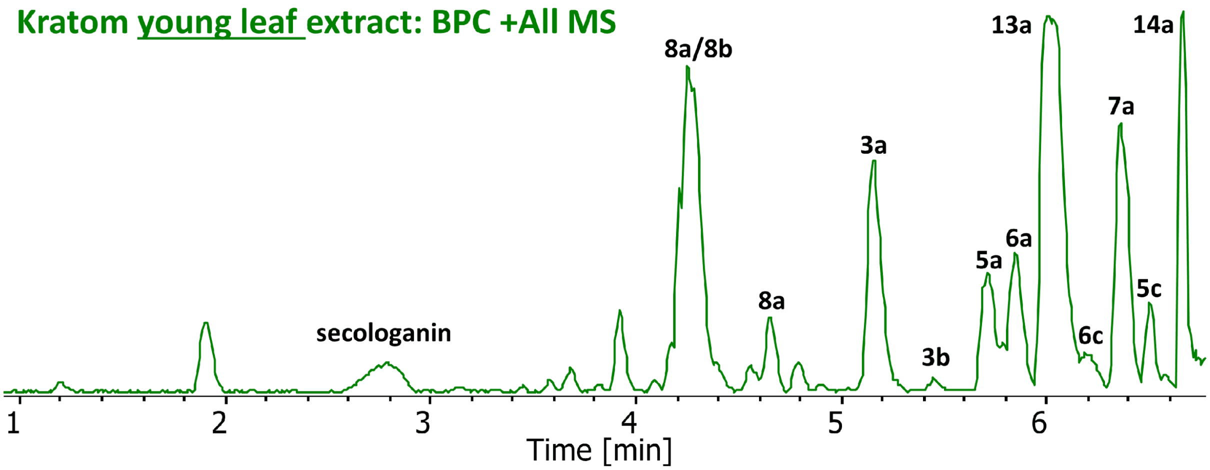


**Figure S4:** Kratom young leaf base peak chromatogram (BPC) with identified alkaloids. Designation of **8a/8b** represents mixtures of corynoxine (**8a**)/rhynchophylline (**8b**) isomers.

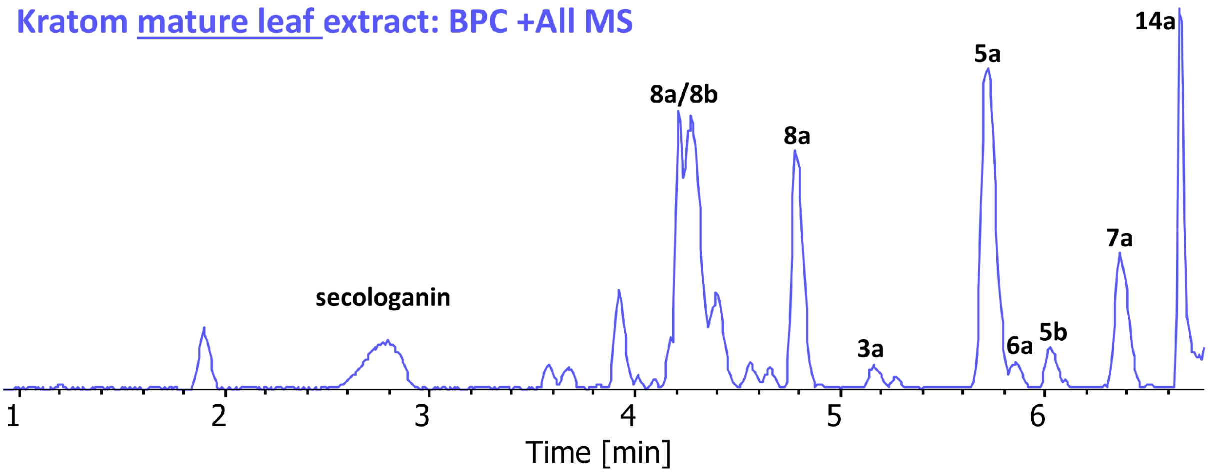


**Figure S5:** Kratom stem base peak chromatogram (BPC) with identified alkaloids. Designation of **8a/8b** represents mixtures of corynoxine (**8a**)/rhynchophylline (**8b**) isomers.

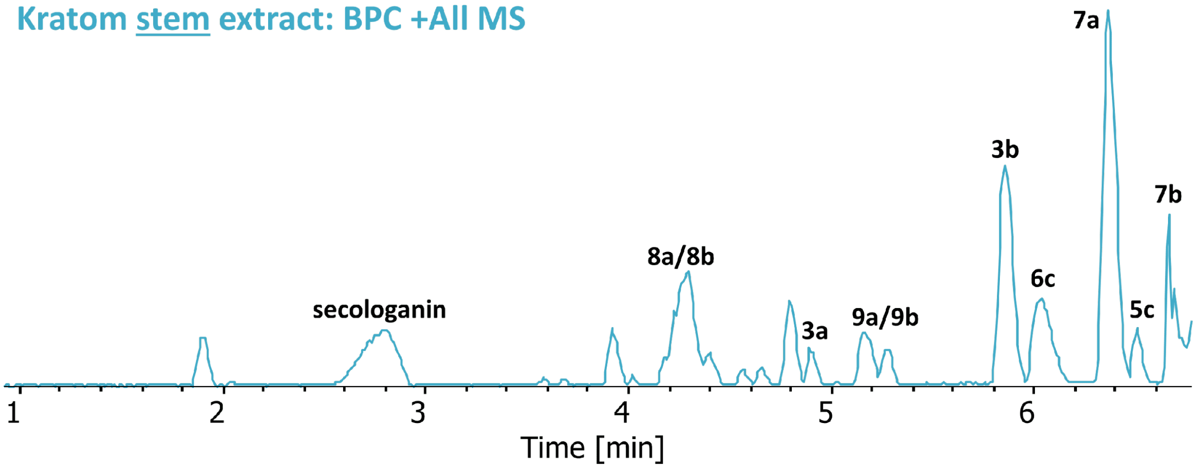


**Figure S6:** Kratom ML base peak chromatogram (BPC) with identified alkaloids. Designation of **8a/8b** represents mixtures of corynoxine (**8a**)/rhynchophylline (**8b**) isomers, and designation of **9a/9b** represents unknown peaks corresponding to mitragynine oxindole (**9a**)/speciogynine oxindole (**9b**) isomers.

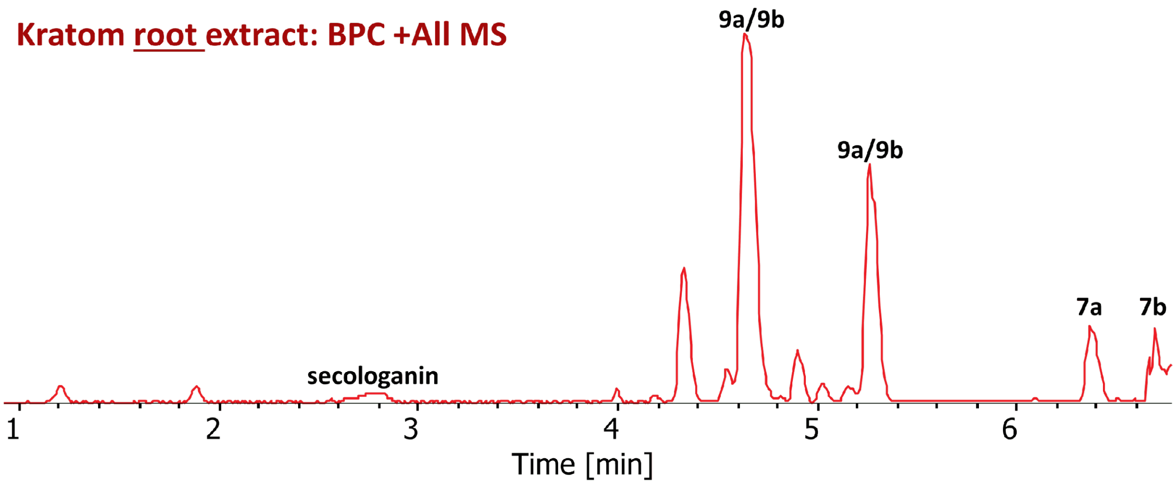


**Figure S7:** Kratom root base peak chromatogram (BPC) with identified alkaloids. Designation of **9a/9b** represents unknown peaks corresponding to mitragynine oxindole (**9a**)/speciogynine oxindole (**9b**) isomers.

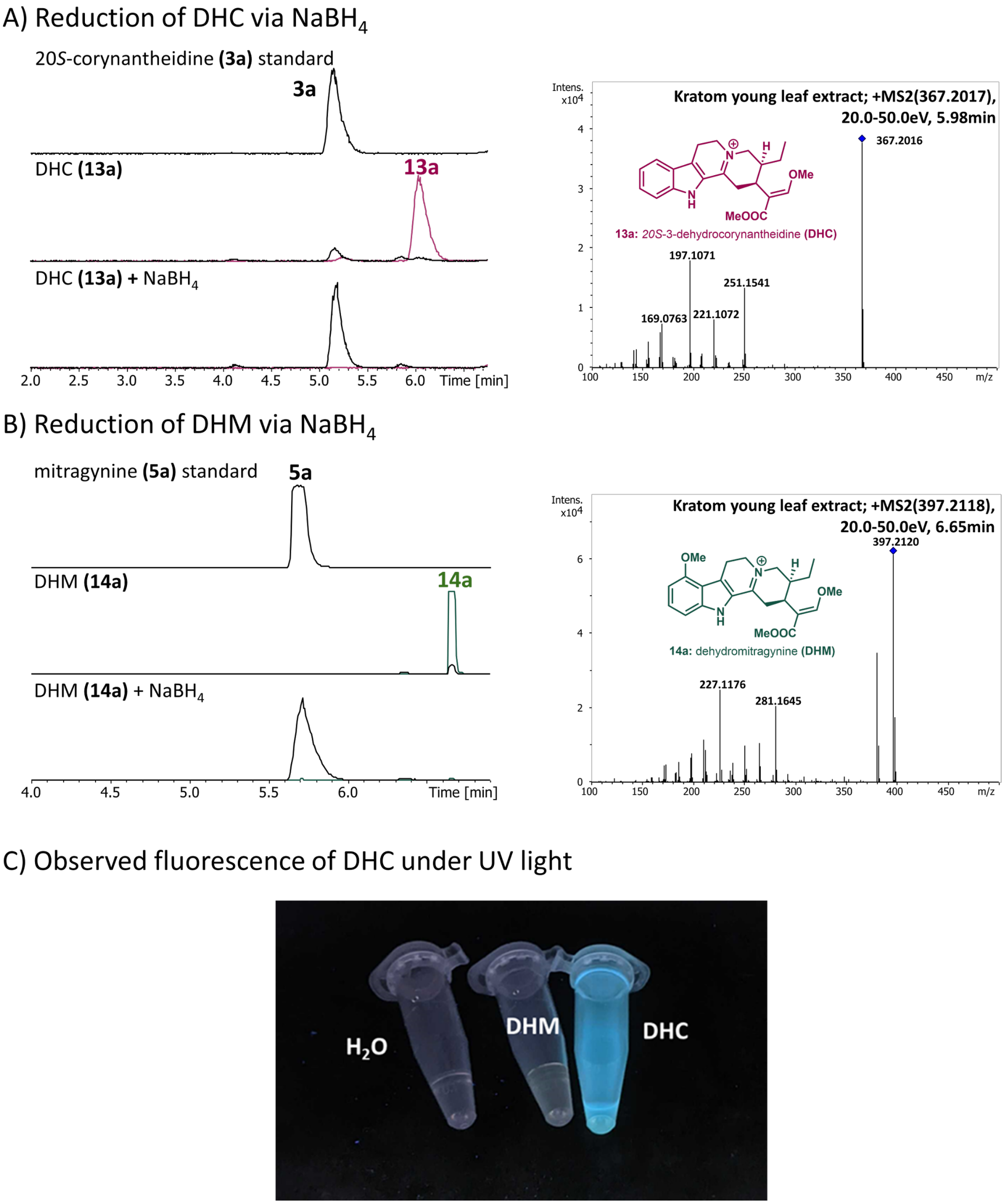


**Figure S8:** A) Reduction of 100 µM isolated DHC with 10 mM $NaBH_4$ at r.t. for 1 h. Full conversion to 20*S*-corynantheidine is observed with no observable formation of other products. MS/MS analysis of isolated DHC. NMR structural characterization of DHC is detailed in Figures S57-S61. B) Reduction of 500 µM DHM with 10 mM $NaBH_4$ at r.t. for 1 h. Full conversion to mitragynine is observed with no observable formation of other products. MS/MS analysis of isolated DHM. C) Observed fluorescence of DHC under UV light. NMR structural characterization of DHM is detailed in Figure S56.

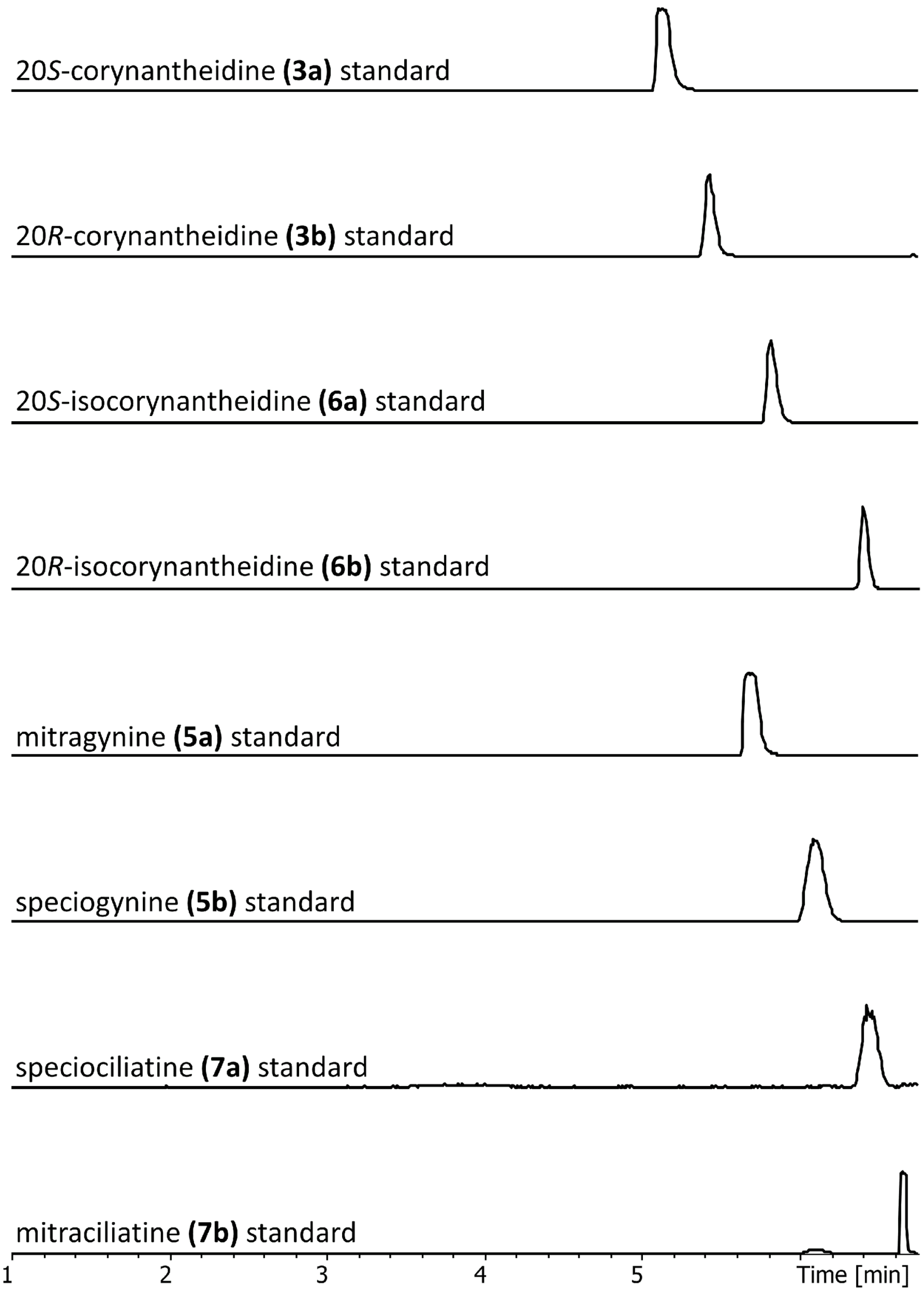


**Figure S9:** Retention times and elution order of corynantheidine and mitragynine stereoisomers with y-axis denoting ionization intensity. We observed a consistent elution order that corresponded to relative stereochemistry at C3 and C20 for both methoxylated and non-methoxylated corynantheidines. In this way, relative retention times assisted in identification of corynantheidine/mitragynine stereoisomers. NMR structural characterization for each isomer is detailed in Figures S48-S55.

A) Young leaf $d_5$-tryptamine feeding results in corynantheidine incorporation

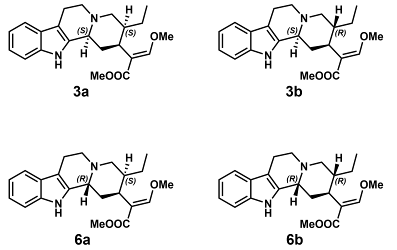


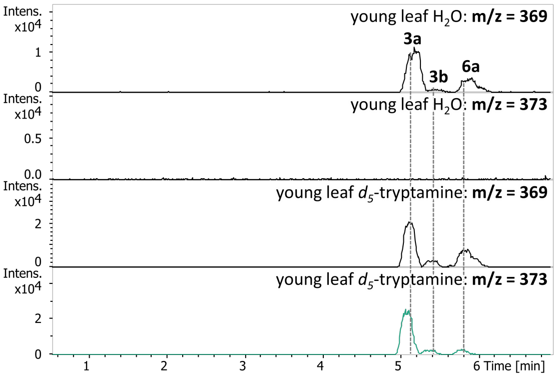


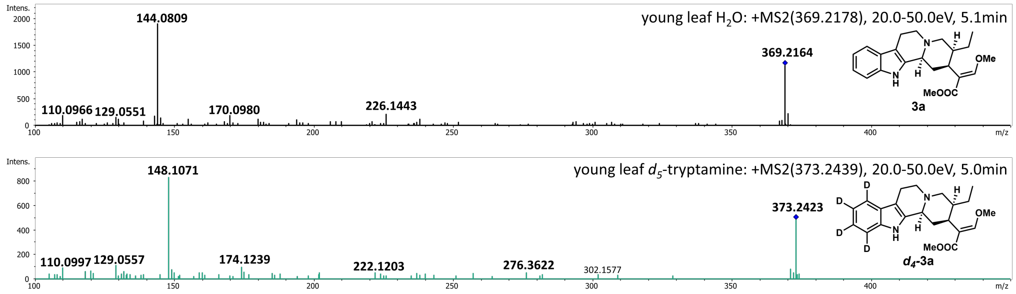


B) Young leaf $d_5$-tryptamine feeding results in mitragynine incorporation

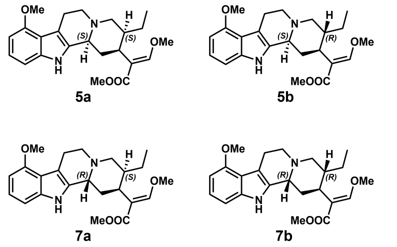


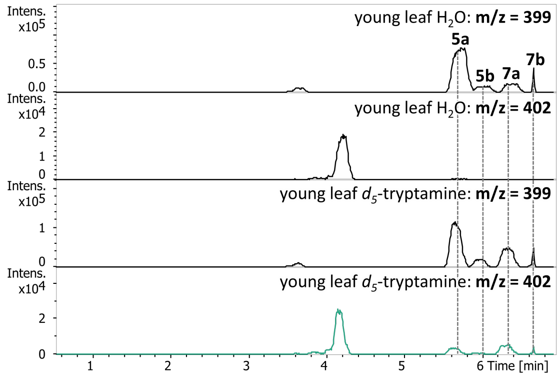


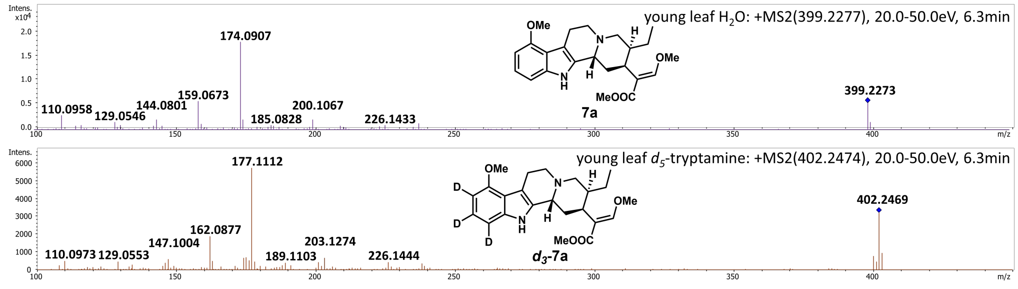


**Figure S10:** Feeding of $d_5$-tryptamine to young leaf tissue. All products were determined via comparison with authentic standards as detailed in Figure S9. A) LC-MS traces of control ($H_2O$) *m/z*+0 and *m/z*+4 and of fed (*d*5-tryptamine) *m/z*+0 and *m/z*+4. Below is the MS2 analysis showing a *m/z*+4 shift relative to 20*S*-corynantheidine (**3a**). B) LC-MS traces of control ($H_2O$) *m/z*+0 and *m/z*+3 and of fed (*d*5-tryptamine) *m/z*+0 and *m/z*+3. Below is the MS2 analysis showing a *m/z*+3 shift relative to speciociliatine.

### Kratom tissue feeding results for $d_4$-strictosidine/$d_4$-vincoside

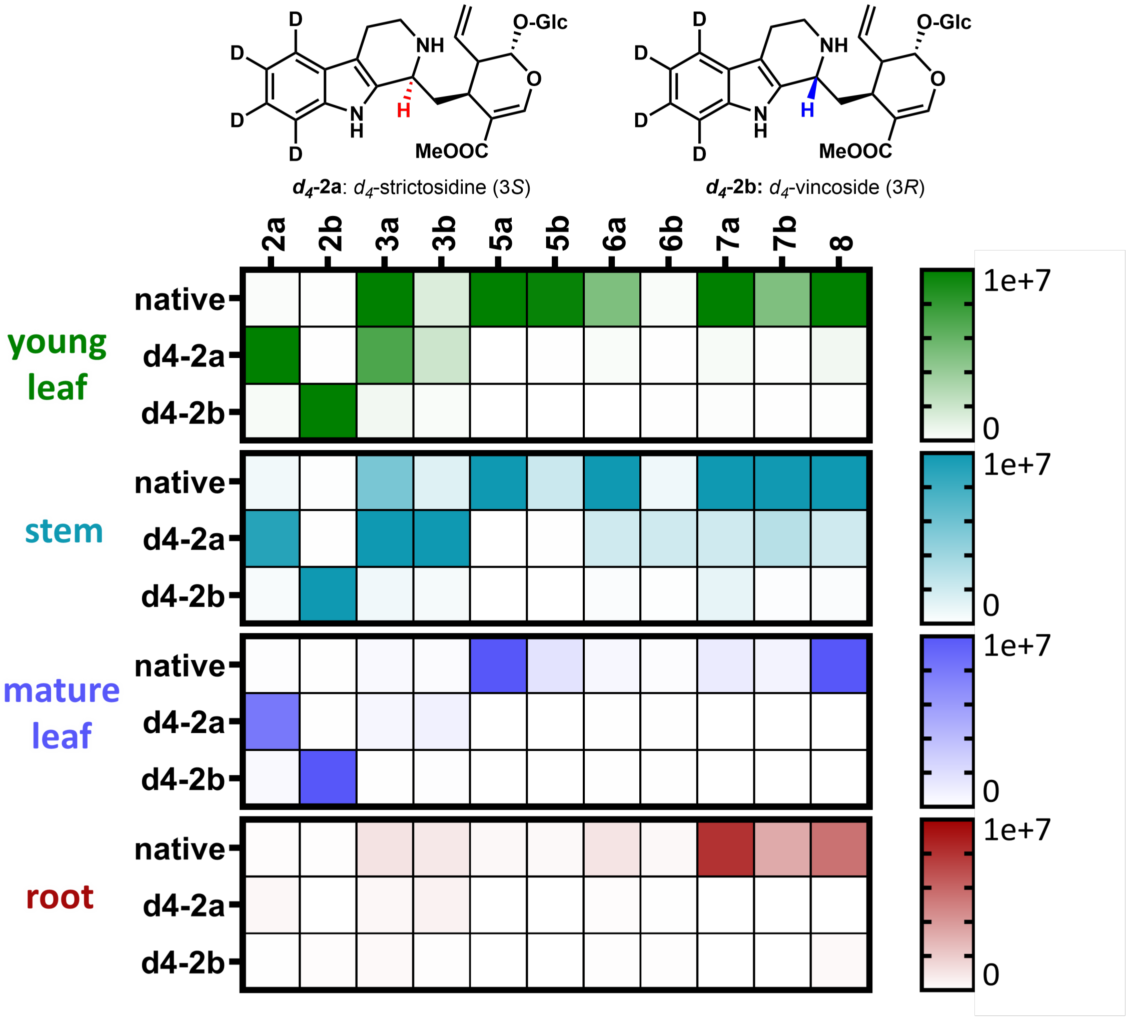


**Figure S11:** Native metabolite distribution of Kratom tissues and incorporation from labeled $d_4$-strictosidine (***d*₄-2a**) and $d_4$-vincoside (***d*₄-2b**).

A) Screening results from BBE candidates

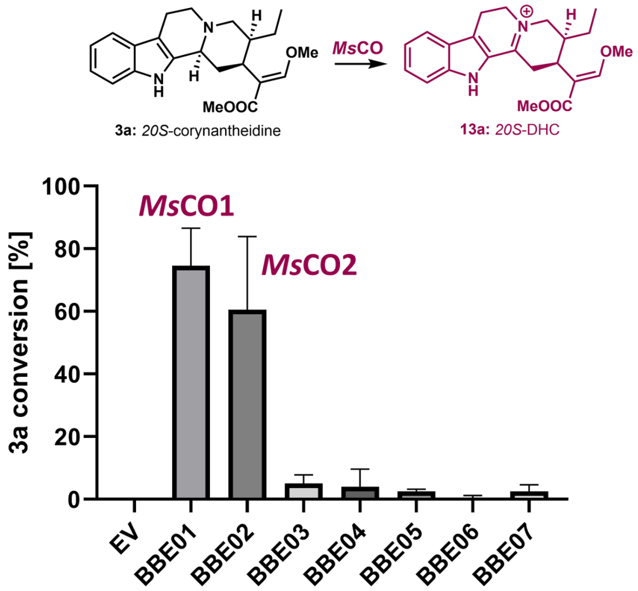


B) Identity matrix of BBEs and *Cr*PAS

| | *Ms*CO1 | *Ms*CO2 | BBE03 | BBE04 | BBE05 | BBE06 | BBE07 | CrPAS |
|---|---|---|---|---|---|---|---|---|
| *Ms*CO1 | | 93.7 | 81.6 | 46.5 | 46.4 | 42.2 | 42.4 | 53.4 |
| *Ms*CO2 | 93.7 | | 81.3 | 47.3 | 46.9 | 42.9 | 43.2 | 53.5 |
| BBE03 | 81.6 | 81.3 | | 47.7 | 45.9 | 42.6 | 42.0 | 55.7 |
| BBE04 | 46.5 | 47.3 | 47.7 | | 58.6 | 44.9 | 44.1 | 42.3 |
| BBE05 | 46.4 | 46.9 | 45.9 | 58.6 | | 42.0 | 42.5 | 42.3 |
| BBE06 | 42.2 | 42.9 | 42.6 | 44.9 | 42.0 | | 73.9 | 39.9 |
| BBE07 | 42.4 | 43.2 | 42.0 | 44.1 | 42.5 | 73.9 | | 42.1 |
| CrPAS | 53.4 | 53.5 | 55.7 | 42.3 | 42.3 | 39.9 | 42.1 | |

C) Identity matrix of *Ms*DCR

| | *Ms*DCR | *Ca*IFR | *Ms*DCS1 | *Cr*THAS1 | *Cr*DPAS |
|---|---|---|---|---|---|
| *Ms*DCR | | 73.3 | 8.5 | 6.4 | 14.3 |
| *Ca*IFR | 73.3 | | 17.0 | 12.8 | 16.3 |
| *Ms*DCS1 | 8.5 | 17.0 | | 62.5 | 56.5 |
| *Cr*THAS1 | 6.4 | 12.8 | 62.5 | | 52.6 |
| *Cr*DPAS | 14.3 | 16.3 | 56.5 | 52.6 | |

**Figure S12:** Newly discovered Kratom epimerase enzymes. A) Activity of berberine bridge-like enzyme (BBE) candidates on corynantheidine (**3a**). Assay conditions: 16 h 100 µM **3a** incubation into *N. benthamiana* leaves infiltrated with the indicated gene. Background activity from EV control is subtracted from final conversion values. B) Sequence alignment matrix of BBE candidates with each other and *Catharanthus roseus* precondylocarpine acetate synthase (*Cr*PAS). C). Sequence alignment matrix of *Ms*DCR, *Coffea arabica* isoflavone reductase (*Ca*IFR), and reported MIA iminium dehydrogenases: *Ms*DCS1, *Cr*THAS1, and *Cr*DPAS.

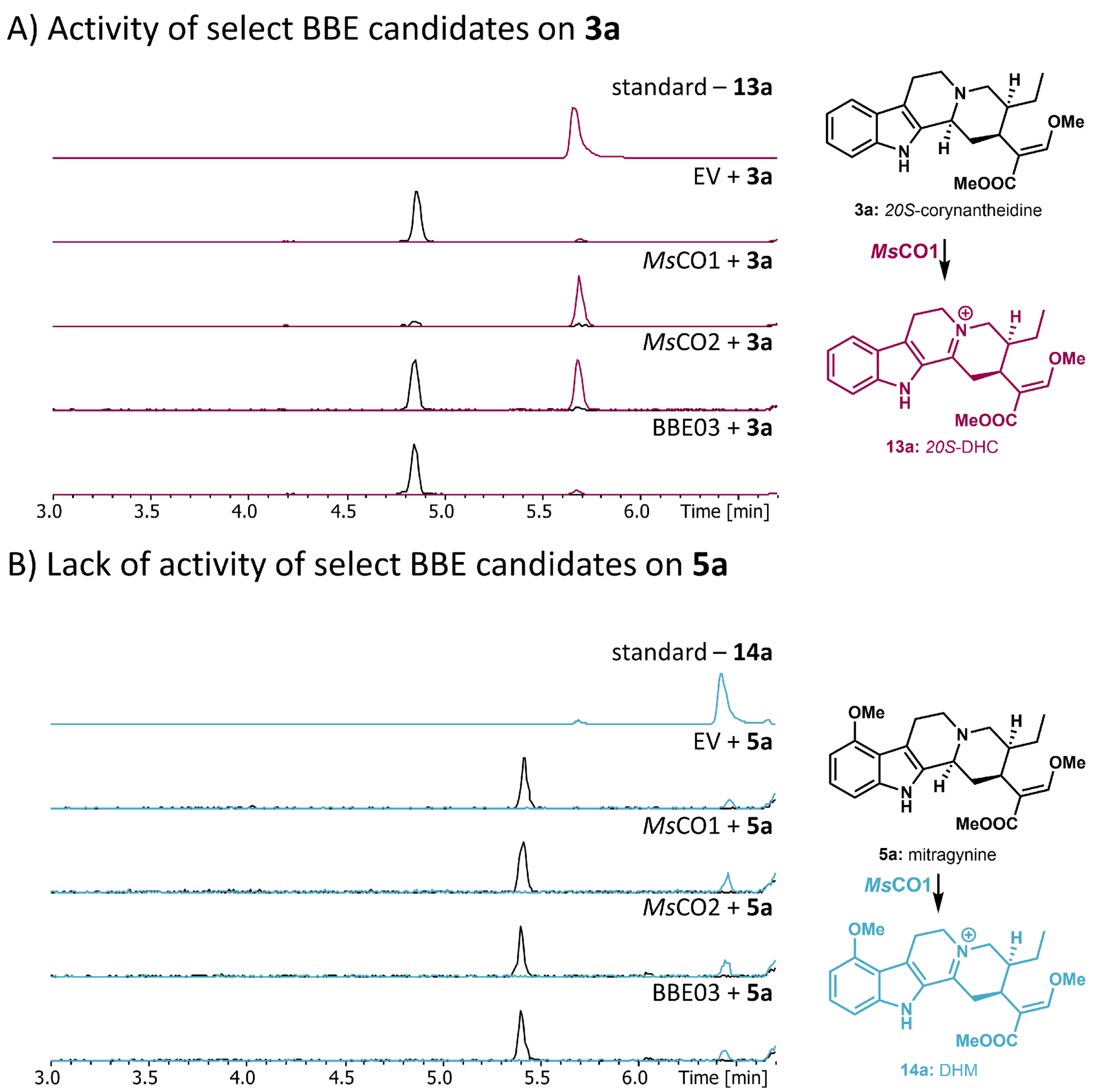


**Figure S13:** Activity differences of *Ms*CO1 and *Ms*CO2 with A) **3a** or B) **5a**. BBE03 is included as an example of a candidate with no activity on **3a**. Assays were performed as described in Figure S12.

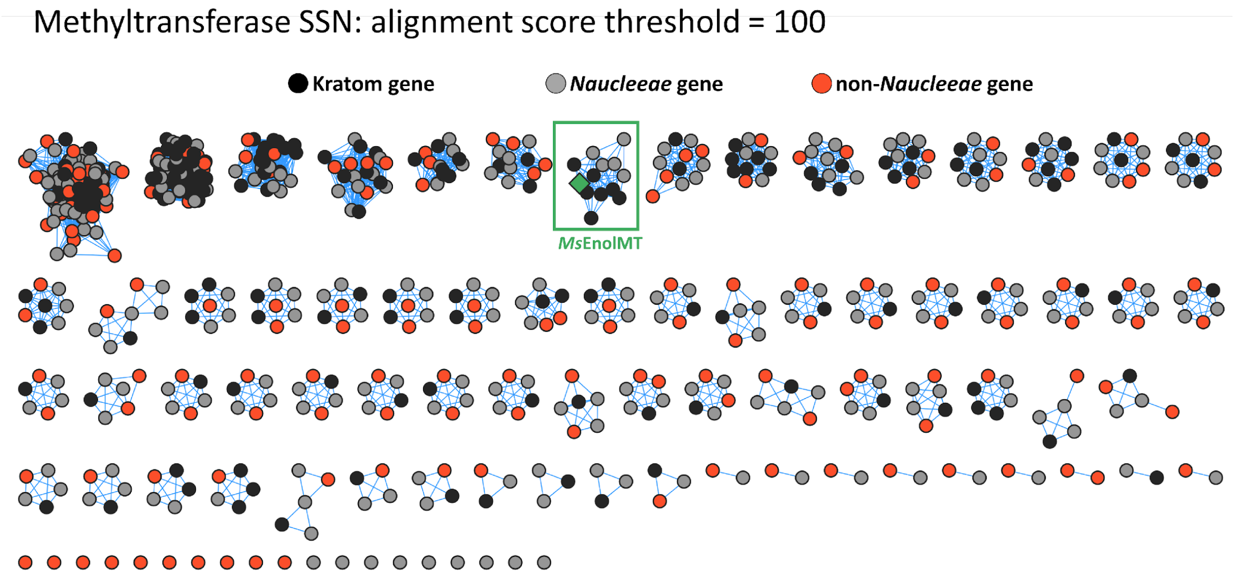


**Figure S14:** Sequence similarity network (SSN) to identify *Naucleeae*-specific methyltransferases with alignment score threshold of 100. Kratom genes are shown in black, *Naucleeae* genes in gray, and non-*Naucleeae* genes in orange. The *Naucleeae*-specific cluster containing the previously identified *Ms*EnolMT is highlighted in green.

Reductase SSN: alignment score threshold = 145

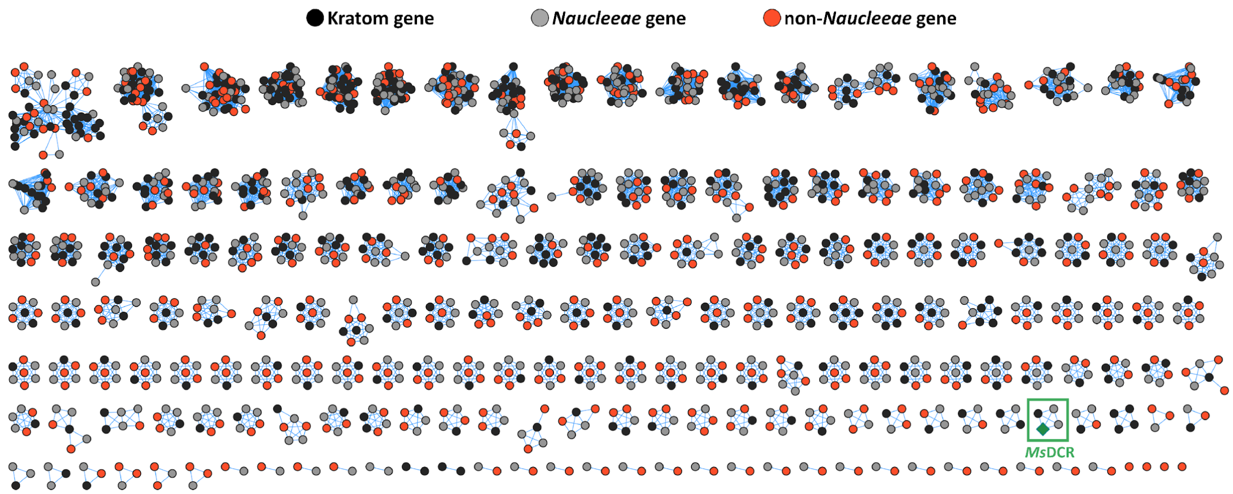


**Figure S15:** Sequence similarity network (SSN) to identify *Naucleeae*-specific reductases/dehydrogenases with alignment score threshold of 150. Kratom genes are shown in black, *Naucleeae* genes in gray, and non-*Naucleeae* genes in orange. The *Naucleeae*-specific cluster containing *Ms*DCR is highlighted with a box (green).

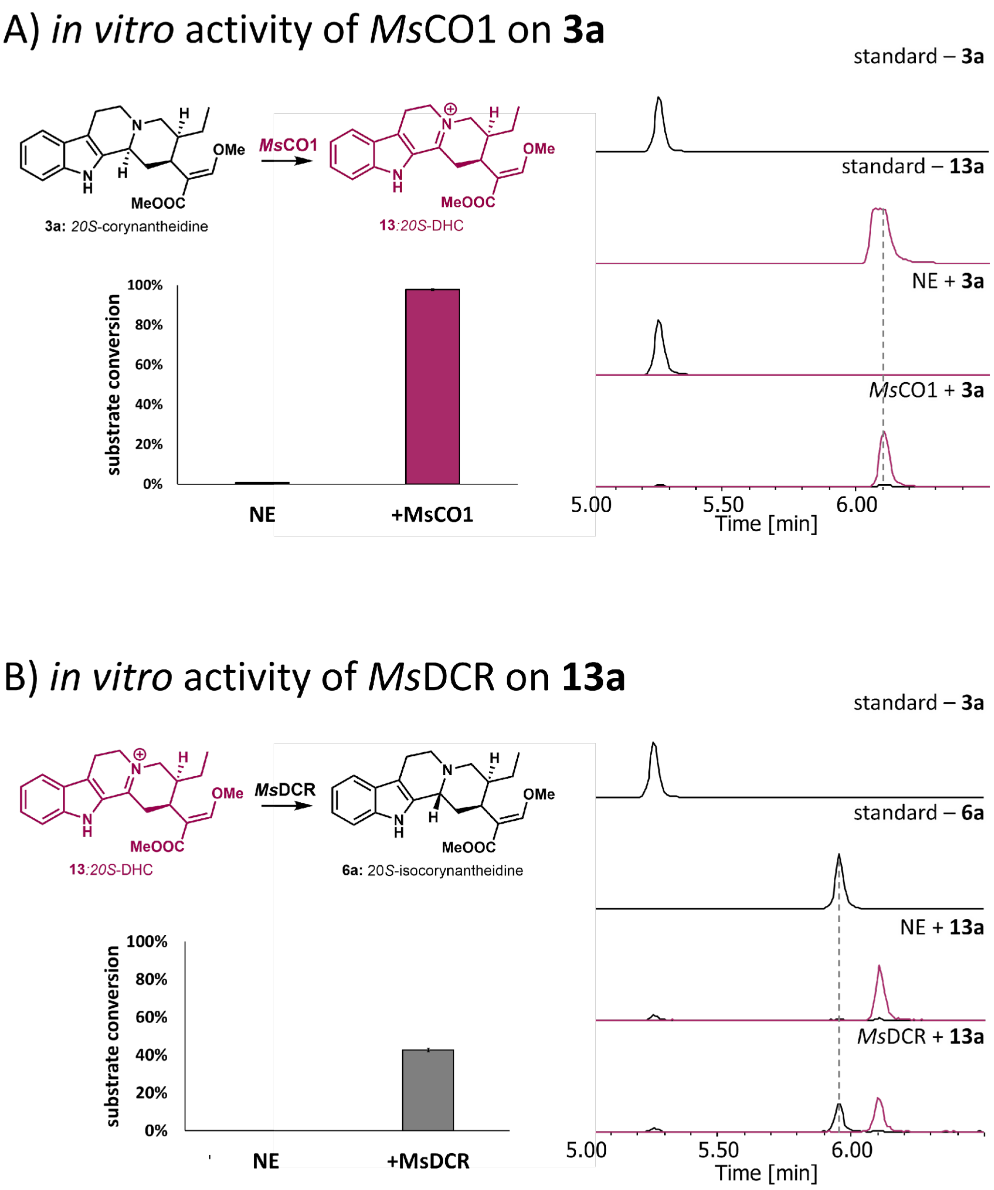


**Figure S16:** *in vitro* verification of *Ms*CO1 and *Ms*DCR activity. A) Reaction of *Ms*CO1 converting **3a** to **13a**. NE = no enzyme control. Reaction conditions: 100 µM **3a**, 0.5 µM *Ms*CO1/EV control, 50 mM Tris-HCl pH = 7.5, 25 ˚C for 16 h. B) Reaction of *Ms*DCR converting **13a** to **6a**. NE = no enzyme control. Reaction conditions: 100 µM **3a**, 1 mM NADPH 5 µM *Ms*DCR, 50 mM Tris-HCl pH = 7.5, 25 ˚C for 16 h.

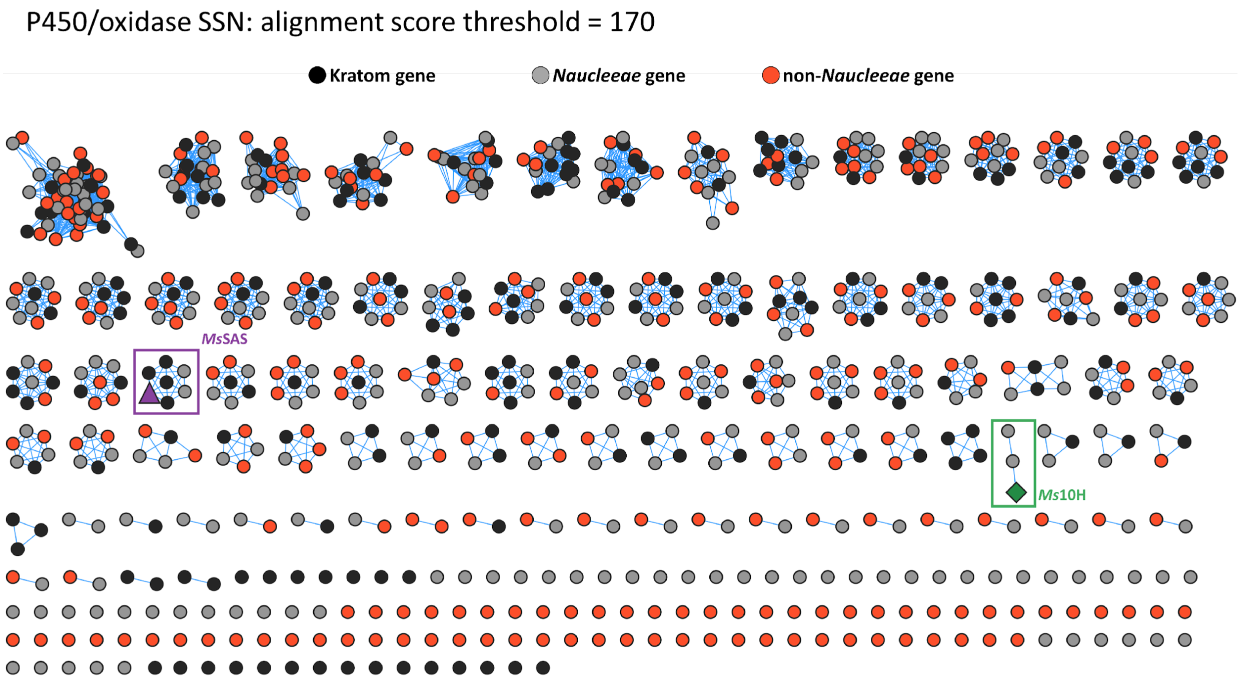


**Figure S17:** Sequence similarity network (SSN) to identify *Naucleeae*-specific P450s/oxidases with alignment score threshold of 170. Kratom genes are shown in black, *Naucleeae* genes in gray, and non-*Naucleeae* genes in orange. The *Naucleeae*-specific cluster containing *Ms*10H is highlighted with a green box and the Naucleeae-specific cluster containing *Ms*SAS is highlighted with a purple box.

#### Discovery of *Ms*10H activity on DHC (**13a**)

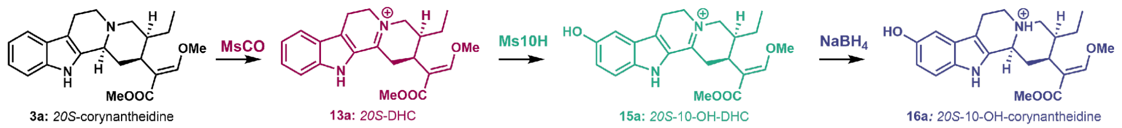


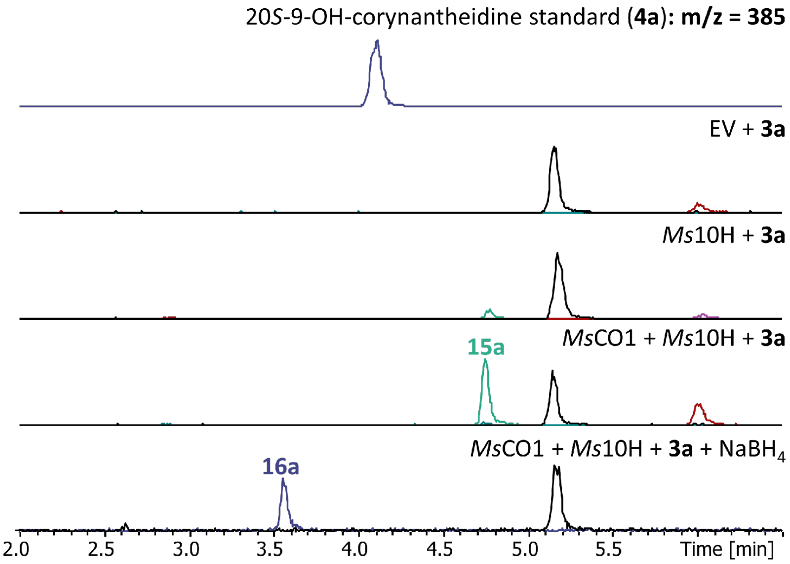


**Figure S18:** Activity of *Ms*10H. LC-MS traces showing *Ms*10H is only active on DHC and is reduced via $NaBH_4$ to a compound that does not match the standard for 20*S*-9-OH-corynantheidine (**4a**).

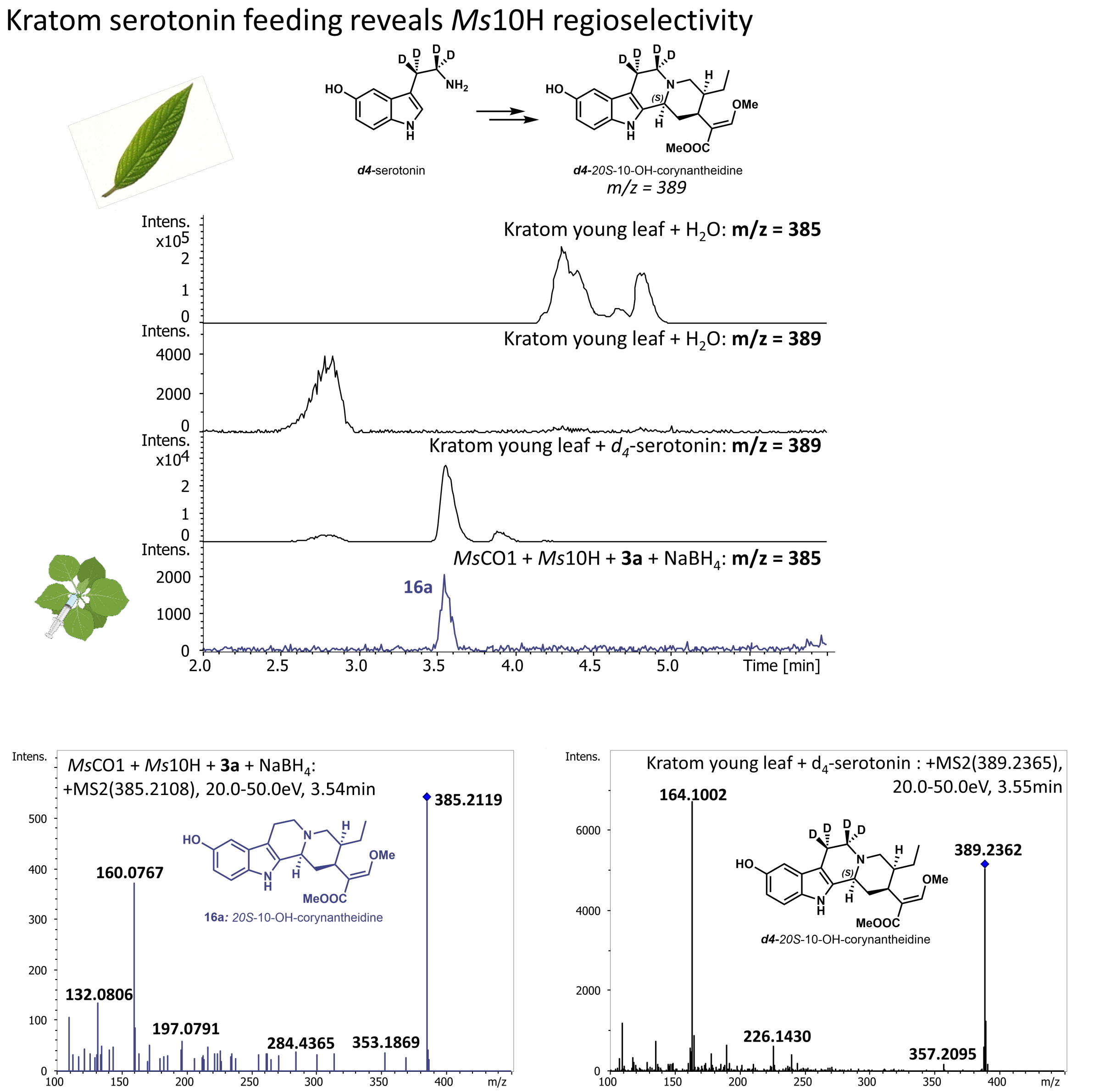


**Figure S19:** Production of 10-OH-corynantheidine standard by feeding isotopically labelled serotonin to Kratom tissue. A) Feeding of 1 mM $d_4$-serotonin into Kratom young leaf tissue produced the corresponding 10-OH-corynantheidine, which co-eluted with the product from *Ms*10H following $NaBH_4$ reduction. Below shows the comparison between the MS2 of the *Ms*10H-derived compound and the produced labelled 10-OH-corynantheidine.

Overview of enzymatically produced spirooxindole compounds

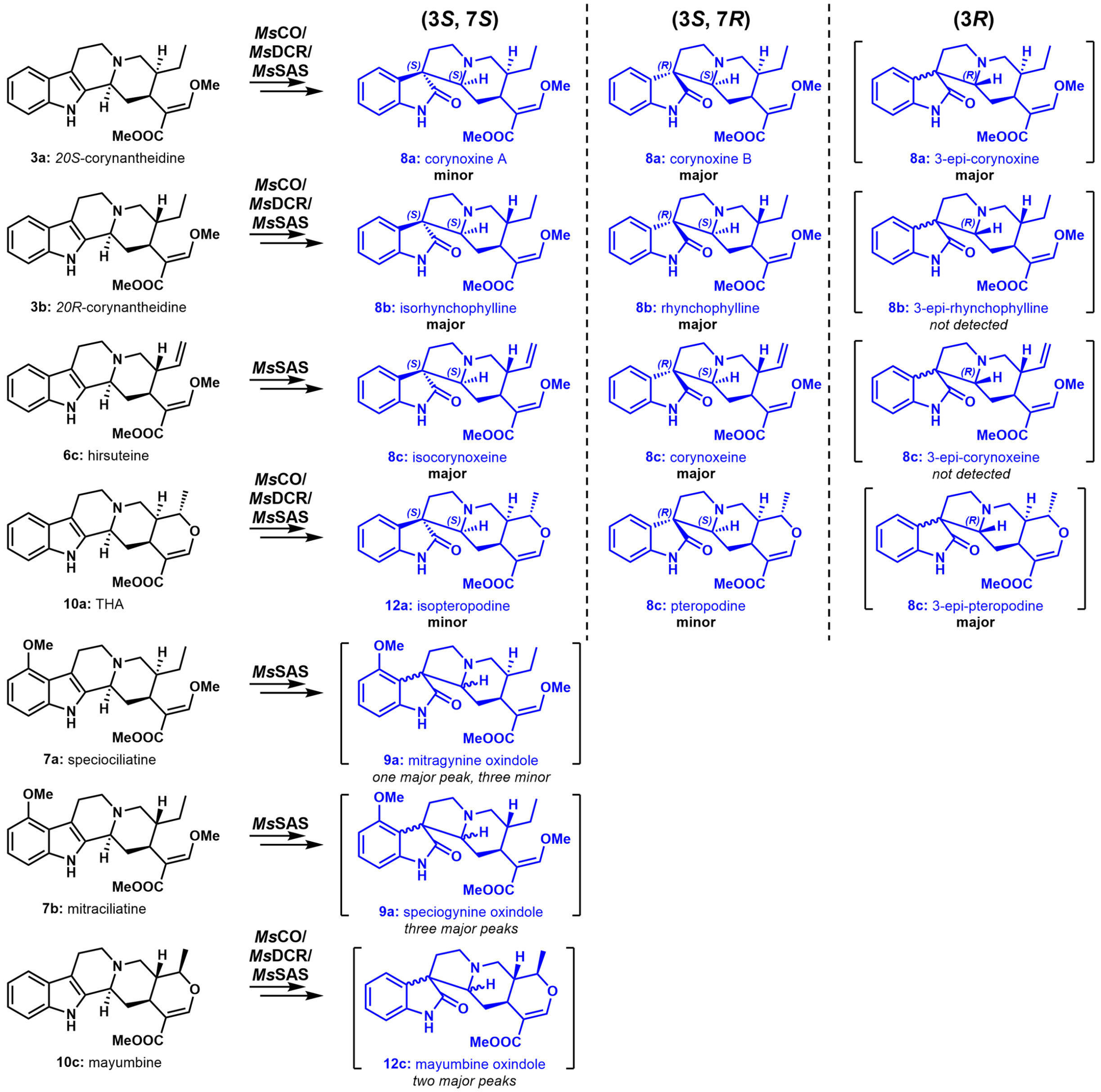


**Figure S20:** Summative figure of substrate scope results (Figures S19-S31) with observed spirooxindole formation via *Ms*SAS and available standards. Compounds in brackets represent putatively identified compounds based on mass fragmentation and chemical reasoning. No standards for 3*R* spirooxindole compounds were available, identification between 3*R* isomers was not possible. **Major**/**minor**/*not detected* modifiers indicate relative peak abundance observed via LC-MS.

#### Activity of *Ms*SAS on hirsuteine (**6c**)

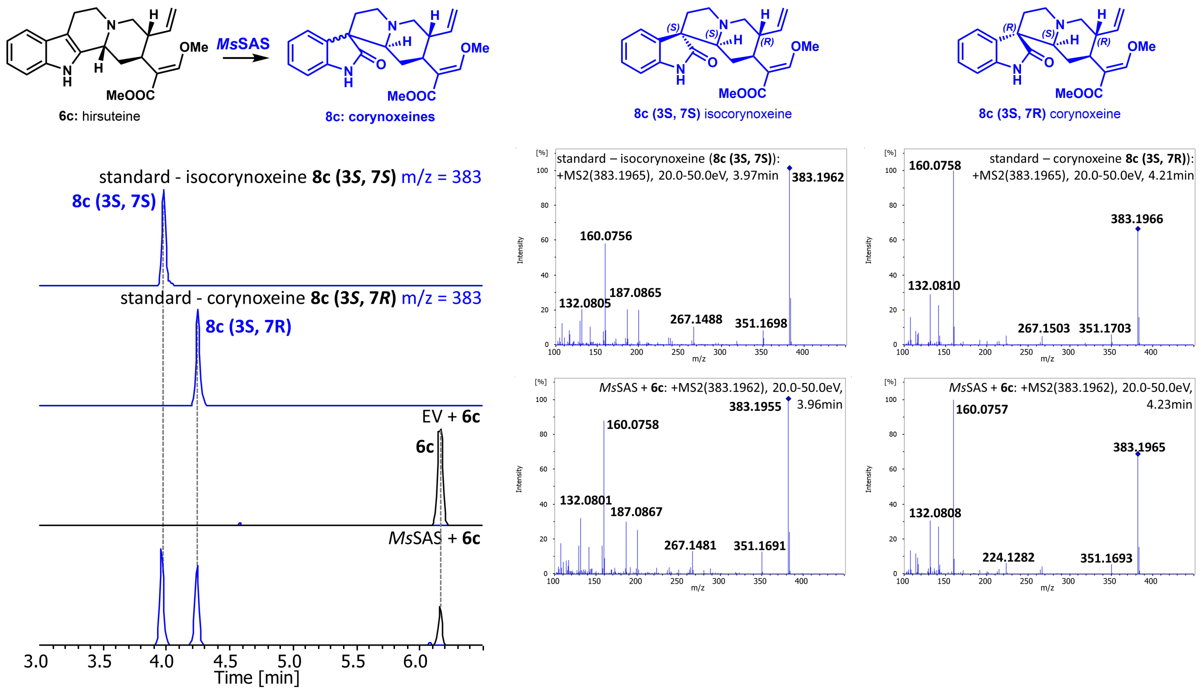


**Figure S21:** Representative UPLC-MS traces and analysis of compounds produced from hirsuteine (**6c**) following reaction with *Ms*SAS in *N. benthamiana*. We note coelution and comparable MS2 fragmentation with authentic standards of isocorynoxeine (**8c (**3S, 7S**)**) and corynoxeine (**8c (**3S, 7R**)**). In contrast to the previous report,[1] we assigned the products as corynoxeine and isocorynoxeine, not epi-corynoxeine (3*R*) based on NMR data. Assays were performed in biological triplicate for all reaction conditions.

#### Activity of *Ms*CO1/*Ms*DCR/*Ms*SAS on 20*S*-corynantheidine (3**a**)

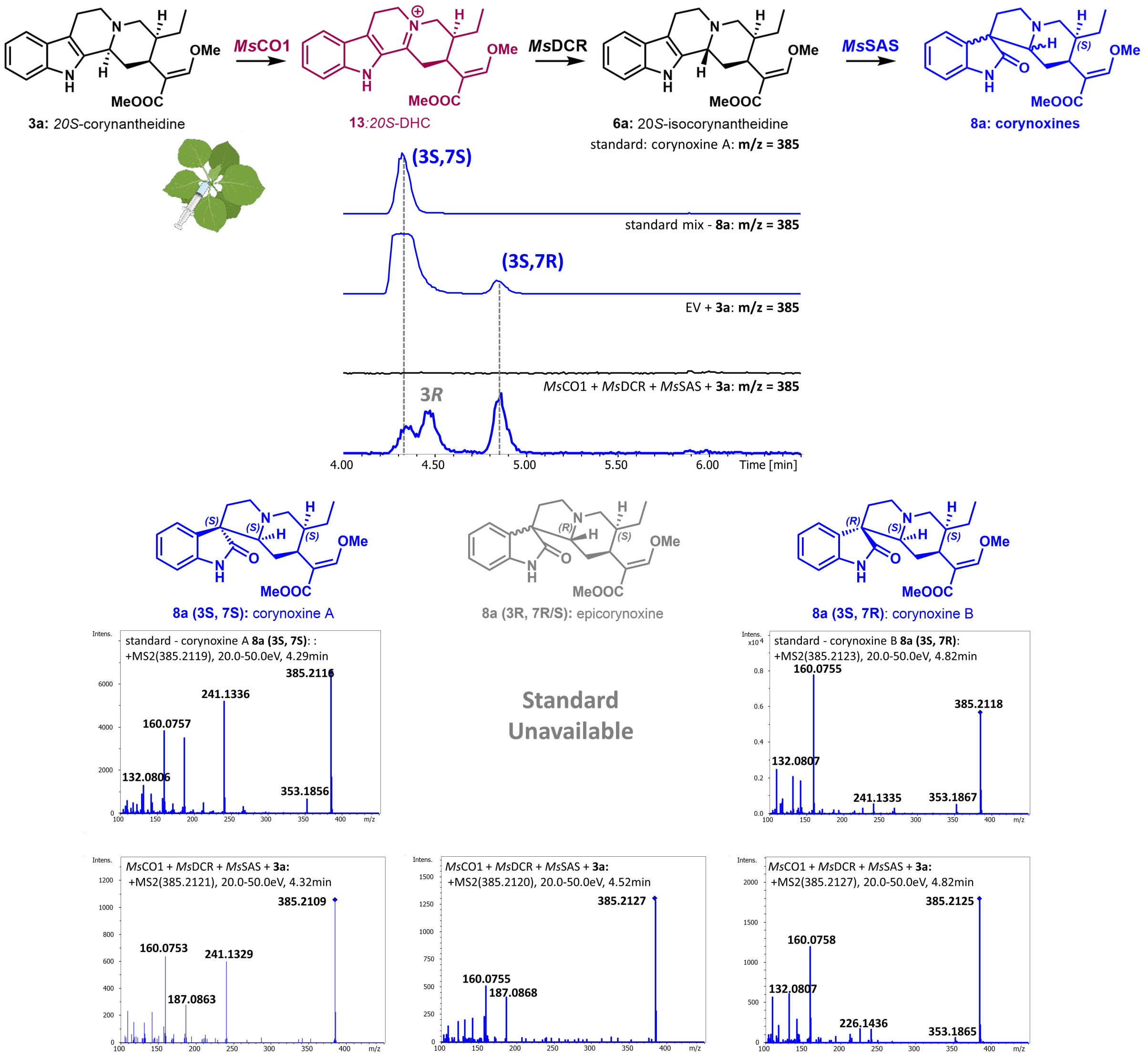


**Figure S22:** Evaluation of spirooxindole alkaloids produced via *Ms*CO1/*Ms*DCR/*Ms*SAS cascade reaction. We note the appearance of a peak corresponding to corynoxine A (**8a** (3*S*, 7*S*)) under these assay conditions. Assays were performed in biological triplicate for all reaction conditions. The 8a standard mix was overloaded to be able to visualize the minor isomer.

Activity of *Ms*CO/*Ms*DCR/*Ms*SAS on 20*S*-9-hydroxycorynantheidine (**4a**)

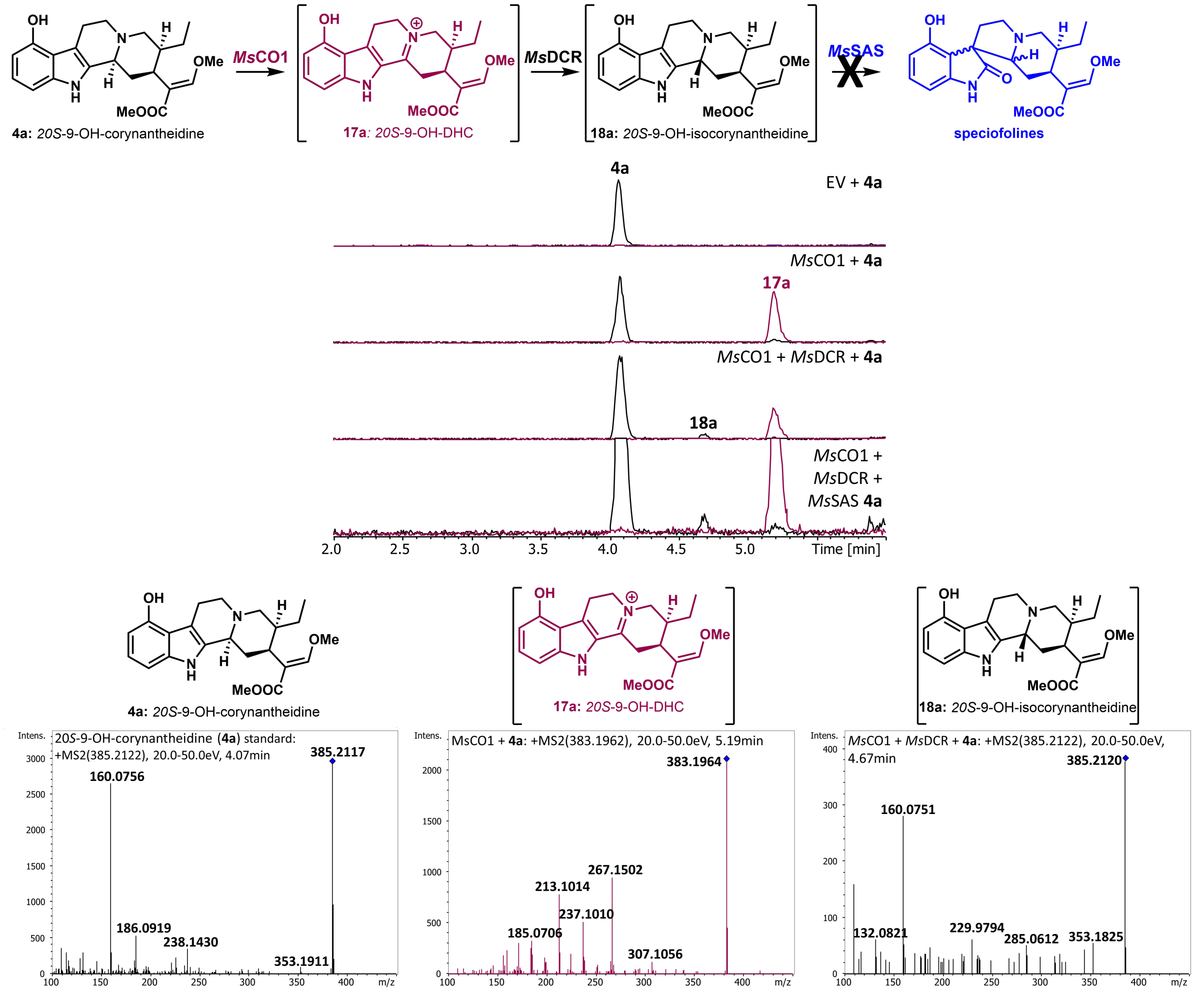


**Figure S23:** Representative UPLC-MS traces and analysis of compounds produced from a 20*S*-9-hydroxycorynantheidine (**4a**) standard following reaction with *Ms*CO1-*Ms*DCR-*Ms*SAS in *N. benthamiana*. A minor peak for a compound with a mass corresponding to 20*S*-9-OH-isocorynantheidine was detected, with similar fragmentation to the 20*S*-9-OH-corynantheidine standard. Assays were performed in biological triplicate for all reaction conditions. Compounds in brackets were putatively assigned without an authentic standard based on mass fragmentation. The bottom chromatogram is overloaded to be able to visualize the lack of spirooxindole formation.

### Activity of *Ms*CO/*Ms*DCR/*Ms*SAS on DHM (**14a**)

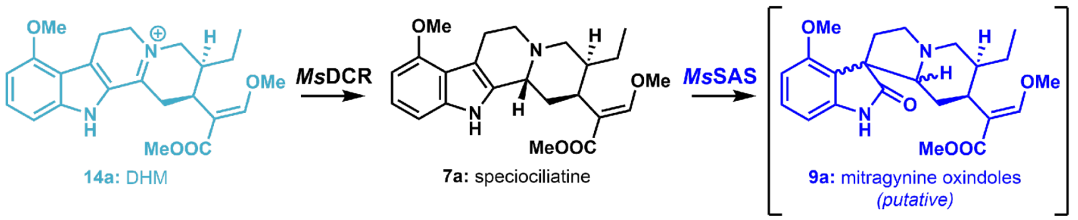


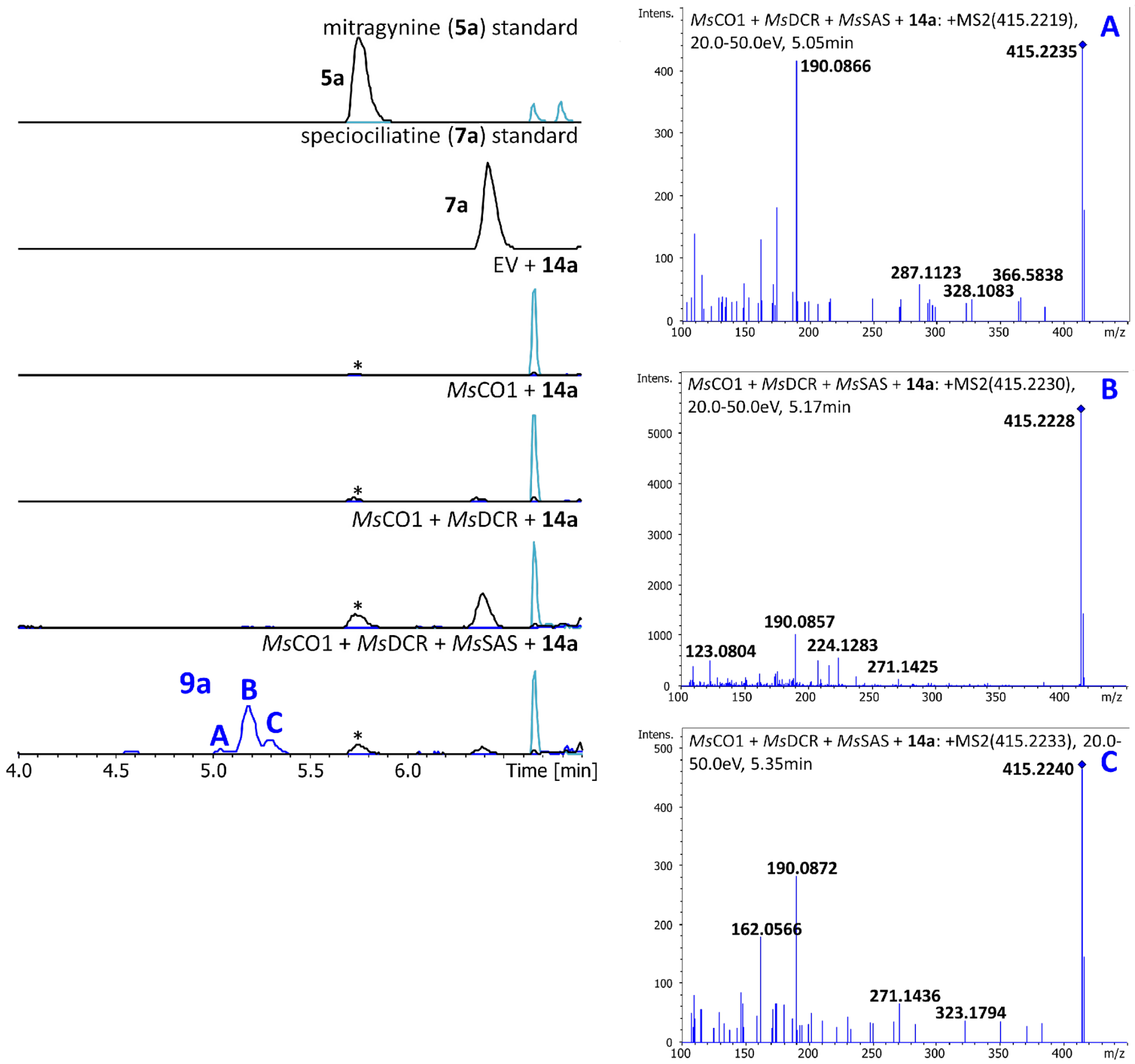


**Figure S24:** Representative UPLC-MS traces and analysis of compounds produced from synthesized dehydromitragynine (DHM, **14a**) following reaction with *Ms*CO1-*Ms*DCR-*Ms*SAS in *N. benthamiana*. Due to lack of standards, we did not attempt to assign structures to the observed mitragynine oxindoles (**9a**), but mass fragmentation is consistent with methoxylated oxindoles. *A small mitragynine (**5a**) impurity was present in the original DHM substrate. Assays were performed in biological triplicate for all reaction conditions.

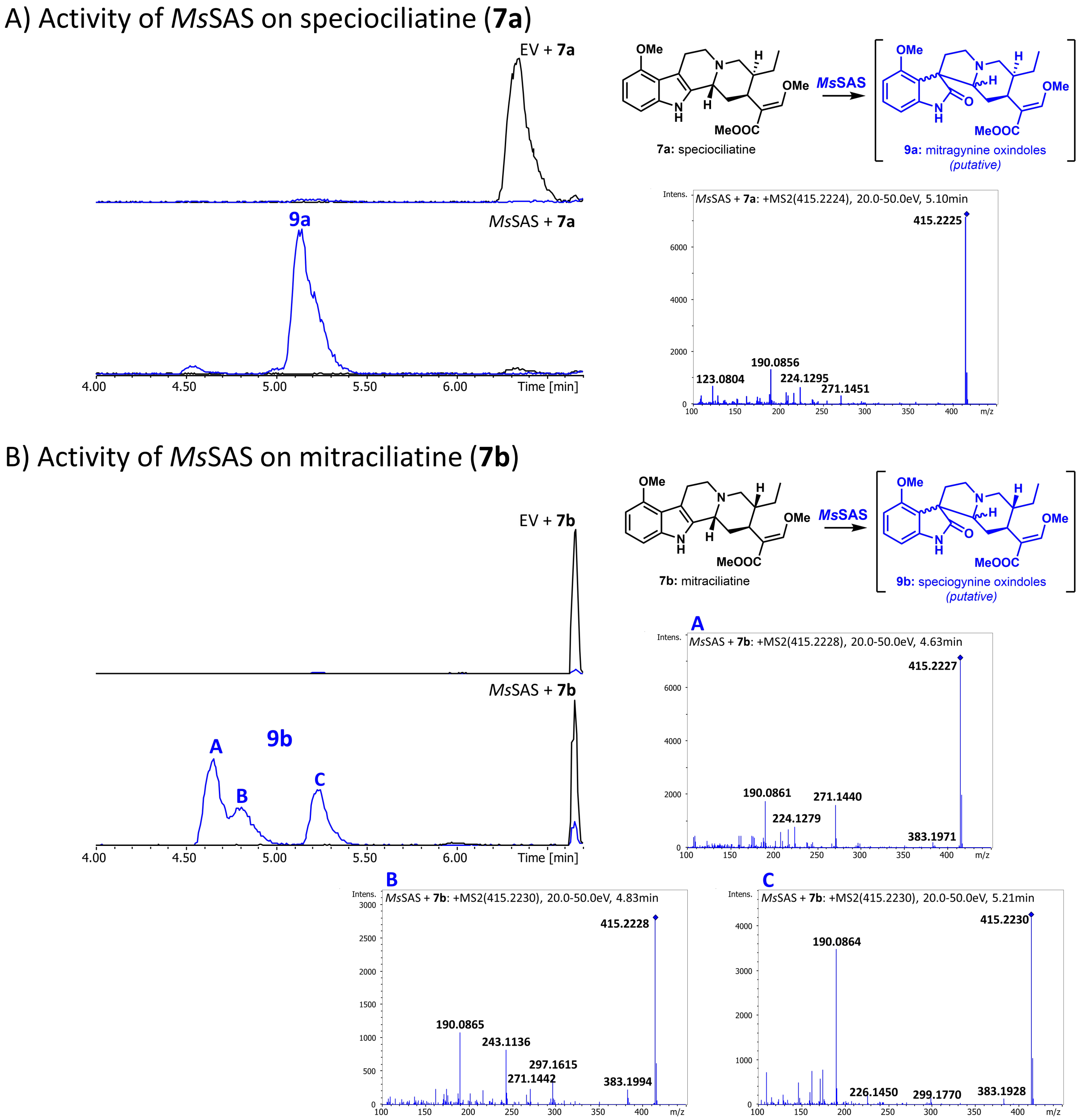


**Figure S25:** A) Representative UPLC-MS traces and analysis of compounds produced from speciociliatine (**7a**) following reaction with *Ms*SAS in *N. benthamiana*. We note the same major putative mitragynine oxindole product (**9a**) (designated in brackets) as from reaction with DHM (**14a**) with matching mass fragmentation patterns (see **Figure S24**). In contrast to *Ms*SAS products from non-methoxylated 20*S* substrates, reaction with speciociliatine (**7a**) produces primarily only a single major peak, although minor peaks are also observed. B) Representative UPLC-MS traces and analysis of compound produced from mitraciliatine (**7b**) following reaction with *Ms*CO1-*Ms*DCR-*Ms*SAS in *N. benthamiana*. Due to lack of standards, we did not attempt to assign structures to the observed speciogynine oxindoles (**9b**) (designated in brackets), but mass fragmentation is consistent with methoxylated oxindoles. Again, in contrast to *Ms*SAS products from non-methoxylated 20*R* substrates, reaction with mitraciliatine (**7b**) produces three major peaks, producing a mixture of 3*S*/3*R* isomers. Assays were performed in biological triplicate for all reaction conditions.

#### Activity of *Ms*CO/*Ms*DCR/*Ms*SAS on tetrahydroalstonine (**10a**)

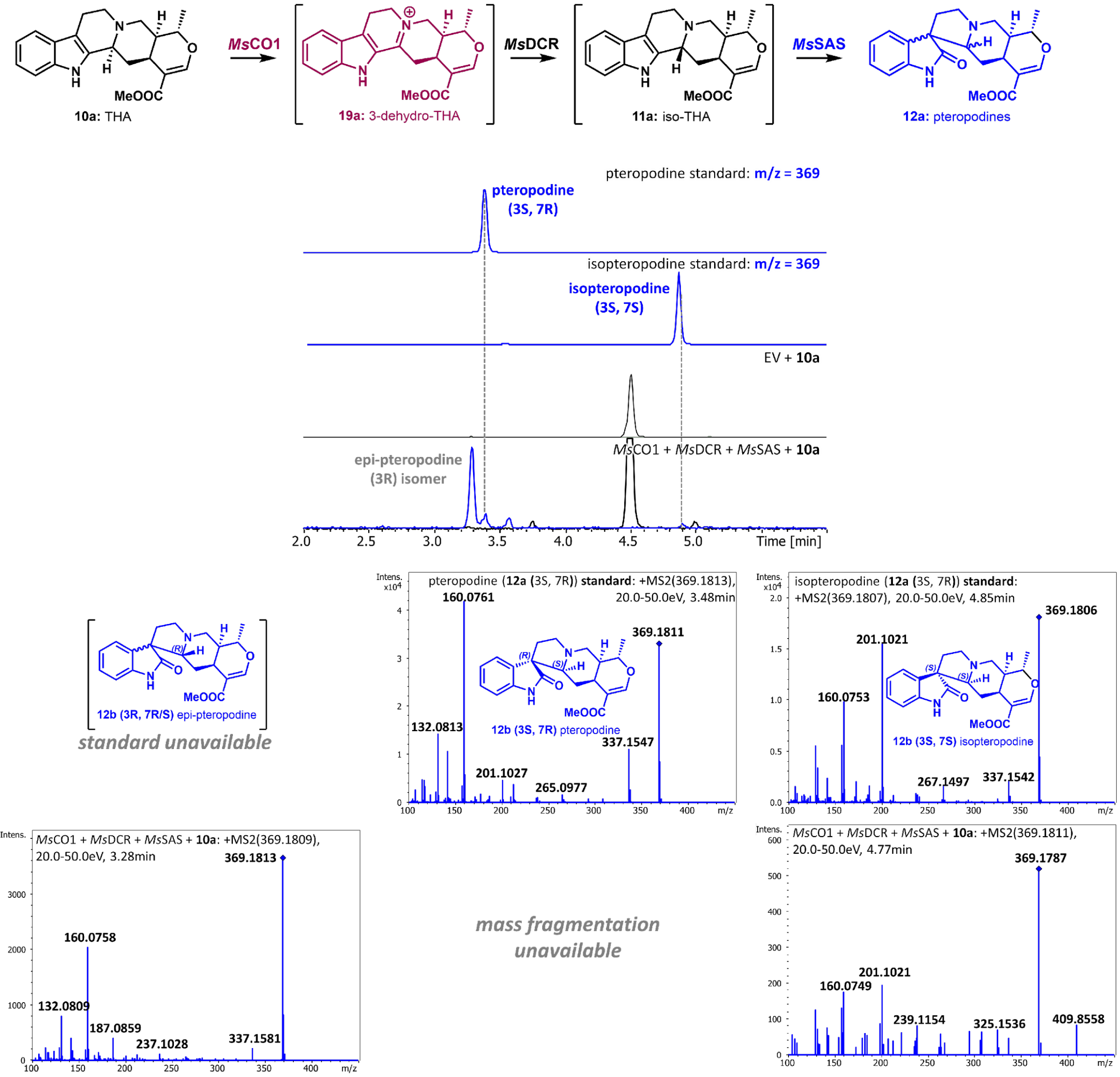


**Figure S26:** Representative UPLC-MS traces and analysis of pteropodine isomers (**12b**) produced from commercially available tetrahydroalstonine (THA) following reaction with *Ms*CO1-*Ms*DCR-*Ms*SAS in *N. benthamiana*. Compounds in brackets were putatively assigned without an authentic standard based on mass fragmentation. Assays were performed in biological triplicate for all reaction conditions.

### *in vitro* activity of *Ms*CO/*Ms*DCR/*Ms*SAS on ajmalicine (**10b**)

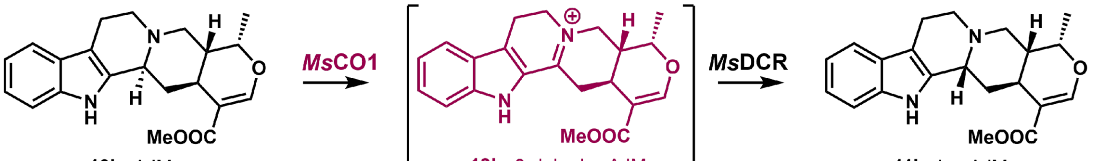


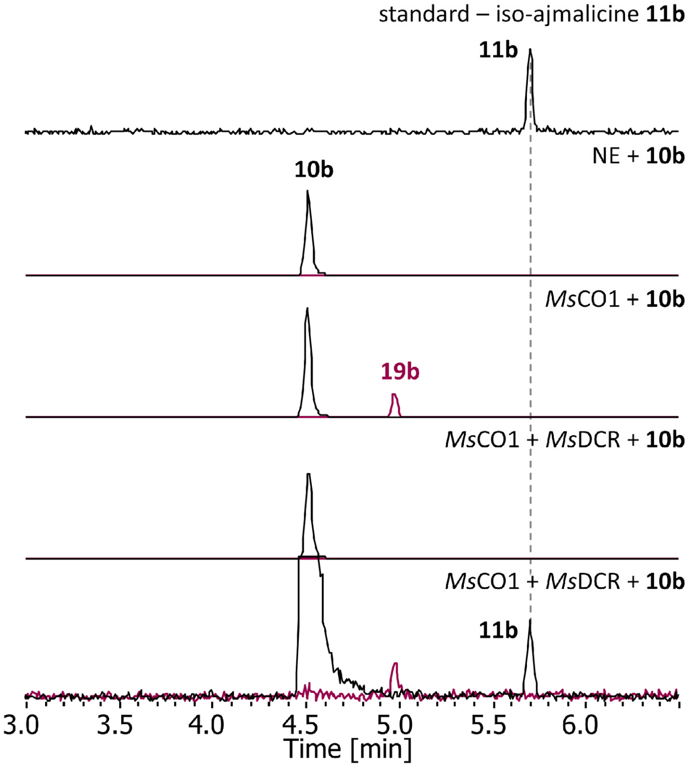


**Figure S27:** Representative UPLC-MS traces and analysis of reaction of *Ms*CO1 and *Ms*DCR with commercially available ajmalicine (**10b**, AJM). Formation of iso-ajmalicine (**11b**, iso-AJM) was poor, but the produced peak co-elutes with an authentic **11b** standard. Compounds in brackets were putatively assigned without an authentic standard based on mass fragmentation. This reaction only worked in *in vitro* reaction conditions. Reaction conditions: 25 µL total volume containing 100 µM ajmalicine (**10b**), 1 mM NADPH, 0.5 µM *Ms*CO1 and 5 µM *Ms*DCR, and 50 mM Tris-HCl pH = 7.5. Reactions were allowed to proceed at 25 ˚C for 16 h at ˚C.

#### Activity of *Ms*CO/*Ms*DCR/*Ms*SAS on mayumbine (**10c**)

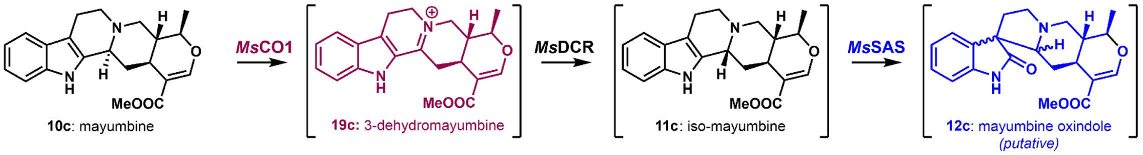


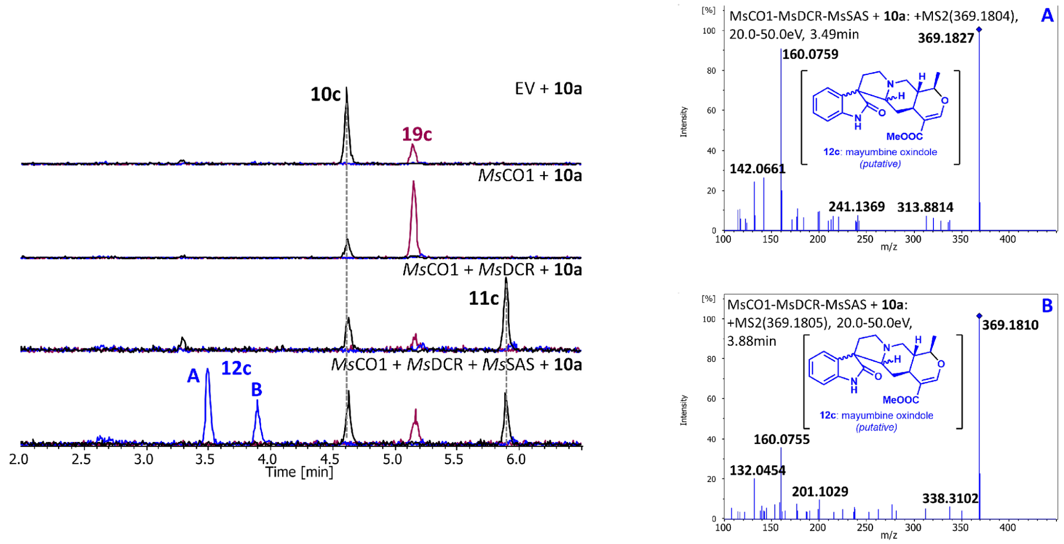


**Figure S28:** Representative UPLC-MS traces and analysis of compounds produced from commercially available mayumbine (**10c**) following reaction with *Ms*CO1-*Ms*DCR-*Ms*SAS in *N. benthamiana*. Only two putative mayumbine oxindole products (**12c**) were observed, but mass fragmentation is consistent with oxindoles: (*m/z* $[M+H]^+ = 160$). Compounds in brackets were putatively assigned without an authentic standard based on mass fragmentation. Assays were performed in biological duplicate for all reaction conditions.

#### Activity of *Ms*CO/*Ms*DCR/*Ms*SAS on yohimbine (**20a**)

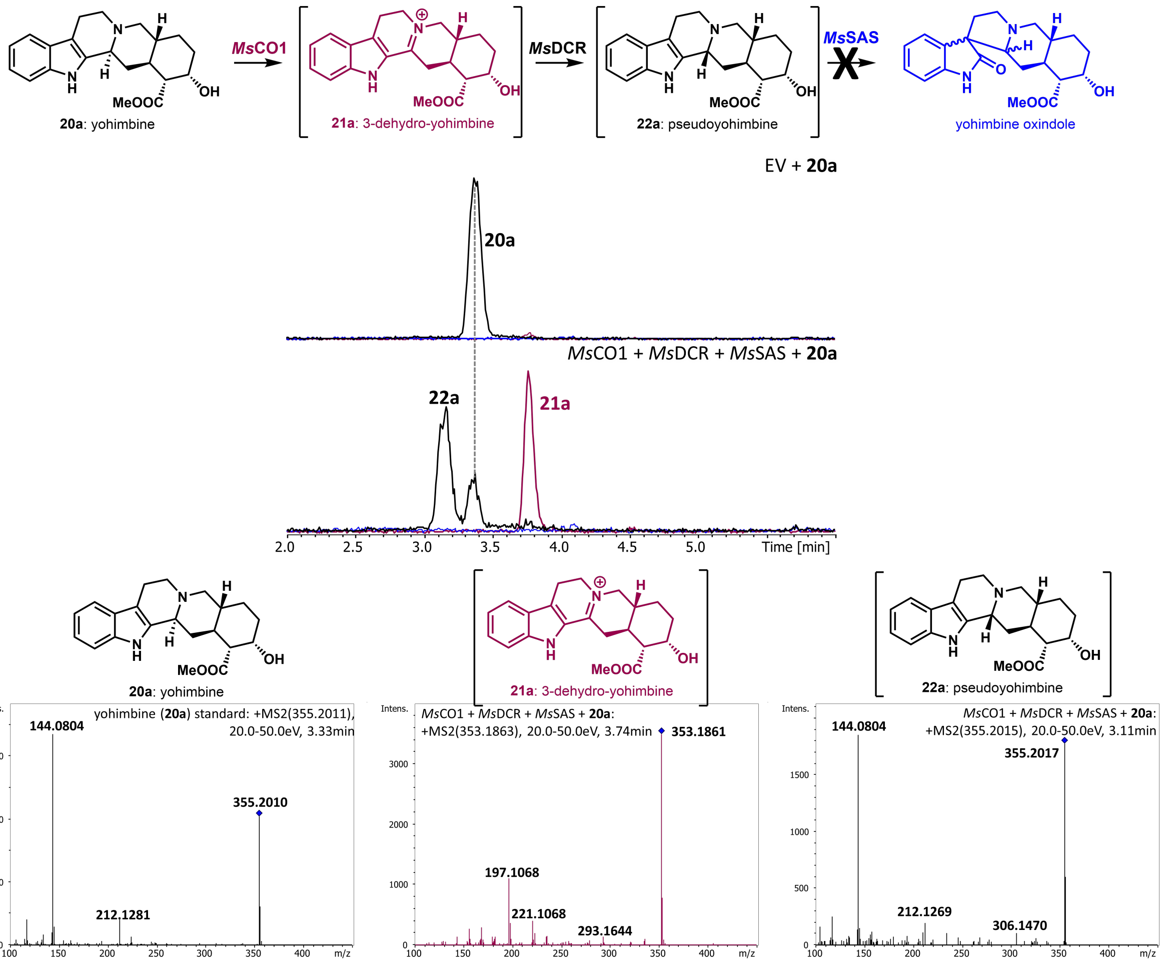


**Figure S29:** Representative UPLC-MS traces and analysis of compounds produced from commercially available yohimbine (**20a**) following reaction with *Ms*CO1-*Ms*DCR-*Ms*SAS in *N. benthamiana*. We note formation of a putative 3*R* epimer of yohimbine, pseudoyohimbine (**22a**) with comparable mass fragmentation to **20a**. Compounds in brackets were putatively assigned without an authentic standard based on mass fragmentation. Assays were performed in biological triplicate for all reaction conditions.

### Activity of *Ms*CO/*Ms*DCR/*Ms*SAS on rauwolscine (**20b**)

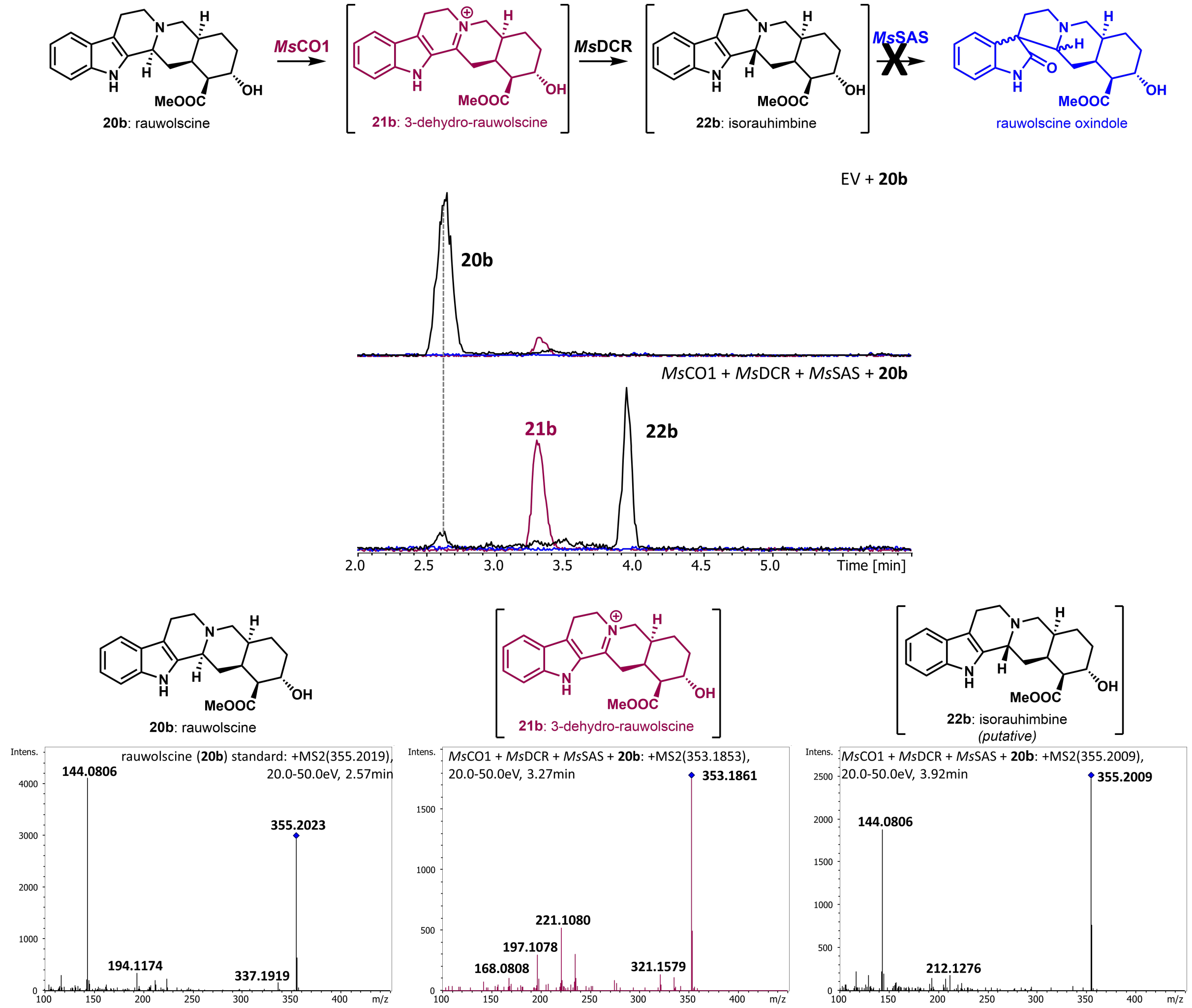


**Figure S30:** Representative UPLC-MS traces and analysis of compounds produced from commercially available rauwolscine (**20b**) following reaction with *Ms*CO1-*Ms*DCR-*Ms*SAS in *N. benthamiana*. We note formation of a putative 3*R* epimer of rauwolscine, isorauhimbine (**22b**). Compounds in brackets were putatively assigned without an authentic standard based on mass fragmentation. Assays were performed in biological triplicate for all reaction conditions.

#### Activity of *Ms*CO/*Ms*DCR/*Ms*SAS on corynanthine (**20c**)

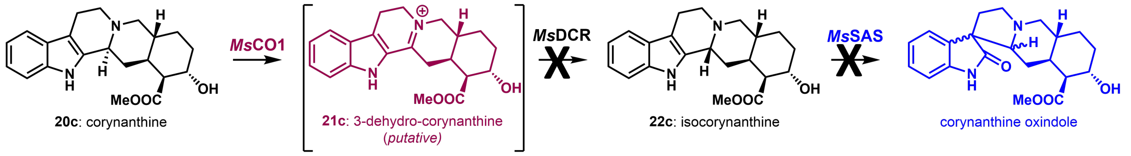


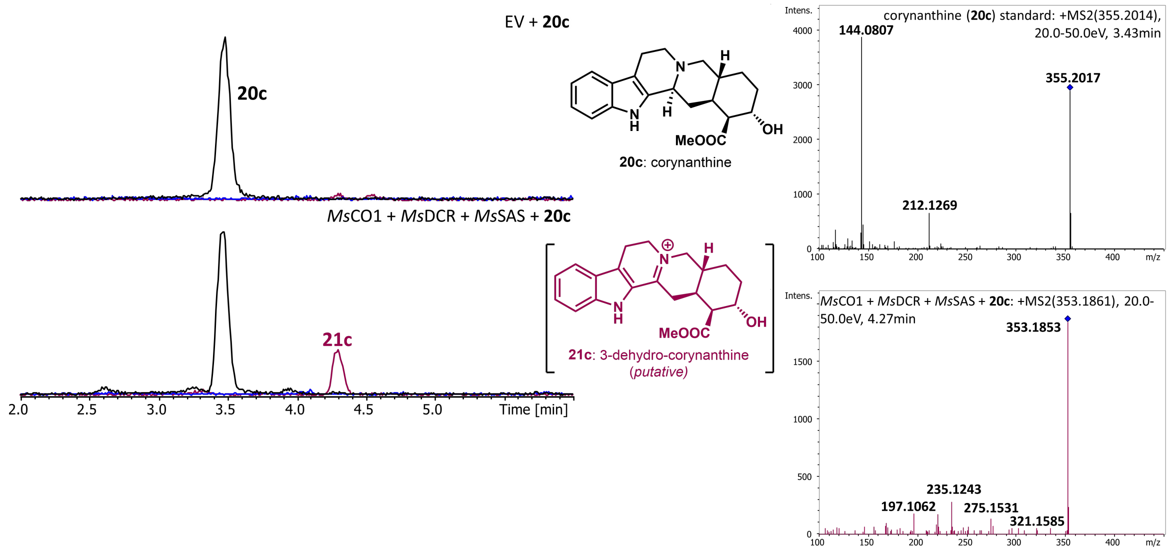


**Figure S31:** Representative UPLC-MS traces and analysis of compounds produced from commercially available corynanthine (**19c**) following reaction with *Ms*CO1-*Ms*DCR-*Ms*SAS in *N. benthamiana*. We note formation of a compound corresponding to a putative 3-dehydro-corynanthine (**21c**), but no significant formation of isocorynanthine (**22c**). Compounds in brackets were putatively assigned without an authentic standard based on mass fragmentation. Assays were performed in biological triplicate for all reaction conditions.

A) Biosynthesis of (20*S*)-spirooxindole alkaloids from tryptamine

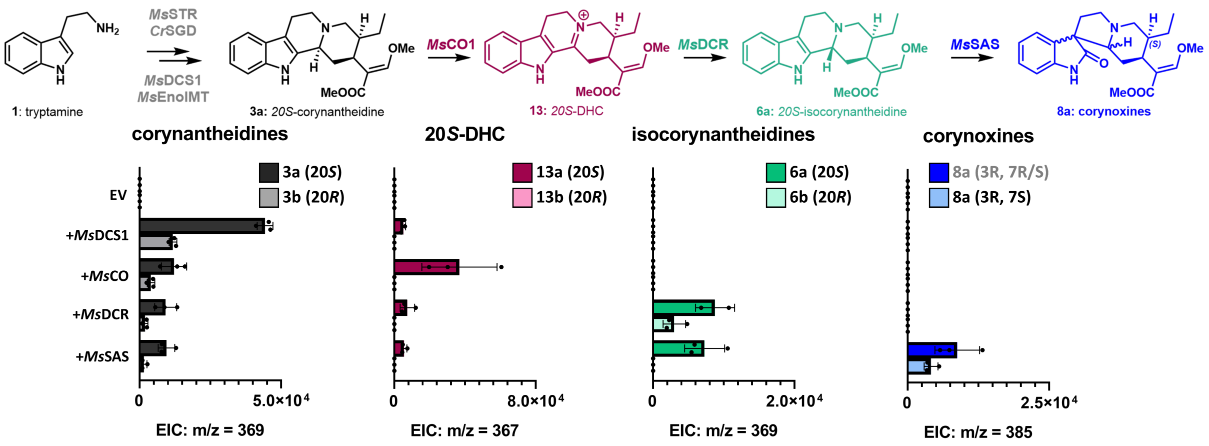


B) Biosynthesis of (20*R*)-spirooxindole alkaloids from tryptamine

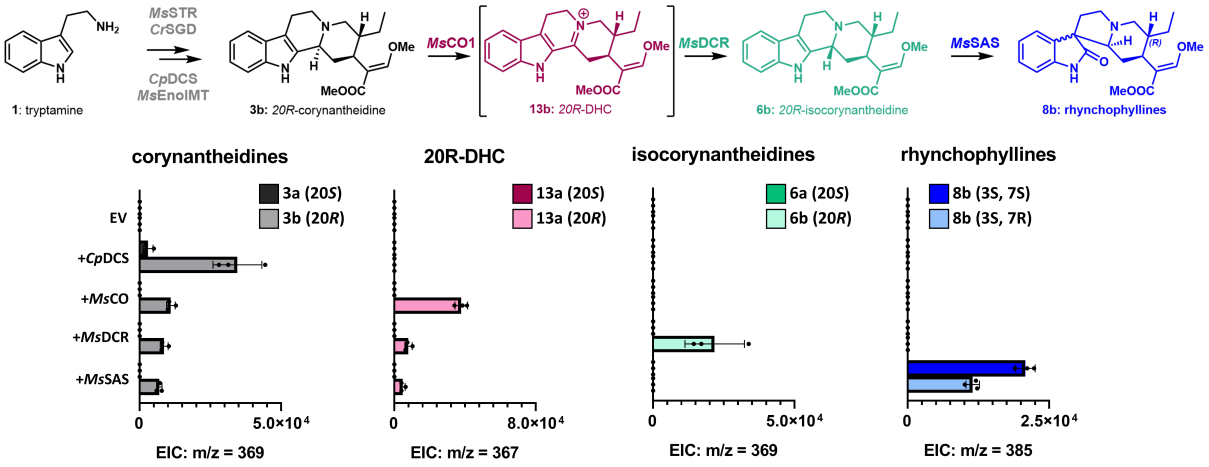


**Figure S32:** Pathway reconstruction in *N. benthamiana* from tryptamine to spirooxindole alkaloids with peak areas from extracted ion chromatograms (EICs) for provided *m/z* values. A) 20*S* biosynthetic pathway utilizing *Ms*DCS1. B) 20*R* biosynthetic pathway utilizing *Cp*DCS. Compounds in brackets were putatively assigned without an authentic standard based on mass fragmentation. Assays were performed in biological triplicate for all reaction conditions.

#### Biosynthesis of spirooxindole alkaloids from tryptamine: (20*S*)

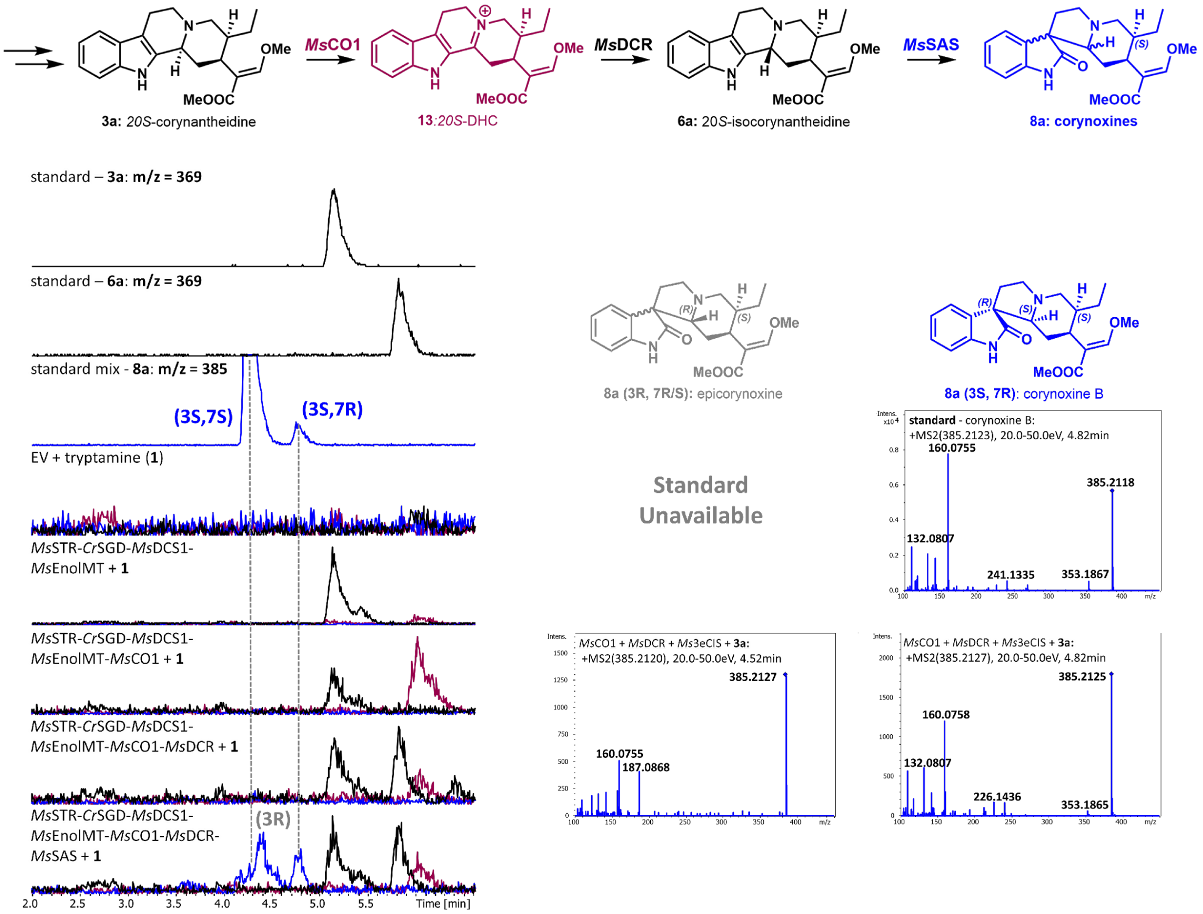


**Figure S33:** Representative UPLC-MS traces from pathway reconstruction for *Ms*DCS1 cascade corresponding to **Figure S32**. The major identified corynoxine (**8a**) peaks following *Ms*SAS infiltration are the putatively assigned 3*R* epicorynoxine isomer and corynoxine B (**8a (**3*S*, 7*R*)). Corynoxine A (**8a** (3*S*, 7*S*)) was not identified as an observable product. Additionally, the corynoxine B standard was found to isomerize relatively quickly to corynoxine A, as previously described.[2] Assays were performed in biological triplicate for all reaction conditions. LC-MS running conditions were optimized for separation/retention of **8a** peaks, resulting in less-defined peak areas corresponding to **3a**, **13a**, and **6a**. See **Figure S22** for additional spectra detailing *Ms*CO1/*Ms*DCR/*Ms*SAS activity on 20*S*-corynantheidine (**3a**).

Biosynthesis of spirooxindole alkaloids from tryptamine: (20*R*)

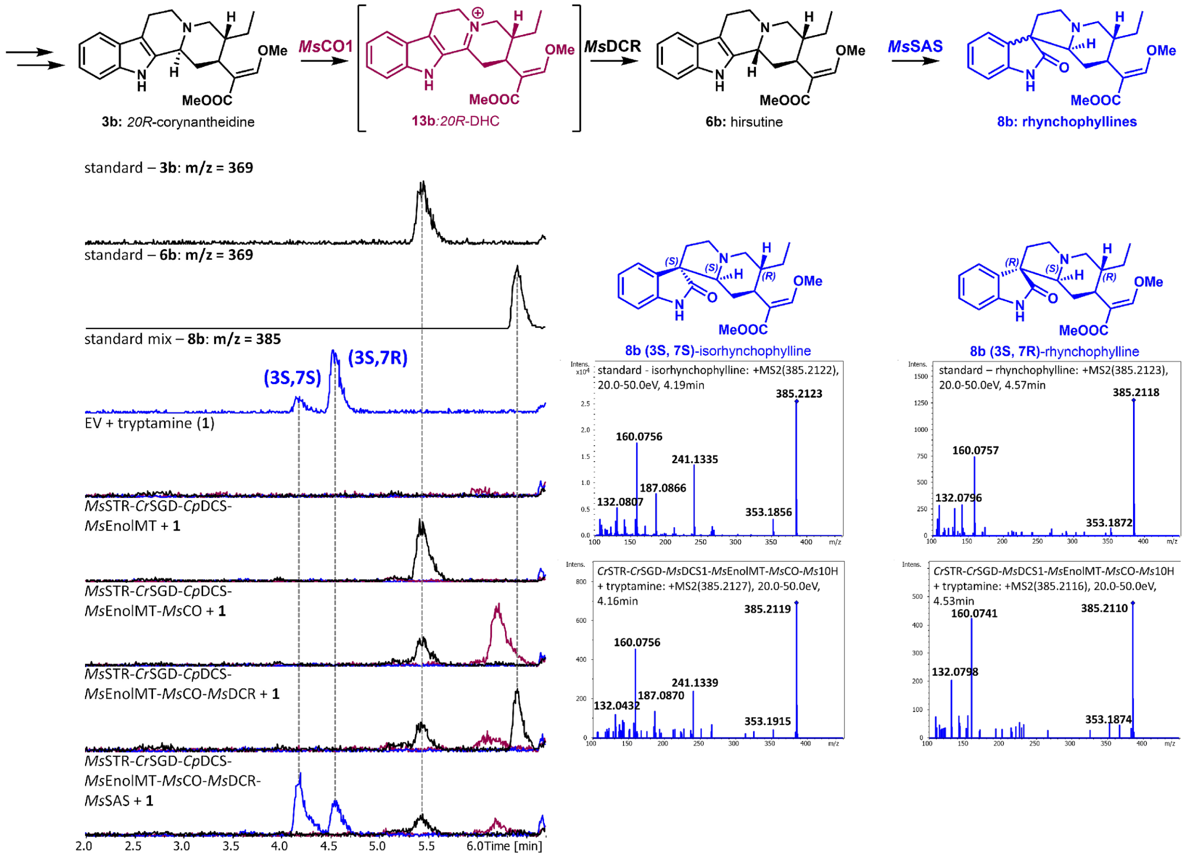


**Figure S34:** Representative UPLC-MS traces from pathway reconstruction for *Cp*DCS cascade corresponding to **Figure S32**. Two major spirooxindole peaks could be observed, corresponding to isorhynchophylline (**8b** (3*S*, 7*S*)) and rhynchophylline (**8b** (3*S*, 7*R*)) standards. The rhynchophylline standard was found to isomerize to isorhynchophylline. Assays were performed in biological triplicate for all reaction conditions. Compounds in brackets were putatively assigned without an authentic standard based on mass fragmentation. LC-MS running conditions were optimized for separation/retention of **8b** peaks, resulting in less-defined peak areas corresponding to **3b**, **13b**, and **6b**.

20*S*-9-hydroxy-dehydrocorynantheidine (**17a**) identified in Kratom young leaves

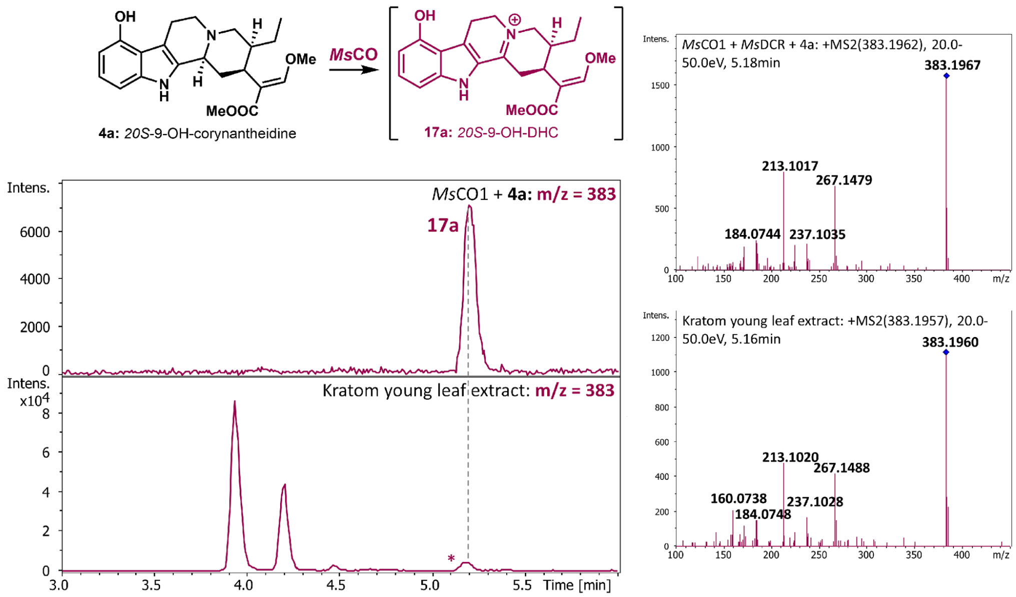


**Figure S35:** Comparison of enzymatically produced 20*S*-9-hydroxy-dehydrocorynantheidine (**17a**) with Kratom young leaf extract.

Isomerization of rhynchophyllines: **8b**

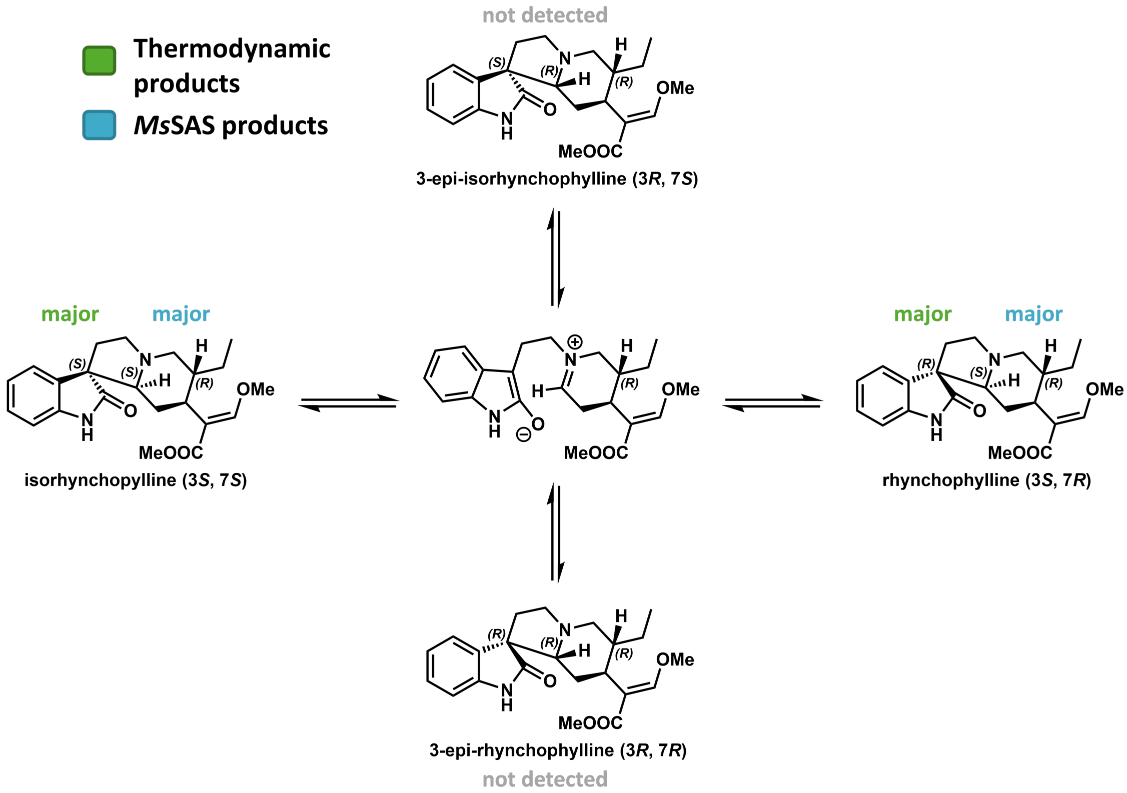


**Figure S36:** Comparison of observed products from *Ms*SAS originating from hirsutine (**6b**) and proposed thermodynamic distribution of product isomerization (see Laus 1996).[2]

#### Isomerization of corynoxines: **8a**

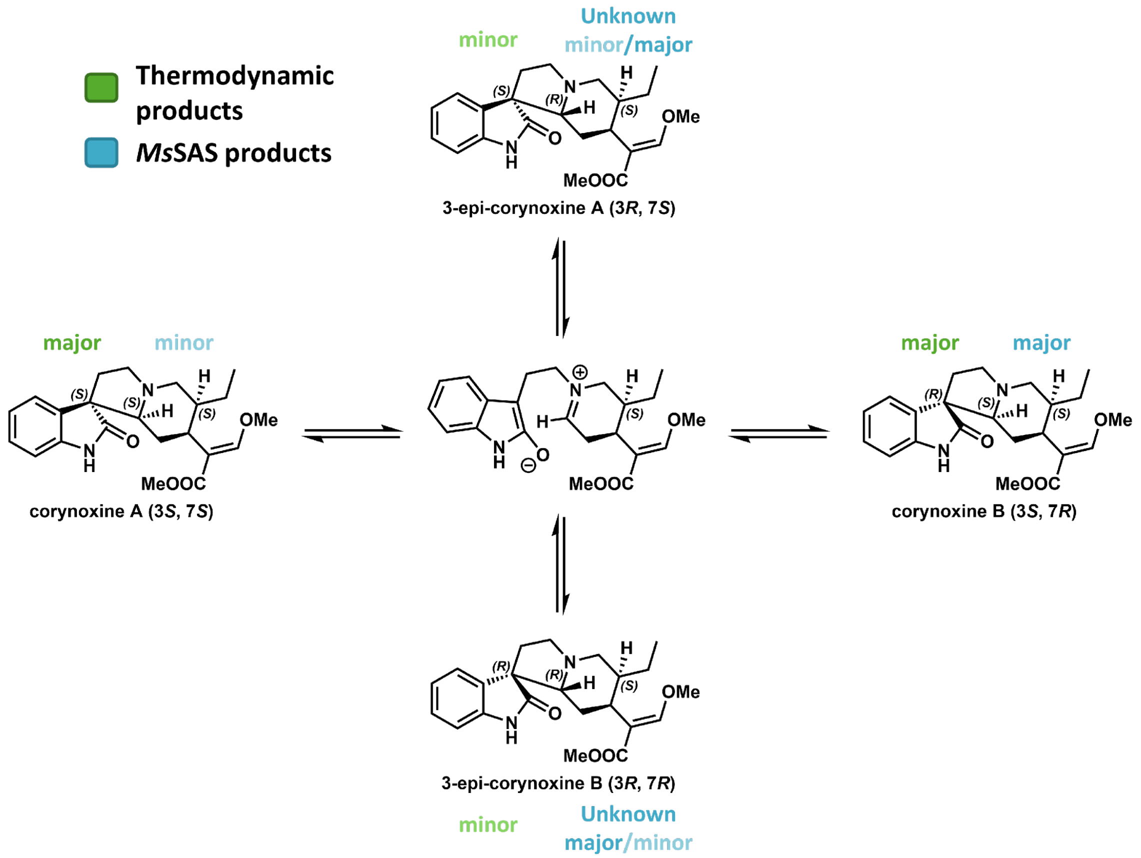


**Figure S37:** Comparison of observed products from *Ms*SAS originating from 20*S*-isocorynantheidine (**6a**) and proposed thermodynamic distribution of product isomerization (see Laus 1996).[2] *Since standards of 3-epi-corynoxine A or B could not be obtained, the precise identity of the remaining major product(s) in the enzymatic reaction could not be conclusively assigned.*

#### Supplementary Tables

**Table S1A:** Peak areas of identified Kratom metabolites from labelled substrate feeding. All values are averages of biological triplicates. *NA = not measured, ND = not detected.*

| METABOLITE AREAS | | 2a | 2b | 3a | 3b | 6a | 6b | 13 | 14 | 5a | 5b | 7a | 7b | 8 |
|---|---|---|---|---|---|---|---|---|---|---|---|---|---|---|
| **Young leaf** | **native** | **2E+05** | **5E+04** | **1E+07** | **1E+06** | **5E+06** | **2E+05** | **4E+07** | **6E+07** | **6E+07** | **1E+07** | **2E+07** | **5E+06** | **1E+08** |
| | d5-1 | 7E+05 | 3E+04 | 1E+07 | 1E+06 | 1E+06 | 2E+04 | 8E+04 | ND | 2E+06 | 4E+05 | 2E+06 | 3E+05 | 4E+06 |
| | d4-2a | 1E+07 | 9E+03 | 7E+06 | 2E+06 | 2E+05 | 3E+04 | NA | NA | ND | 3E+04 | 3E+05 | 7E+04 | 5E+05 |
| | d4-2b | 3E+05 | 1E+07 | 5E+05 | 2E+05 | ND | 3E+04 | NA | NA | ND | ND | 1E+05 | 2E+04 | 5E+04 |
| | d2-9OS | 2E+02 | 2E+04 | 4E+04 | ND | 2E+04 | 3E+04 | NA | NA | 4E+05 | 4E+05 | 1E+05 | 8E+04 | 3E+04 |
| **Stem** | **native** | **6E+05** | **7E+04** | **6E+06** | **1E+06** | **6E+07** | **6E+05** | **2E+07** | **5E+07** | **1E+07** | **2E+06** | **2E+08** | **1E+07** | **2E+08** |
| | d5-1 | 3E+07 | 8E+04 | 2E+07 | 1E+07 | 2E+06 | 2E+06 | 9E+03 | ND | ND | 9E+04 | 2E+06 | 4E+06 | 1E+06 |
| | d4-2a | 9E+06 | 2E+04 | 1E+07 | 2E+07 | 2E+06 | 2E+06 | NA | NA | ND | 3E+04 | 2E+06 | 3E+06 | 2E+06 |
| | d4-2b | 3E+05 | 3E+07 | 6E+05 | 4E+05 | 2E+05 | 6E+04 | NA | NA | ND | ND | 1E+06 | 1E+05 | 2E+05 |
| | d2-9OS | 9E+03 | 3E+04 | ND | ND | 1E+05 | 4E+04 | NA | NA | 6E+05 | 2E+06 | 1E+06 | 7E+04 | 2E+05 |
| **Mature leaf** | **native** | **9E+04** | **5E+04** | **3E+05** | **2E+05** | **4E+05** | **2E+05** | **ND** | **4E+06** | **2E+07** | **2E+06** | **1E+06** | **7E+05** | **3E+07** |
| | d5-1 | 1E+06 | 1E+05 | 4E+05 | 2E+05 | 1E+05 | 7E+04 | ND | ND | ND | 6E+03 | ND | 3E+04 | 1E+05 |
| | d4-2a | 8E+06 | 2E+04 | 5E+05 | 8E+05 | 2E+04 | 5E+04 | NA | NA | ND | ND | ND | ND | ND |
| | d4-2b | 3E+05 | 1E+07 | 3E+04 | 7E+04 | 9E+03 | 5E+04 | NA | NA | ND | ND | ND | ND | ND |
| | d2-9OS | 9E+03 | 2E+03 | ND | 6E+04 | ND | 3E+04 | NA | NA | ND | 9E+04 | ND | ND | ND |
| **Root** | **native** | **1E+05** | **5E+04** | **1E+06** | **9E+05** | **1E+06** | **3E+05** | **ND** | **1E+06** | **3E+05** | **2E+05** | **8E+06** | **3E+06** | **6E+06** |
| | d5-1 | 4E+05 | 1E+04 | 4E+04 | 1E+05 | 2E+04 | 8E+04 | ND | ND | ND | ND | ND | 2E+04 | ND |
| | d4-2a | 3E+05 | ND | 3E+05 | 5E+05 | 8E+04 | 3E+04 | NA | NA | ND | ND | ND | ND | ND |
| | d4-2b | 2E+04 | 1E+05 | 2E+05 | 4E+04 | 6E+04 | 4E+04 | NA | NA | ND | ND | ND | ND | 2E+05 |
| | d2-9OS | 2E+04 | 8E+03 | 1E+04 | 3E+04 | 6E+04 | 3E+05 | NA | NA | ND | ND | ND | 6E+04 | ND |

**Table S1B:** Corresponding standard deviations for above observed metabolite areas. *NA = not measured, ND = not detected.*

| STDEV | | 2a | 2b | 3a | 3b | 6a | 6b | 13 | 14 | 5a | 5b | 7a | 7b | 8 |
|---|---|---|---|---|---|---|---|---|---|---|---|---|---|---|
| **Young leaf** | **native** | **3E+04** | **2E+04** | **3E+06** | **3E+05** | **1E+06** | **2E+04** | 3E+05 | 1E+05 | **1E+07** | **2E+06** | **8E+06** | **6E+05** | **1E+07** |
| | d5-1 | 3E+05 | 1E+04 | 4E+06 | 7E+05 | 5E+05 | 6E+04 | 1E+06 | NA | 6E+05 | 2E+05 | 1E+06 | 6E+04 | 1E+06 |
| | d4-2a | 4E+06 | 8E+03 | 2E+06 | 3E+05 | 1E+05 | 2E+04 | NA | NA | ND | 5E+04 | 1E+05 | 8E+03 | 3E+05 |
| | d4-2b | 6E+04 | 6E+06 | 3E+04 | 1E+04 | ND | 4E+04 | NA | NA | ND | ND | 1E+04 | 9E+03 | 9E+04 |
| | d2-9OS | 3E+04 | 2E+04 | 4E+04 | ND | 2E+04 | 4E+04 | NA | NA | 3E+05 | 2E+05 | 7E+04 | 3E+04 | 4E+04 |
| **Stem** | **native** | **3E+05** | **2E+04** | **6E+05** | **2E+05** | **5E+06** | **7E+04** | **2E+05** | **1E+05** | **1E+06** | **1E+06** | **9E+06** | **8E+05** | **1E+07** |
| | d5-1 | 1E+07 | 4E+04 | 7E+06 | 5E+06 | 1E+06 | 6E+05 | 2E+06 | ND | ND | 7E+04 | 3E+05 | 1E+06 | 4E+05 |
| | d4-2a | 9E+05 | 9E+03 | 6E+06 | 6E+06 | 2E+06 | 7E+05 | NA | NA | ND | 4E+04 | 4E+05 | 6E+05 | 2E+06 |
| | d4-2b | 7E+04 | 1E+06 | 1E+05 | 6E+04 | 9E+04 | 4E+03 | NA | NA | ND | ND | 4E+04 | 2E+04 | 5E+04 |
| | d2-9OS | 2E+04 | 2E+04 | 2E+04 | 4E+04 | 9E+04 | 2E+04 | NA | NA | 7E+05 | 7E+05 | 1E+05 | 3E+04 | 1E+05 |
| **Mature leaf** | **native** | **9E+03** | **1E+04** | **ND** | **ND** | **4E+04** | **8E+03** | **ND** | **4E+04** | **2E+06** | **2E+05** | **1E+05** | **7E+04** | **3E+06** |
| | d5-1 | 5E+05 | 8E+04 | 5E+04 | 1E+05 | 1E+05 | 2E+04 | ND | ND | ND | 7E+04 | ND | 2E+04 | 1E+05 |
| | d4-2a | 3E+06 | 2E+04 | 5E+05 | 5E+05 | 3E+04 | 3E+04 | NA | NA | ND | ND | ND | ND | ND |
| | d4-2b | 8E+04 | 6E+05 | 6E+04 | 3E+04 | 1E+04 | 6E+04 | NA | NA | ND | ND | ND | ND | ND |
| | d2-9OS | 3E+04 | 2E+04 | ND | 3E+04 | ND | 2E+04 | NA | NA | ND | 2E+04 | ND | ND | ND |
| **Root** | **native** | **1E+04** | **2E+04** | **3E+05** | **3E+05** | **2E+05** | **1E+05** | **ND** | **2E+04** | **7E+04** | **3E+04** | **5E+05** | **2E+05** | **8E+05** |
| | d5-1 | 5E+05 | 2E+04 | 7E+04 | 2E+05 | 4E+04 | 8E+04 | ND | ND | ND | ND | ND | 1E+05 | ND |
| | d4-2a | 9E+04 | ND | 2E+05 | 1E+04 | 5E+04 | 5E+04 | NA | NA | ND | ND | ND | ND | ND |
| | d4-2b | 3E+04 | 5E+04 | 1E+05 | 5E+04 | 6E+04 | 2E+04 | NA | NA | ND | ND | ND | ND | 2E+05 |
| | d2-9OS | 1E+04 | 5E+03 | 4E+04 | 2E+04 | 3E+04 | 1E+05 | NA | NA | ND | ND | ND | 4E+04 | ND |

### Materials and Methods

#### Plants and Plant Growth

*M. speciosa* *"*Green Thai*"* plants were a generous gift from Satya Swathi Nadakuduti (University of Florida) and were purchased from a supplier in Thailand. Plants were kept on a standard soil mix in the greenhouse (Jena, Germany) at 24-32 °C during the day and 22-24 °C during the night (summer) and at 18-23 °C during the day and 18-20 °C during the night (winter). Relative humidity was kept between 60% and 80%. Plants were propagated via cuttings.

*M. speciosa* *"*Rifat*"* plants were grown on a standard soil mix in the greenhouse (Athens, GA). Culture conditions were set to 28 °C (day) and 19 °C (night) following a 15-h light/9-h dark photoperiod with 23 DLI.

*Nicotiana benthamiana* plants were grown on a standard soil mix in the greenhouse. Culture conditions were set to 22 °C, 60% relative humidity and followed a 16-h light/8-h dark photoperiod. Tobacco plants were usually grown for at least 3 weeks but no longer than four weeks prior to infiltration with *Agrobacterium tumefaciens* GV3101. Plant watering was performed as needed.

#### Chemicals

All chemicals used in this study were purchased as molecular biology grade or higher from commercial vendors (*Sigma Aldrich*, *Thermo Fischer*, etc.) unless denoted differently. Kratom alkaloid standards were obtained from the following sources: mitragynine (**5a**) from *Biosynth Ltd.*; speciogynine (**5b**), paynantheine (**5c**), hirsutine (**6b**), hirsuteine (**6c**), speciociliatine (**7a**), corynoxine A (**8a (3*S*, 7*S*)**), corynoxine B (**8a (3*S*, 7*R*)**), rhynchophylline (**8b (3*S*, 7*S*)**), and isorhynchophylline (**8b (3*S*, 7*R*)**) were obtained from *Cayman Chemical*. Yohimbine (**20a**), rauwolscine (**20b**), and corynanthine (**20c**) were obtaind from *Extrasynthese*. Isopteropodine (**12a (3*S*, 7*S*)**) and pteropodine (**12a (3*S*, 7*R*)**) were purchased from *Phytolab*. Corynoxeine (**8c (3*S*, 7*R*)**) and isocorynoxeine (**8c (3*S*, 7*S*)**) were purchased from *MedChemExpress*. Mitraphylline (**12b (3*S*, 7*R*)**), 20*S*-9-hydroxycorynantheidine (**4a**), (20*S*)-corynantheidine (**3a**) were kindly gifted to us by Christopher McCurdy and iso-ajmalicine (**11b**) and isomitraphylline (**12b (3*S*, 7*S*)**) were provided by Prof. Adrianna Lopez. (20*R*)-corynantheidine (**3b**), mitraciliatine (**7b**), and 20*S*-isocorynantheidine (**3b**) standards were isolated from Rifat Kratom leaf tissue provided by Prof. Robin Buell. Unless otherwise denoted, all reagents were obtained from commercial sources and used without any further purification. Thin layer chromatography (TLC) and preparative TLC (PTLC) were carried out on aluminum-backed silica gel 60 F254 plates (Merck) and visualized using UV254 nm light detection.

**Molecular biology kits**

All molecular biology kits were used according to the manufacturer´s instructions, unless specified. The RNeasy Mini Kit (*Qiagen*) was used for RNA extraction (*vide infra*). cDNA was subsequently prepared using iScript reverse transcriptase (*Bio-Rad*), following manufacturer's instructions. For genes or gene fragments destined for downstream applications the SuperFi polymerase was used for amplification. Gene fragments were purified by agarose gel electrophoresis (1% agarose; 120 V, 40 min) and extracted from the gel using a Zymoclean™ Gel DNA Recovery Kit (*Zymo*). All oligonucleotide primers were synthesized by and obtained from *Sigma Aldrich*. Gene cloning was routinely performed using an In-Fusion kit (Clontech *Takara*) or via Gibson assembly. Plasmid DNA was isolated from bacterial cultures using the Wizard® *Plus* SV Minipreps DNA Purification System kit (*Promega*).

**Isotopologue feeding of *M. speciosa* tissue**

Tissue samples of *M. speciosa* were collected from recently pruned, four-year-old plants (pruning greatly increased the availability of fresh young leaf tissue). Samples for metabolomic analysis were freshly extracted in MeOH prior to LC-MS analysis. Samples for isotopologue feeding were prepared in the following way: 5 mg leaf disks, 5 mm cut stem disks, or 5 mg cut roots were obtained fresh from plants. These samples were incubated with 200 µl $H_2O$, 1 mM $d_5$-trypamine (***$d_5$*-1**), $d_4$-strictosidine (***$d_4$*-2a**), $d_4$-vincoside (***$d_4$*-2b**), $d_2$-9-methoxystrictosidine (***$d_2$*-9OS**) or $d_4$-serotonin for 24 h at 25 °C. Any remaining liquid was removed, and the samples were extracted 1 h with 500 ml MeOH, filtered, and then analyzed via LC-MS. The remaining issue was immediately snap frozen in liquid nitrogen and stored at –80 °C indefinitely. For samples with positive feeding results (or negative for mature leaves and roots), the non-cut material was used for subsequent RNA extraction and RNA-sequencing.

**RNA purification and sequencing**

Total RNA of *M. speciosa* (roots, stem, young leaves, mature leaves) was extracted using the RNeasy Mini Kit (*Qiagen*) according to manufacturer´s instructions with additional on-column DNAse incubation in biological triplicate. The quality of obtained RNA was analysed using an *Implen* NanoPhotometer® N60. All samples except roots satisfied the necessary requirements for total RNA sequencing (≥ 100 ng; A260/280 = 1.8-2.2; A260/230 ≥ 1.8) and were submitted to Novogene (https://en.novogene.com/) for total RNA sequencing using the company´s standard protocols for library preparation and RNA-Seq. ≥ 30 M raw sequencing reads (Illumina, 150 bp paired-end) were acquired per sample. Pac-BIO *de novo* transcriptome assembly was prepared from a pooling of the above RNA samples using the PACBIO_SMRT platform (Pacbio Sequel II CSS mode, single end).

**Coexpression/homology analysis for gene discovery**

The above Pac-BIO assembled transcriptome was used for transcript analysis with predicted functions annotated via BLASTing to the SwissProt database. FPKM counts were used to evaluate transcript expression abundances. Pearson correlation coefficients were calculated using Microsoft Excel using the expression profile of *Ms*EnolMT or *Ms*CO1. For reductase candidates, annotated reductases/ dehydrogenases with high homology to *Cr*THAS1 and *Ms*DCS1 were chosen as candidates for screening.

**Cross-species transcriptome analysis**

Three Naucleeae and three non-Naucleeae transcriptomes were used for cross-species transcriptome analysis. These transcriptomes were obtained from the following sources. Publicly available RNA - sequencing data was obtained from the European Nucleotide Archive (https://www.ebi.ac.uk/ena/browser/home). The following run was downloaded: *Coffea arabica* (SRR17345234). The *Catharanthus roseus* transcriptome was obtained from BioProject accession PRJNA847226. The *Cinchona pubescens* transcriptome was obtained from Lombe *et al.* 2024.[3]

The *Uncaria rhynchophylla* transcriptome was obtained from BioProject (https://ngdc.cncb.ac.cn/bioproject). The following runs were downloaded from project PRJNA792441: leaf-1 (SRX13561691) and leaf-2 (SRX13561691). Trinity v2.15.1 was used for the assembly on the Galaxy server, with the parameters "paired-end data" and "--no_normalize_reads" for the assembly.[4]
For the *Uncaria guianensis* transcriptome, libraries were prepared using the KAPA Stranded RNA-Seq Library Preparation Kit (KR0934) with NEBNext Multiplex Oligos for Illumina (E7335). Sequencing was performed on an Illumina HiSeq 2500 to 150 nt in paired end mode by the RTSF Genomics Core at Michigan State University. Raw reads were trimmed using Cutadapt v1.1[5] with the parameters, "--trim-n --quality-cutoff 20,20 --minimum-length 30 --times 3 -a AGATCGGAAGAGCACACGTCTGAACTCCAGTCACNNNNNNATCTCGTATGCCGTCTTCTGCTTG -A AGATCGGAAGAGCGTCGTGTAGGGAAAGAGTGTAGATCTCGGTGGTCGCCGTATCATT". Trimmed reads were used for transcriptome assembly using Trinity v2.3.2 with the parameters "--SS_lib_type RF" and "--no_normalize_reads" for the assembly.[4]

**Identification of closest homologs to Kratom genes in available transcriptomes and sequence similarity network analysis**

Methyltransferases:

Based on SwissProt gene annotation, the amino acid sequences of all Kratom transcripts in the Pac-BIO assembly that were annotated as a 'methyltransferase' were obtained (205 transcripts). Using an in-house Python script, for each of the six additional transcriptomes (see above), the ORF of every transcript (starting with ATG) with a minimum amino acid length of 100 residues was translated. The peptides were then aligned to the Kratom methyltransferase list, and the gene from each transcriptome with the highest scoring alignment (using Python Bio.Align's scoring metric) was obtained as a peptide sequence. These genes along with the list of Kratom genes was aligned using the Enzyme Function Initiative-Enzyme Similarity Tool (EFI-EST) server (https://efi.igb.illinois.edu/efi-est/).[6] Clusters were generated with an initial alignment score threshold of 50. Using CytoScape, clusters were visualized, and the alignment score threshold increased until clusters not containing *C. roseus*, *C. arabica*, or *C. pubescens* genes emerged (*Naucleeae*-specific clusters (NSCs)). We observed a clear NSC-cluster containing *Ms*EnolMT emerging with an alignment score threshold of 100.

Reductases:

As detailed above, the amino acid sequences of all Kratom transcripts in the Pac-BIO assembly that were annotated as either a 'reductase' or 'dehydrogenase' were obtained (605 transcripts). Kratom reductase candidates and their closest homologs from the other transcriptomes (see above for details) were then aligned using EFI-EST and clusters were generated with an initial alignment score threshold of 50. Using CytoScape, clusters were visualized, and the alignment score threshold increased until NSCs emerged. We started observing NSCs with an alignment score threshold of 145, and identified the Kratom genes within these clusters as candidates for screening.

P450s/oxidases:

As detailed above, the amino acid sequences of all Kratom transcripts in the Pac-BIO assembly that included keywords 'P450,' 'oxidase,' 'hydroxylase,' or 'monooxygenase' were obtained (215 transcripts). Kratom oxidase candidates and their closest homologs from the other transcriptomes (see above for details) were then aligned using EFI-EST and clusters were generated with an initial alignment score threshold of 50. Using CytoScape, clusters were visualized, and the alignment score threshold increased until NSCs emerged. We started observing NSCs with an alignment score threshold of 170, and identified the Kratom genes within these clusters as candidates for screening. In addition, genes in Kratom-specific clusters and unclustered Kratom genes were also possible candidates for screening.

**Cloning of gene candidates**

Primers containing overhanging ends homologous to a modified 3Ω1 vector[7] were used to amplify full-length genes via PCR. *M. speciosa* cDNA from either stem or young leaf was used as the template. Resultant amplicons were purified via gel electrophoresis. Empty 3Ω1 vector was digested via BsaI (*Thermo Fischer*) and purified via the DNA Clean & Concentrator™-5 (*Zymo*) kit. In-Fusion cloning (*Clontech Takara*, manufacturer´s instructions) was fused with the gene of interest to assemble the final plasmid. The In-Fusion crude reaction was transformed into chemically competent *E. coli* TOP10 cells (*Thermo Fischer*) and plated onto LB agar supplemented with spectinomycin (200 µg/mL) and incubated at 37 °C for 16 h. Plasmids of positive transformants were isolated from overnight cultures (37 °C, 225 rpm, 2 mL LB + spectinomycin) and correct cloning was confirmed by Sanger sequencing (*Azenta Life Sciences*).

**Transformation of *Agrobacterium tumefaciens* GV3101**

Electrocompetent cells of *Agrobacterium tumefaciens* GV3101 (*Goldbio*) were thawed on ice and mixed with plasmid DNA (~500 ng) that had been verified by Sanger sequencing. Immediately post-thawing, the cell suspension was transferred to pre-chilled electroporation cuvette and cells were electroporated using a MicroPulser™ (BioRad) at 2.2 kV. Cells were mixed with 0.6 mL LB medium and recovered at 28 °C/225 rpm for 3 h prior to plating on selective LB agar plates (supplemented with 20 µg/mL rifampicin, 50 µg/mL gentamycin and 200 µg/mL spectinomycin). Plates were kept at 28 °C for 2 d. Single colonies were used to inoculate liquid cultures. Liquid cultures were prepared as 5 mL cultures (supplemented with 20 µg/mL rifampicin, 50 µg/mL gentamycin and 200 µg/mL spectinomycin) and cultivated at 28 °C and 250 rpm for up to 24 h. 50 % glycerol stocks were prepared thereof, snap frozen in liquid nitrogen and stored at –80 °C indefinitely. The remaining culture was then used for transient expression in *Nicotiana benthamiana*.

**Transient expression of gene candidates in *Nicotiana benthamiana***

Transient expression of gene candidates in *N. benthamiana* was performed as previously reported by Sparkes et al. 2006.[8] The cells containing the gene of interest in the above cultures were collected by centrifugation (4000 x g, 5 min). Cells were resuspended in 5 mL infiltration buffer (27.8 mM glucose, 100 µM acetosyringone, 50 mM MES, 2 mM $Na_3PO_4$, pH = 6.0) to an optical density $OD_{600}$ of ~ 0.6. Upon infiltration of multiple Agrobacterium strains the strains were diluted so that the final

$OD_{600}$ was < 1 (equal concentration for each strain). Resulting suspensions were incubated at 25 °C for 1 h and then infiltrated into the underside of 3-4 week old *N. benthamiana* leaves using a needleless 1 mL syringe. After 2 days the substrate(s) were infiltrated (50 – 100 µL) into the underside of the same leaves previously infiltrated with the Agrobacterium strains of choice in a pre-demarcated area. Substrate concentrations were 100 µM or 500 µM (tryptamines + secologanin). Each individual infiltration experiment was tested at least 2x times for candidate screening and 3x for activity characterization, with biological replicates consisting of leaves from different tobacco plants.

At 2 days post-infiltration, 5 mg leaf disks were excised from the sites of substrate injection. Leaf disks were extracted in 400 µL MeOH for 1 h with 300 RPMs. The MeOH supernatant was subsequently filtered through 0.45 µm low-binding hydrophilic PTFE spin-filter plates (Millipore). Filtered samples were directly analyzed by high-resolution LC-MS and individual metabolites were identified based on comparison of retention times and MS2 spectra with authentic standards. DataAnalysis Version 5.3 (Bruker) was used to analyze LC-MS data.

**Reduction of Kratom iminiums using $NaBH_4$**

Kratom young leaf disks (5 mg) were extracted with 500 µL MeOH for 16 h at r.t. The extract was then divided into two equal portions, with 10 mM $NaBH_4$ (dissolved in MeOH) added to one. Both extract portions were incubated at 40 °C for 1 h, then quenched via addition of 20 mM HCl. The extracts were subsequently filtered and analyzed via HPLC-MS.

Reduction of isolated iminium standards was performed as follows: 10 mM $NaBH_4$ or $H_2O$ was added to 100 µM 20*S*-3-dehydrocorynantheidine or 500 µM 3-dehydromitragynine and incubated 1 h at 40 °C, then quenched via addition of 20 mM HCl. The reactions were then filtered and analyzed via HPLC-MS.

**Isolation of 20*S*-3-dehydrocorynantheidine (13a) and dehydromitragynine (14a) from Kratom leaves**

Young leaves mixed with some mature leaves (62.2 g) were harvested from multiple Kratom plants. The leaves were blended in 500 mL of MeOH, and the resultant slurry filtered and washed with additional MeOH. The solution was evaporated down to ~50 mL, turning an orange color as the chlorophyll precipitated. This mixture was diluted 1:4 in $H_2O$ and run through a 10 g C18 SPE column. The column was washed with 10% MeOH until flow-through became clear (originally pink-ish). Alkaloids were eluted with 50% MeOH, resulting in an orange eluent. The MeOH was evaporated and the resulting aqueous solution was dried via lyophilization.

This powder was dissolved in 5 mL of 25% MeOH and injected onto a preparative HPLC for compound isolation with 300 µL injections. Peaks corresponding to 20*S*-3-dehydrocorynantheidine (DHC, **13a**) and dehydromitragynine (DHM, **14a**) were identified and fractions pooled. These solutions were then dried to obtain 12 mg of crude DHC (**13a**) and 62 mg of crude DHM (**14a**). Since the compounds were not pure, some of these crude mixtures were re-purified using semi-preparative HPLC, resulting in 0.63 mg DHC (**13a**) and 2.5 mg of DHM (**14a**). The identities and structures were verified using NMR (DHM: Figure S56; DHC (Figures S57-S61).

**Isolation of alkaloid standards from Rifat Kratom tissue**

A mixture of dried young and mature leaves (5 g) from Rifat Kratom were blended in 200 mL of MeOH, and the resultant slurry was filtered and washed with additional MeOH. The solution was evaporated down to ~5 mL. This mixture was diluted 1:4 in $H_2O$ and run through a 1 g C18 SPE column. The column was washed with 10% MeOH until flow-through became clear and then alkaloids were eluted with 50% MeOH, resulting in an orange flow-through. The MeOH was evaporated and the resulting aqueous solution was dried via lyophilization. The resulting solution was resuspended in 10% MeOH and compounds purified using semi-preparative HPLC. Fractions containing compounds of interest were pooled and dried via lyophilization. The following isolated compounds were obtained: **3a**: 0.11 mg, **3b**: 0.09 mg, **6a**: 0.17 mg, **6b**: 0.12 mg, **5b**: 0.31 mg, **7a**: 0.10 mg, **7b**: 0.08 mg, **7c**: 0.12 mg.

**Liquid Chromatograph – Mass Spectrometry (LC-MS) data acquisition**

**Compound identification**

For LC-MS data acquisition an UltiMate 3000 ultrahigh performance liquid chromatography system (UHPLC; *Thermo Fischer*) connected to an Impact II UHR-Q-ToF (Ultra-High Resolution Quadrupole-Time-of-Flight) mass spectrometer (*Bruker*) was used. Compound separation was achieved using reverse-phase liquid chromatography on a Phenomenex Kinetex XB-C18 (100 x 2.1 mm, 2.6 µm; 100 Å) column operated at 40 °C. Mobile phases: (A) water with 0.1 % formic acid; (B) acetonitrile; flow rate = 0.6 ml/min. 2 µL sample was injected in each run; authentic standards were prepared as methanol solutions in concentration ranges between 20-100 µM. Chromatography conditions: 10% B from 0 – 1 min, 10 – 30% B from 1 – 6 min, 30 – 100% B from 6 – 6.1 min, 100% B from 6.1 – 7.5 min, 100 – 10% B from 7.5 – 7.6 min, and 10% B from 7.5 – 10 min. Mass spectrometry conditions: mass spectrometry was performed in positive electrospray ionization mode (capillary voltage = 3500 V; end plate offset = 500 V; nebulizer pressure = 2.5 bar; drying gas: nitrogen at 250 °C and 11 L/min). Mass spectrometry data was recorded at 12 Hz ranging from 80 to 1000 m/z using data dependent MS2 and an active exclusion window of 0.2 min. Tandem mass spectrometry settings: fragmentation was triggered on an absolute threshold of 400 and restricted to a total cycle time range of 0.5 s; collision energy was deployed in a stepping option model (20- 50 eV). To calibrate MS spectrum recording each run was initiated with the direct source infusion of a sodium formate-isopropanol calibration solution (operated by an external syringe pump at

0.18 ml/min using a 5 mL syringe with an ID of 10.3 mm). The initial 1 min of the chromatographic gradient was directed to waste.

**Compound purification using preparative HPLC**

For preparative HPLC, an Agilent 1260 Infinity II system was used, equipped with a Phenomenex Kinetex XB-C18 column (250 x 10 mm, 5 µm; 100 Å) and coupled to a multiple wavelength detector and fraction collector. Mobile phases A (water + 0.1 % formic acid) and B (acetonitrile) were used. The flow rate was set to 9 mL/min with the following gradient: 10 – 30% B from 0 – 12 min, 30 – 40% B from 12 – 18 min, 40 – 95% B from 18 – 20 min, 95% B from 20 – 24 min, 95 – 10 % B from 24 – 27 min, and 10% B from 27 – 30 min. Samples were prepared in MeOH as concentrated solutions filtered through a 0.22 µm PTFE syringe filter and injected successively (injection volume: 300 µL). All fractions were assessed by UPLC-MS and fractions containing the desired product were concentrated *in vacuo* and dried using lyophilization.

**Compound purification using semi-preparative HPLC**

For semi-preparative HPLC, an Agilent 1260 Infinity II was used, equipped with a Phenomenex LC column (Luna® 5 µm C18 (2) 100A, 250 x 30 mm, AXIA™ Packed, Ea) and coupled to a multiple wavelength detector and fraction collector. As mobile phases A (water + 0.1 % formic acid) and B (acetonitrile) were used. The flow rate was set to 9 mL/min with the following gradients:

**DHC (13a)/DHM (14a) purification**:

20% B from 0 – 6 min, 20 – 30% B from 6 – 14 min, 30 – 100% B from 14 – 16 min, 100% B from 16 – 18 min, 100 – 20 % B from 18 – 19 min, and 20% B from 19 – 22 min.

**Rifat Kratom alkaloid purification**:

20% B from 0 – 6 min, 20 – 30% B from 6 – 14 min, 30 – 35% B from 14 – 16 min, 35 – 100% B from 16 – 18 min, 100% B from 18 – 20 min, 100 – 20 % B from 20 – 22 min, and 20% B from 22 – 24 min.

Samples were prepared in MeOH as concentrated solutions filtered through a 0.22 µm PTFE syringe filter and injected successively (injection volume: 20 µL). All fractions were assessed by UPLC-MS and fractions containing the desired product were concentrated *in vacuo* and dried using lyophilization.

**Purification of *Ms*CO1 from *Nicotiana benthamiana* leaves**

For the purification of *Ms*CO1, expression in *Escherichia coli* or *Saccharomyces cerevisiae* failed to yield active enzyme. Therefore, the procedure from Caputi et al. 2018[9] was adapted for expression and purification from *N.benthamiana*. Using the previously mentioned modified 3Ω1 plasmid backbone, a C-terminal $His_6$-tag (LEKHHHHHH) was appended to *Ms*CO1. Since native enzymes present in *N. benthamiana* are able to catalyze corynantheidine oxidation (albeit with low efficiency), non-*Ms*CO1 transformed tissue was also investigated as a control. Using our standard agroinfiltration/expression strategy, 25 *N. benthamiana* plants were infiltrated with an *Agrobacterium tumefaciens* GV3101 strain containing this plasmid and another 25 *N.benthamiana* plants were infiltrated with an *A. tumefaciens* GV3101 strain containing an empty vector (EV) plasmid. After four days post-infiltration, infected leaves were harvested, comprising 21.2 g (*Ms*CO1) and 22.5 g (EV). These leaves were homogenized in 100 mL 50 mM Tris-HCl buffer pH = 7.5 containing EDTA-free protease inhibitors and 1% insoluble polyvinylpolypyrrolidone (PVPP) using a blender. The homogenates were then filtered, washed with additional buffer, and centrifuged at 4,000 *g* for 10 min to pellet the insoluble PVPP and tissue debris. The supernatants were further clarified by centrifugation at 35,000 *g* for 20 min. *Ms*CO1 and the EV lysate were treated identically, with both lysates poured over a 1 mL Ni-NTA column, washed with 20 mL 50 mM Tris-HCl buffer pH = 7.5 buffer containing 20 mM imidazole, and eluted with 2.5 mL of 50 mM Tris-HCl buffer pH = 7.5 buffer containing 250 mM imidazole. The eluted fractions were spin-concentrated/buffer-exchanged using centrifuge filtration with a 30 kDa cutoff (Amicon, *Millipore*) into 50 mM Tris-HCl buffer pH = 7.5. *Ms*CO1 and the EV control were then snap-frozen at –70 ˚C. Protein concentration was estimated using absorbance at 280 nm (no protein was detected for EV control).

**Purification of *Ms*DCR from *Escherichia coli***

The coding sequence for *Ms*DCR was cloned into pOPINF and transformed into BL21 (DE3) *E. coli* cells for expression. This strain was grown for 16 h in 2 mL LB + 100 mg/mL carbenicillin (CARB) at 37 ˚C. 1 mL of this saturated culture was used to inoculate 100 mL LB + CARB, which was grown for a further 3.5 h at 37 ˚C until $OD_{600}$ = 0.6. The culture was moved to 18 ˚C for 1 h, after which expression was induced via addition of 20 µL 1 M isopropyl ß-D-1-thiogalactopyranoside (IPTG, final concentration = 200 µM). Expression then continued at 18 ˚C for another 16 h.

Cells were harvested via centrifugation, and the supernatant removed. A cell pellet of 2 g of was obtained, and was frozen at –20 ˚C for 1 h. Upon thawing, cells were resuspended in 10 mL 50 mM Tris-HCl buffer pH = 7.5 + 1 mg/mL lysozyme, 0.1 mg/mL DNAse, and 1 mM $MgCl_2$ and incubated with shaking at 37 ˚C for 1 h. The lysed cells were moved to ice for 30 min and then sonicated on ice for 5 min (1 s on; 1 s off). The lysate was then clarified by centrifugation at 35,000 *g* for 15 min. The resulting supernatant was then poured over a 1 mL Ni-NTA column, washed with 20 mL 50 mM Tris-HCl buffer pH = 7.5 buffer containing 20 mM imidazole, and eluted with 2.5 mL of 50 mM Tris-HCl buffer pH = 7.5 buffer containing 250 mM imidazole. The eluted protein was spin-concentrated/buffer-exchanged using centrifuge filtration with a 10 kDa cutoff (Amicon, *Millipore*) into 50 mM Tris-HCl buffer pH = 7.5. The protein was then snap-frozen at –70 ˚C. Protein concentration was estimated using absorbance at 280 nm.

##### *In vitro* reactions

*Ms*CO1, *Ms*DCR, and the EV control for *Ms*CO1 were thawed from –70 ˚C, and centrifuged for 5 min at 10,000 g to pellet any aggregated enzyme. The supernatants were then removed and used for the following assays: 25 µL total volume containing 100 µM substrate (20*S*-corynantheidine (**3a**), ajmalicine (**10b**), or DHC (**13a**)), 1 mM NADPH, 0.5 µM *Ms*CO1/EV control and/or 5 µM *Ms*DCR, and 50 mM Tris-HCl pH = 7.5. Reactions were allowed to proceed at 25 ˚C for 16 h at ˚C prior to quenching via 20:1 addition of MeOH. Solutions were then filtered, and analyzed via LC-MS.

##### Purification of *Rgn*TDC variants and *Pf*TrpB$^{2B9}$ H275E from *Escherichia coli*

The coding sequences for *Rgn*TDC variants (wt, L355A, L355M) and *Pf*TrpB$^{2B9}$ H275E[10] were cloned into pET28b and transformed into BL21 (DE3) *E. coli* cells for expression. This strain was grown for 16 h in 2 mL LB + 100 mg/mL kanamycin (KAN) at 37 ˚C. 1 mL of this saturated culture was used to inoculate 100 mL 2x yeast extract broth (YEB) + KAN, which was grown for a further 3.5 h at 37 ˚C until $OD_{600}$ > 1.0. The culture was moved to 18 ˚C for 1 h, after which expression was induced via addition of 100 µL 1 M isopropyl ß-D-1-thiogalactopyranoside (IPTG, final concentration = 1 mM). Expression then continued at 18 ˚C for another 16 h.

Cells were harvested via centrifugation, and the supernatant removed. Cell pellets were frozen at –20 ˚C for 1 h. Upon thawing, cells were resuspended in 10 mL 50 mM sodium phosphate buffer pH = 8.0 + 1 mg/mL lysozyme, 0.1 mg/mL DNAse, 1 mM $MgCl_2$, and 400 µM PLP and incubated with shaking at 37 ˚C for 1 h. The lysed cells were moved to ice for 30 min and then sonicated on ice for 5 min (1 s on; 1 s off). The lysate was then clarified by centrifugation at 35,000 *g* for 15 min. The resulting supernatant was then poured over a 2 mL Ni-NTA column, washed with 20 mL 50 mM sodium phosphate buffer pH = 8.0 containing 20 mM imidazole, and eluted with 4 mL of 50 mM sodium phosphate buffer pH = 8.0 buffer containing 250 mM imidazole. The eluted protein was spin-concentrated/buffer-exchanged using centrifuge filtration with a 10 kDa cutoff (Amicon, *Millipore*) into 50 mM 50 mM sodium phosphate buffer pH = 8.0. The protein was then snap-frozen at –70 ˚C. Protein concentration was estimated using a Bradford assay.

#### Coding Sequences used for this study

***Cr*STR:**
atggcacaccatcaccaccatcacagcagcggtctggaagttctgtttcagggcccgtcgcctattctgaaaaagattttcattgagtctcctagttacgctc
ctaacgcgtttactttcgacagcactgataaagggttctatacaagcgttcaagacggacgtgtgatcaaatatgagggaccgaactctggttttacagattt
cgcttatgcaagtccattctggaacaaggctttctgtgaaaacagtactgacccagaaaaacgccctttgtgtgggcgtacgtacgacatttcatacgattat
aagaattctcagatgtacatcgtcgatggtcactaccatctttgtgtggtaggaaaggagggaggttatgctactcaactggcaacgtctgtccaaggtgta
ccgttcaagtggttatatgcagtcaccgttgatcaacgcactgggatcgtgtactttacggacgtctcgtctatccatgacgactcacctgagggggtcgag
gagattatgaatacgtctgatcgtacgggccgtcttatgaagtacgacccatctactaaggagactacgctgttacttaaagagttacacgttccgggcggg
gctgagatttcggcagatgggagctttgtggtagtagcggagtttctttctaaccgcatcgtcaaatactggttagaaggtcccaaaaaggggagtgcaga
gttcttggtgacgattccgaatcccggtaacattaagcgtaactcagacgggcatttttgggtgtcttcaagcgaggaactggatggaggacagcatggcc
gtgtggttagtcgcgggatcaaattcgacggttttggaaatatcttgcaagtcatccctttgcctcctccttatgaaggcgaacactttgagcagatccaaga
acatgacgggttactgtatattggcagcctgttccattcatcagtcggaatcttagtctacgatgaccatgacaacaaggggaactcttacgtgagcagcta
a

***Ms*STR1:**
atgctatcgaaaatcacacctaacatacacacttctgaaagtatggttgcgttaaccattttctttaccctgttcttgttccctctttcagttgttctatcttctgcgg
aatttttccagttcctcaagtcaccctacggccccaacgccttcgcttttaactccgctggtgaactctacgctgccgtcgaagatggcagaattgtcaagta
caaaggatcaagcaatcacgggttttcgacccacgctgttgcctccccattctggaacagaaaagtttgtgagaattataccgaacttcagctgaaaccctttt
tgtgggaggacatatgaccttggatttcaccatgaaactcggcagttgtacattgctgattgctattacggtcttggggtagttggacctgaaggaggccgt
gccactcaagttgccaggagtgcagatggagtggacttcaagtggctctatgccttggccgtggaccaacaaactggctttgtttacctcactgatgttagc
acaaaatatgatgacagaggtgttcaagacatcatgaggataaatgatacaacaggaagattaattaaatatgatccctcaactaatgaagctagagttttga
tgaatgggctgaatgtaccaggtggcaccgaagttagcaaagatggctcatttcttgttgtcgctgaattcttgagccacagaattctcaagtattggttaaag
ggtcctaaggcaaatacttctgaggtattattgaaagtgaggggcccaggaaacataaaaaggaccaaagctggtgaattttgggtggcctctagtgaca
ataatggaattactgttacgcctagagctataaagtttgacgactttggcaacattttacaagtcgtgcctgtccctccaccatataaaggtgaacatttcgaac
aggctcaagagcataatggtgctctttacatcgggacactgttccacgactttgtgggcatattacacaataacgaaggttcatctgaacccaaggaaaata
atgcacatggggtcagtggatctttgaatggggtggcttcttctgtctga

***Cr*SGD:**
atgaacacttctgaaagtatggttgcgttaaccattttctttgctctgttcttgtcccctctttcagttgttttatcttctgcggaatttttccagttcctcaagtcaccc
tacggccccaacgccttcgcttttaactcagctggtgaactctacgctgccgtcgaagacggcagaattgtcaagtacaaaggatcaagcaatcacgggtt
ttccacccacgctgttgcctccccattctggaacagaaaagtttgtgagaattataccgaacttcagctgaaaccgttttgtgggaggacatatgaccttgga
tttcactatgaaactcagcagttgtacattgctgattgctattacggtcttggggtggttggacctgaaggaggccgtgccactcaagttgccaggagtgca
gatggagtggacttcaagtggctctatgccttggccgtggaccagcaaactggctttgtttacctcactgatgttagcataaaatatgatgacagaggtgttc
aagacatcctgaggataaatgatacaacaggcagattaattaaatatgatccgtcaactaatgaagctagagttttgatgaatgggttgaatgtaccaggtg
gtaccgaagttagcaaagatggctcatttcttgttgttgctgaattcttgagccacagaatactcaagtattggttaaagggtcctaaagcaaatacttctgag
gtattattgaaagtgaggggcccaggtaacataaaaaggaccaaagctggtgaattttgggtggcctctagtgacaataatggaattactgttacgcctaga
gctattaagttcgacgactttggcaacattttacaagtcgtgcctgtccctccaccatacaaaggtgaacatttcgaacaagctcaagagcataatggttctct
ttacatcgggacactgttccacgactttgtgggtatattacacaactacgagggttcatctgatcccaaagaaaataatgtagatggggtcgatggatctttg
aatggagtggcttcttctgtctga

***Ms*SGD:**
atggaagctcaaagaactgccactgttgtttccaacgatgcaagcaagatcaaccgcggtgattttgctgaggatttcatttttggagcagcttcatctgctta
tcagacggaaggcggtgcaagtgaaggtggtcgaggtcctagcatatgggacactttcacccagagacgaccaggtatgataaaggagggcggcaat
ggaaatatggctgttgattcatatcatcagtataaggaagatgtcaagattttgaagaacatggggctagatgcctatcggttctcaatatcatggtcgagagt
actgccaggtgggaatttaaatgctggcgtaaataaggaaggaatcaactattacaacaatctcattgatgagctcctagccaatggtatcgagccatatgt
aactctgtttcactgggatgttccccaagcattggaagataaatatggtggctttttaagtcctcaaattgtggacgacttccgcgagtacgtagagctttgctt
ttgggagtttggagatcgagtgaaacactggataacactgaatgaaccatggagctttagtgttggtggatatgtaaacggcacgtttgcaccgggccgag
gtgcctcttcatcagatcaagaaaacgaccatccagctgcactaccgagcagatgttctccatggcaatcacaaagtatttctagcaatggaaatccaggg
acagagccatatgtggtgactcacaatcagcttcttgctcatgcagctgctgtcgaattgtataagagcaactttcagaaatcacaaaatggcaagattggg

attacacttgtgtctcagtggatggaacctttggacgaaaacagtaaagctgatgtcgaagccgcaaagagagctcttgatttcatgcttggatggtttatgg
agcctttaacgaccggtgattatcccaaaactatgagaaaattagttggatctcgtctcccaaaattttcagccgagcaatctaagcaactcaagggatcata
tgattttcttggattaaattattacactgctgactatgtcacaagtgcatcaagctccactactggaggaaatttgagttacactacggattctcaagtgacccat
acaactgatcgaaatggagtgccaattggtccacagggtggctcagaatggttgcatatttatccagaagggattcgcaaactattggtttacgtaaagaag
acatacaatgttccgctcatttacataacagagaatggagttgatgaagtgaatgatacaagcttaacactttctgaagctcgagttgataacaccagaataa
agtatatccaagaccaccttttaaatattcgactagcaatcagtgatggagtaaatgttaagggctacttcgtttggtcattgttggataatttcgagtggagcg
aaggatacactgttcgttttgggtttattcacattgattacacaaacaattttgcaagatacccaaaagactcagccatatggtttatgaattcttttcacaagga
gtaccctaaaaaattcctcaagagaactctggaagatcacgaagatttcgtttcgaagaaaaggttgcgccagtag

***Ms*DCS1:**
atggcaggaaaatgtgcccaagaagagcacacagtgaaggcttttggatgggccgctagagaagcctccggcgctctatctccttacgggttctcaaga
agggcaacaggagagcgtgatgttcgggttaaaattttgtattgtggaatctgtagaacagacgcagaaatgatcagcgacaaattttgctttactaagtatc
ctcatgtgcctgggcatgagatcgtgggtgtggtatctgaagttggtaacaaggtgcaaaaattcaaggttggagctaaagtcggtgtgacaggcataatt
ggatgttgtcgaacttgttatagctgtaccaatggtcttgagagttactgcccaaatgttgcactaacagaagcaggtgaaggtggttgctctaacttcatagt
tttggatgaagactttgtgtttcgttggcctgagaaattacctcttgatcttggagctcctctcctgtgtgctggagccgcttcttacagccctttgaaaaattttg
gacttgataaacctggattgcatattggtatagctggtcttggtggcatgggccatgtagctgtaaaatttgctaaggcttttggggcaaaggtgacagtaatt
agtacatcagataacaaaaaggaggaagccattaaaaaatatggtgcagacgcatttttgaatagtagtaatcctgagcagatgcgggctgcagctggta
cactggctgccatcgttgatactatcccttcgcctcactctctagtgccattgctcgatttattgttgcctcatgggaaggttattgtattaggggcacccagtg
agccatttgtgttgccggttatgcccctgcttcaaggtggaagagtagtcgctgggagttccggtgcaagtttgaagcaaatccaagaaatgctcgattttg
ctgcagaacacaacatagtagctgatgctgaggttatcccaattgactatataaacactgcaataaagcgcattgagaagggcgatatcaaataccgatttg
tcgttgacatcgggaatacactgaaatcggcttaa

***Ms*DCS2:**
atggcagagaagagtcccgaggaagaacatccggttaaagcattcgggctcgcggccaaagatagctcaggcatcttatctccgtttaattttagtcgtcg
cgctacgggcgatcacgacgttcaactgcgtgttctgtactgcggattgtgctactacgacactaagatgattaagaataagagaggcgtgaccagatacc
cgtttgtattcggacacgaaatcgttggagaggttaccgaaataggccgtgaggttcagaaatttaaggtaggtgacaaggttggagtcggttgtatggttg
cttcctgccgtagttgcgagtcttgcgcgaataactgcgagaattattgtccgaacgtatccgtgacggacggtgcgttcttctttaaaacgggtgaggtgtt
gtacggcggctgctccgatattatggtcgcggacgagaacttcgttattcggtggccagagaatttcccacttgacgcaggagcgccattactgtgcgca
ggcattacaacgtattcgccattacgcaactttggcctggacaagccgggcatccacgtgggcatctacggactcggtggtctcggtcatgttgcggttca
gttcgcgaaagccttcggcgcgaaggttacagtcatatcttctagtgaccggaagcggatggaagcgatcgagaagctgggcgctgatagcttcctggt
gaatagcaacttagaagagatgcaggcagctatggggaccatgcacggaattatcgacaccgttccggcgaaccacagtttagtcccgttactggacct
gttaaaaccacagggtaaactgatagtggttggtggtcctgagaagccgttcgagctgccagtattcccgttactgcagggtggccgtctggttgctggaa
gcgccacaggtggcattaaacagacgcaggagatgatagacttcgccgcggaacacaatattctgccccacgtcgaggtggttagcgtcgactacgtca
atacggcaattgaacgaacggaacgtggagacgttaagtaccgttttgtaatcgatattggcaacaccctgtactga

***Cp*DCS:**
atggccggaaaatctcaagaagatgggcagacggtaaaggctctaggatgggccgctagggaagtttctggggcgatctctcctttcgatttctcaagaa
gggccccaggagagcgcgatgtgcaggttaaaatactatattgtggaatctgtagttttgacacagaaatgatcaataacaagtttggctttaccagatatcc
ctttgtactcgggcatgagattgtgggagtggtatctgaagttggtagaaaggtgcaaaaattcaagattggggataaagttggtgtaggaaccatgattgg
atcttgtcgcacttgttatagctgcactcacaatctcgaaaattactgcccaaaagttacattaacagaagcaacttctggtggttgttctaatcttgtgatagca
gatgaagactttgtgttccattggccggtgaatttgcctcttgatcttggagctcctctcctttgtgctgggattactgtttatagccctttgaaaaattttgaactt
gataagcctggattgcgtattggtgtggttggtcttggtggtattggccatatagctgtaaaatttgccaaggcttttggggctaaggtgacagtgattagttca
tcagaaagtaaaaaggttgaagccattgaaaaatatggtgcagattccttttggttagcagtgatccagggcagatgctggcagctgccggaaccttgga
tggtgtcattgataccgtcccagcacctcactctattttgccattccttgatttactcttgcctcgtggaaagctaattatattaggtgcaccaatggagccatttg
tactgccaatctatcccctgcttcaaggtgggagagtagttgctgggagtgccactggaggattgaaacaaatccaagaaatgcttcattttgcagcagag
cacaacatagtagcagatggcgaggttatcccaatcgacgacattaacactgcgataaagcgcattgagaaaggcgatgtcaaatatcgatttgtggttga
cattggcaataccttaaaatctgcttga

***Ms*EnolMT:**
atgcaaccacagagagggagaaagagagagagagagagagagatagagaagagatggaatccgtgcagagcaacagtagttcttctgatcaattcgcaa
tgaaaggtggagatgacgacttcagttacacaaagaattccacctggcagagagatgcaattcaagcaaccaaatttttcattcaagaatctattgctgaga
agcttgacgtcaataaattttgtggaaaggcattttgcgttgctgatttgggatgctcagttggacctaacactttgatagcaatgcagaacattgttgaagctg
tggagcttaaattcaaaaatagaaaaggattccattctcccactatccctgaatttcaagtcttctttaacgatcatacggtgaatgatttcaataccctctttaga
tctctcccaactggtcacgacaagcgctattacggcgttggggttccgggttccttttacggtcgattatttccttgtgactctattcacataatgcacacttcatt
ttctacaccgtttctttctcaagtaccaaaagaggtgattgacaaaaattcagctgcgtggaataaaggaaggattcatcacaattatgctaaagcagatgttt
tgaaggcttatgaagcacaacatgctgaggatatcgactgctttttgacggctagagctaaagaactggtccatggaggattattgatggatgtgacttcatt
ccgcccagatggggtccctcatacccatgtcttgactaacatagggatggaggtattgggttattgcctcatggacttggctggacttatcgatgaagaaaa
cgtggattcctacaacgttccagtttatcttcaatctcctgaagagttgaaacaagctgttcaacggaacaaatacttcagtatagaaaaaatggagagcgtg
cctatgatgatagattcagatgtttctgccaaagctcaacaatattcattgggaatgagggccgtaatgggggacgtgattagagagcaatttggagcgga
gatagtggataaactctttgatttgttcaagaagaaacttgaagagcatcctaactttgcaaaaggagttgtccttgacatgtttgttctccttaaacgcaatgc
agaggattga

***Ms*CO1:**
atgatcacaaaaatctgcaagattctattgctgatttcaattttctacttagtgatcccatcatcacattcatgcttgattcctcatagtttcatccattgcatttcacg
tacgtttccatcaaacgcttctatacttaatgtcctgtatctccctaacaattcttcttatccatatttattaaagtctaccattcacaatcttagattcttgacgccta
ccacccctaccccgcttgcaatagtcactcctttagaatactctcatgtccaagccactgttaaatgtagcaagctgaacggattaaacatcagaatccgaa
gtggtggccatgactatgaaggcatgtcatatacatctgaagttccgtttgtgatgcttgatctcagaaacctaacgtctatcagcattgacattaagactaata
gtgcatgggttgagactggtgcaacgttaggcgaattgtattattcaattgctaagacaagtcctattcatggctttccagcaggcctttgtccaactgttggtg
ttggtggacactttagtggtggcggtgtaggtaacctgatcagaaagtatggactagctgctgataatgtcatcaatgcgcgcattgttgatgttaatggtcg
aattttagatagaaaatcaatgggagctgatctttttgggctattagaggaggtggaggagcaagttttggagttatagttgcctggaaaatcaagcttgtgc
gtgttccacctgtagttactgttttcaatttaaccaagagttcaagtcaagaagccataggtcttattcacaaatggcaatatgcagcgcccaagttgagtgac
gatttgattgtttacattacaatatcatcgataaatgagaagggaggaggacttaccgcaacatttaattcattgttcctcggtagatctggtcagctcttgaaa
atgttggaggagagcttccccgaacttaacctcagaaaagaagattgtgttgagatgagttggattgagtcagtactccattttgcagcatatcaaaatgtgg
aaactatagaggccctaaagaaaagaattaactcgcaacctgataattacttcaaggctaagtcagacttggttcgcaagcctctaccatatgaagcattag
aagagttctggaaatggggttcagatatgaatgctccttacttttatgcagaattgcgtccttatggtggaagaatgaatgagatatcggaatcagaaactcc
atatccacacaggaaaaatgttctctatgaaattctctacgtggtgtcttggacgaaggataaagatgatggatcttccaaaaagaacatcaattggctaaga
ggattatacgagttcatgaccccttacgtgtcaaaagggccaagaggtgctgtttggaattgtagagatcttgatttaggtgctaatggtgcttcagaaacta
cttattctaaagccaaggcatggggatcaaggtatttcaggaacaattttaagaggctggcggttattaaaggtgaagttgatccaaataatttttcaactatg
agcaaagcattccccctcttgttttccatggacaaagtgtggaagagcagatgtgtgagttttcttga

***Ms*CO1-C-His$_6$:**
atgatcacaaaaatctgcaagattctattgctgatttcaattttctacttagtgatcccatcatcacattcatgcttgattcctcatagtttcatccattgcatttcacg
tacgtttccatcaaacgcttctatacttaatgtcctgtatctccctaacaattcttcttatccatatttattaaagtctaccattcacaatcttagattcttgacgccta
ccacccctaccccgcttgcaatagtcactcctttagaatactctcatgtccaagccactgttaaatgtagcaagctgaacggattaaacatcagaatccgaa
gtggtggccatgactatgaaggcatgtcatatacatctgaagttccgtttgtgatgcttgatctcagaaacctaacgtctatcagcattgacattaagactaata
gtgcatgggttgagactggtgcaacgttaggcgaattgtattattcaattgctaagacaagtcctattcatggctttccagcaggcctttgtccaactgttggtg
ttggtggacactttagtggtggcggtgtaggtaacctgatcagaaagtatggactagctgctgataatgtcatcaatgcgcgcattgttgatgttaatggtcg
aattttagatagaaaatcaatgggagctgatctttttgggctattagaggaggtggaggagcaagttttggagttatagttgcctggaaaatcaagcttgtgc
gtgttccacctgtagttactgttttcaatttaaccaagagttcaagtcaagaagccataggtcttattcacaaatggcaatatgcagcgcccaagttgagtgac
gatttgattgtttacattacaatatcatcgataaatgagaagggaggaggacttaccgcaacatttaattcattgttcctcggtagatctggtcagctcttgaaa
atgttggaggagagcttccccgaacttaacctcagaaaagaagattgtgttgagatgagttggattgagtcagtactccattttgcagcatatcaaaatgtgg
aaactatagaggccctaaagaaaagaattaactcgcaacctgataattacttcaaggctaagtcagacttggttcgcaagcctctaccatatgaagcattag
aagagttctggaaatggggttcagatatgaatgctccttacttttatgcagaattgcgtccttatggtggaagaatgaatgagatatcggaatcagaaactcc
atatccacacaggaaaaatgttctctatgaaattctctacgtggtgtcttggacgaaggataaagatgatggatcttccaaaaagaacatcaattggctaaga
ggattatacgagttcatgaccccttacgtgtcaaaagggccaagaggtgctgtttggaattgtagagatcttgatttaggtgctaatggtgcttcagaaacta
cttattctaaagccaaggcatggggatcaaggtatttcaggaacaattttaagaggctggcggttattaaaggtgaagttgatccaaataatttttcaactatg
agcaaagcattccccctcttgttttccatggacaaagtgtggaagagcagatgtgtgagttttcttgactcgagaaacaccaccaccaccaccac

***Ms*CO2:**
atgctcacaaaaatgtgtaaggtactattgctgatttcaattttctacttagtgatcccatcatcacattcaagcttgattcctcatagtttcatccattgcatttcag
gtacttttccatcaaacacttctatacttgatgtcctgtatctccctaacaattcttcttatccatatttattaaagtctaccattcacaatcttagattcttgacgccta
ccacccctaccccgcttgcaatagtcactcctttagaatactctcatgtccaagccactgttaaatgtagcaagctgaacggattaaacatcagaatccgaa
gtggtggccatgactatgaaggcatgtcatatacatccgaagttccgtttgtgatgcttgatctcagaaacctaacgtctatcagcattgacattaagactaat
agtgcatgggttgagactggtgcaacgttaggcgaattgtattattcaattgccaagacaagtcctattcatggctttccagcaggcctttgtccaactgttgg
tgttggtggacactttagcggtggcggcgtaggtaacctgatcagaaagtatggactagctgctgataatgtcatcaatgcgcgcattgttgatgttaatggt
cgaattttagatagagaaaatcaatgggagctgatctttttgggctattagaggaggtggaggagcaagtttcggagttatagttgcctggaaaatcaagcttg
tgcgtgttccacctgtagttactgttttcaatttaaccaagagttcaagtcaagaagccataggtcttattcacaaatggcaatatgcagcgcacaagctgagt
aacgatttgatggtttacattgcaatatcatccataaatgagaagggaggaggaattgccgcaacatttaattcattgttcctcggtagatctgatcaactcttg
aaaatgatggaggagagcttccctgaacttgacctgagaaaagaagattgtattgagatgagttggatcgagtcagtactccattttgcagcatatcaaaat
gtggaaactatagaggccctaaagaaaagaattaactcgctacctgatagttacttcaaggctaagtcagacttggttcacaagcctataccatatgaagca
ttagaagaattctggaaaaggtgttcagatatgaaagctccttacttttatgcagaattgcgtccttatggtggaagaatgaatgagacatcggaatcagaaa
ctccatatccacacaggaaaaatgtcctctacgaaattctctacatggtgttttggacgaaggataaagatgatggatcttcaaaaaagaacatcaattggct
aagaagattatacgagttcatgactccttatgtatcaaaagggccaagagctgctgttttgaattgcagagatcttgatttaggtgcaaatggtgcttcagaaa
ctacttattctaaagccaagacatggggatcaaggtatttcaggaacaattttaagaggttggcggttattaaaggtgaagttgatccaaataatttttcaact
atgagcaaagcattccacctctggttttccatggacgaagtgtcgaagagcaaatgtga

***Ms*DCR:**
atgggaagcaaaagcaagattttgataattgggggcacagggtacattggaaaattcgtagttgaagccagtgtaaaagaagggcacccaacttttgcatt
ggttagagaaagcgcagtctcagatcctaaaaaggctgcaattgtagaaagcttcaagagctcaggagtcacaattctttatggagatttaaacaatcatca
gcagttggttaatgcaatcaaacaagtggatattgtcatctctgctctgggtggagatttgacagtgatgcctgagcaagtgaagatcattgcggctattaaa
gaagctggaaccatcaaaagatttttacctgctgaattcgagattgatgtggatcgtatgcatgctgttgagcctgctgcaagctatttgaggtcgaaggcga
agattcgcaaaattgttgaggctgaaggaatacctcatacttacttggtatctaacggttttattggttatttgccccatttctttaatctctttgaagtccttagccc
cacaactcttcccagagacaaagttgttgttcttggtgatggaaatccaaaagttgttttcaataaggaagaggacatagctacgtataccatcaaagcagc
agatgacccaaggactctgaacaagagcgtgtacgttagacctcctgccaacactttgtccttcaatgaaataatctcattgtgggaaaagaaaattggcaa
gaccctcgaaaggatttatgttccagaggaagaacatcttaagaaaattcaagaggcttcaatgccattaaacgcgatcctagctatgggatactcggctat
cgtgaagggagatactgcaaactatgagattgtagctacttttggagcggaggcaactgagctttatcctgatgtgaaatataccacaactgacgagttcctt
gaccagtttgtacgagacatatcaagaaat

***Ms*SAS (*Ms*3eCIS):**
atggcaagcttcccagcatttcaggaagaagcgaagagatggctctcgaaccattcagtggcagtctctttgcttcttccttcgttcatgatcttcattctatttc
tttccaaatggtggttcctcactaccagtaagaaaaagaaccttccaccttcacctccgagactgccaataattggaaaccttcatcaaattggttcttttgccc
accgctctttcaaatccttggccaaaaagcacggtccgataatgctacttcacgttggcaagcaaagactgttagttgtctcctccgccgacgctgctcgtg
aggctctcaaaacacatgatgctgtcttttgcgatagacctgattcagaggttacacgaagaattttctataaccataagaacatatctctctccacctatggtg
attattggaagctagtgaggagcatcgctgtgaatcaaattctaaatcagaagagggttcagtcgttcagaagtgtaagagaagaagaggtttcattactgg
tggaaagaatcaaagaatcttgctcttcttcctccgtaatatgcatgaatacgttgttgacaacgcttgtaaatgacatagtgtcaaggataaccattgggaag
aggtcctcaggaaaagatggaagcagatttgcagagtttttgtttgcatccgtcgacttattaggttcttttcttgcaggggacttcttcccagggctttggtgg
ctcggttacataactggattcgaagcaaagatcaagaaagtttccaaagatttggatcaatgtttggagaacttaattgaagaggaaataaacaggaacaaa
agaggagatgaccaaggtaaagacaatcagaatttcctccaggttttgcttgaaatccagagaaccgactcatctggccatgctttggatcgagaatccatt
aaggctgtcataatggacatggttgccggtgcatttgactcgtatacacttttggagtgggcaacgtcattgctagtaaaacatccagatgccatgaaaaaat
tgcaaaatgaggtgaaagaagttgctggatccaaatcattcatatcggaggatgatttaagtaaactgcaatacttgaaagcagtaataaaagaaactttcc
gattctgtcccggtgcaattcttgcaaggataccaatcaaggatgtcaaattaatgggctatgatattgcagcaggcactcaactcctcgtcaatacatgggc
aattggaagggatccaacgttgtgggaaaaacctgaggagtttcggccagaaaggttcttaaatagttcaatagatttgaaaggacatcattttgagctgctt
ccatttggcacaggaagaaggtcttgccccggtatgacatttgccttagtcatggatgagctcgtactagcaaatttggtgtgcaactttaattatgcattgcct
ggtggaacaagagtcgaggacttggacatgagtgaagttcctggaatcacgcctcgtaggagaacccctctgctgctcgttccatctctttgttaa

***Ms*10H:**
atggatgtcctcttgtactttctggccctgattttctctctattcttcattcaaaaccttctcagaaaacagaaagaaagactaccaccaagtccaccatccctac
ctattatcggccatctccacctcctaagaaatccgaaacacagagctctccaacatctctccaacaagtatggccccgtagtttaccttcgtttaggaacccg
tcaaacccttctcgtctcctctccatcaatcatagaggaatgcttcacaaagaatgacataattttcgcaaacagacccgattcacttatcagcaaatacttca

gctatcaaaataatgatcttacattcgccccatatggagatcgctggcgtaatctccgccggctggccaccatccatgtgttctcttcagccaatcttaaacg
gttctccgccattaggacggaagaagtttttgatgctgtgcaagaattgttgtccatctctcatattgaatcccacaaagtaaacctgagatctttgttttccaaa
atggtactcaatgtggctatgaagatgctttctggaaaaaaatttcgtggctctcctgaaatgactgaaatgttttcttcagatatgccgatgggcatttgtgatt
atcttccaatactgggatggcttggttttgggggttttgagaagagcttagtgaaatatcaaaagctgattgatgagttcctgcaagatctgcttgaggagag
ccgcaagattggggaaggttcagttgtgcaaaataaaaccattattcagtcattattgtctctgcaagaagctgaccccgagcgctatcatgatgatgttatc
aaaggaattatgatagatgttttcacagctgggacagatacagttgcatacaccatggaatggatcatgtccctcttactgaaccaccctgaggtgttgcag
aaaataaggagcgaaatagattttcatgtcatcccaggatgtttgattcaggattcagatctccctaaactgtcttatctacgttgtgtgatcaatgagacattaa
gactccttgcccccgtgccccttctgctgccccatttcagctccgaagactgcatagtaaatggattcaatgtgcctcgagggacaactttgttagtagatgtt
tggtccgttcatagagacccaaatgtgtgggaagagccattgaagtttaagcccgaaaggttcgaaggaatcagagtggaagatgaaaaaaagttcaagt
ttattccattcggagtggggagaagggcatgtccgggctcaggaatggctatgagattgatgggattggcattgggatccttgattcagtgctttgaatggg
agagggttgggactgagctagtggacttggaagagggagatgggctagttttgccgaaggttgaacctctagaagcactgtgcaagcctcgcccatcca
tggcccatctcatctctcatatt

**BBE03:**
atggtcatacaattctgtaagattctattgctaacttttattttctacttggtgatcccatcatcagattcaatcttgattcctcatagttttatcgattgcatttcacgtt
cgtttttatcaaacatttctattcttaatgtcctgtatctccccaacaattcttcttatccatttatctttctgtctaccattcacaatcttagattcttgacacataccag
ccccacccctcttgcaataatcactcctttagattactctcatgtccaagctactgttaaatgtagcaaattgaatggattaaatatcagaatccgaagtggtgg
ccatgactatgaaggcatgtcttatacatctgaagttccatttgtcttgcttgacctcaataacctgaggtctattagcattgacattaagaaaaatagtgcatgg
gttgaatctggtgcaacaataggagaattgtattattcgattgctgaaagaagtccaattcatggctttccagcaggcctttgccccactattggcgtcggtgg
acactttagtgggggcggtgtaggtaacctgatcagaaagtatggactagctgctgataatgtcatcaatgcgtgcattgttgatgttaatggccgaattcta
gatagaaaatcaatgggaactgatctttttgggccataagaggaggtggtggagcaagtttcggagttatagttgcctggaaaatcaggcttgtgcgcgtt
ccacctgtagtaactgttttcgaattaactaagagttcgaatcaagaagccataggtcttattcacaaatggcaatatgtagcgcacaagctgagtgaagattt
gctgtttagaatcacaatatcatcgataaatgggaaggaaggaggaattgtcgcaacatttgattcattattcctcggtaaagctggtcaccttttgaaaatga
tggaggagagcctccccgaaattcacctgagaaaagaagattgtattgagatgagttggattgagtccgtcctccattttgcagcgtatcagaaaggggaa
aatacagatgccctaaagaatagaattagcccgctgcctaatgcttatttcaagggaaagtcagacttggttcacaagcctataccttatgaggcattagaa
gagttctggaaatggtgttcgcataaaaattctcctactcttcacatagaattgcatccttatggtggaagaatgaaagaaatatcggaatcagaaattccata
cccacacaggaaagatgtgctctacgaaatcctctacatggtgttatggatgaaggataaagatggtgaatcttcggaaaagaacatcaattggctaagag
gattatatgagttcatgactccttatgtgtcaaaagggccaagaggtgctatttggaatattagggatcttgatttaggtgcaaatggtgtttctaaaactactta
ctctaaagccaaggcatggggatcaaggtatttcaagaataattttaagaggctggcatttattaaaggtgaagttgatccaaataatttttcaactatgagca
aagcattccacctttggttttacatgcaaaa

**BBE04:**
atgagaaagctcagtatcattgtcgcttcattgctctcaaccttgcttatagtctcatttgcaacttctgattctgttcatgaggcatttgttcaatgtctagaacaa
cattcccaaaccaatatttcttcagtaatatacacccctaacaactcttcatttccatctgttttgcaagcttacattagaaacctacgattcaatgagtcctcaac
ccgaaaaccgttccttatcctcactgctttggatgtttctcatatacgggcagccgttatctgtgcaaaagcacatggcttgcagatgaaaatccgaagcgga
ggccatgactacgagggcgtttcctacgtctctgaagtccctttctttatcctagacttgttcaatcttcgatcgatcagtgtgaacatagccgaagagactgc
ttgggttcaggttggagcaacccttggtgaagtatactatagaattgctgagaaaagtaatgtccatggcttccctgcaggtgtttgtcccaccgtgggcgtt
ggtgggcattttggtggaggtggatatggtaacatgatgaggaaatatggtctttctgtggataacatcattgatgcacagatcattgatgggaatggtcgcg
tccttgatcgagcatcaatgggcgaagacttgttctgggccattactggaggtgggggttcgagttatggagttgtccttgcgtacaaaatcaacttagttcg
tgtcccacctcaagtaacagtgttccgggtggagaggacttacgaacaaaatgctacataccttgtacgccgttggcaagaaattgctgacaaattggata
atgatatcttcattaggatgatcattgatgtgattaataatactcgtaccgaagggaaaacaataagatctgcatatttcgcattattcctcggagattcggcaa
ggcttctttctctcatgaaccaaagttttcccgaattgggattgcagcaaaaagattgcattgagatgagttgggctgagtcagttgtctattatacaagctttc
cccttggaactcctgttgatgctcttcttagcagagttcctcaggtaatgactcatcttaaaaggaagtccgattatttgaaaaagcctatgccaatagaaggc
atagaattcatcttcaagaaaatgattgaattgcaaactccagtcttgactttcaatccatatgggggaaggatggctgaaattgcatcctcagcaaaaccctt
tccccatagagctgggaacattgcaaagatccagtatgcaacaaactgggatcaaaatggtgtcgagacggcagaacattacataaatttgacccgagct
ttatacaaatatatgactccttttgtctccaagtttccgagggaagcatttctaaactacagagatcttgacttgggaatcaaccataacggcaaggatagtttc
cttgaaggaactgtttatggaattaagtacttcaaggaaaattttaacagattggtaaaagttaagaccgcggttgatcctgataatttcttcagaaatgaacaa
agcatccccgtgtttccatccaagaaa

**BBE05:**
atggaaaacagctttcagtctccaaattacacgtctcttctacttcttcttcctcttttcagtttgaattttcagccctttgttgctgcctcagattcaatctatgaaaa
ttttgctagttgtctaacaaaaaatgaaatcccaagtgaccaaatcggtaagattctttatagcccatcaaacacttcattcaactctgttttagaagcttatgttc
gaaaccttaggctcaatacttccagtaccagaaaaccatcaataattgtcacacctttggaaatccaacatgttcaagcaacaattttatgcaccaaaggaac
agggctgcaactaaaaatcagaagcggcggtcatgattttgaaggtatctcatatgtttctgatgtcccttttatcattcttgacatgtttaatctgaggtccatc
agtgttaatatcccaagtgaaactgcttgggtacaagccggggcgacactcggggaactttattacagaatttgggaaaagagcaatgtatatggataccc
agctggtgtttgcccaactgtaggcgtcggtgggcatattagtgggggtggttatggtgcaatgttgcgaaaatttggcctcacagttgataatgttcttgatg
cacaaattgttgatgtaaaaggccaggttttggatagaaaagcaatgggggaagatctttttgggctattagaggcggtggtggtgccagttttggtgttgtt
ttggcgtacaaaattaagatagtccaagtgcctcaaacagtaactgtctttcgggttgaaaagactgaggcagagaatgcaacagatactcttgttcagtgg
caaaatgttgcagacaaaattgacaatgatcttttcataagagttcttgtccaaccaatcactgcaaaaagtggcaaaagtaagggtcagaagatcattaggt
taacattcattgcattatttcttggagattcaaataggcttatctctgtcatgaatgctggatttcctaaattggggttgaagaaaacggattgtcaagaaatgag
ttggatagagtccatgctatattgggcaaattttgacaacacgacaaaacctgaagctcttcttagcagacattacgataccaatttcttgaaaagaaaatcag
attatgtccagaccccaattcctaaggatgcactgaattcaatattcgcgaaaatggttcagcttggaaaaacaggttttgttttcaatccttatggaggaaga
atgagtgaaattcctgaaaacgaaacgccatttcctcatcgggctggtatcatttacaagcttcagtattctgtgaattgggatgatgcagacaccaacttag
caaaacaatatattgggcaagcaagggaactctacagtttcatgacccccctacgtatcaaagaatccaaggcaggcttttctcaactatagggatctagata
ttggtataactaataatggaaagaatagttataatgaaggaaaagtttatggactcaagtatttcaagggtaattatgatagattggttaaagtgaagactattg
tagatcctgaaaatttcttcaggaatgagcaaagtatcccacctctgacctccggggcggcctatcgtgggaggaggaag

**BBE07:**
atgaagcttcaaatttcttttgctcttcttctactaagcatttcttgtgcaatttcttcagctcagaccgataactccacagcttcttttctcagctgcatcttcagttc
caatgacacctcaatttcaagcatcgtttacaccccagaaaattcttctttcttatcaatcttggatttctacatacaaaattcacgattcctaaatccagaaacac
ccaagcctaaagtgattctgacaccagttaacgaatcacaaattcatttagccatatcttgcggtaacagttctggtttacaaatgagagtacgaagtggagg
ccatgattttgctggcagttcctatatttctgtagtcccatttttcatcctcgatatgttcaactttcgttcaatttctgtcgatgctgaaaatcgcaccgcatgggtt
ggagctggtgcaacccttggcgaaacatactacagcatttatgaaaagaacaggtctttgggtttcacagccggctattggcctactgttgctattggtggg
cacatcagcggaggaggttatggtgcattgaccaggcaatatggtctagctgctgacaatgtcattgatgctcgggtgataactgctaatggaacatatctt
gatcgagcctccatgggtgaggatctttttgggctattagaggtggggttggttcaaatttcgtagtcattcttgcgttccaactatctttagttgatgttccgga
aaatgttactgcattttctgtcactaggaccctggaacaaaacgcgattcaacttgtctacaagtggcaacatgtagcccccactttacctataaatcttaccat
ttccctccaatttactagcaacatttcgagtgaaacaggtaatgtaaccataaatgctgcatttatatctgtttaccgtggtggagttgatgaattactttcaataa
tgggagaatatttccctgaattgggtttaacaagagaggattgcagagaaatgctttggatccaatacttcccttttcatattaaccatccaatagataacgttat
agagtttttgaccaacagaactcctcctagcaaaccttatttcacaggcaaagctgattttgtcaaggaccctatcccagtagagggtcttgaaaagatacta
tacaagctttttgatgttcctccacttattggacagatggaatggactgtttttggaggagggataatggatgaaatccctgaatctgaaataccattcccacat
agagggaacctcttcattatgtttgaggtggtttattggtacgaaaacgatacgtctgccgtgattcaaaatcataccaattggctgagggaacttcatgaagt
tattggaaattatgttcctagcaaccccagggccgcatatgctgattatcgagatcttgacttgggggtgaacaacgttgaaggtgaaacaagcattgcaca
agcacgaatttggggtgctccatatttcaaggataactttgacaggttagtccaagtgaaaactgaggttgatcctgaaaactatttcaagaatgaacaaagt
tttccaccccttccatcttatgcctcatct

**BBE08:**
atgatgaagccaaaaactatgaagcttcaaattgcttttgcttttctacttgccttttcattaattgcatttgcatcaaagcctaatgagaaagcttttcttcaatgc
ctattgaaacattcagatgacaccacaatttcaagcatcatttacacacccaaaaactctactttcttatcagtcttggacttctacatccaaaattcacgctttct
aaatccggaaactcccaagcccaaagtcattctcaccccactcactgaatcacacattcaaactgcaatttactgtggcaagaagcataacatgcaaatga
gagttcgaagtggtggccatgactttgctggtagttcttacattgctgaggtacccttttttcgtacttgacatgttcaactttcgatccatctcagttgatgctaat
agccgcactgcttgggttggagctggtgccacccttggtgaaacatactacagcatttatgaaaagaatagctctcttggcttccctgctggttattggccta
ccgtttgcattggtgggcacatcagtggcggaggctacggtgcattgacaagggaatatggtcttgccgctgatcatgtcattgatgctcgcataattgatg
ccaccggggcaattcttgacagagcatccatgggtgaggatctcttttgggctatcagaggtggaattggagctaacttcgtagttattctttcctaccagctc
actctagttgacgttccagaaaaagttacagcattttctgtcccaagaaccctcgaacaagatgcaattcagctcgttcacaagtggcaacacgtcgccccc
aagttgccacaacagctcacgatttcagtccaatttacgagtgatgtttcagctgaaactggaaagagaactataattgccacatttatctctgtataccgtgg
tggggctgatcaacttctttcaataatgggagcacagttccctgagttgggcttgaccaaagctgattgcaaggaaatgctttggattcaatacttccccttcc
atattggccactcaatcgataacattaaagatttcttgaccagcagggtccctccaagtaagccttatttcacagccaaggctgattttgccaaggaccccat
ctcagtaaagggacttgaaaggatactgaacaatcttttcgatgtcggtccacttcttggacaaatggaatggaccattttcggaggaggtgtgatggataa
aattccggaatcggaaattccattcccacatagaggaagattgctgattatgtttcaggttgtttattggactgcaaatgatacctcatcagtgattcaatcacg
tatcgattggctaagaaagcttcacaaatctatcggaggctatgttcctaagaatccaagggctgcatatgctgattatcgtgaccttgacttgggtgtgaac

aatgttattggggaaacaagcatcgaacaagcaaagaaatggggtgctccatatttcgcgaacaattttgacaggttagtgcaagtgaaaactcaagttgat
cctgacaattacttcaagaatgaacaaagctttccaaccctcagtccttatacgacctcc

***Rgn*TDC-C-His$_6$** (protein sequence)**:**
MSQVIKKKRNTFMIGTEYILNSTQLEEAIKSFVHDFCAEKHEIHDQPVVVEAKEHQEDKIKQIKIP
EKGRPVNEVVSEMMNEVYRYRGDANHPRFFSFVPGPASSVSWLGDIMTSAYNIHAGGSKLAPM
VNCIEQEVLKWLAKQVGFTENPGGVFVSGGSMANITALTAARDNKLTDINLHLGTAYISDQTHS
SVAKGLRIIGITDSRIRRIPTNSHFQMDTTKLEEAIETDKKSGYIPFVVIGTAGTTNTGSIDPLTEISA
LCKKHDMWFHIDGAYGASVLLSPKYKSLLTGTGLADSISWDAHKWLFQTYGCAMVLVKDIRNL
FHSFHVNPEYLKDLENDIDNVNTWDIGMELTRPARGLKLWLTLQVLGSDLIGSAIEHGFQLAVW
AEEALNPKKDWEIVSPAQMAMINFRYAPKDLTKEEQDILNEKISHRILESGYAAIFTTVLNGKTV
LRICAIHPEATQEDMQHTIDLLDQYGREIYTEMKKALEKHHHHHH

***Rgn*TDC L355A-C-His$_6$** (protein sequence)**:**
MSQVIKKKRNTFMIGTEYILNSTQLEEAIKSFVHDFCAEKHEIHDQPVVVEAKEHQEDKIKQIKIP
EKGRPVNEVVSEMMNEVYRYRGDANHPRFFSFVPGPASSVSWLGDIMTSAYNIHAGGSKLAPM
VNCIEQEVLKWLAKQVGFTENPGGVFVSGGSMANITALTAARDNKLTDINLHLGTAYISDQTHS
SVAKGLRIIGITDSRIRRIPTNSHFQMDTTKLEEAIETDKKSGYIPFVVIGTAGTTNTGSIDPLTEISA
LCKKHDMWFHIDGAYGASVLLSPKYKSLLTGTGLADSISWDAHKWLFQTYGCAMVLVKDIRNL
FHSFHVNPEYLKDLENDIDNVNTWDIGME**A**TRPARGLKLWLTLQVLGSDLIGSAIEHGFQLAVW
AEEALNPKKDWEIVSPAQMAMINFRYAPKDLTKEEQDILNEKISHRILESGYAAIFTTVLNGKTV
LRICAIHPEATQEDMQHTIDLLDQYGREIYTEMKKALEKHHHHHH

***Rgn*TDC L355M-C-His$_6$** (protein sequence)**:**
MSQVIKKKRNTFMIGTEYILNSTQLEEAIKSFVHDFCAEKHEIHDQPVVVEAKEHQEDKIKQIKIP
EKGRPVNEVVSEMMNEVYRYRGDANHPRFFSFVPGPASSVSWLGDIMTSAYNIHAGGSKLAPM
VNCIEQEVLKWLAKQVGFTENPGGVFVSGGSMANITALTAARDNKLTDINLHLGTAYISDQTHS
SVAKGLRIIGITDSRIRRIPTNSHFQMDTTKLEEAIETDKKSGYIPFVVIGTAGTTNTGSIDPLTEISA
LCKKHDMWFHIDGAYGASVLLSPKYKSLLTGTGLADSISWDAHKWLFQTYGCAMVLVKDIRNL
FHSFHVNPEYLKDLENDIDNVNTWDIGME**M**TRPARGLKLWLTLQVLGSDLIGSAIEHGFQLAVW
AEEALNPKKDWEIVSPAQMAMINFRYAPKDLTKEEQDILNEKISHRILESGYAAIFTTVLNGKTV
LRICAIHPEATQEDMQHTIDLLDQYGREIYTEMKKALEKHHHHHH

***Pf*TrpB$^{2B9}$ H275E-C-His$_6$** (protein sequence)**:**
MWFGEFGGQYVPETLVGPLKELEKAYKRFKDDEEFNRQLNYYLKTWAGRPTPLYYAKRLTEKI
GGAKVYLKREDLVHGGAHKTNNAIGQALLAKLMGKTRLIAETGAGQHGVATAMAGALLGMK
VDIYMGAEDVERQKMNVFRMKLLGANVIPVNSGSRTLKDAINEALRDWVATFEYTHYLIGSVV
GPHPYPTIVRDFQSVIGREAKAQILEAEGQLPDVIVACVGGGSNAMGIFYPFVNDKKVKLVGVEA
GGKGLESGKHSASLNAGQVGVSEGMLSYFLQDEEGQIKPSHSIAPGLDYPGVGPEHAYLKKIQR
AEYVAVTDEEALKAFHELSRTEGIIPALESAHAVAYAMKLAKEMSRDEIIIVNLSGRGDKDLDIV
LKASGNVLEKHHHHHH

***Cr*DPAS** (protein sequence)**:**
MAGKSAEEEHPIKAYGWAVKDRTTGILSPFKFSRRATGDDDVRIKILYCGICHTDLASIKNEYEFL
SYPLVPGMEIVGIATEVGKDVTKVKVGEKVALSAYLGCCGKCYSCVNELENYCPEVIIGYGTPY
HDGTICYGGLSNETVANQSFVLRFPERLSPAGGAPLLSAGITSFSAMRNSGIDKPGLHVGVVGLG
GLGHLAVKFAKAFGLKVTVISTTPSKKDDAINGLGADGFLLSRDDEQMKAAIGTLDAIIDTLAVV
HPIAPLLDLLRSQGKFLLLGAPSQSLELPPIPLLSGGKSIIGSAAGNVKQTQEMLDFAAEHDITANV
EIIPIEYINTAMERLDKGDVRYRFVVDIENTLTPPSEL

***Cr*THAS1** (protein sequence)**:**
MAMASKSPSEEVYPVKAFGLAAKDSSGLFSPFNFSRRATGEHDVQLKVLYCGTCQYDREMSKN
KFGFTSYPYVLGHEIVGEVTEVGSKVQKFKVGDKVGVASIIETCGKCEMCTNEVENYCPEAGSID
SNYGACSNIAVINENFVIRWPENLPLDSGVPLLCAGITAYSPMKRYGLDKPGKRIGIAGLGGLGH
VALRFAKAFGAKVTVISSSLKKKREAFEKFGADSFLVSSNPEEMQGAAGTLDGIIDTIPGNHSLEP
LLALLKPLGKLIILGAPEMPFEVPAPSLLMGGKVMAASTAGSMKEIQEMIEFAAEHNIVADVEVIS
IDYVNTAMERLDNSDVRYRFVIDIGNTLKSN*

***Ca*IFR** (protein sequence)**:**
MAVKSKILIVGATGYIGKHIVEASAKAGHPTFALVRESTISDPKRAAIIESFKSLGVTSLYGDLYN
HQQLVNAIKQVDIVISTVGGGTEVVAHQVKIIAAIKEAGNIKRFLPSEFGGDADRWHCVEPAASW
YRTKAEIRRAVEAAGIPYTYLVSNGFAGYLNCFLNFGDFSSATPPKDKIGILGDGNSKVVFSKEE
DIAAYTIKAADDPRTLNKIVHLRPPANTLSCNEIVSLWEKKIGKTLEKIYLPEKEVLEKIQEATMPL
NLFLSVGYTIFVKGEMANFEIEASFGVEASELYPDVKYTTLDEYLNQFVSD

#### Synthetic Methods

##### *d₅*-tryptamine

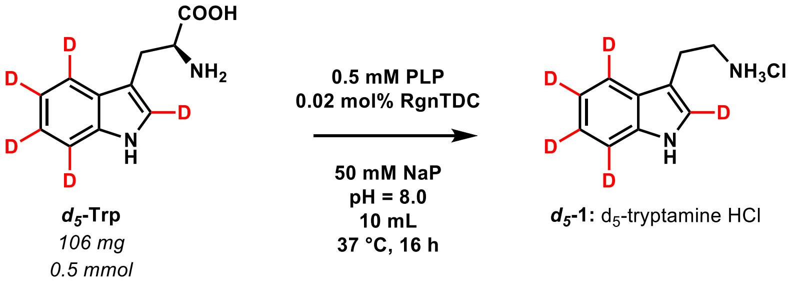


For the synthesis of $d_5$-tryptamine: to 10 mL of 50 mM sodium phosphate buffer pH = 8.0 was added 106 mg $d_5$-tryptophan (*Sigma-Aldrich*) (0.51 mmol, [final] = 51 mM), 250 µL 20 mM PLP ([final] = 0.51 mM), and 8.0 µM *Rgn*TDC (0.02 % mol catalyst).[10] The solution was then incubated for 16 h with shaking at 37 °C.

The reaction was quenched via addition of 30 mL ACN, followed by centrifugation to pellet any aggregated protein. The supernatant was removed and the pellet washed with 10 mL ACN, followed again by centrifugation. The combined supernatant fractions were pooled and the ACN evaporated. 0.5 mL 10 M NaOH was used to alkalize the resulting aqueous suspension, which was then extracted three times using 15 mL ethyl acetate (EtOAc). The combined organic layers were then concentrated to 20 mL via evaporation. The EtOAc layer was extracted three times with 15 mL 10 mM HCl in $H_2O$, the aqueous layers combined, frozen using liquid $N_2$, and dried using lyophilization. A tannish powder (92 mg) corresponding to $d_5$-tryptamine HCl was isolated in 90% yield.

##### *d₄*-strictosidine/d₄-vincoside

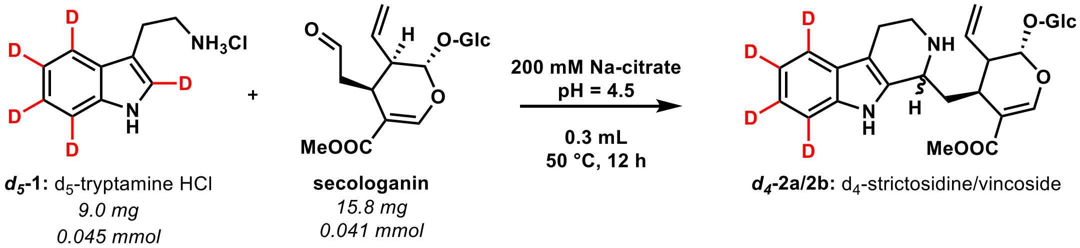


For the synthesis of $d_4$-strictosidine/vincoside: to a 1.5 mL microcentrifuge tube was added 9.0 mg $d_5$-tryptamine HCl (0.045 mmol, [final] = 150 mM) and 15.8 mg secologanin (0.041 mmol, [final] = 136 mM). 0.3 mL 200 mM sodium citrate buffer pH = 4.5 was then added, and the reaction heated to 50 °C with shaking for 12 h. Full conversion of secologanin was observed. The mixture was injected onto preparative HPLC for purification. Fractions corresponding to $d_4$-strictosidine and $d_4$-vincoside were collected and pooled. These pooled fractions were evaporated to remove ACN, frozen, and lyophilized to remove solvent. 5.1 mg $d_4$-strictosidine (white powder, 23% yield), and 3.6 mg $d_4$-vincoside (white powder, 17% yield) were obtained.

**4-methoxytryptamine**

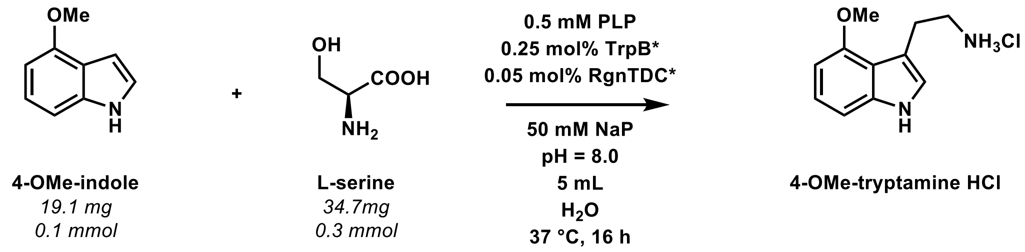


For the synthesis of 4-methoxytryptamine: to 5 mL of 50 mM sodium phosphate buffer pH = 8.0 was added 19.1 mg 4-methoxyindole (0.13 mmol, [final] = 26 mM), 34.7 mg serine (0.33 mmol, [final] = 66 mM), 163 µL 20 mM PLP ([final] = 0.65 mM), 65 µM *Pf*TrpB$^{2B9}$ H275E (0.25 % mol catalyst), and 13 µM *Rgn*TDC L355M (0.05 % mol catalyst).[9] The mixed solution was incubated for 16 h with shaking at 37 °C.

The reaction was quenched via addition of 5 mL ACN, followed by centrifugation to pellet aggregated protein. The supernatant was removed and the pellet washed with 10 mL of 10 mM HCl in MeOH, followed again by centrifugation. The combined supernatant fractions were pooled and the ACN/MeOH evaporated. 0.5 mL 10 M NaOH was used to alkalize the resulting aqueous suspension, which was then extracted three times using 15 mL EtOAc. The combined organic layers were then concentrated to 10 mL via evaporation. The EtOAc layer was extracted three times with 5 mL 10 mM HCl in $H_2O$, the aqueous layers combined, frozen using liquid $N_2$, and dried using lyophilization. 10.7 mg of an off-white powder corresponding to 4-methoxy-tryptamine HCl was isolated in a 36% yield. $^1$H-NMR analysis matched previous reports of 4-methoxytryptamine.[9]

#### $d_2$-4-methoxytryptamine

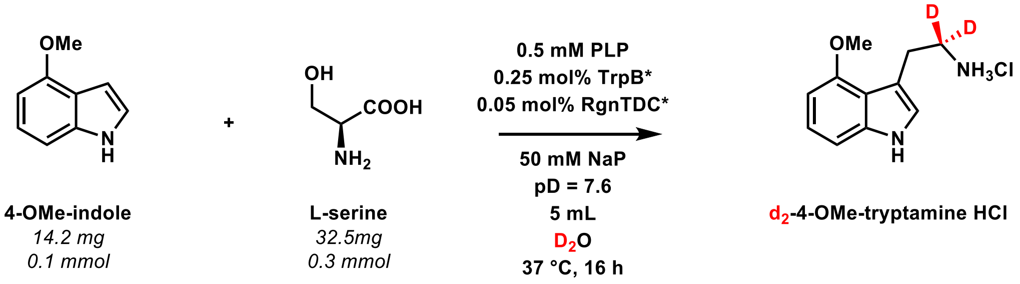


For the synthesis of $d_2$-4-methoxytryptamine: to 5 mL of 50 mM sodium phosphate buffer pD = 7.6 was added 14.2 mg 4-methoxyindole (0.10 mmol, [final] = 19 mM), 32.5 mg serine (0.31 mmol, [final] = 62 mM), 120 µL 20 mM PLP ([final] = 0.48 mM), 48 µM *Pf*TrpB$^{2B9}$ H275E (0.25 % mol catalyst),[10] and 10 µM *Rgn*TDC L355A (0.05 % mol catalyst).[10] All solutions were prepared in $D_2O$, and enzyme solutions were buffer-exchanged into 50 mM sodium phosphate buffer pD = 7.6. The solution was then incubated for 16 h with shaking at 37 °C.

The reaction was quenched via addition of 5 mL ACN, followed by centrifugation to pellet aggregated protein. The supernatant was removed and the pellet washed with 10 mL of 10 mM HCl in MeOH, followed again by centrifugation. The combined supernatant fractions were pooled and the ACN/MeOH evaporated. 0.5 mL 10 M NaOH was used to alkalize the resulting aqueous suspension, which was then extracted three times using 15 mL EtOAc. The combined organic layers were then concentrated to 10 mL via evaporation. The EtOAc layer was extracted three times with 5 mL 10 mM HCl in $H_2O$, the aqueous layers combined, frozen using liquid $N_2$, and dried using lyophilization. 13.4 mg of an off-white powder corresponding to $d_2$-4-methoxy-tryptamine HCl was isolated in a 61% yield. $^1$H-NMR analysis matched previous reports of 4-methoxytryptamine[9] and revealed a deuterium labeling ratio of 7:3 for $d_2$:$d_1$.

#### 9-methoxystrictosidine

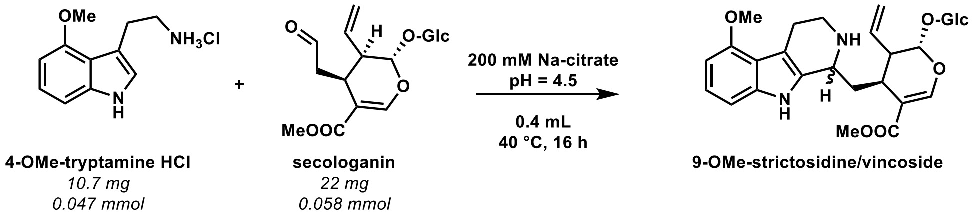


For the synthesis of 9-methoxystrictosidine: to a 1.5 mL microcentrifuge tube was added 10.7 mg 4-methoxy-tryptamine HCl (0.047 mmol, [final] = 120 mM) and 22 mg secologanin (0.058 mmol, [final] = 140 mM). 0.4 mL 200 mM sodium citrate buffer pH = 4.5 was then added, and the reaction heated to 40 °C with shaking for 16 h. Full consumption of tryptamine was was observed, and so the mixture was injected onto preparative HPLC for purification. Fractions corresponding to 9-methoxystrictosidine and 9-methoxyvincoside were collected and pooled. These pooled fractions were evaporated to remove ACN, frozen, and lyophilized to remove solvent. 3.0 mg 9-methoxystrictosidine (**9OS**, white powder, 21% yield), and 2.9 mg 9-methoxyvincoside (white powder, 24% yield).

#### *d₂*-9-methoxystrictosidine

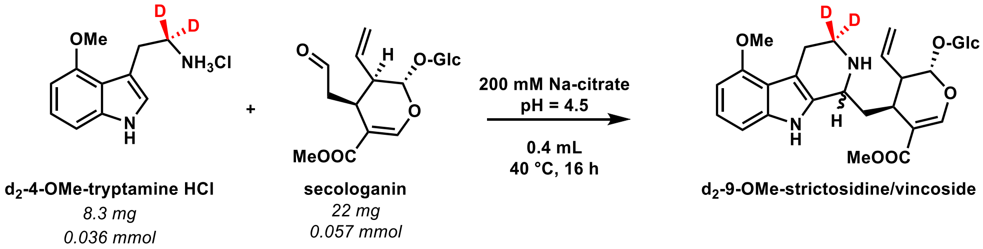


For the synthesis of $d_2$-9-methoxystrictosidine: to a 1.5 mL microcentrifuge tube was added 8.3 mg $d_2$-4-methoxytryptamine HCl (0.036 mmol, [final] = 91 mM) and 22 mg secologanin (0.057 mmol, [final] = 140 mM). 0.4 mL 200 mM sodium citrate buffer pH = 4.5 was then added, and the reaction heated to 40 °C with shaking for 16 h. Full tryptamine conversion was observed, and so the mixture was injected onto preparative HPLC for purification. Fractions corresponding to unreacted secologanin, $d_2$-9-methoxystrictosidine, and $d_2$-9-methoxyvincoside were collected and pooled. These pooled fractions were evaporated to remove ACN, frozen, and lyophilized to remove solvent. 7.4 mg of secologanin was recovered (34% of original), along with 4.4 mg $d_2$-9-methoxystrictosidine (***d₂*-9OS**, white powder, 14% yield), and 4.9 mg $d_2$-9-methoxyvincoside (white powder, 13% yield).

**Synthesis of Dehydromitragynine trifluoroacetate (14a).**

Dehydromitragynine trifluoroacetate (**14a**) was prepared over two steps from mitragynine (**5a**) according to methods described previously.[11,12]

**7-Hydroxymitragynine (24a)**

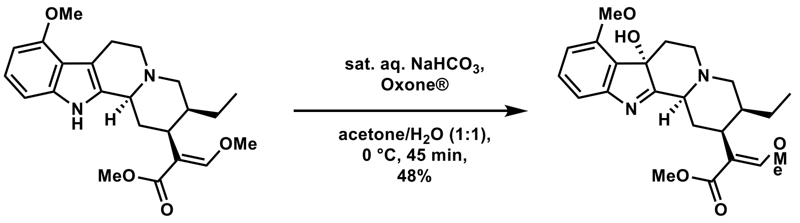


**5a**: mitragynine
*4 mg*
*0.010 mmol*

**24a**: 7-hydroxymitragynine

To a 1.5 mL vial containing (mitragynine (**5a**, 4 mg, 10 μmol) dissolved in anhydrous acetone (0.33 mL) was added sat. aq. $NaHCO_3$ (0.2 mL). The resulting amber suspension was cooled to 0 °C and then treated dropwise with a solution of Oxone® (6.16 mg, 10 μmol) in $H_2O$ (0.1 mL). After stirring at 0 °C for 45 min, the mixture was diluted with $H_2O$ (0.5 mL) and extracted with EtOAc (0.25 mL x 5). The combined organic layers were dried over anhydrous $Na_2SO_4$ and concentrated under a gentle stream of Ar to give an amber residue that was further purified by PTLC (2% [v/v] $NEt_3$ in EtOAc/hexane [2:3]) to afford title compound **24a** as a pale amber residue (2 mg, 48%): $R_f$ 0.17 (2% [v/v] $NEt_3$ in EtOAc/hexane [2:3]).

**Dehydromitragynine trifluoroacetate (14a)**

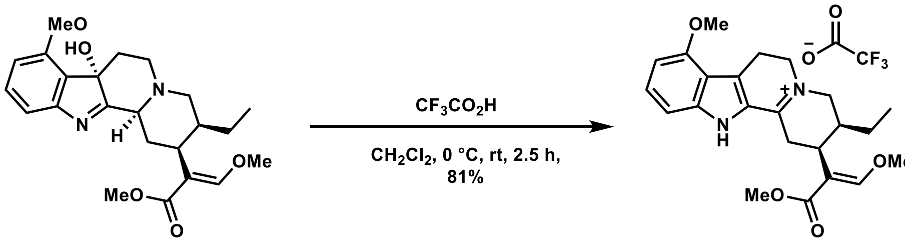


**24a**: 7-hydroxymitragynine
*2 mg*
*0.005 mmol*

**14a**: DHM-TFA

To a 1.5 mL vial containing 7-hydroxymitragynine (**24a**, 2 mg, 4.8 μmol) was added anhydrous $CH_2Cl_2$ (0.5 mL). Upon cooling to 0 °C, the solution was charged with trifluoroacetic acid (2.2 μL, 0.28 μmol) and allowed to stir for 2.5 h whilst slowly warming to rt. The reaction mass was concentrated under a gentle stream of argon to afford title compound **14a** as a bright yellow residue (2 mg, 81%).

### Structural Data

#### General Methods

NMR measurements were carried out on a 400 MHz Bruker Avance III HD spectrometer (Bruker Biospin GmbH, Rheinstetten, Germany), a 700 MHz Bruker Avance III HD spectrometer (Bruker Biospin GmbH, Rheinstetten, Germany) and a 500 MHz Bruker Avance III HD spectrometer (Bruker Biospin GmbH, Rheinstetten, Germany) using standard pulse sequences as implemented in Bruker Topspin ver. 3.6.1. 700 MHz and 500 MHz NMR were equipped with a TCI cryoprobe. Chemical shifts were referenced to the residual solvent signals of $CDCl_3$ ($\delta_H$ 7.26/$\delta_C$ 77.16), MeOD and MeOH-$d_3$ ($\delta_H$ 3.31/$\delta_C$ 49.0), $D_2O$ ($\delta_H$ 4.70) respectively. Chemical shift values ($\delta_H$) are reported in parts per million (ppm) and coupling constants ($J$) are expressed in Hertz (Hz), in the following format; chemical shift value (multiplicity, coupling constant, integration). $^1$H NMR spectral data are described, using the following abbreviations; s (singlet), d (doublet), t (triplet), dd (doublet of doublets), appbrs (apparent broad singlet), and m (multiplet). All spectra were recorded at 298 K.

#### NMR data

**tryptamine HCl (1)**: $^1$H-NMR (400 MHz, $D_2O$) $\delta$ ppm: 7.60 (*brd, J*= 7.9 Hz, 1H), 7.46 (*brd, J*= 8.1 Hz, 1H), 7.22 (*s*, 1H), 7.20 (*brdd, J*= 7.9, 7.2 Hz, 1H), 3.27-3.21 (*m*, 2H), 3.11-3.05 (*m*, 2H). Corresponding spectra are shown in Figure S38.

***d*$_5$-tryptamine (1- *d*$_5$):** $^1$H-NMR (400 MHz, $D_2O$) $\delta$ ppm: 3.20-3.14 (*m*, 2H), 3.05-2.99 (*m*, 2H). Corresponding spectra are shown in Figure S39.

***d*$_4$-strictosidine (*d*$_4$-2a):** $^1$H-NMR (500 MHz, MeOH-$d_3$) $\delta$ ppm: 7.78 (*s*, 1H), 5.85 (*ddd, J*= 17.4, 10.6, 7.7 Hz, 1H), 5.84 (*d, J*= 9.0 Hz, 1H), 5.34 (*ddd, J*= 17.4, 1.2, 1.2 Hz, 1H), 5.26 (*brd, J*= 10.6, 1.2, 1.2 Hz, 1H), 4.79 (*d, J*= 8.0 Hz, 1H), 4.53 (*brd, J*= 11.1, 1H), 3.97 (*dd, J*= 11.9, 1.8 Hz, 1H), 3.78 (*s*, 3H), 3.69-3.61 (*m*, 2H), 3.43-3.33 (*m*, 3H), 3.26-3.19 (*m*, 2H), 3.10-3.02 (*m*, 2H), 2.99 (*ddd, J*= 16.2, 4.9, 4.5 Hz, 1H), 2.73 (*m*, 1H), 2.28 (*ddd, J*= 14.8, 11.9, 2.9 Hz, 1H), 2.18 (*ddd, J*= 14.8, 11.4, 3.7 Hz, 1H). $^{13}$C-NMR (126 MHz, $CDCl_3$) $\delta$ ppm: 171.0, 156.6, 138.2, 135.4, 131.0, 127.4, 122.7, 119.9, 119.6, 118.5, 111.8, 109.1, 107.2, 100.3, 97.2, 78.8, 78.1, 74.7, 71.8, 63.0, 52.9, 52.4, 45.4, 42.8, 35.0, 32.5, 19.9. The chemical shifts were in agreement with published data of strictosidine (Misa *et al*. 2022).[13] Corresponding spectra are shown in Figure S40.

**vincoside (2b):** $^1$H-NMR (400 MHz, MeOD) $\delta$ ppm: 7.52 (*s*, 1H), 7.44 (*brd, J*= 7.8 Hz, 1H), 7.34 (*brd, J*= 8.1 Hz, 1H), 7.12 (*brdd, J*= 8.1, 7.5 Hz, 1H), 7.03 (*brdd, J*= 7.8, 7.5 Hz, 1H), 5.98 (*ddd, J*= 17.4, 10.6,

9.0 Hz, 1H), 5.67 (*d, J*= 7.0 Hz, 1H), 5.40 (*brd, J*= 17.4 Hz, 1H), 5.35 (*brd, J*= 10.6 Hz, 1H), 4.75 (*d, J*= 7.9 Hz, 1H), 4.60 (*brs*, 1H), 3.98 (*dd, J*= 11.9, 2.0 Hz, 1H), 3.68 (*dd, J*= 11.9, 6.6 Hz, 1H), 3.65 (*m*, 1H), 3.64 (*s*, 3H), 3.43-3.34 (*m*, 3H), 3.29-3.20 (*m*, 2H), 3.12 (*m*, 1H), 3.07-2.91 (*m*, 2H), 2.81 (*m*, 1H), 2.40 (*m*, 1H), 2.14 (*m*, 1H). Corresponding spectra are shown in Figure S41.

**$d_4$-vincoside ($d_4$-2b):** $^1$H-NMR (400 MHz, MeOD) $\delta$ ppm: 7.53 (*s*, 1H), 5.98 (*ddd, J*= 17.4, 10.5, 8.7 Hz, 1H), 5.67 (*d, J*= 7.1 Hz, 1H), 5.41 (*brd, J*= 17.4 Hz, 1H), 5.36 (*brd, J*= 10.5 Hz, 1H), 4.76 (*d, J*= 7.9 Hz, 1H), 4.65 (*brs*, 1H), 3.99 (*dd, J*= 11.8, 2.1 Hz, 1H), 3.68 (*dd, J*= 11.8, 6.6 Hz, 1H), 3.65 (*m*, 1H), 3.64 (*s*, 3H), 3.43-3.34 (*m*, 3H), 3.29-3.20 (*m*, 2H), 3.12 (*m*, 1H), 3.07-2.92 (*m*, 2H), 2.81 (*m*, 1H), 2.43 (*m*, 1H), 2.14 (*m*, 1H). Corresponding spectra are shown in Figure S42.

**4-methoxytryptamine HCl:** $^1$H-NMR (400 MHz, $D_2O$) $\delta$ ppm: 7.11-7.00 (*m*, 3H), 6.56 (*brd, J*= 7.5 Hz, 1H), 3.84 (*s*, 3H), 3.23-3.18 (*m*, 2H), 3.12-3.06 (*m*, 2H). Corresponding spectra are shown in Figure S43.

**$d_2$-4-methoxytryptamine HCl:** $^1$H-NMR (400 MHz, $D_2O$) $\delta$ ppm: 7.11-7.00 (*m*, 3H), 6.56 (*brd, J*= 7.5 Hz, 1H), 3.84 (*s*, 3H), 3.21-3.17 (*m*, 0.3H), 3.11-3.06 (*m*, 2H). Corresponding spectra are shown in Figure S44.

**9-methoxystrictosidine (9OS)**: $^1$H-NMR (500 MHz, MeOH-$d_3$) $\delta$ ppm: 7.76 (*s*, 1H), 6.98 (*dd*, *J*= 8.1, 7.8 Hz, 1H), 6.88 (*d*, *J*= 8.1 Hz, 1H), 6.46 (*d*, *J*= 7.8 Hz, 1H), 5.84 (*ddd*, *J*= 17.2, 10.7, 7.7 Hz, 1H), 5.82 (*d*, *J*= 8.9 Hz, 1H), 5.33 (*ddd*, *J*= 17.2, 1.2, 1.2 Hz, 1H), 5.25 (*ddd*, *J*= 10.7, 1.2, 1.2 Hz, 1H), 4.78 (*d*, *J*= 7.9 Hz, 1H), 4.42 (*brd*, *J*= 11.6 Hz, 1H), 3.97 (*dd*, *J*= 11.8, 1.4 Hz, 1H), 3.85 (*s*, 3H), 3.78 (*s*, 3H), 3.64 (*dd*, *J*= 11.8, 6.8 Hz, 1H), 3.55 (*ddd*, *J*= 12.0, 5.0, 5.0 Hz, 1H), 3.40 (*dd*, *J*= 9.0, 8.7 Hz, 1H), 3.35 (*m*, 1H), 3.26 (*m*, 1H), 3.23 (*dd*, *J*= 9.2, 9.0 Hz, 1H), 3.22 (*dd*, *J*= 8.7, 7.9 Hz, 1H), 3.19 (*m*, 2H), 3.05 (*ddd*, *J*= 11.8, 4.5, 3.8 Hz, 1H), 2.72 (*m*, 1H), 2.22 (*ddd*, *J*= 14.5, 11.8, 2.7 Hz, 1H), 2.13 (*ddd*, *J*= 14.5, 11.6, 3.8 Hz, 1H). $^{13}$C-NMR (126 MHz, MeOH-$d_3$) $\delta$ ppm: 170.8, 156.3, 155.6, 139.5, 135.5, 129.9, 123.8, 119.5, 117.7, 109.3, 107.3, 105.5, 100.3, 100.3, 97.3, 78.7, 78.1, 74.7, 71.8, 63.0, 55.4, 52.6, 52.3, 45.4, 43.0, 35.4, 32.5, 22.3. Corresponding spectra are shown in Figure S45.

**$d_2$-9-methoxystrictosidine ($d_2$-9OS)**: $^1$H-NMR (500 MHz, MeOH-$d_3$) $\delta$ ppm: 7.75 (*s*, 1H), 6.98 (*dd*, *J*= 8.1, 7.8 Hz, 1H), 6.87 (*d*, *J*= 8.1 Hz, 1H), 6.45 (*d*, *J*= 7.8 Hz, 1H), 5.84 (*ddd*, *J*= 17.3, 10.7, 7.7 Hz, 1H), 5.82 (*d*, *J*= 8.8 Hz, 1H), 5.33 (*ddd*, *J*= 17.3, 1.2, 1.2 Hz, 1H), 5.25 (*ddd*, *J*= 10.7, 1.2, 1.2 Hz, 1H), 4.78 (*d*, *J*= 7.9 Hz, 1H), 4.41 (*brd*, *J*= 11.2 Hz, 1H), 3.97 (*dd*, *J*= 11.8, 1.8 Hz, 1H), 3.85 (*s*, 3H), 3.78 (*s*, 3H), 3.64 (*dd*, *J*= 11.8, 6.9 Hz, 1H), 3.52 (*dd*, *J*= 5.0, 5.0 Hz, 0.3H), 3.39 (*dd*, *J*= 9.0, 8.7 Hz, 1H), 3.35 (*m*, 1H), 3.23 (*m*, 1H), 3.22 (*m*, 1H), 3.17 (*m*, 2H), 3.05 (*ddd*, *J*= 11.5, 4.3, 3.9 Hz, 1H), 2.71 (*ddd*, *J*= 8.8, 7.7, 4.3 Hz, 1H), 2.21 (*ddd*, *J*= 14.7, 11.5, 2.7 Hz, 1H), 2.13 (*ddd*, *J*= 14.7, 11.2, 3.9 Hz, 1H). $^{13}$C-NMR (126 MHz, MeOH-$d_3$) $\delta$ ppm: 170.8, 156.3, 155.6, 139.5, 135.5, 129.9, 123.8, 119.5, 117.7, 109.4, 107.3,

105.5, 100.3, 100.3, 97.3, 78.7, 78.1, 74.7, 71.8, 63.0, 55.4, 52.6, 52.3, 45.4, 42.8, 35.4, 32.5, 22.1. Corresponding spectra are shown in Figures S46-S47.

**20*S*-corynantheidine (3a)**: $^1$H-NMR (500 MHz, $CDCl_3$) $\delta$ ppm: 7.71 (*brs*, NH, 1H), 7.46 (*brd, J*= 7.8 Hz, 1H), 7.44 (*s*, 1H), 7.29 (*brd*, *J*= 7.8 Hz, 1H), 7.11 (*ddd*, *J*= 7.8, 7.2, 1.4 Hz, 1H), 7.07 (*ddd*, *J*= 7.8, 7.2, 1.2 Hz, 1H), 3.73 (*s*, 3H), 3.71 (*s*, 3H), 3.20 (*m*, 1H), 3.08-2.94 (*m*, 4H), 2.70 (*m*, 1H), 2.57 (*m*, 1H), 2.53 (*m*, 1H), 2.49 (*ddd, J*= 11.5, 3.1, 0.7 Hz, 1H), 1.84 (*m*, 1H), 1.78 (*m*, 1H), 1.64 (*m*, 1H), 1.21 (*m*, 1H), 0.87 (*t*, *J*= 7.5 Hz, 3H). $^{13}$C-NMR (126 MHz, $CDCl_3$) $\delta$ ppm: 169.3, 160.7, 136.0, 135.8, 127.7, 121.3, 119.5, 118.3, 111.6, 110.8, 108.3, 61.7, 61.4, 57.9, 53.7, 51.5, 40.9, 40.1, 30.1, 22.1, 19.3, 13.0. The chemical shifts were in agreement with published data (Stærk *et al*. 2000, Wanner *et al*. 2011).[14,15] Corresponding spectra are shown in Figure S48.

**20*R*-corynantheidine (3b)**: $^1$H-NMR (700 MHz, $CDCl_3$) $\delta$ ppm: 7.46 (*brd*, *J*= 7.6 Hz, 1H), 7.36 (*brs*, 1H), 7.27 (*d*, *J*= 7.8 Hz, 1H), 7.11 (*brdd*, *J*= 7.8, 7.4 Hz, 1H), 7.07 (*brdd*, *J*= 7.6, 7.4 Hz, 1H), 3.92-3.61 (*m*, 6H), 3.33 (*m*, 1H), 3.25-3.13 (*m*, 2H), 3.06 (*m*, 1H), 2.75 (*m,*1H), 2.70-2.61 (*m*, 2H), 2.34-1.87 (*m*, 4H), 1.43 (*m*, 1H), 1.06 (*m*, 1H), 0.88 (*t*, *J*= 7.0 Hz, 3H). $^{13}$C-NMR (176 MHz, $CDCl_3$) $\delta$ ppm: 168.3, 160.1, 136.3, 127.6, 121.7, 119.6, 118.4, 108.1, 61.8, 60.9, 60.5, 53.2, 40.1, 38.7, 33.8, 24.5, 21.6, 11.4. Due to low amount (0.09 mg) and broad signals only partial assignment was possible. The chemical shifts were in agreement with published data (Ren *et al*. 2024).[16] Corresponding spectra are shown in Figure S49.

**20*S*-isocorynantheidine (6a)**: $^1$H-NMR (700 MHz, $CDCl_3$) $\delta$ ppm: 7.46 (*d*, *J*= 7.8 Hz, 1H), 7.45 (*s*, 1H), 7.32 (*d*, *J*= 7.6 Hz, 1H), 7.14-7.07 (*m*, 2H), 4.45 (*brs*, 1H), 3.81 (*s*, 3H), 3.67 (*s*, 3H), 3.36 (*m*, 2H), 3.15 (*m,*1H), 3.13 (*m,*1H), 2.74 (*m,*1H), 2.68 (*m,*1H), 2.07 (*m,*1H), 1.62 (*m*, 1H), 1.25 (*m*, 1H), 0.89 (*t*, *J*= 7.3 Hz, 3H). $^{13}$C-NMR (176 MHz, $CDCl_3$) $\delta$ ppm: 169.3, 160.9, 136.2, 127.3, 121.9, 119.8, 118.1, 111.3, 110.7, 107.6, 61.9, 55.3, 52.4, 51.6, 38.9, 20.8, 12.5. Due to low amount (0.17 mg) and broad signals only partial assignment was possible. The chemical shifts were in agreement with published data (Lounasmaa *et al*. 1998).[17] Corresponding spectra are shown in Figure S50.

**hirsutine (6b)**: $^1$H-NMR (500 MHz, $CDCl_3$) $\delta$ ppm: 8.91 (*brs*, NH, 1H), 7.48 (*brd, J*= 7.7 Hz, 1H), 7.44 (*d*, *J*= 8.2 Hz, 1H), 7.34 (*s*, 1H), 7.22 (*dd*, *J*= 8.2, 7.4 Hz, 1H), 7.15 (*dd*, *J*= 7.7, 7.4 Hz, 1H), 4.87 (*brs*, 1H), 3.79 (*brs*, 3H), 3.69 (*s*, 3H), 3.43 (*m*, 2H), 3.10 (*dd*, *J*= 11.4, 3.1 Hz, 1H), 3.01 (*m*, 1H), 2.76 (*m*, 1H), 2.69 (*m*, 1H), 2.61 (*dd*, *J*= 11.4, 11.4 Hz, 1H), 2.47 (*m*, 1H), 2.30 (*brdd, J*= 11.5, 11.5 Hz, 1H), 2.17 (*brd, J*= 13.4 Hz, 1H), 1.35 (*m*, 1H), 0.82 (*m*, 1H), 0.75 (*t*, *J*= 7.2 Hz, 3H). $^{13}$C-NMR (126 MHz, $CDCl_3$) $\delta$ ppm: 168.6, 160.5, 136.8, 129.2, 127.0, 122.4, 120.0, 118.3, 111.8, 110.2, 106.3, 61.9, 54.3, 51.6, 50.2,

49.7, 37.0, 33.5, 30.3, 24.1, 16.4, 10.9. The chemical shifts were in agreement with published data (Ren *et al*. 2024).[16] Corresponding spectra are shown in Figure S51.

**mitragynine (5a)**: $^{1}$H-NMR (500 MHz, $CDCl_3$) $\delta$ ppm: 7.68 (*brs*, NH, 1H), 7.43 (*s*, 1H), 6.99 (*dd*, *J*= 8.1, 7.7 Hz, 1H), 6.90 (*dd*, *J*= 8.1, 0.6 Hz, 1H), 6.45 (*brd*, *J*= 7.7 Hz, 1H), 3.87 (*s*, 3H), 3.73 (*s*, 3H), 3.71 (*s*, 3H), 3.16 (*brd*, *J*= 11.4 Hz, 1H), 3.12 (*m*, 1H), 3.06-2.89 (*m*, 4H), 2.53 (*ddd*, *J*= 11.5, 11.5, 4.2 Hz, 1H), 2.51 (*m*, 1H), 2.45 (*m*, 1H), 1.80 (*m*, 1H), 1.78 (*m*, 1H), 1.62 (*m*, 1H), 1.20 (*m*, 1H), 0.87 (*t*, *J*= 7.4 Hz, 3H). $^{13}$C-NMR (126 MHz, $CDCl_3$) $\delta$ ppm: 169.4, 160.7, 154.7, 137.4, 133.8, 122.0, 117.8, 111.7, 108.1, 104.3, 99.9, 61.7, 61.4, 57.9, 55.5, 53.9, 51.5, 40.8, 40.1, 30.1, 24.1, 19.2, 13.0. The chemical shifts were in agreement with published data (Flores-Bocanegra *et al*. 2020).[18] Corresponding spectra are shown in Figure S52.

**speciogynine (5b)**: $^{1}$H-NMR (700 MHz, $CDCl_3$) $\delta$ ppm: 7.37 (*brs*, 1H), 7.01 (*brdd*, *J*= 7.7, 7.5 Hz, 1H), 6.92 (*brs*, 1H), 6.43 (*brd*, *J*= 7.5 Hz, 1H), 3.94-3.55 (*m*, 9H), 3.55-3.08 (*m*, 6H), 2.66 (*m*, 1H), 2.27 (*m*, 2H), 2.03 (*m*, 1H), 1.41 (*m*, 1H), 1.04 (*m*, 1H), 0.88 (*t*, *J*= 6.9 Hz, 3H). $^{13}$C-NMR (176 MHz, $CDCl_3$) $\delta$ ppm: 168.5, 160.4, 154.6, 137.9, 122.7, 117.2, 104.8, 99.9, 62.0, 60.6, 59.7, 55.4, 54.6, 39.1, 32.7, 24.1, 22.8, 14.2. Due to low amount (0.31 mg) and broad signals only partial assignment was possible. The chemical shifts were in agreement with published data (Flores-Bocanegra *et al*. 2020).[18] Corresponding spectra are shown in Figure S53.

**speciociliatine (7a)**: $^{1}$H-NMR (700 MHz, $CDCl_3$) $\delta$ ppm: 7.80 (*brs*, NH, 1H), 7.42 (*s*, 1H), 7.00 (*dd*, *J*= 8.1, 7.8 Hz, 1H), 6.91 (*d*, *J*= 8.1, 1H), 6.47 (*d*, *J*= 7.8 Hz, 1H), 4.13 (*brs*, 1H), 3.89 (*s*, 3H), 3.77 (*s*, 3H), 3.66 (*s*, 3H), 3.19 (*m,*1H), 3.12 (*dd*, *J*= 12.7, 5.8 Hz, 1H), 3.00 (*m*, 1H), 2.97 (*m,*1H), 2.90 (*m*, 2H), 2.77 (*dd, J*= 11.3, 5.8 Hz, 1H), 2.62 (*m*, 1H), 1.91 (*m*, 1H), 1.75 (*m*, 1H), 1.63 (*m*, 1H), 1.25 (*m*, 1H), 0.89 (*t*, *J*= 7.4 Hz, 3H). $^{13}$C-NMR (176 MHz, $CDCl_3$) $\delta$ ppm: 169.6, 160.1, 154.5, 137.2, 132.1, 121.9, 118.0, 111.6, 108.1, 104.5, 99.8, 61.5, 55.4, 54.7, 52.7, 51.5, 50.3, 39.8, 33.8, 30.1, 20.9, 20.5, 12.7. The chemical shifts were in agreement with published data (Flores-Bocanegra *et al*. 2020).[18] Corresponding spectra are shown in Figure S54.

**mitraciliatine (7b)**: $^{1}$H-NMR (700 MHz, $CDCl_3$) $\delta$ ppm: 7.32 (*s*, 1H), 7.06 (*dd*, *J*= 8.1, 7.7 Hz, 1H), 7.01 (*d*, *J*= 8.1, 1H), 6.50 (*d*, *J*= 7.7 Hz, 1H), 4.61 (*brs*, 1H), 3.90 (*s*, 3H), 3.78 (*brs*, 3H), 3.69 (*s*, 3H), 3.16 (*brd*, *J*= 11.4 Hz, 1H), 3.34 (*m*, 2H), 3.21 (*m,*1H), 2.99 (*brd, J*= 16.4 Hz, 1H), 2.92 (*brd, J*= 10.5 Hz, 1H), 2.56 (*m*, 1H), 2.51 (*brdd, J*= 11.3, 11.3 Hz, 1H), 2.31 (*m*, 1H), 2.25 (*m*, 1H), 2.02 (*m*, 1H), 1.33 (*m*, 1H), 0.80 (*m*, 1H), 0.76 (*t*, *J*= 7.1 Hz, 3H). $^{13}$C-NMR (176 MHz, $CDCl_3$) $\delta$ ppm: 169.1, 160.1, 154.5, 137.6, 129.3, 122.5, 117.8, 111.2, 107.3, 104.8, 99.8, 61.8, 55.4, 54.3, 51.5, 51.3, 50.2, 38.2, 34.4, 31.3, 24.3,

18.9, 11.2. The chemical shifts were in agreement with published data (Flores-Bocanegra *et al*. 2020).[18] Corresponding spectra are shown in Figure S55.

**3-isoajmalicine (11b)**: $^1$H-NMR (500 MHz, $CDCl_3$) $\delta$ ppm: 8.38 (*brs*, NH, 1H), 7.50 (*d, J*= 1.6 Hz, 1H), 7.49 (*brd*, *J*= 8.0 Hz, 1H), 7.41 (*ddd*, *J*= 8.1, 0.9, 0.9 Hz, 1H), 7.19 (*ddd*, *J*= 8.1, 7.2, 0.9 Hz, 1H), 7.12 (*ddd*, *J*= 8.0, 7.2, 0.9 Hz, 1H), 4.65 (*m*, 1H), 4.35 (*qd*, *J*= 6.7, 4.2 Hz, 1H), 3.73 (*s*, 3H), 3.34 (*m*, 2H), 3.22 (*ddd*, *J*= 14.0, 2.6, 2.6 Hz, 1H), 3.01 (*dddd*, *J*= 16.2, 10.4, 8.4, 2.7 Hz, 1H), 2.76 (*dd*, *J*= 10.7, 3.0 Hz, 1H), 2.70 (*m*, 1H), 2.58 (*dd*, *J*= 11.1, 10.7 Hz, 1H), 2.11 (*m*, 1H), 2.00 (*m*, 1H), 1.73 (*ddd*, *J*= 14.0, 12.0, 5.0 Hz, 1H), 0.92 (*d*, *J*= 6.7 Hz, 3H). $^{13}$C-NMR (126 MHz, $CDCl_3$) $\delta$ ppm: 167.5, 154.9, 136.1, 131.7, 127.7, 121.9, 119.7, 118.2, 111.5, 107.6, 106.6, 73.7, 54.2, 51.2, 50.6, 47.1, 40.8, 30.9, 26.0, 16.8, 15.1. The chemical shifts were in agreement with published data (Ren *et al*. 2024).[16] Corresponding spectra are shown in Figure S62.

**Isomitraphylline (12b (3*S*, 7*S*))**: $^1$H-NMR (700 MHz, $CDCl_3$) $\delta$ ppm: 8.02 (*brs*, NH, 1H), 7.39 (*s,* 1H), 7.35 (*m*, 1H), 7.17 (*brdd*, *J*= 7.6, 7.2 Hz, 1H), 7.00 (*m*, 1H), 6.86 (*brd*, *J*= 7.6 Hz, 1H), 4.37 (*m*, 1H), 3.57 (*s*, 3H), 3.29 (*m*, 1H), 3.13 (*m*, 1H), 2.61 (*m*, 1H), 2.54 (*m*, 1H), 2.42 (*m*, 1H), 2.21 (*m*, 1H), 2.18 (*m*, 1H), 2.04 (*m*, 1H), 1.93 (*m*, 1H), 1.91 (*m*, 1H), 1.12 (*d*, *J*= 6.6 Hz, 3H), 0.61 (*m*, 1H). $^{13}$C-NMR (126 MHz, $CDCl_3$) $\delta$ ppm: 181.3, 167.2, 154.1, 140.3, 134.0, 127.7, 125.1, 122.5, 109.7, 107.4, 74.2, 72.0, 56.5, 54.4, 53.5, 51.0, 41.0, 35.6, 30.2, 29.2, 15.0. The $^{13}$C chemical shifts were in agreement with published data (Paradowska *et al*. 2008).[19] Corresponding spectra are shown in Figure S63.

**dehydromitragynine – TFA salt (DHM, 14a)**: $^1$H-NMR (500 MHz, $CDCl_3$) $\delta$ ppm: 12.0 (*s*, 1H), 7.52 (*s*, 1H), 7.29 (*dd*, *J*= 8.3, 7.7 Hz, 1H), 7.13 (*d*, *J*= 8.3 Hz, 1H), 6.42 (*d*, *J*= 7.7 Hz, 1H), 3.95-3.80 (*m*, 2H), 3.91 (*s*, 3H), 3.78 (*s*, 3H), 3.62 (*s*, 3H), 3.60-3.53 (*m*, 4H), 3.53-3.38 (*m*, 2H), 3.32 (*dd*, *J*= 12.3, 11.4 Hz, 1H), 2.12 (*m*, 1H), 1.49 (*m*, 1H), 1.18 (*m*, 1H), 0.98 (*t*, *J*= 7.4 Hz, 3H). $^{13}$C-NMR (126 MHz, $CDCl_3$) $\delta$ ppm: 168.5, 167.2, 162.7, 156.0, 143.5, 130.4, 125.4, 122.6, 107.8, 107.1, 100.2, 62.2, 55.5, 54.6, 53.1, 51.7, 38.6, 31.6, 28.1, 23.2, 21.5, 11.8. The chemical shifts were in agreement with published data (Chakraborty *et al*. 2021).[12] Corresponding spectra are shown in Figure S56.

**dehydrocorynantheidine (DHC, 13a)**: $^1$H-NMR (500 MHz, $CDCl_3$) $\delta$ ppm: 7.80 (*brd*, *J*= 7.9 Hz, 1H), 7.54 (*d*, *J*= 8.1 Hz, 1H), 7.50 (*s*, 1H), 7.37 (*brdd*, *J*= 7.9, 7.6 Hz, 1H), 7.13 (*dd*, *J*= 8.1, 7.6 Hz, 1H), 3.98-3.82 (*m*, 3H), 3.77 (*s*, 3H), 3.74 (*m*, 1H), 3.61 (*s*, 3H), 3.65-3.54 (*m*, 2H), 3.37-3.26 (*m*, 3H), 2.10 (*m*, 1H), 1.49 (*m*, 1H), 1.17 (*m*, 1H), 0.99 (*t*, *J*= 7.3 Hz, 3H). $^{13}$C-NMR (126 MHz, $CDCl_3$) $\delta$ ppm: 168.5, 168.5, 162.6, 142.6, 128.5, 126.7, 123.9, 121.7, 120.6, 115.5, 108.1, 62.2, 54.7, 53.2, 51.7, 38.8, 32.8, 28.1, 23.3, 20.2, 11.8. Detailed assignment shown in Figures S57-S61.

**7-Hydroxymitragynine (24a)**: $^1$H NMR (500 MHz, $CDCl_3$) $\delta$ 7.44 (s, 1H), 7.30 (t, $J$ = 8.0 Hz, 1H), 7.21 (d, $J$ = 7.6 Hz, 1H), 6.74 (d, $J$ = 8.3 Hz, 1H), 3.87 (s, 3H), 3.81 (s, 3H), 3.70 (s, 3H), 3.13–3.09 (m, 1H), 3.05–3.00 (m, 2H), 2.82–2.77 (m, 2H), 2.64–2.62 (m, 2H), 2.48 (dd, $J$ = 11.3, 2.3 Hz, 1H), 2.16 (s, 1H), 1.89 (d, $J$ = 13.6 Hz, 1H), 1.74–1.65 (m, 2H), 1.42 (t, $J$ = 7.3 Hz, 2H), 0.82 (t, $J$ = 7.3 Hz, 3H). Spectral data agreed with those reported previously.[20] Detailed assignment shown in Figures S64.

**Dehydromitragynine trifluoroacetate (TFA) (14a)**: $^1$H NMR (400 MHz, $CDCl_3$)* $\delta$ 11.61 (s, 1H), 7.52 (s, 1H), 7.30 (t, $J$ = 8.1 Hz, 1H), 7.12 (d, $J$ = 8.4 Hz, 1H), 6.43 (d, $J$ = 7.7 Hz, 1H), 3.91 (s, 3H), 3.90–3.84 (m, 2H), 3.78 (s, 3H), 3.62 (s, 3H), 3.60–3.55 (m, 3H), 3.54–3.42 (m, 3H), 3.27 (t, $J$ = 12.6 Hz, 1H), 2.15–2.12 (appbrs, 1H), 1.55–1.44 (m, 1H), 1.20–1.13 (m, 1H), 0.99 (t, $J$ = 7.4 Hz, 3H). Spectral data agreed with those reported previously.[12] *Note: The presence of residual methylene chloride[21] and triethylammonium trifluoroacetate[22] was confirmed upon comparison of $^1$H NMR obtained for **14a** with previously reported spectral data. Detailed assignment shown in Figures S65.

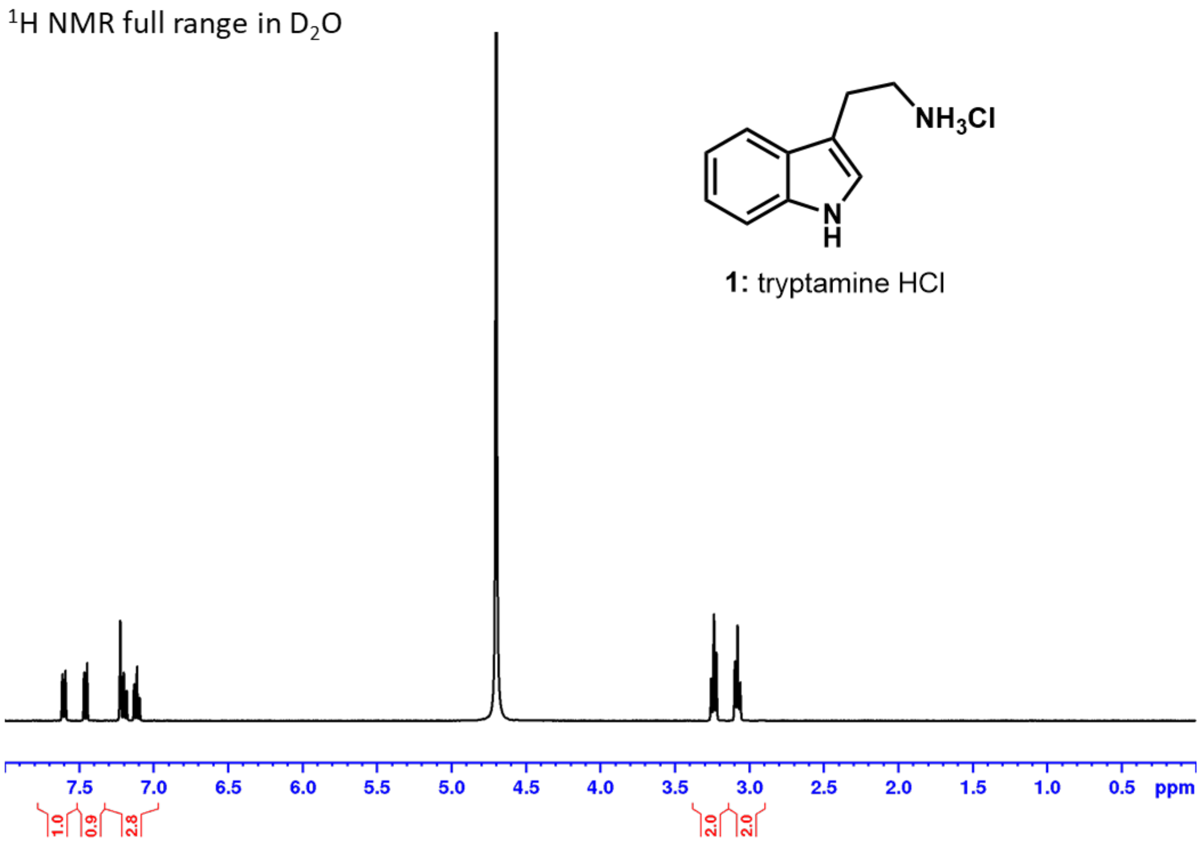


**Figure S38:** NMR spectrum for **1:** tryptamine-HCl.

<sup></sup>$^1$H NMR full range in $D_2O$

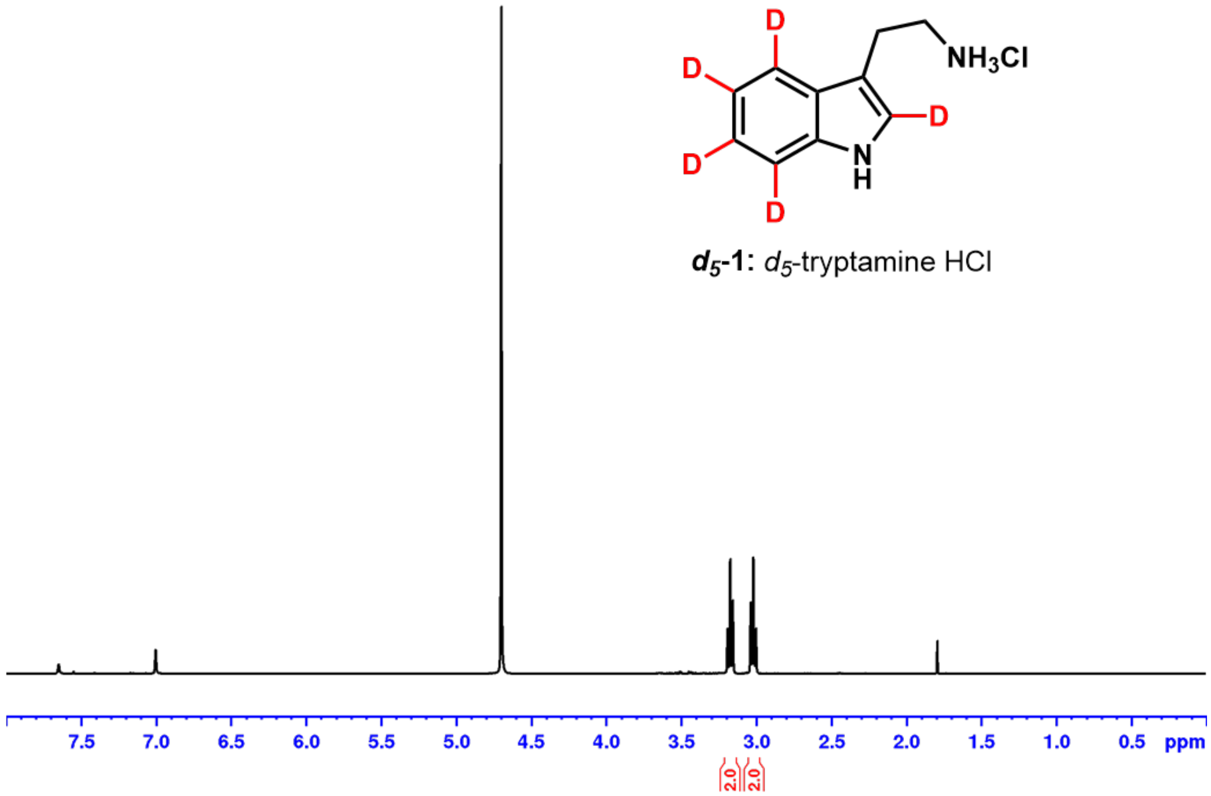


**Figure S39:** NMR spectrum for ***d*$_5$-1:** *d*$_5$-tryptamine-HCl.

$^1H$ NMR full range in $CDCl_3$

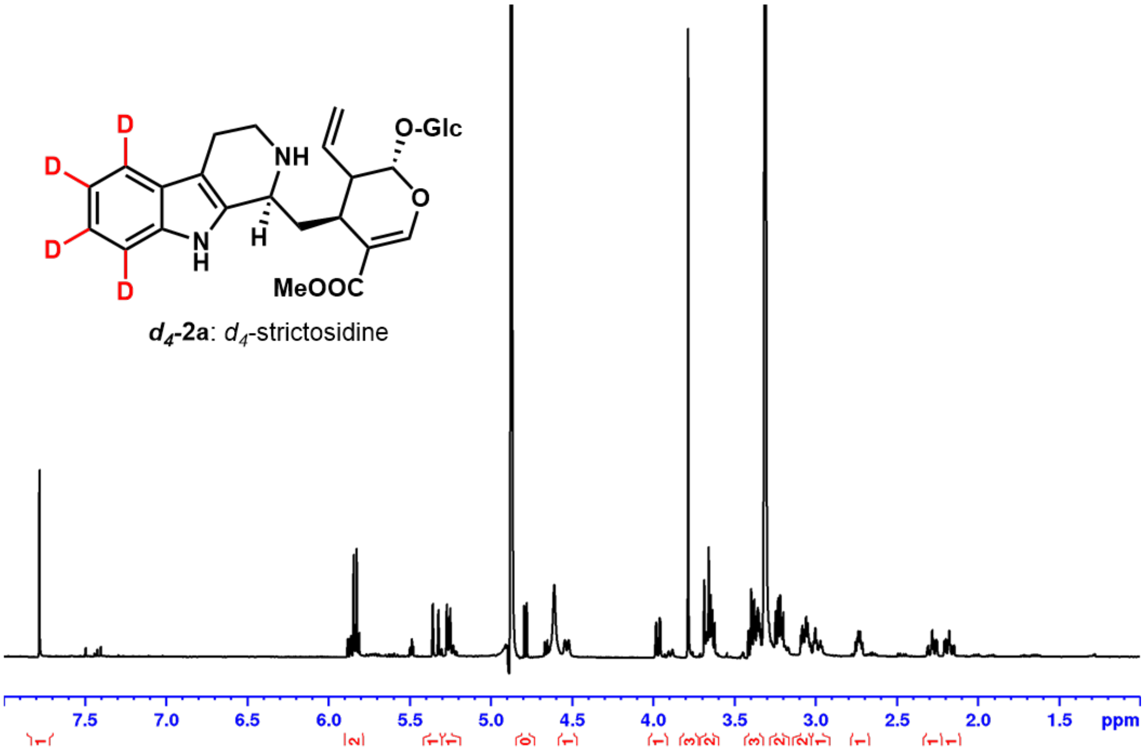


DEPTQ full range in $CDCl_3$

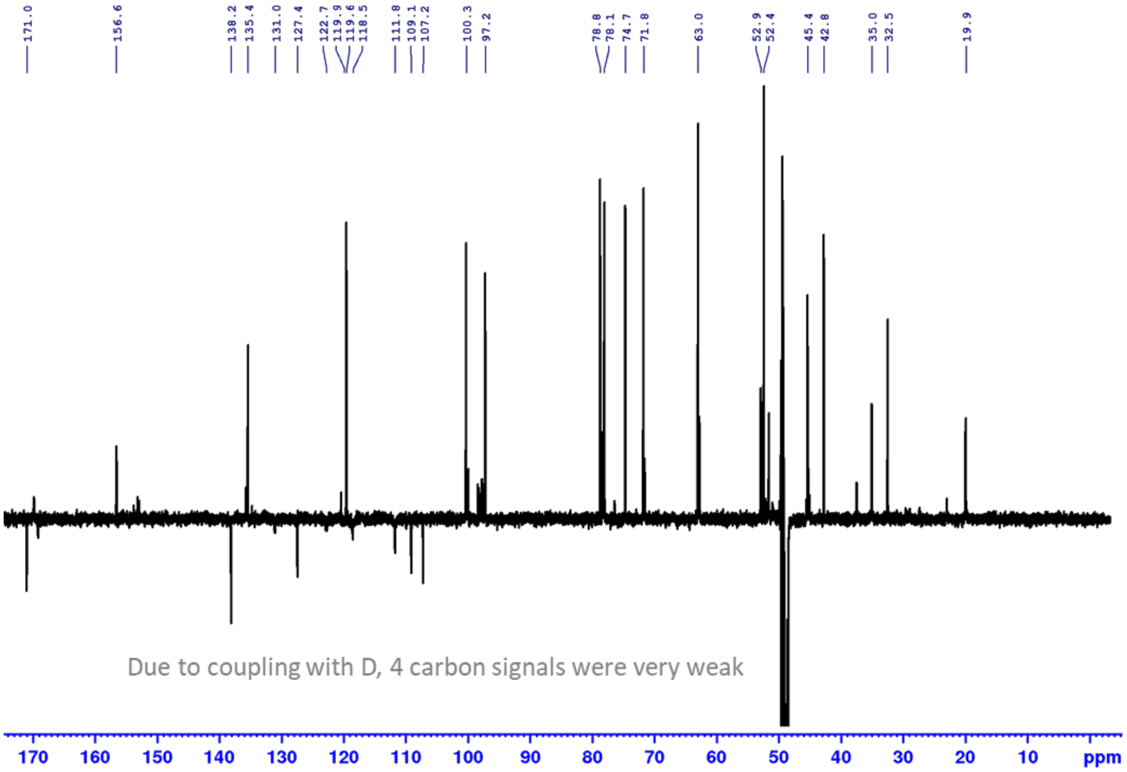


**Figure S40:** NMR spectrum for $d_4$-strictosidine (***d*$_4$-2a**).

¹H NMR full range in $D_2O$

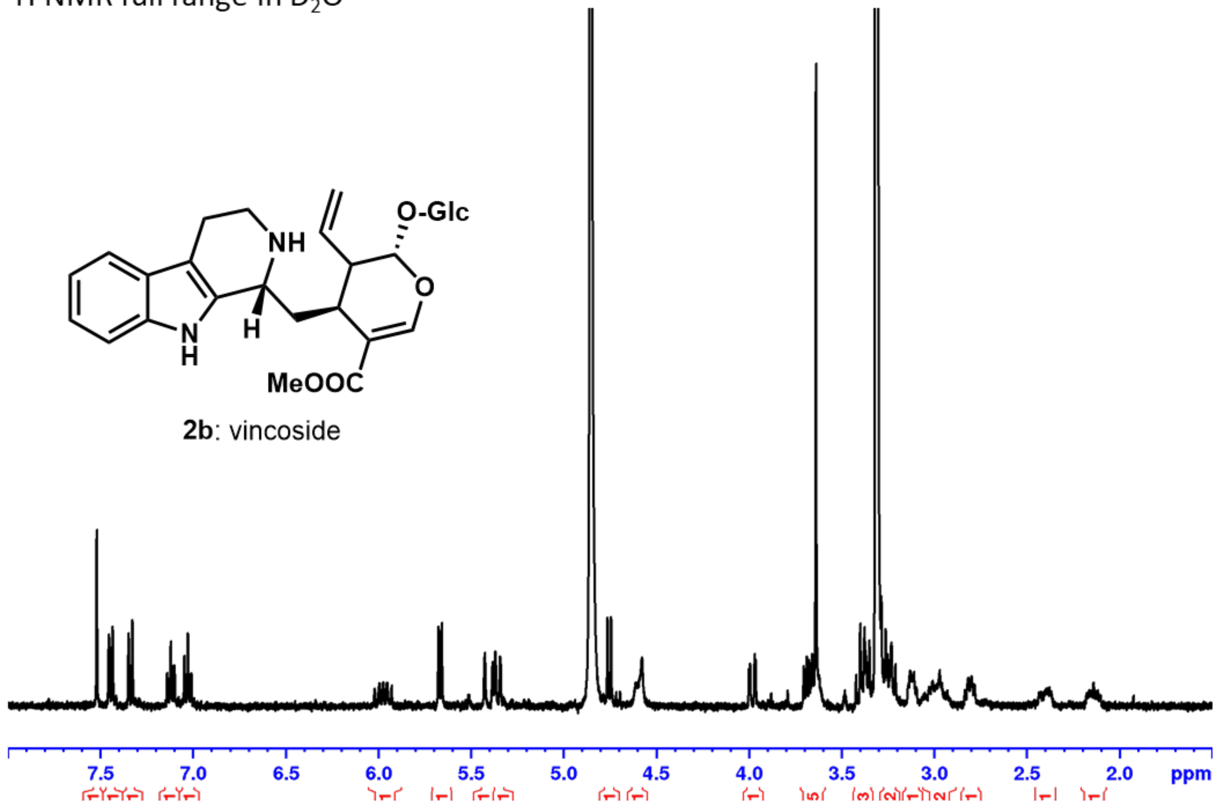


**Figure S41:** NMR spectrum for vincoside (**2b**).

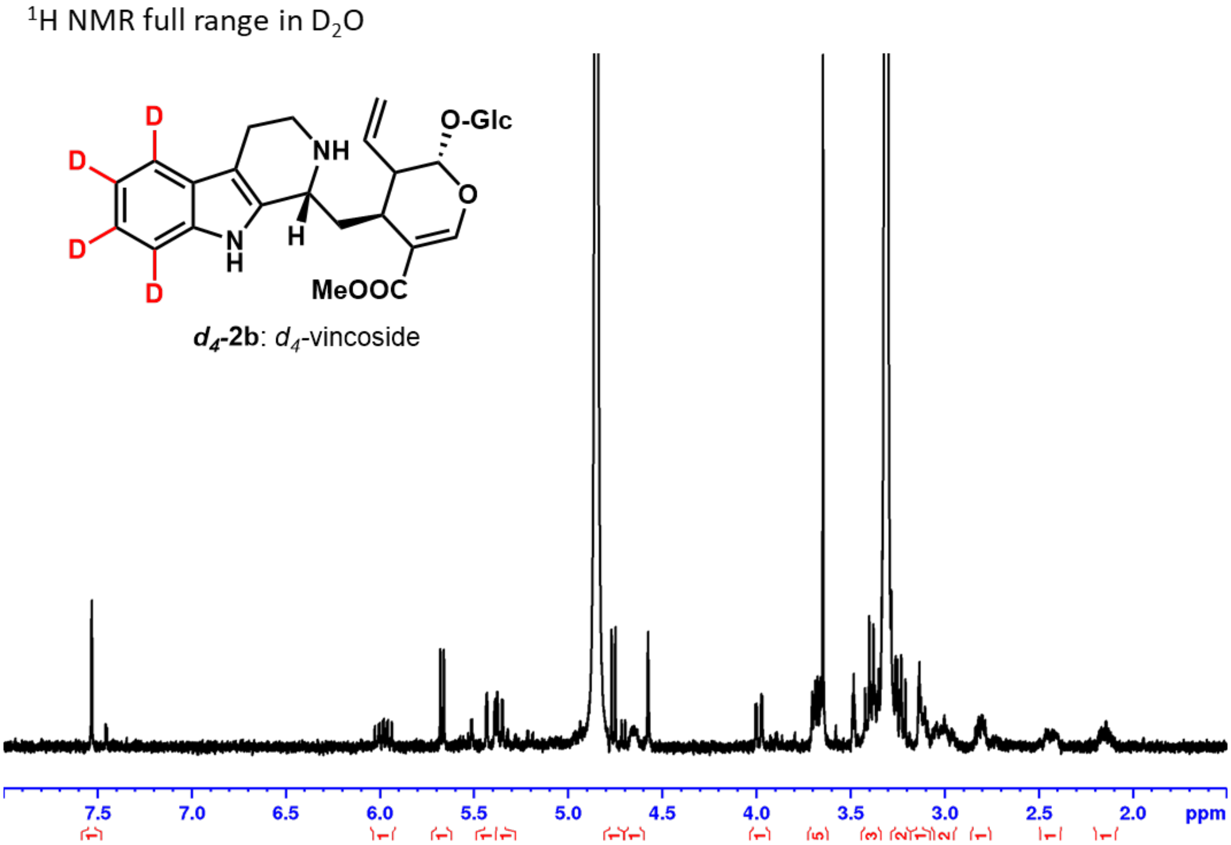


**Figure S42:** NMR spectrum for $d_4$-vincoside (***$d_4$*-2b**).

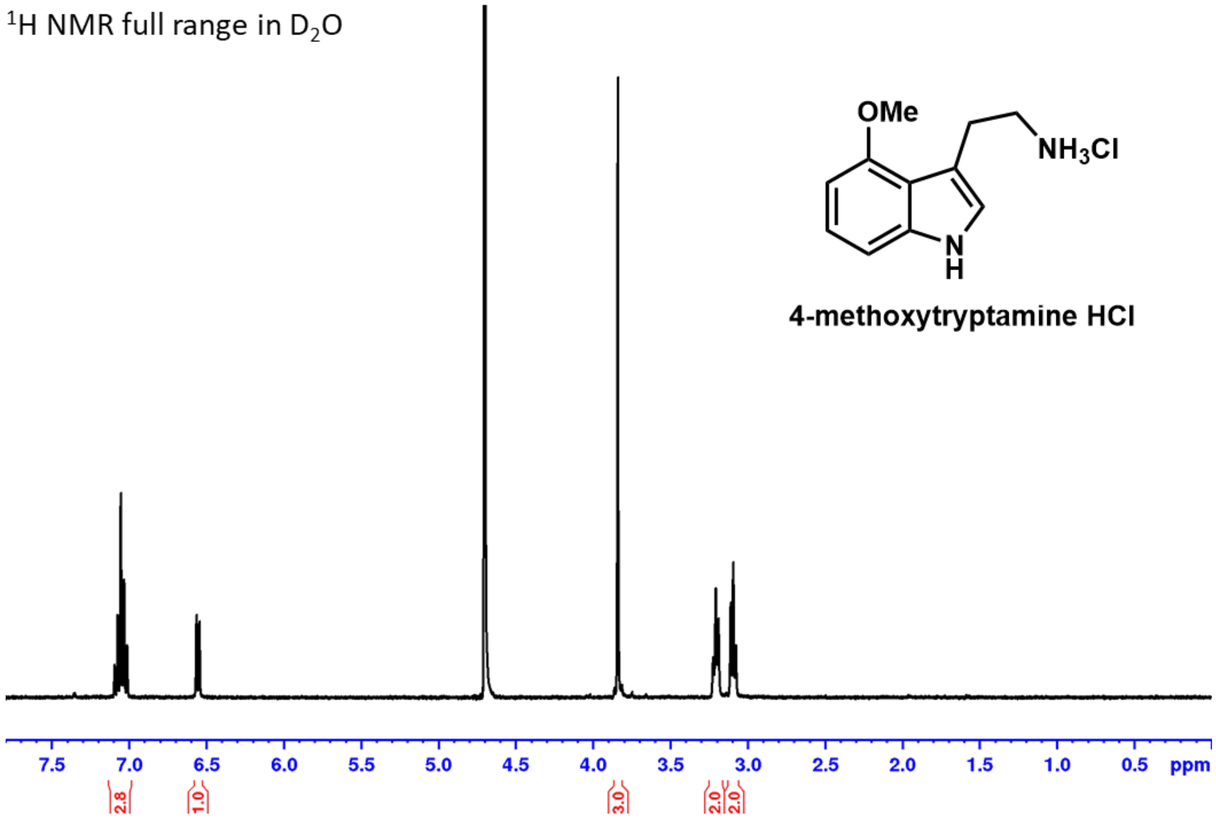


**Figure S43:** NMR spectrum for 4-methoxytryptamine-HCl.

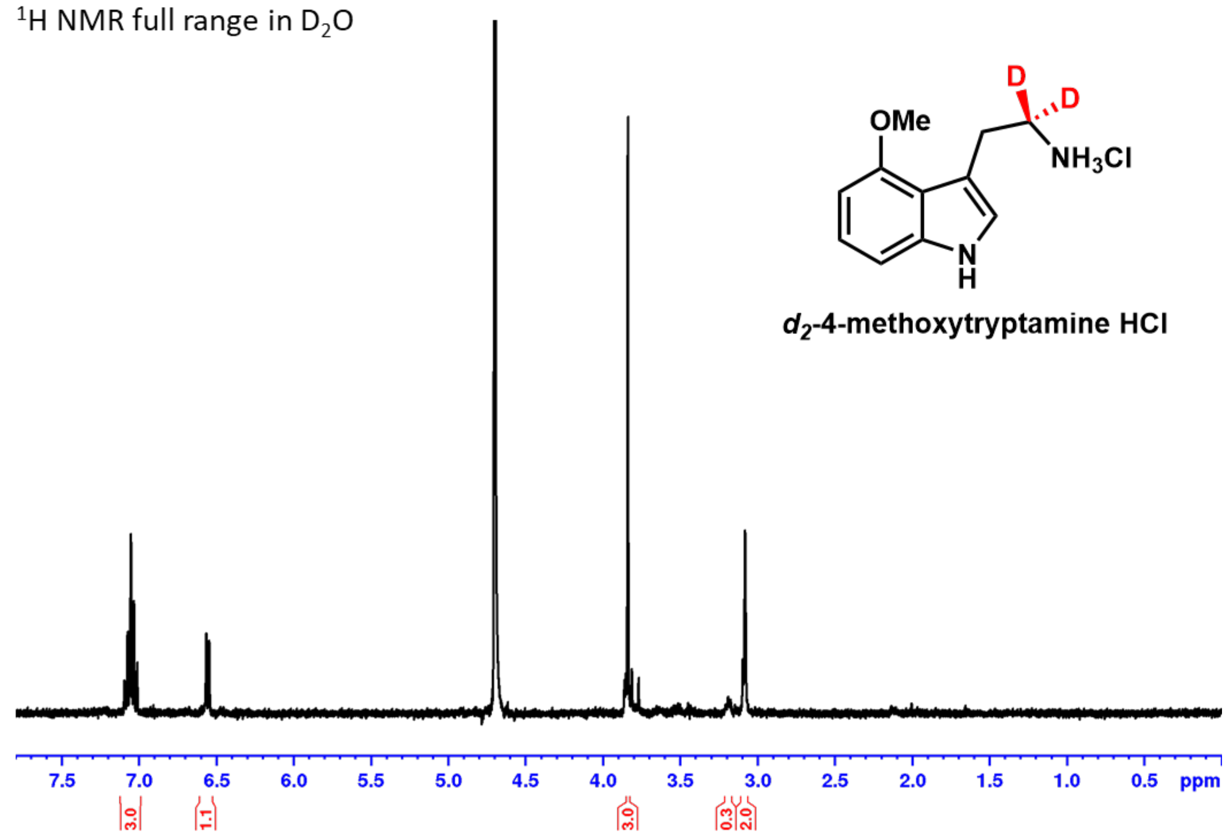


**Figure S44:** NMR spectrum for $d_2$-4-methoxytryptamine-HCl.

$^1$H NMR with water presaturation full range in MeOH-$d_3$

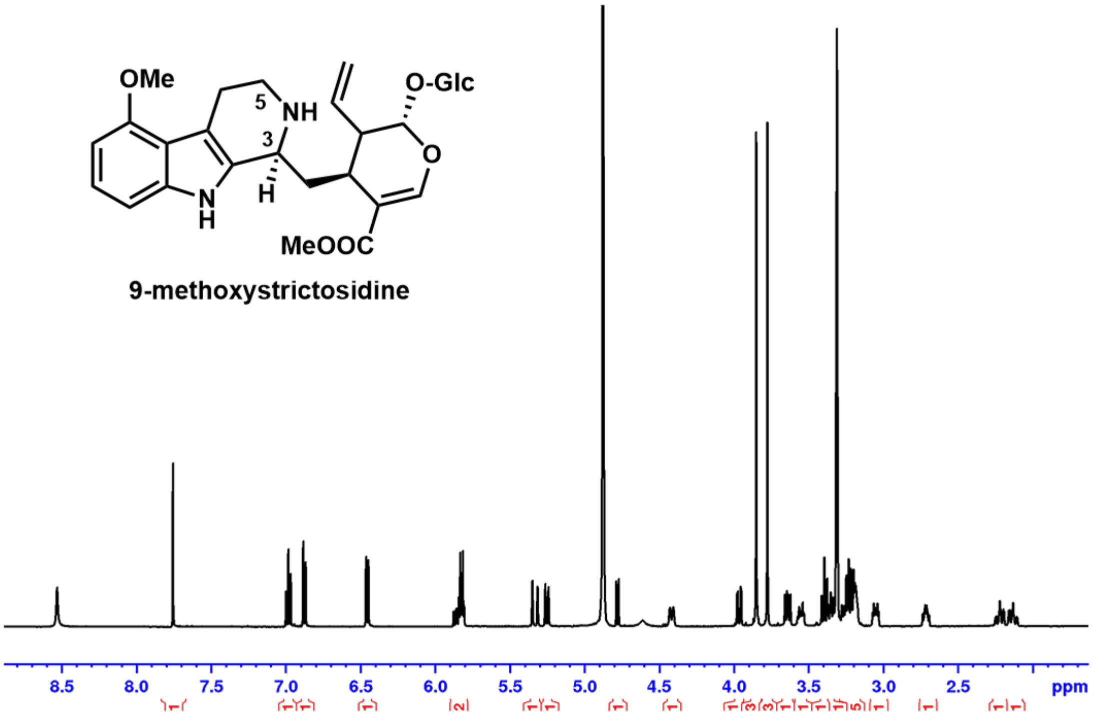


DEPTQ full range in MeOH-$d_3$

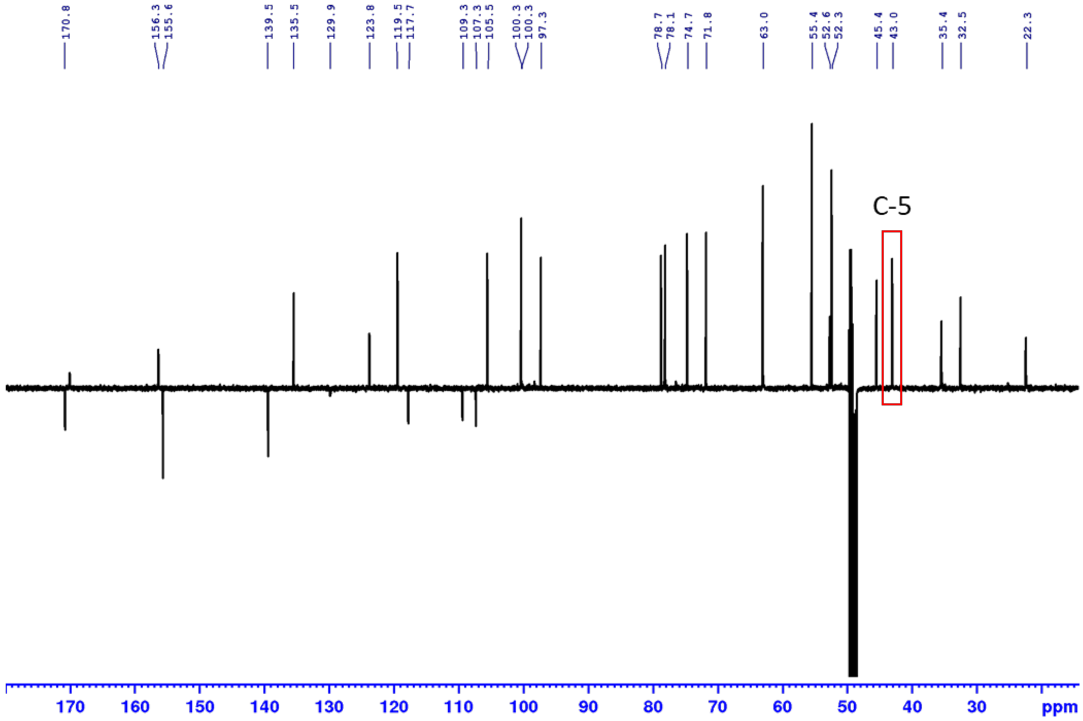


**Figure S45:** NMR spectra for 9-methoxystrictosidine.

[1]H NMR with water presaturation full range in MeOH-$d_3$

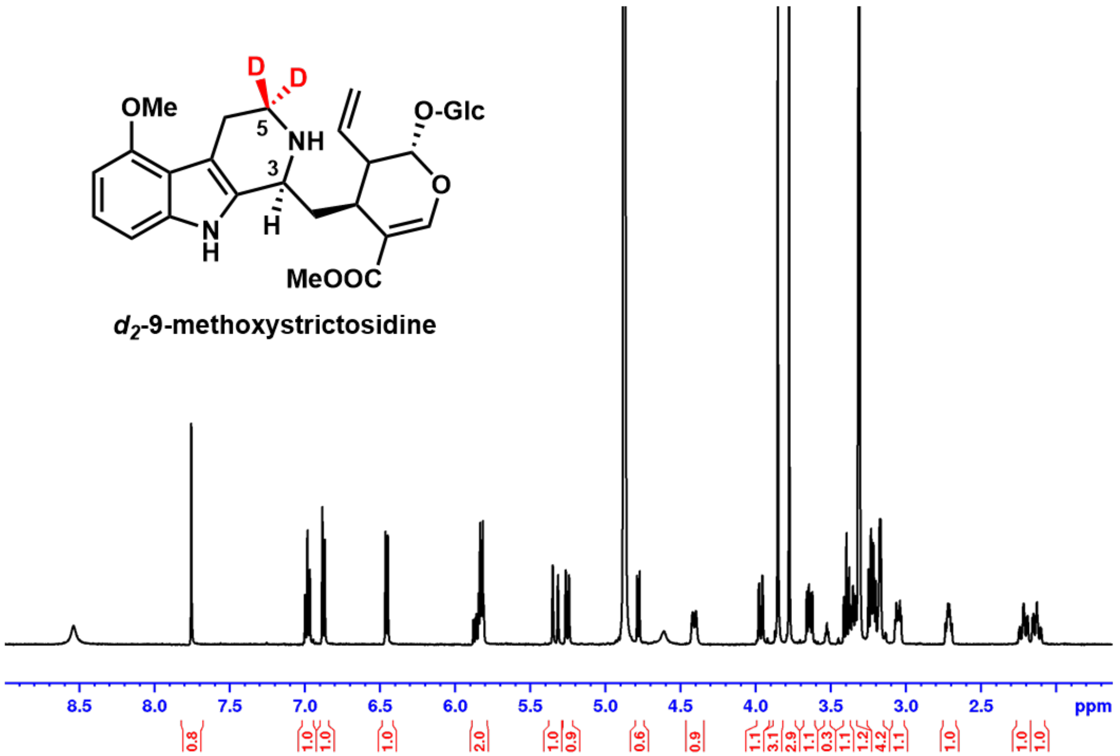


DEPTQ full range in MeOH-$d_3$

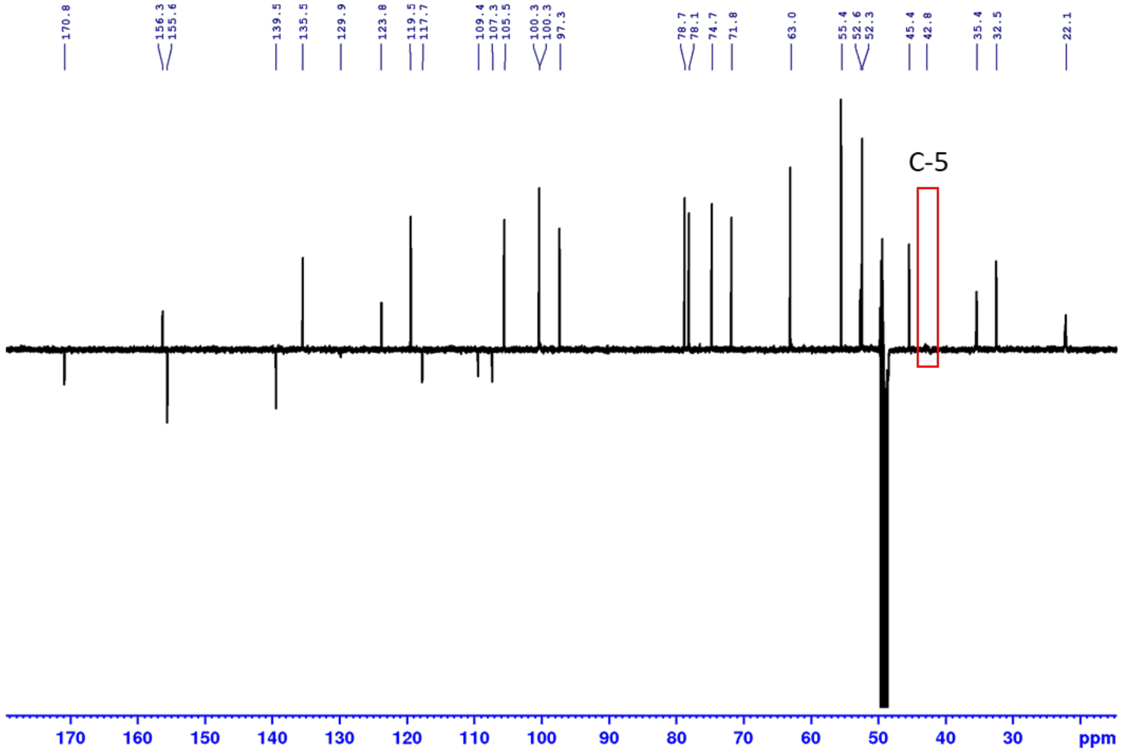


**Figure S46:** NMR spectra for $d_2$-9-methoxystrictosidine.

$^{1}$H NMR with water presaturation of 9-OMe-strictosidine (3-4 ppm)

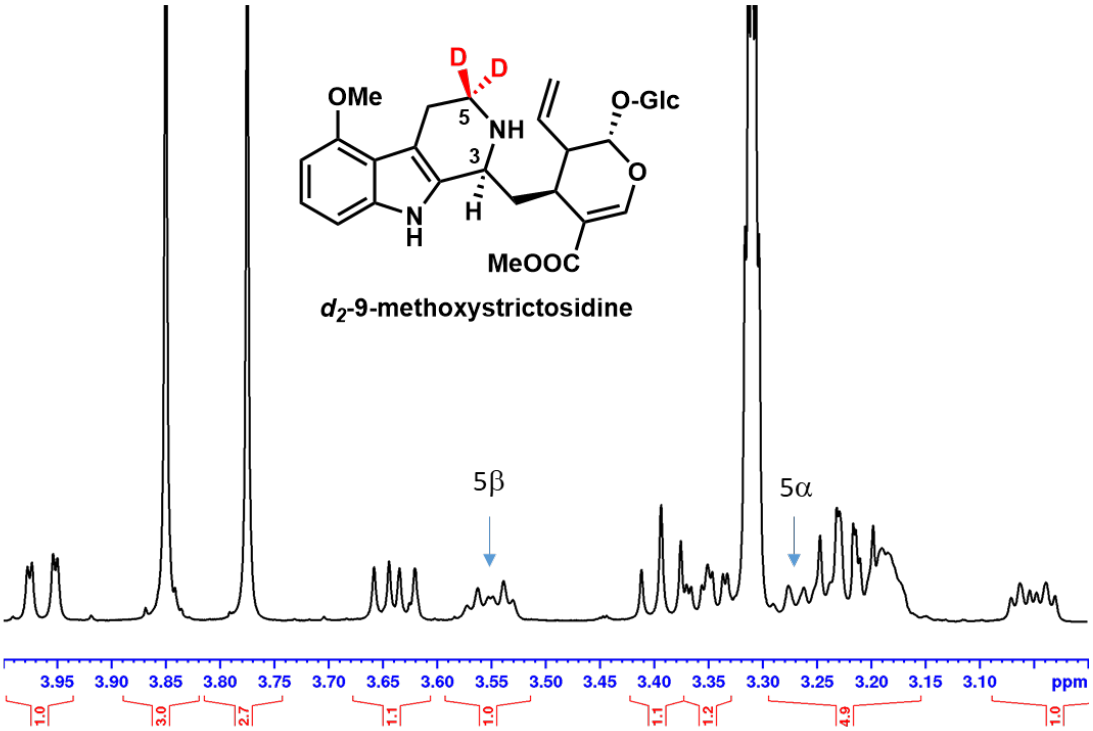


$^{1}$H NMR with water presaturation of $d_2$-9-OMe-strictosidine (3-4 ppm)

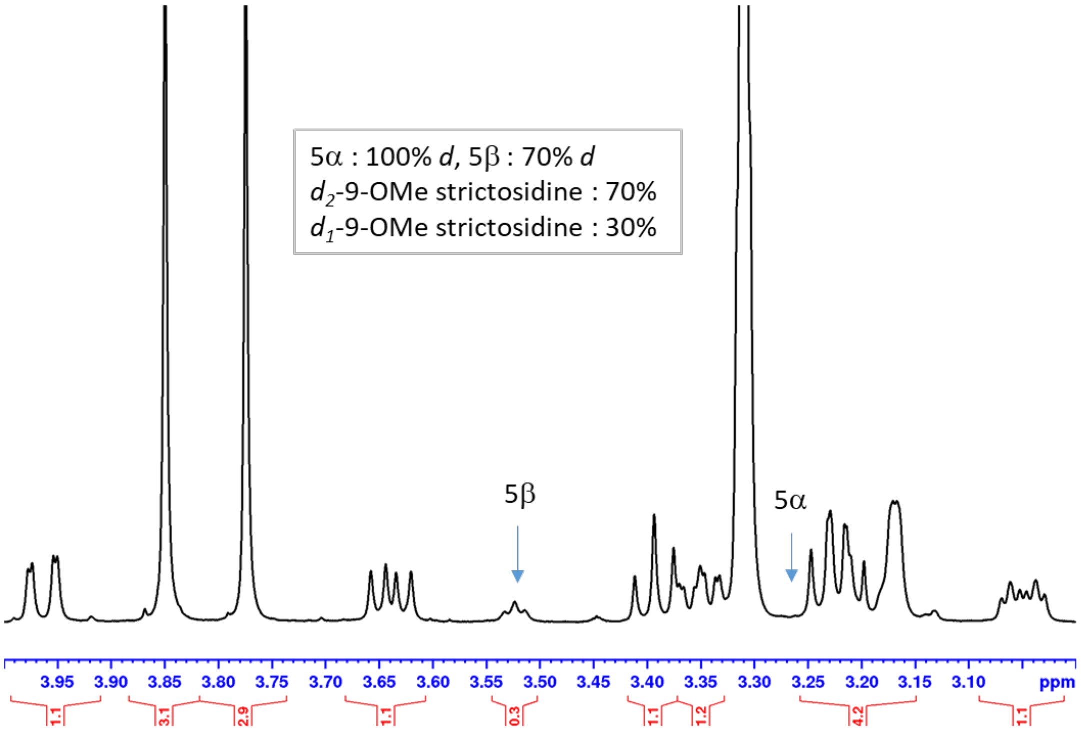


**Figure S47:** NMR spectrum for $d_2$-9-methoxystrictosidine.

$^{1}H$ NMR full range in $CDCl_3$

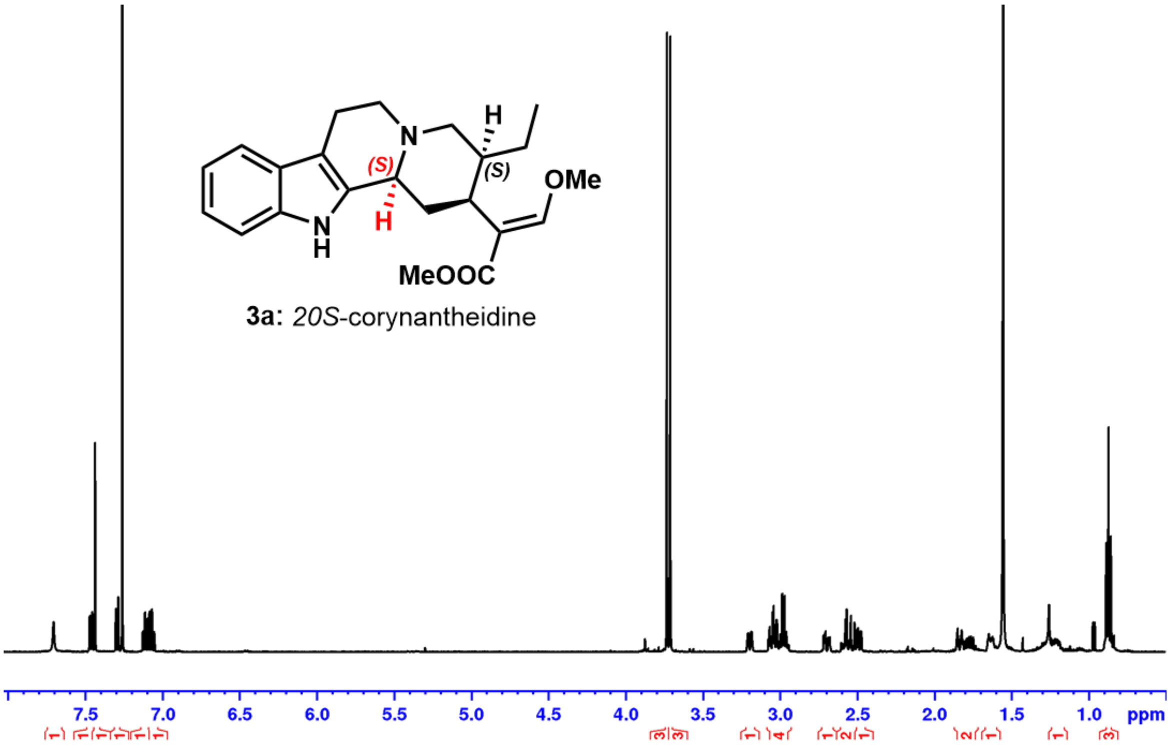


DEPTQ full range in $CDCl_3$

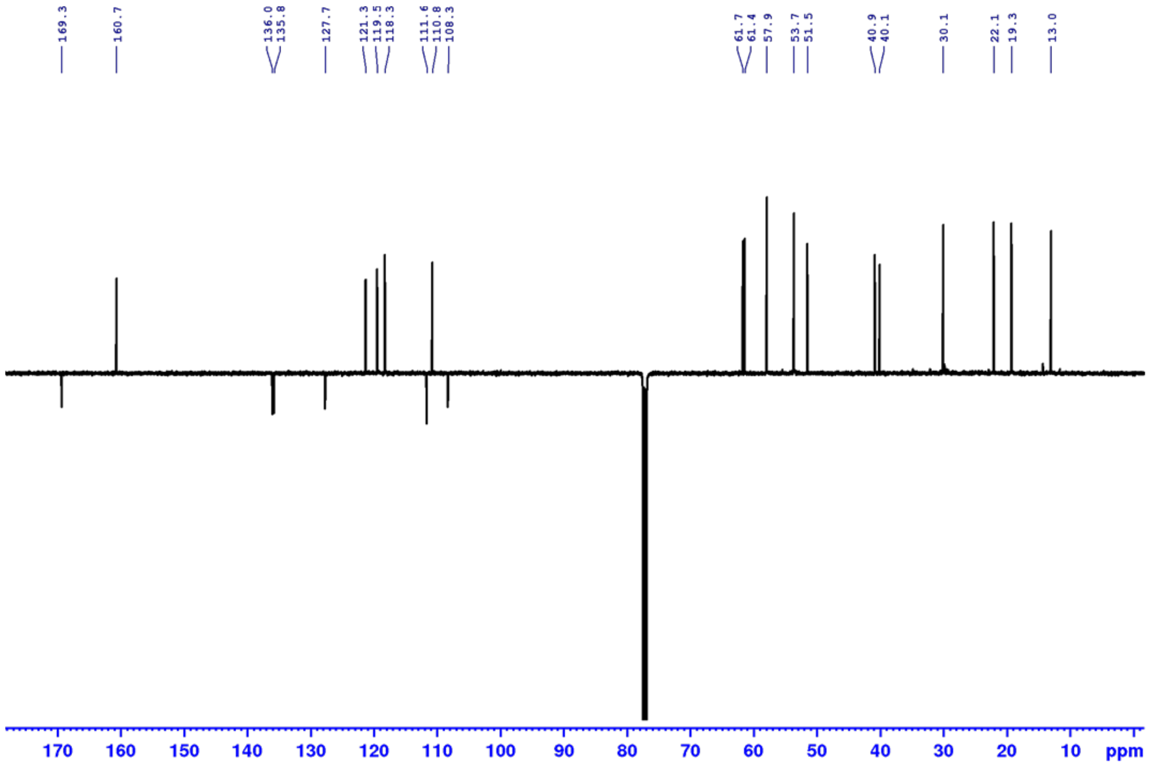


**Figure S48:** NMR spectra for **3a:** 20*S*-corynantheidine.

$^{1}H$ NMR full range in $CDCl_3$

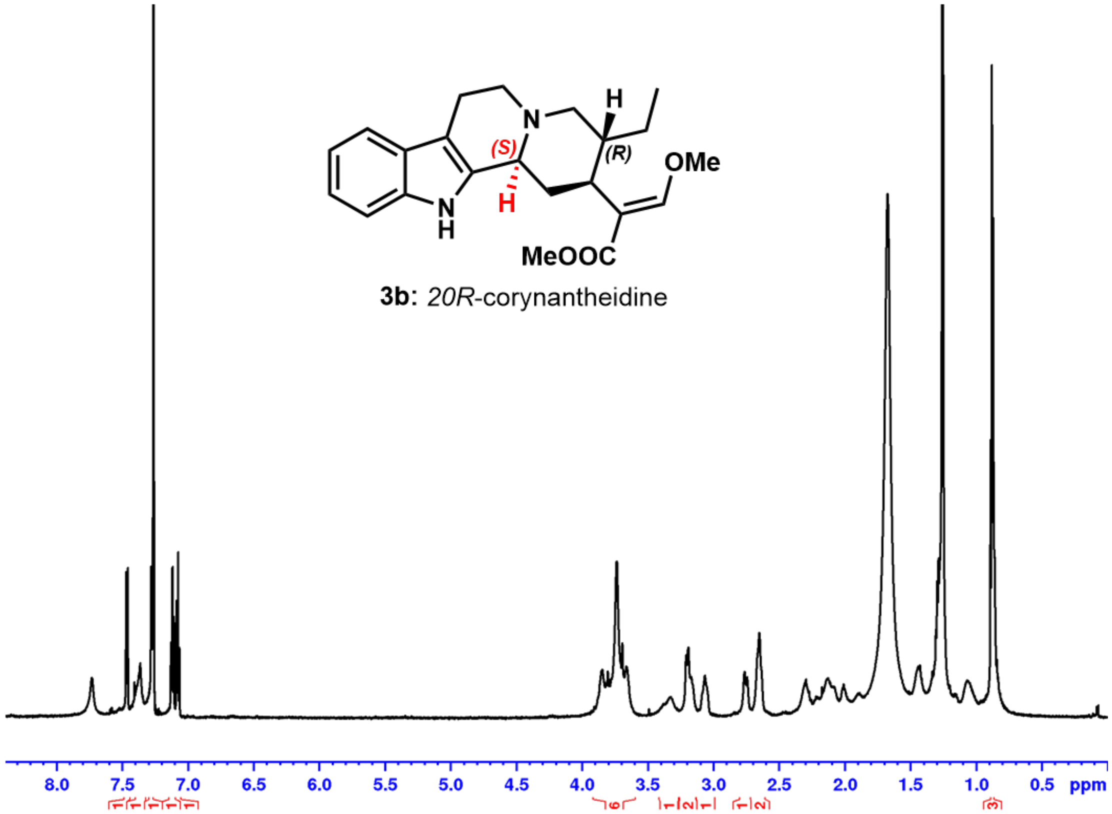


Phase sensitive HSQC (blue: CH, $CH_3$, green $CH_2$)/ HMBC (red) full range in $CDCl_3$

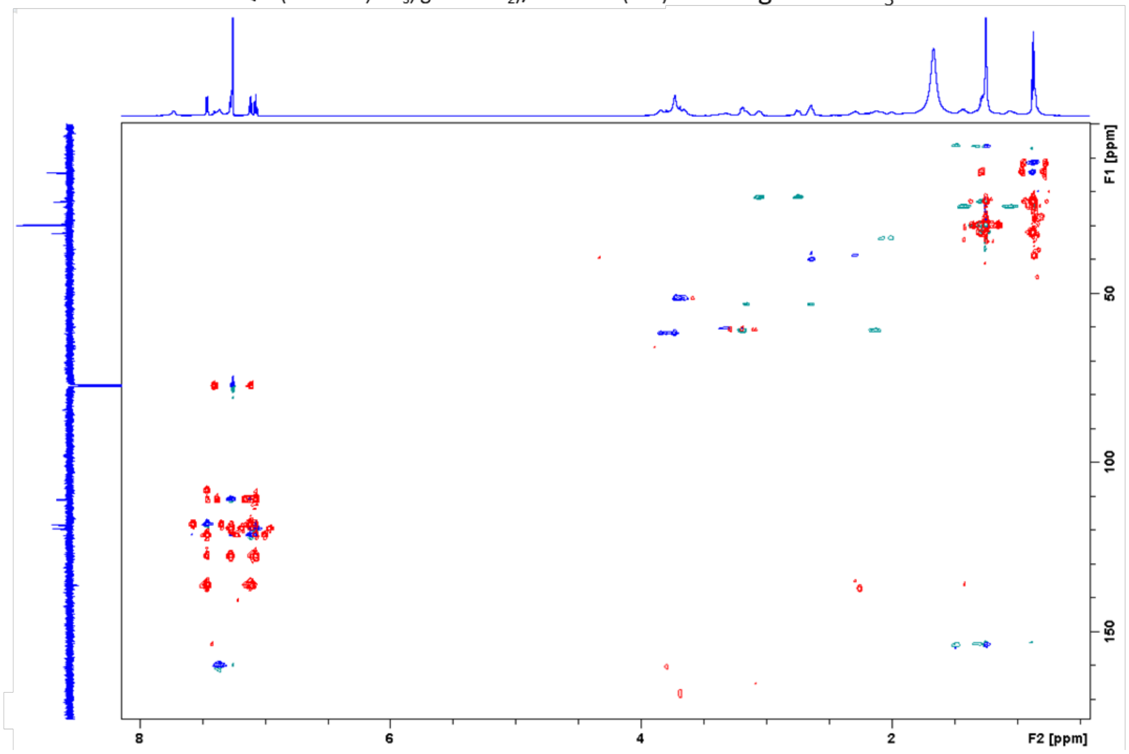


**Figure S49:** NMR spectra for **3b:** 20*R*-corynantheidine.

$^1H$ NMR full range in $CDCl_3$

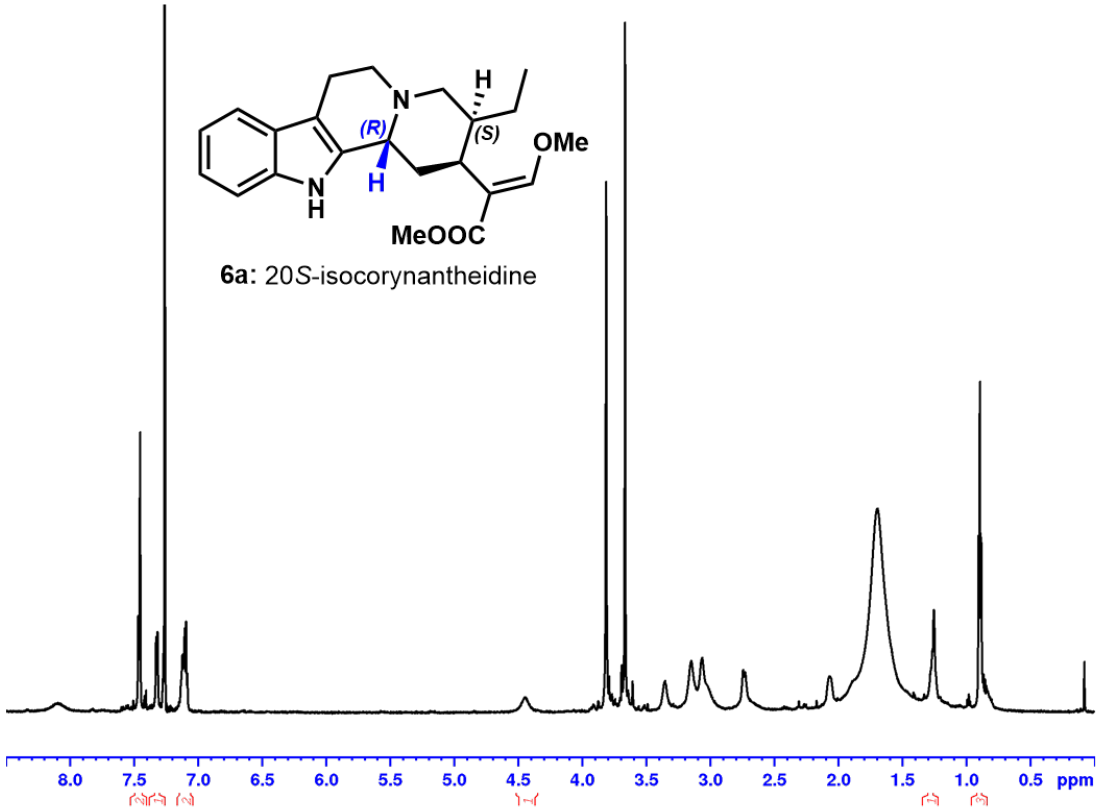


Phase sensitive HSQC (blue: CH, $CH_3$, green $CH_2$)/ HMBC (red) full range in $CDCl_3$

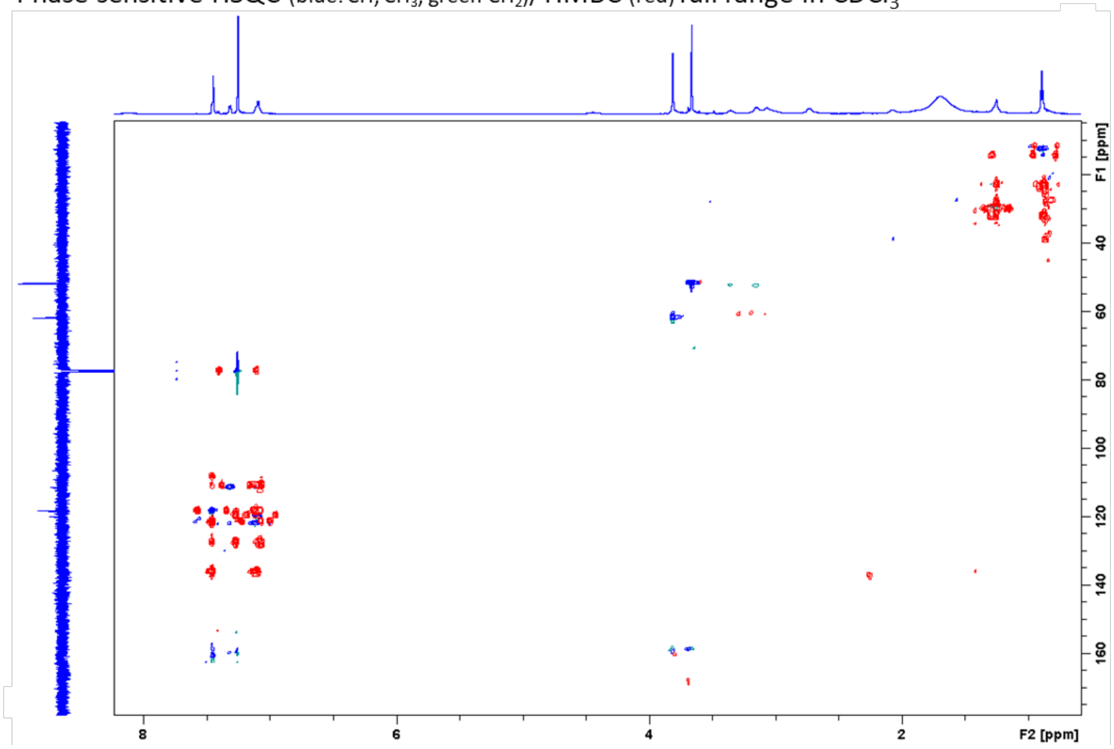


**Figure S50:** NMR spectra for **6a:** 20*S*-isocorynantheidine.

$^{1}H$ NMR full range in $CDCl_3$

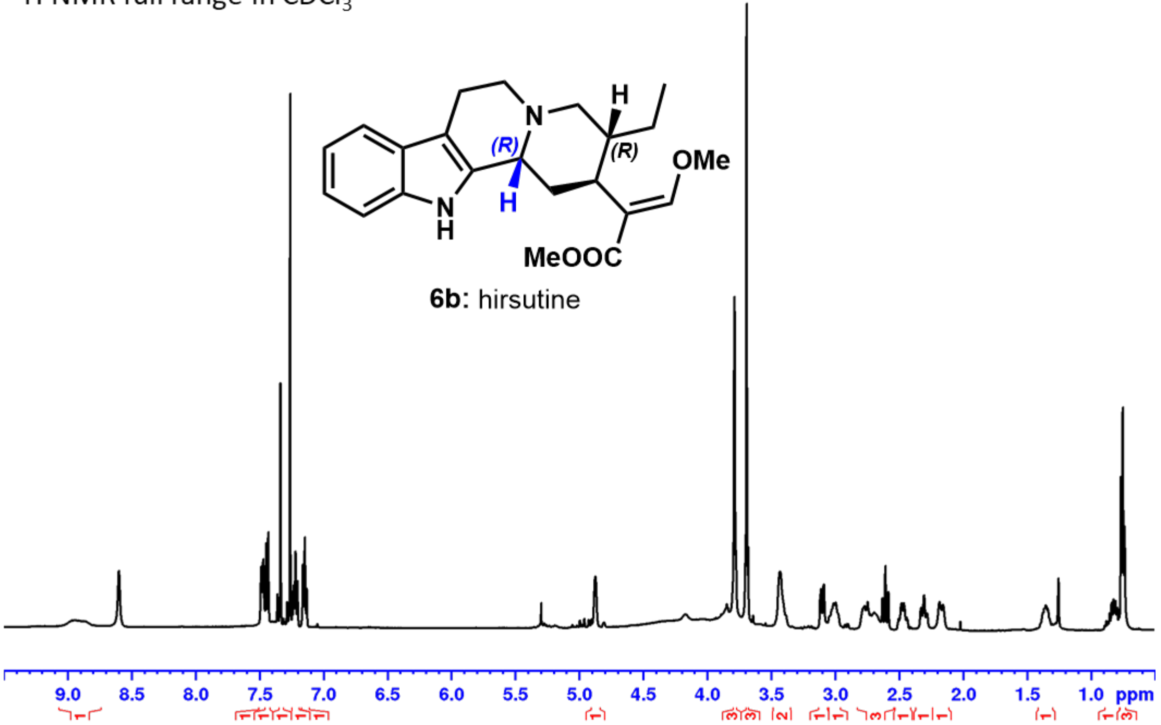


DEPTQ full range in $CDCl_3$

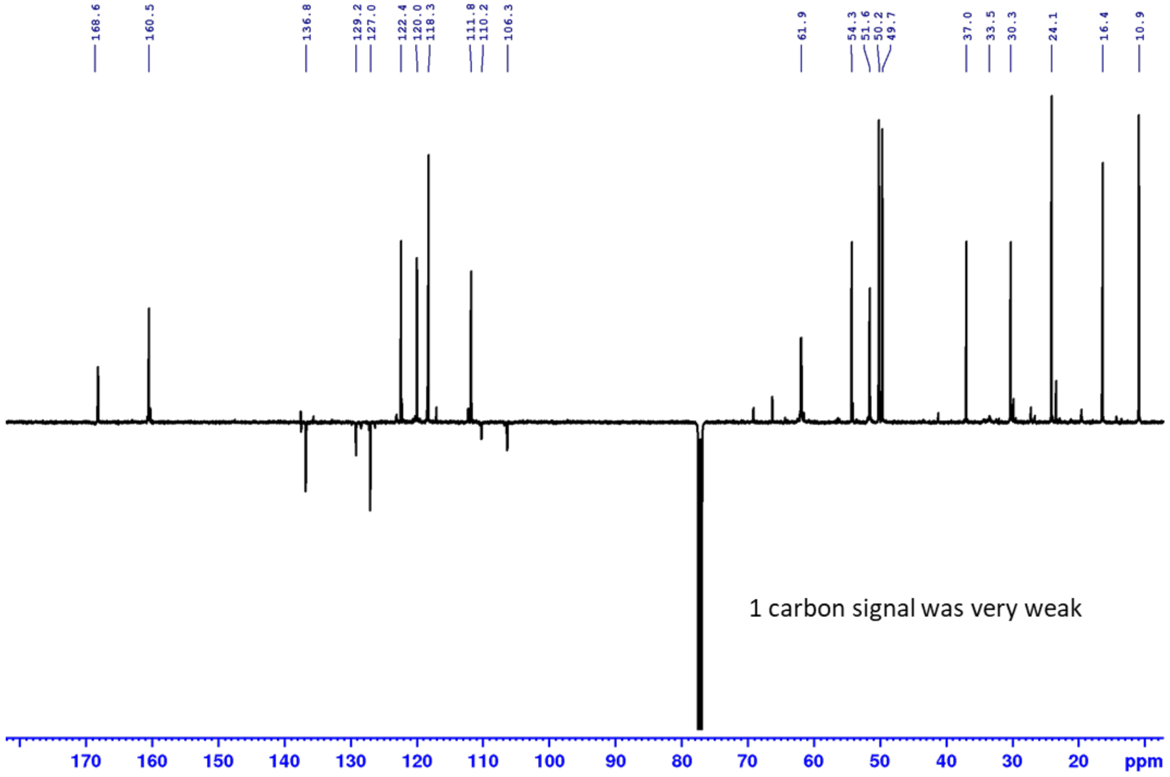


**Figure S51:** NMR spectra for **6b:** hirsutine.

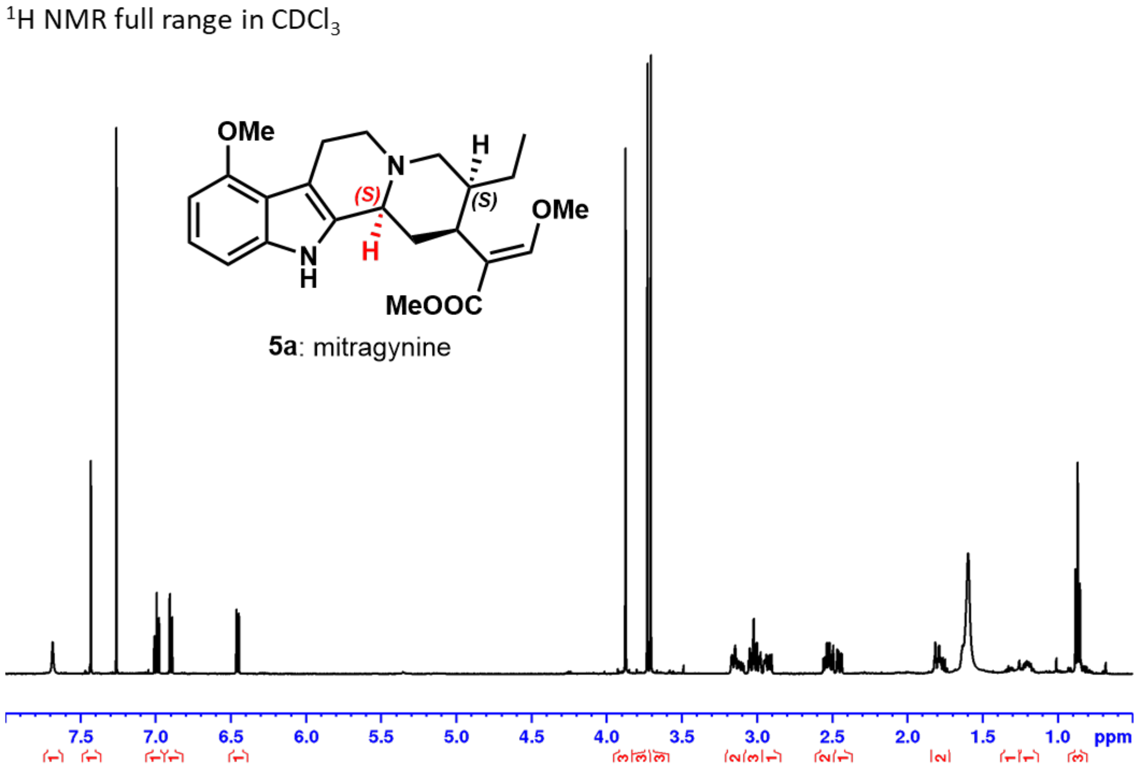


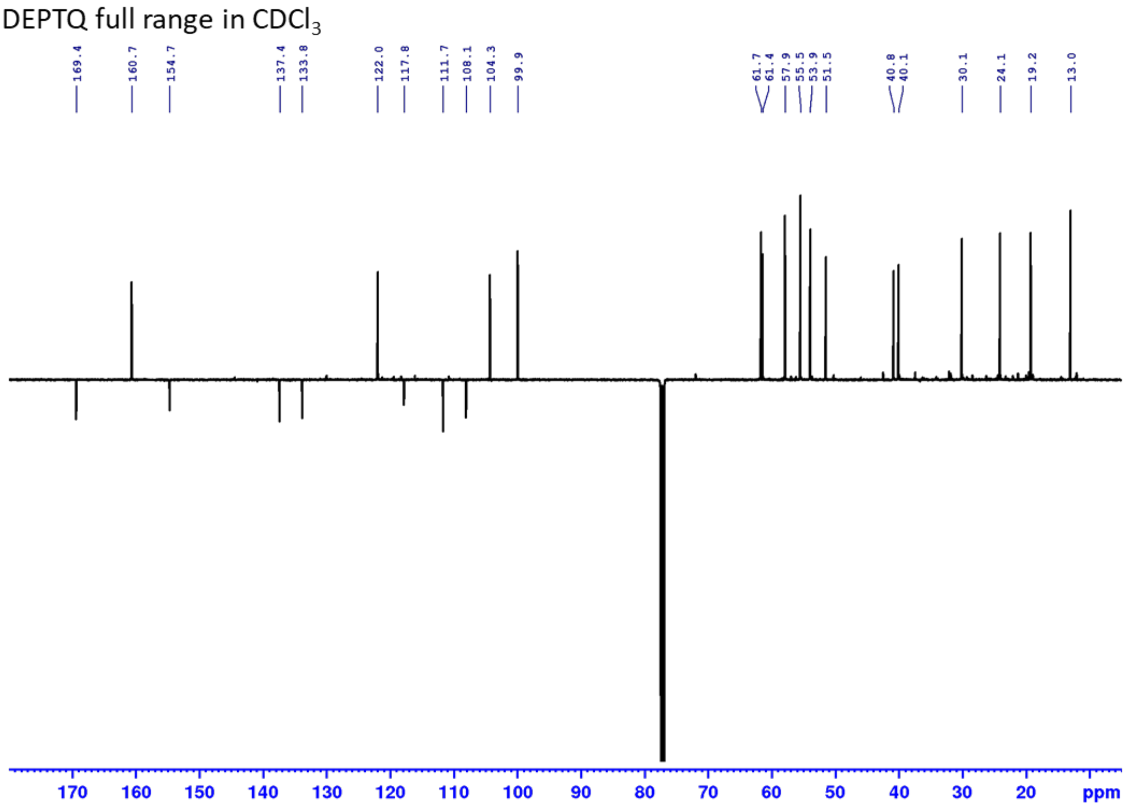


**Figure S52:** NMR spectra for **5a:** mitragynine.

$^1H$ NMR full range in $CDCl_3$

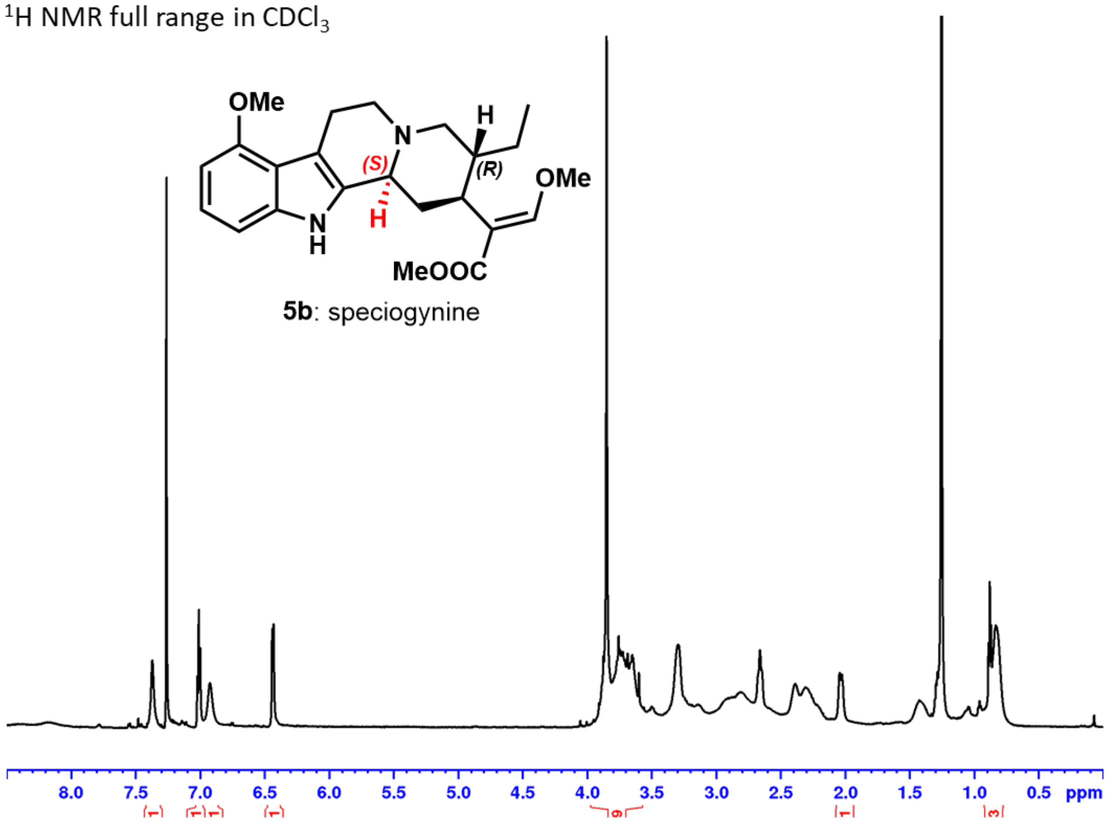


Phase sensitive HSQC (blue: CH, $CH_3$, green $CH_2$)/ HMBC (red) full range in $CDCl_3$

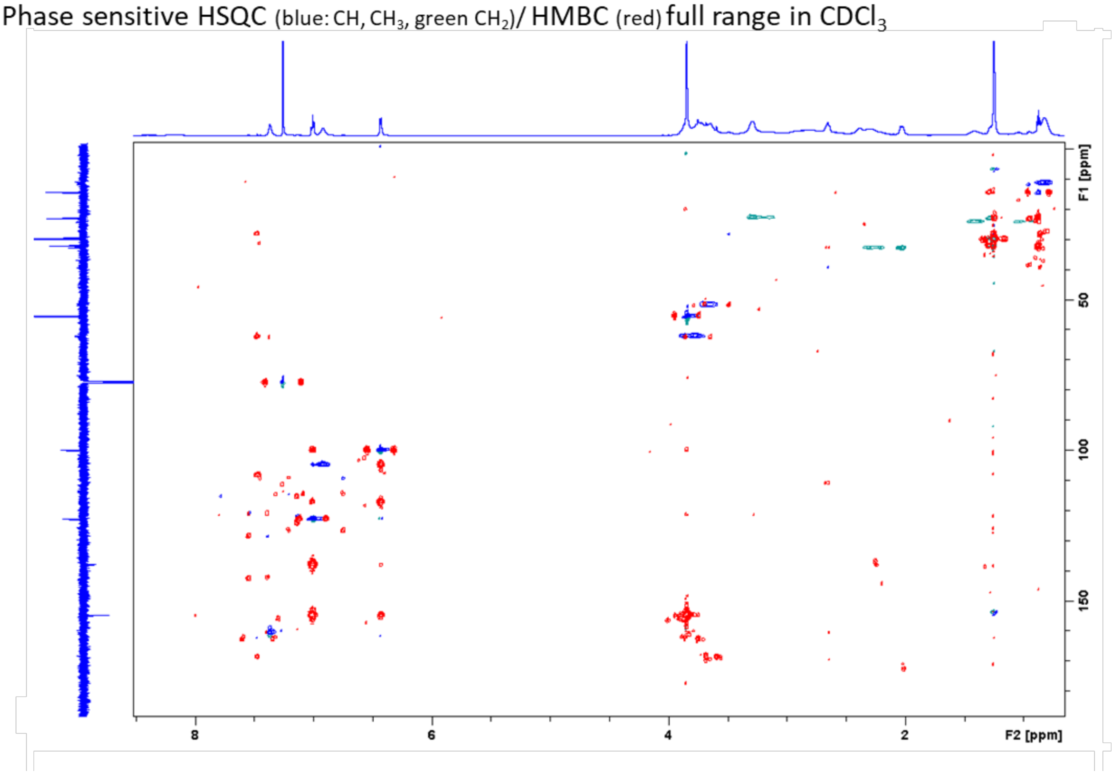


**Figure S53:** NMR spectra for **5b:** speciogynine.

$^{1}H$ NMR full range in $CDCl_3$

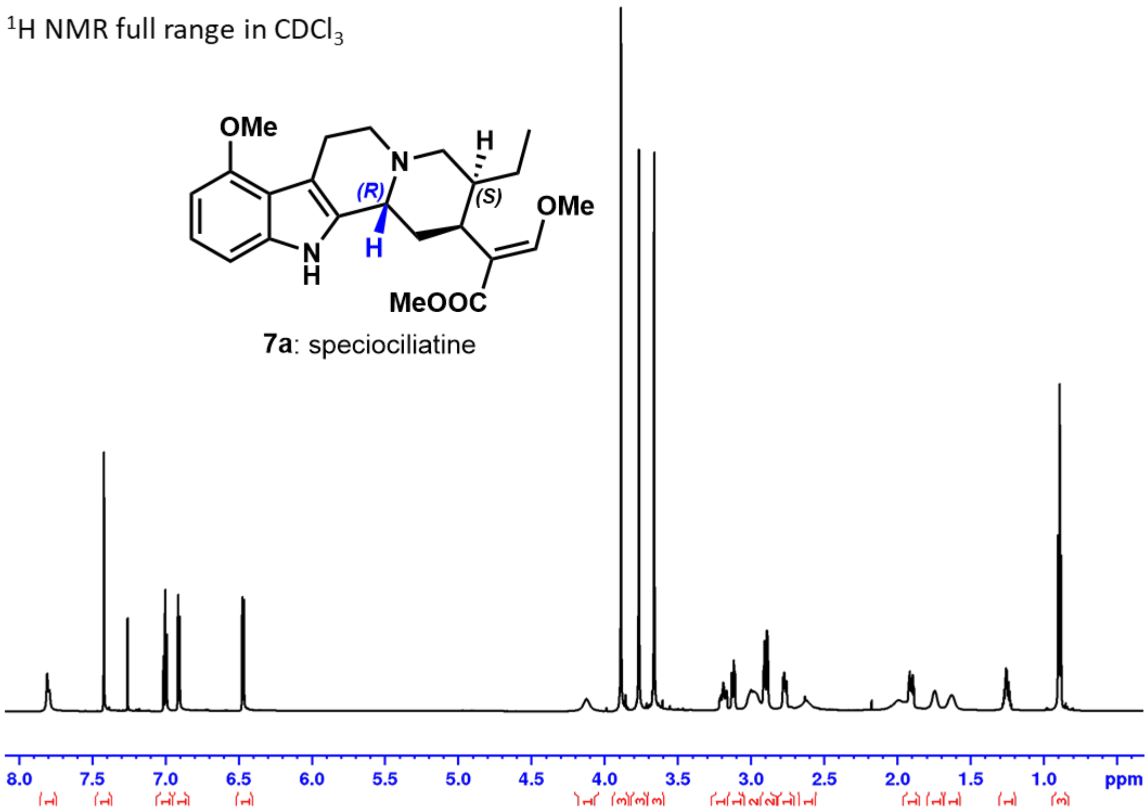


Phase sensitive HSQC (blue: CH, $CH_3$, green $CH_2$)/ HMBC (red) full range in $CDCl_3$

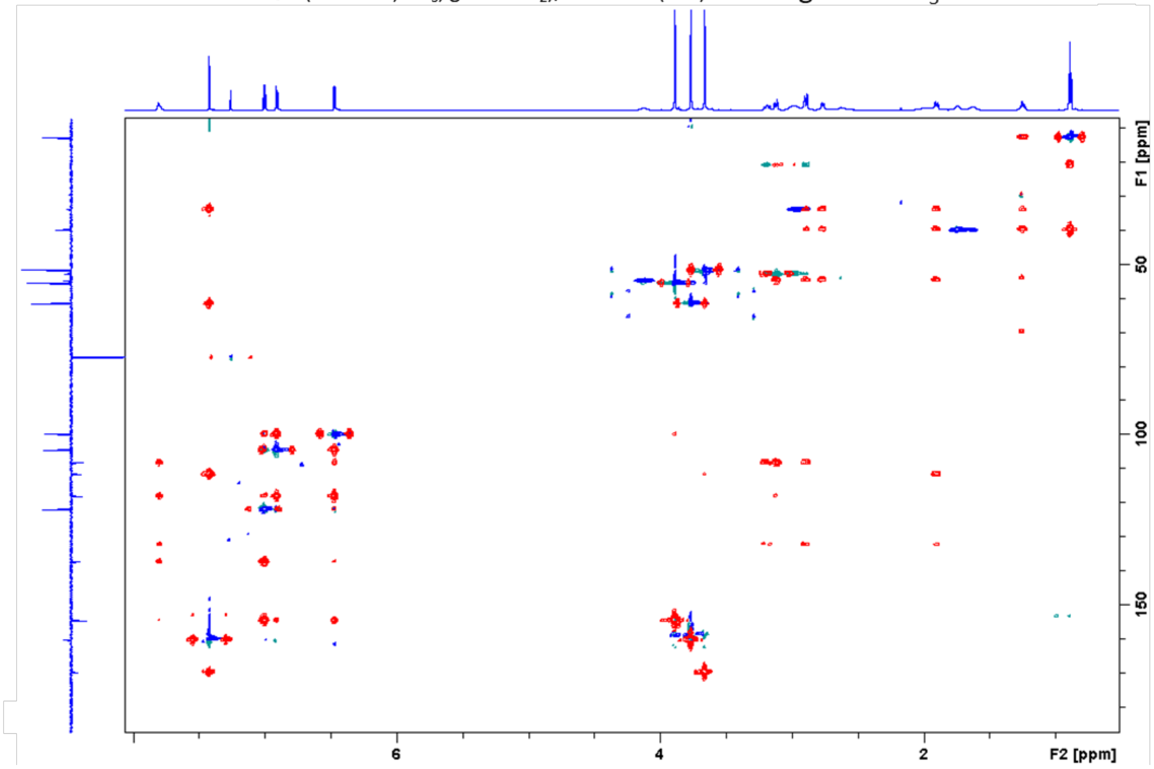


**Figure S54:** NMR spectra for **7a:** speciociliatine.

**Figure SX-1** NMR spectra for **7b:** mitraciliatine

$^1H$ NMR full range in $CDCl_3$

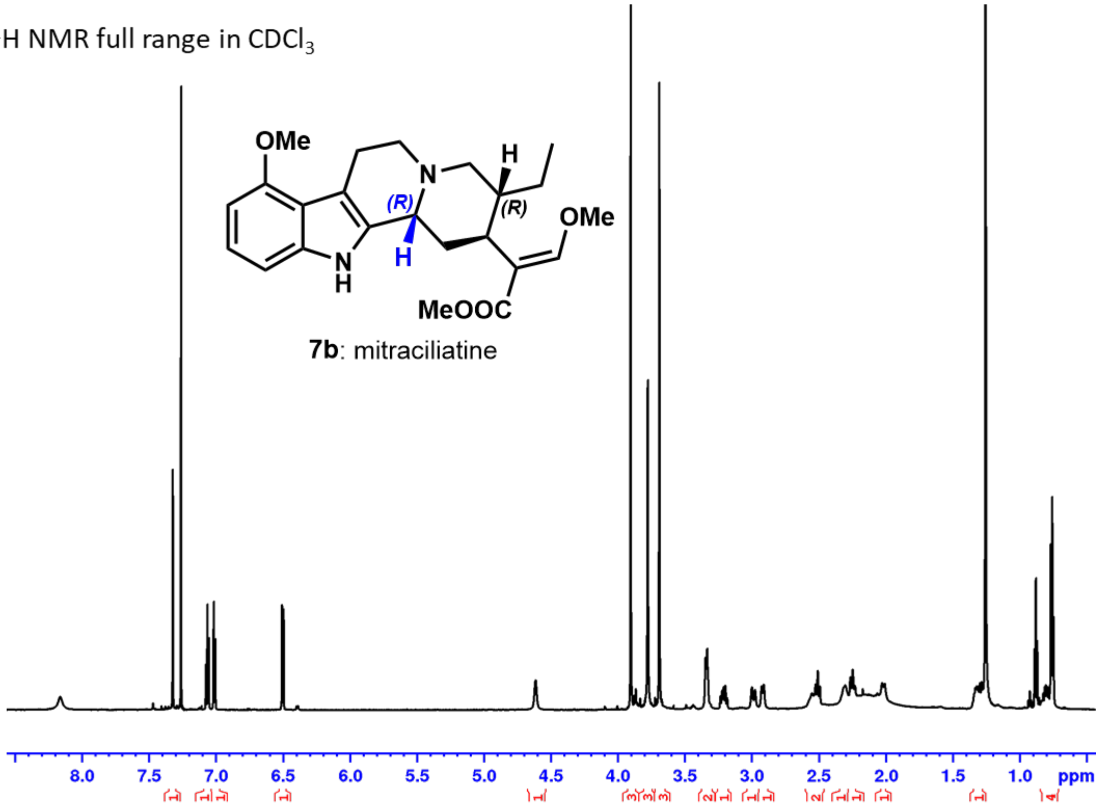


Phase sensitive HSQC (blue: CH, $CH_3$, green $CH_2$)/ HMBC (red) full range in $CDCl_3$

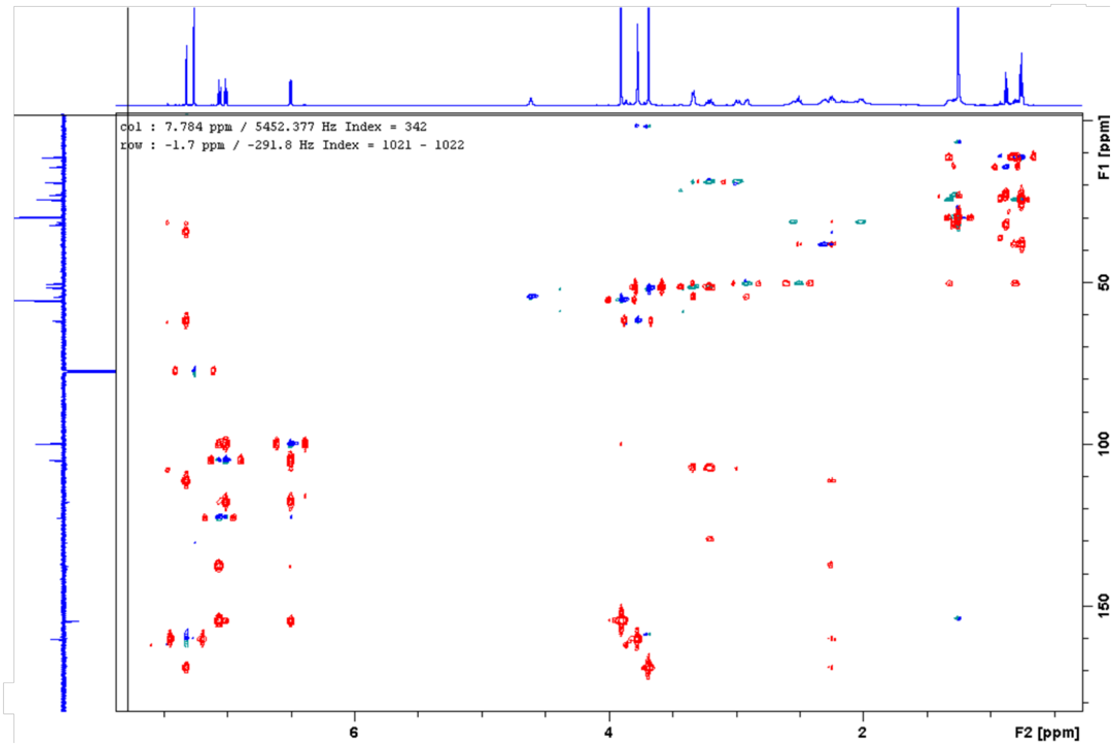


**Figure S55:** NMR spectra for 7b**:** mitraciliatine.

$^1H$ NMR full range in $CDCl_3$

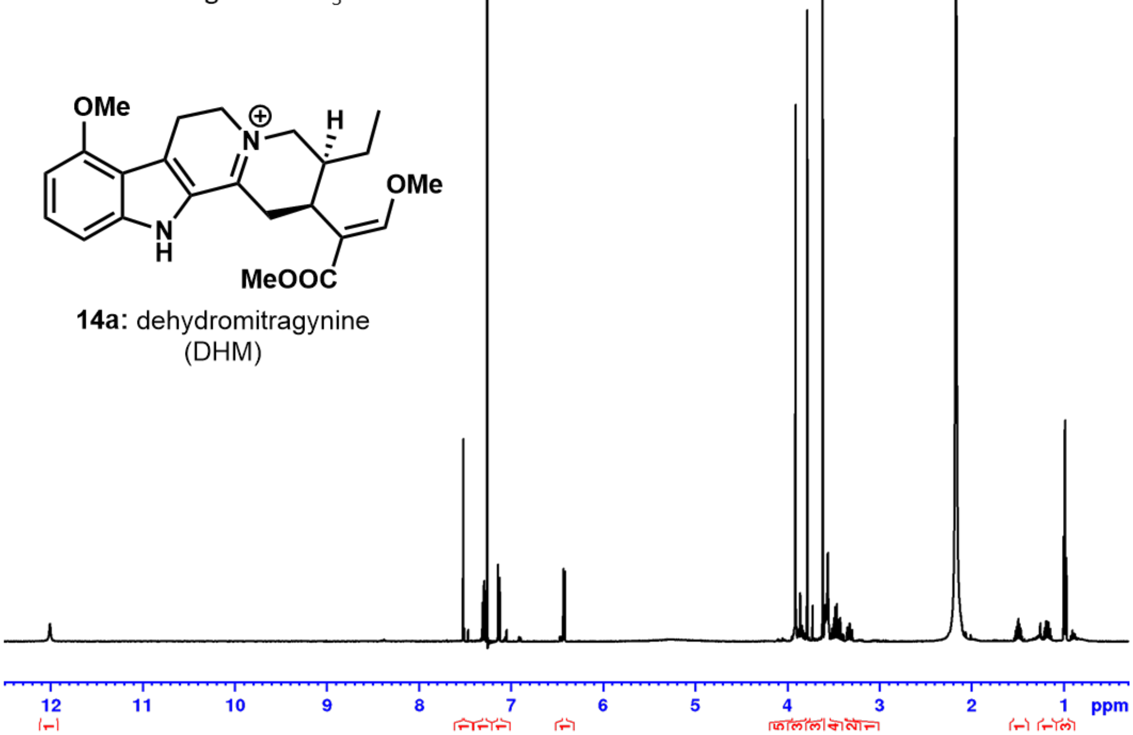


DEPTQ full range in $CDCl_3$

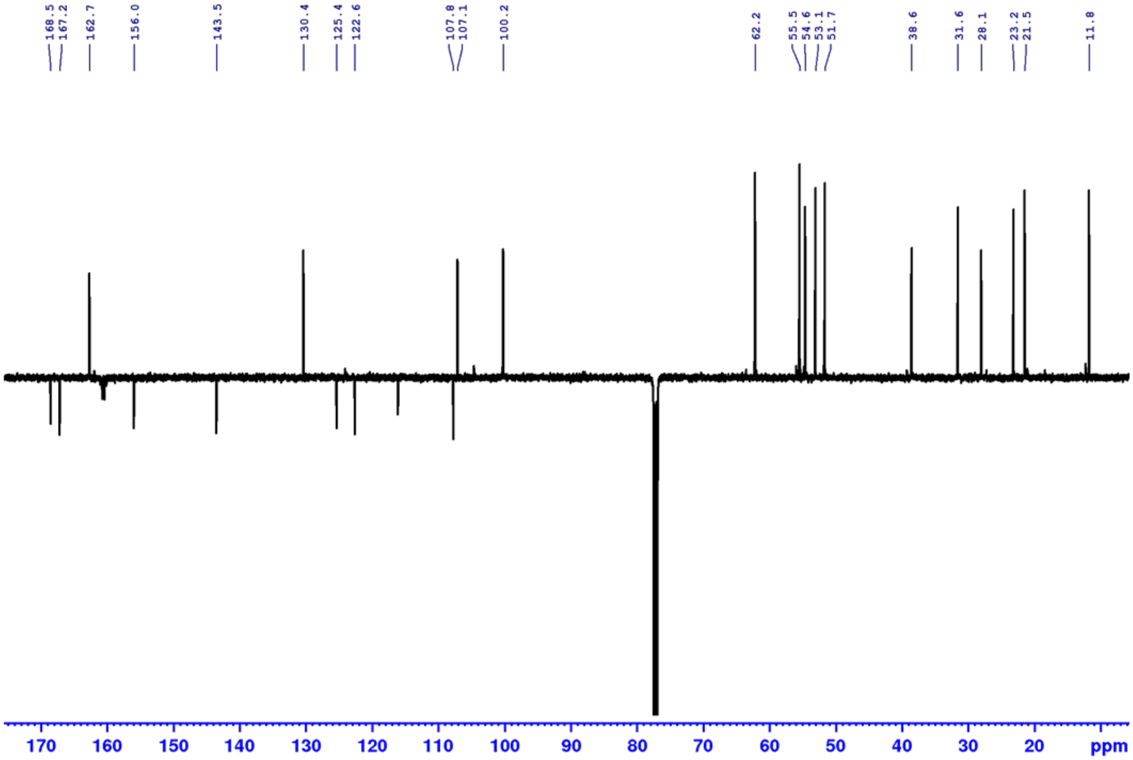


**Figure S56:** NMR spectra for **14a:** dehydromitragynine (DHM).

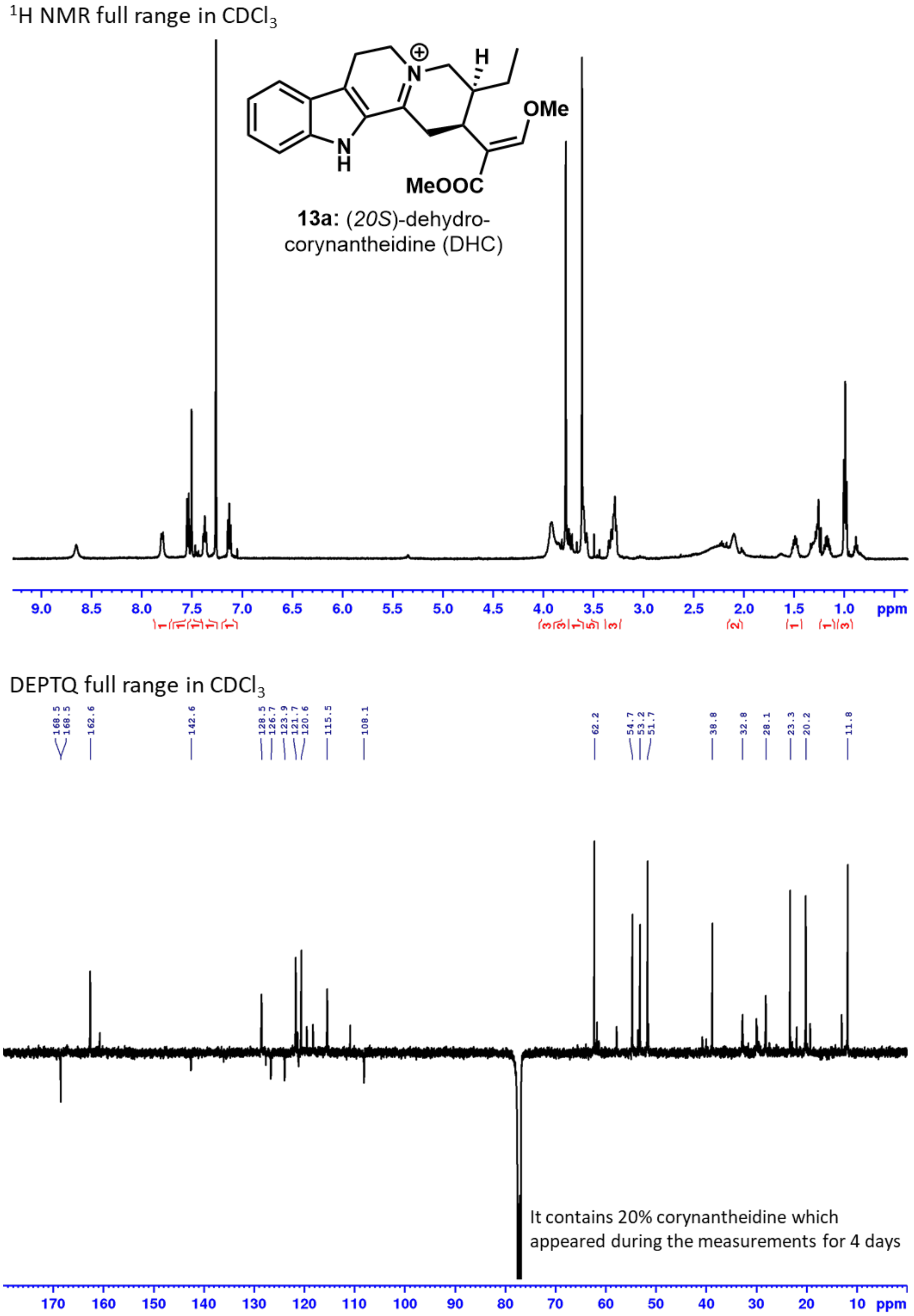


**Figure S57:** NMR spectra for **13a:** 20*S*-dehydrocorynantheidine (DHC).

Phase sensitive HSQC (black: CH, $CH_3$, red: $CH_2$) full range

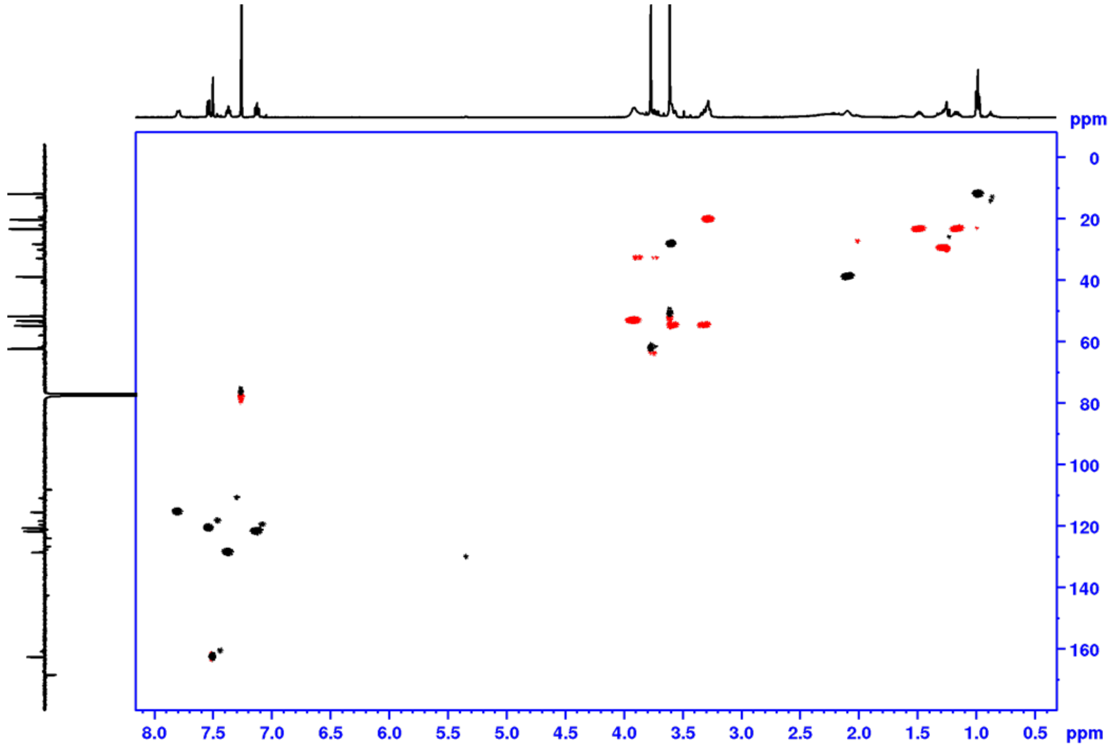


Phase sensitive HSQC (black: CH, $CH_3$, red: $CH_2$) aliphatic range

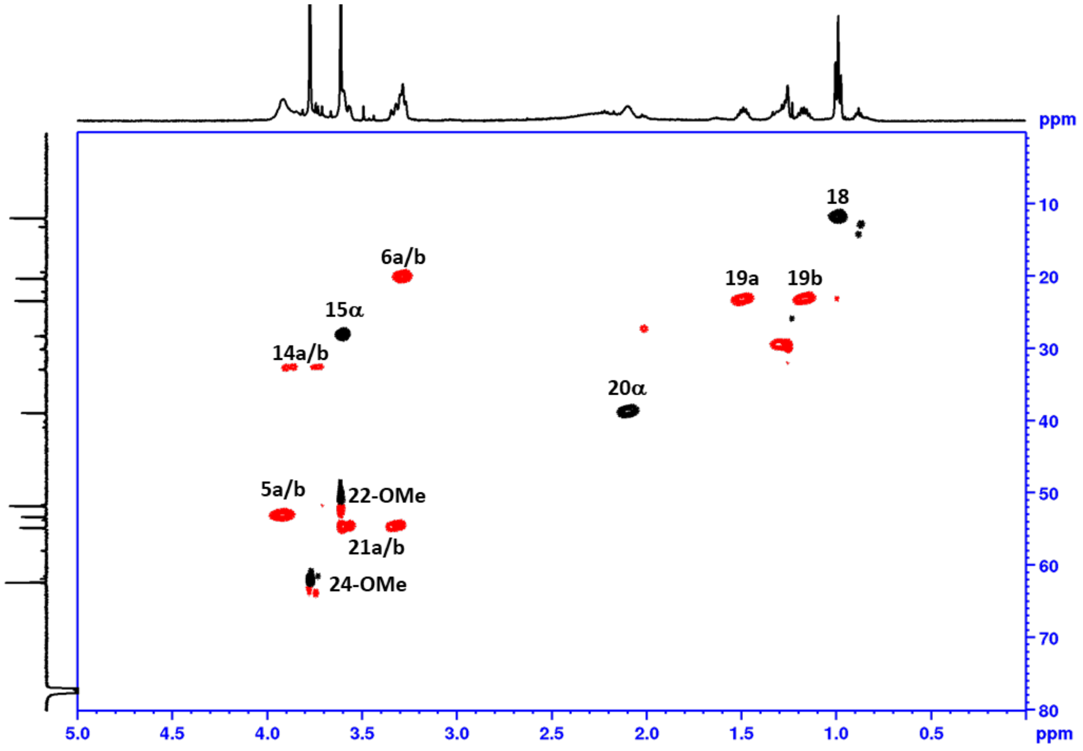


**Figure S58:** NMR spectra for **13a:** 20*S*-dehydrocorynantheidine (DHC).

HMBC

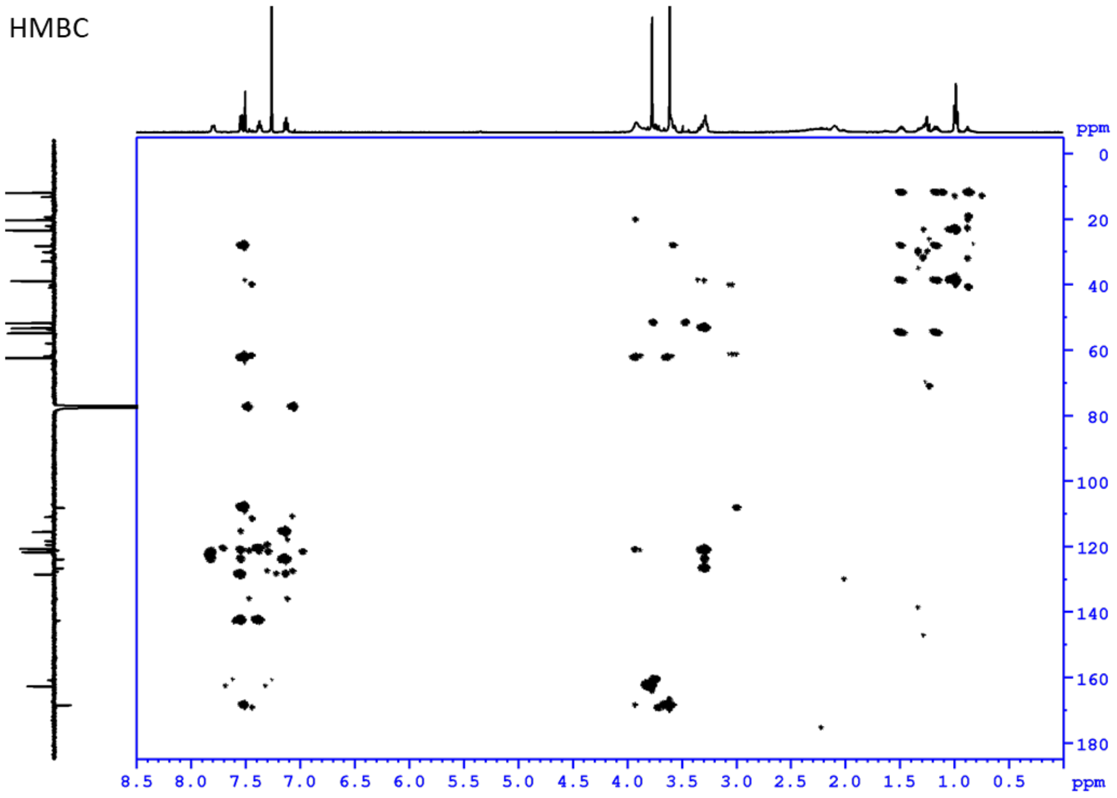


COSY

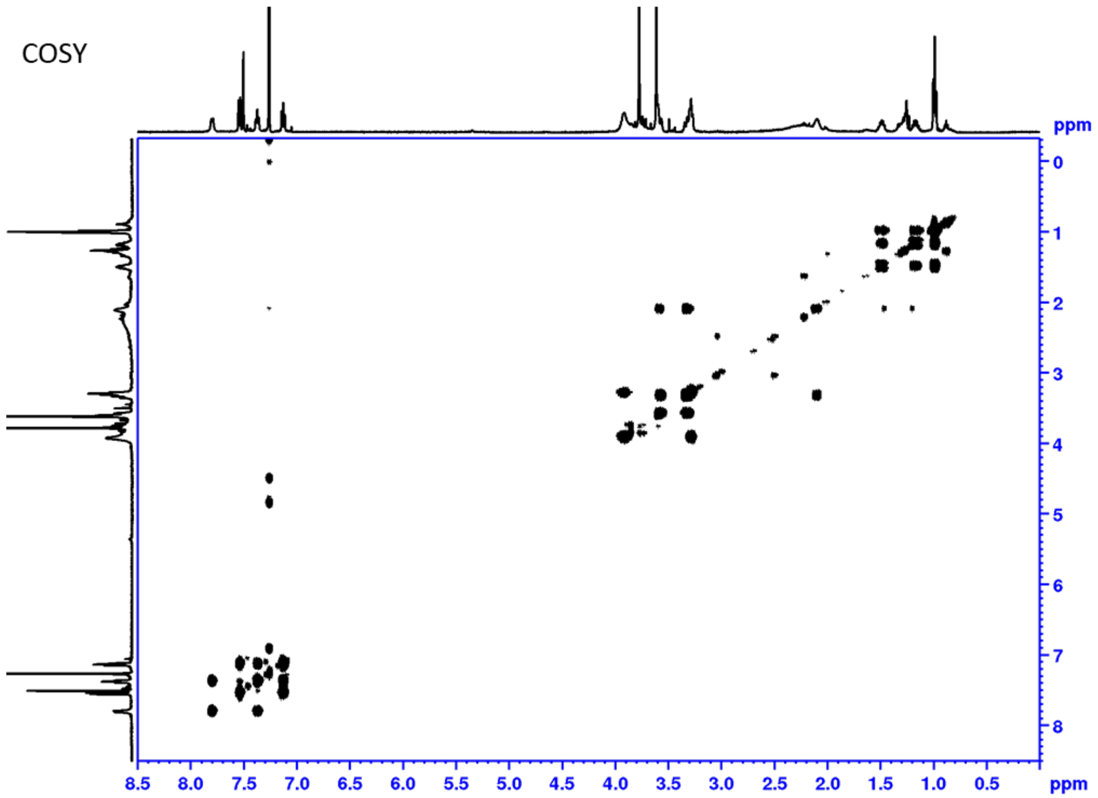


**Figure S59:** NMR spectra for **13a:** 20*S*-dehydrocorynantheidine (DHC).

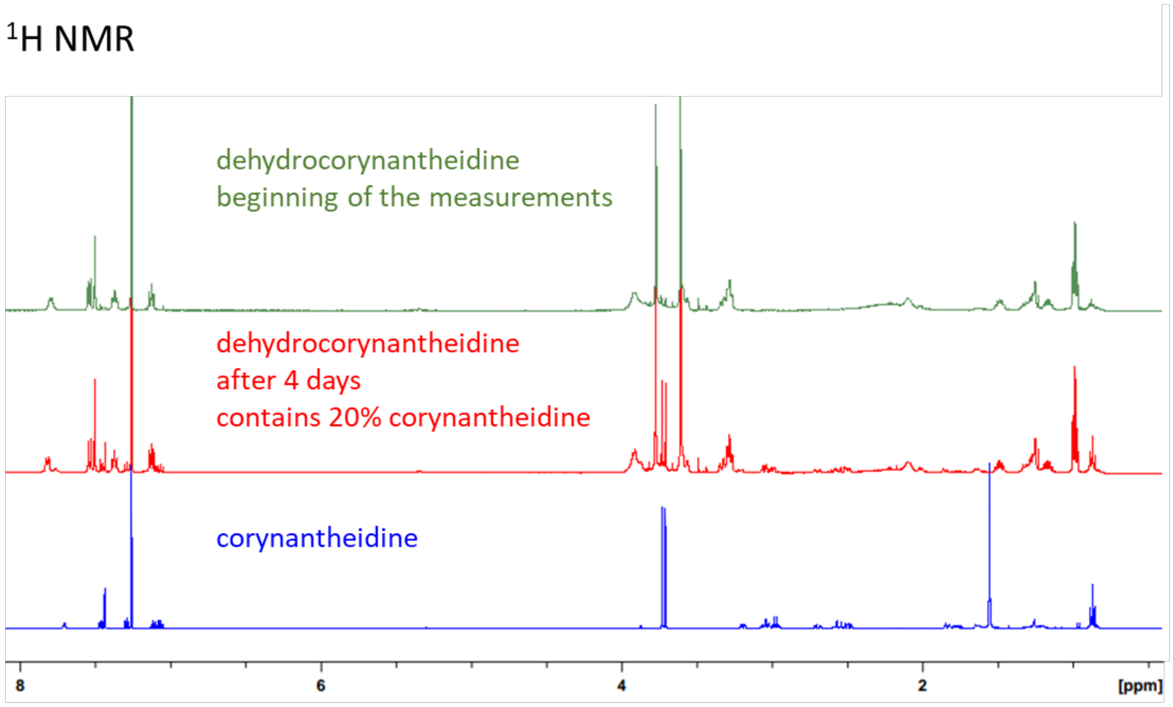


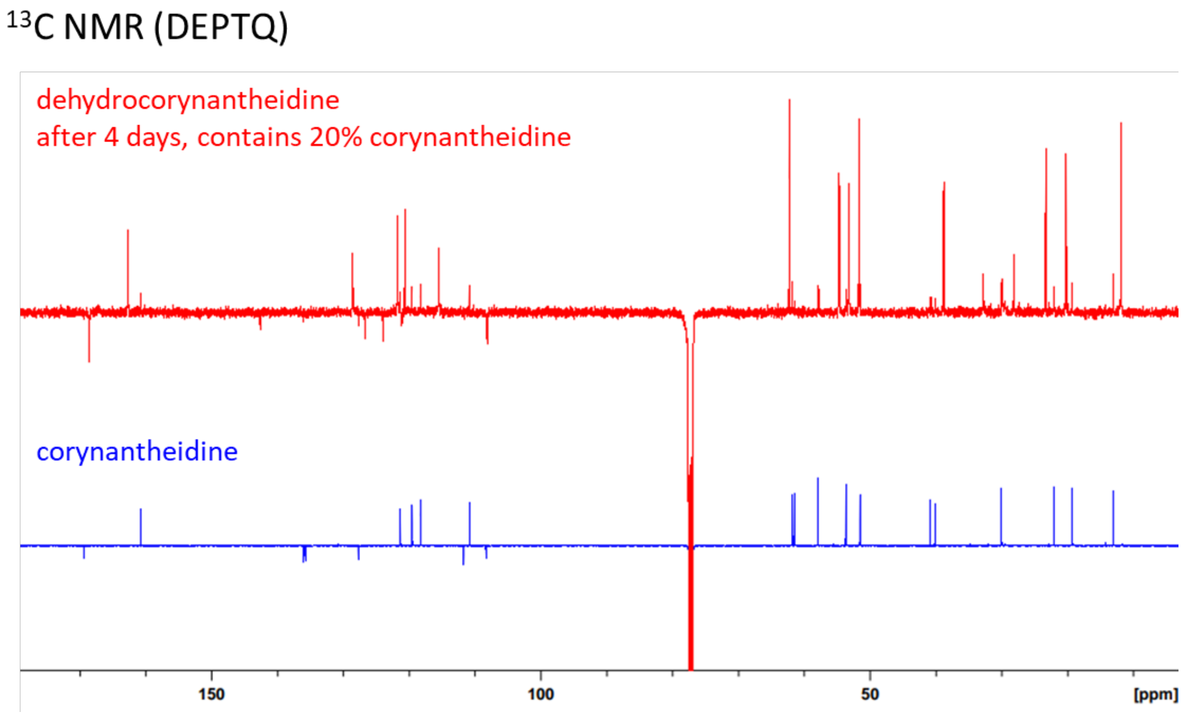


**Figure S60:** NMR spectra for **13a:** 20*S*-dehydrocorynantheidine (DHC) and **3a**: 20*S*-corynantheidine. Some conversion of **13a** to **3a** was observed during NMR analysis.

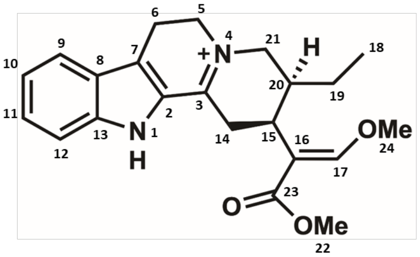


| pos. | $d_H$ | mult. | $J_{HH}$ | $d_C$ |
|---|---|---|---|---|
| 2 | - | - | - | 126.7 |
| 3 | - | - | - | 168.5 |
| 5a | 3.92 | m* | - | 53.2 |
| 5b | 3.92 | m* | - | 53.2 |
| 6a | 3.28 | m* | - | 20.2 |
| 6b | 3.28 | m* | - | 20.2 |
| 7 | - | - | - | 121.1 |
| 8 | - | - | - | 123.9 |
| 9 | 7.54 | d | 8.1 | 120.6 |
| 10 | 7.13 | dd | 8.1/7.6 | 121.7 |
| 11 | 7.37 | brdd | 7.9/7.6 | 128.5 |
| 12 | 7.80 | brd | 7.9 | 115.5 |
| 13 | - | - | - | 142.6 |
| 14a | 3.88 | m* | - | 32.8 |
| 14b | 3.74 | dd | 16.6/10.7 | 32.8 |
| 15a | 3.60 | m* | - | 28.1 |
| 16 | - | - | - | 108.1 |
| 17 | 7.50 | s | - | 162.6 |
| 18 | 0.99 | t | 7.3 | 11.8 |
| 19a | 1.49 | m | - | 23.3 |
| 19b | 1.17 | m | - | 23.3 |
| 20a | 2.10 | m | - | 38.8 |
| 21a | 3.58 | m* | - | 54.7 |
| 21b | 3.32 | m* | - | 54.7 |
| 22(OMe) | 3.61 | s | - | 51.7 |
| 23 | - | - | - | 168.5 |
| 24(OMe) | 3.77 | s | - | 62.2 |

**Figure S61:** NMR data for **13a**: (20S)-dehydrocorynantheidine in $CDCl_3$
**overlapped signal J unresolved, measured by 500 MHz NMR*

**Figure SX-1** NMR spectra for 3-isoajmalicine

$^1H$ NMR full range in $CDCl_3$

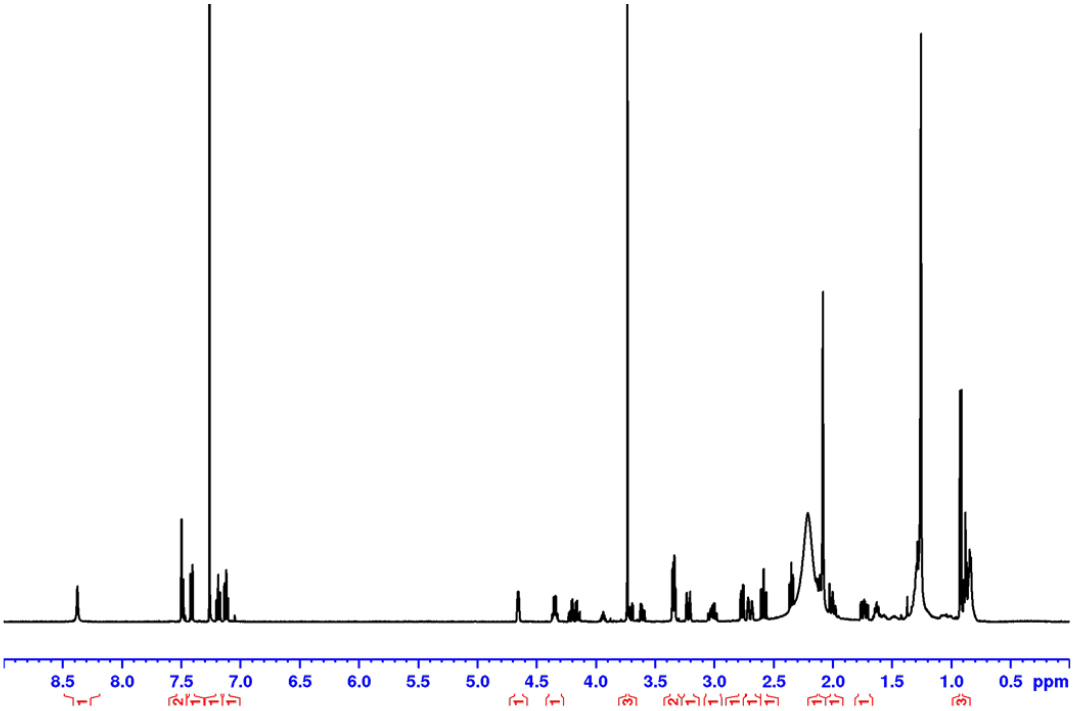


DEPTQ full range in $CDCl_3$

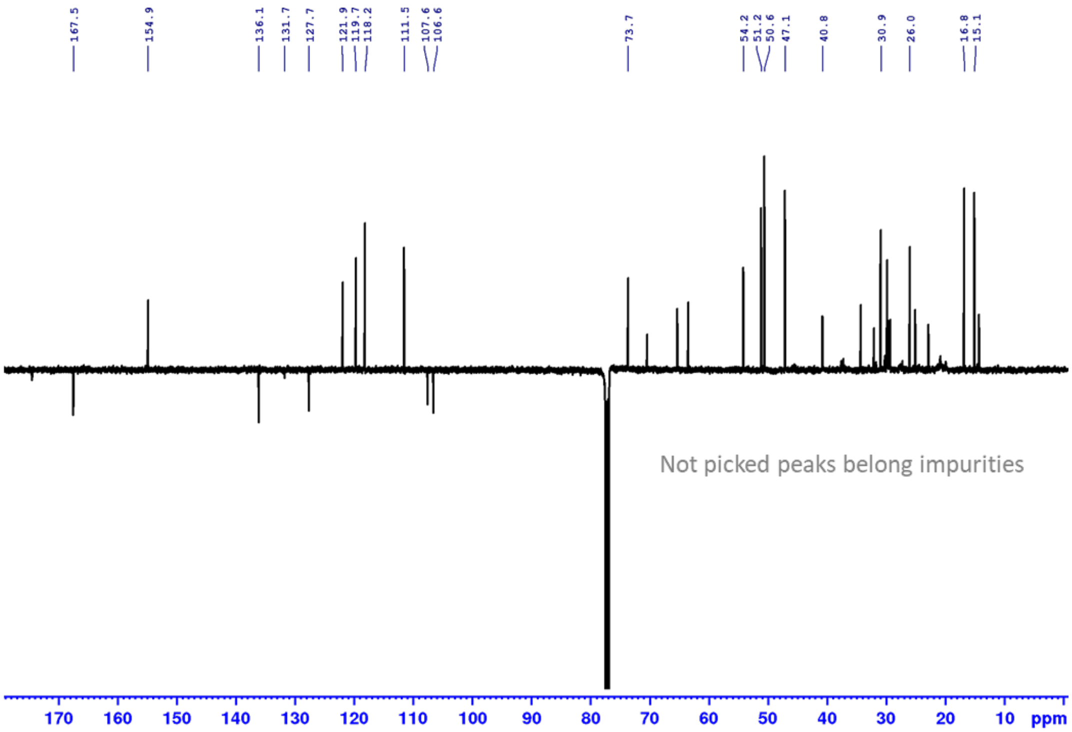


**Figure S62:** NMR spectra for **11b:** 3-epi-ajmalicine.

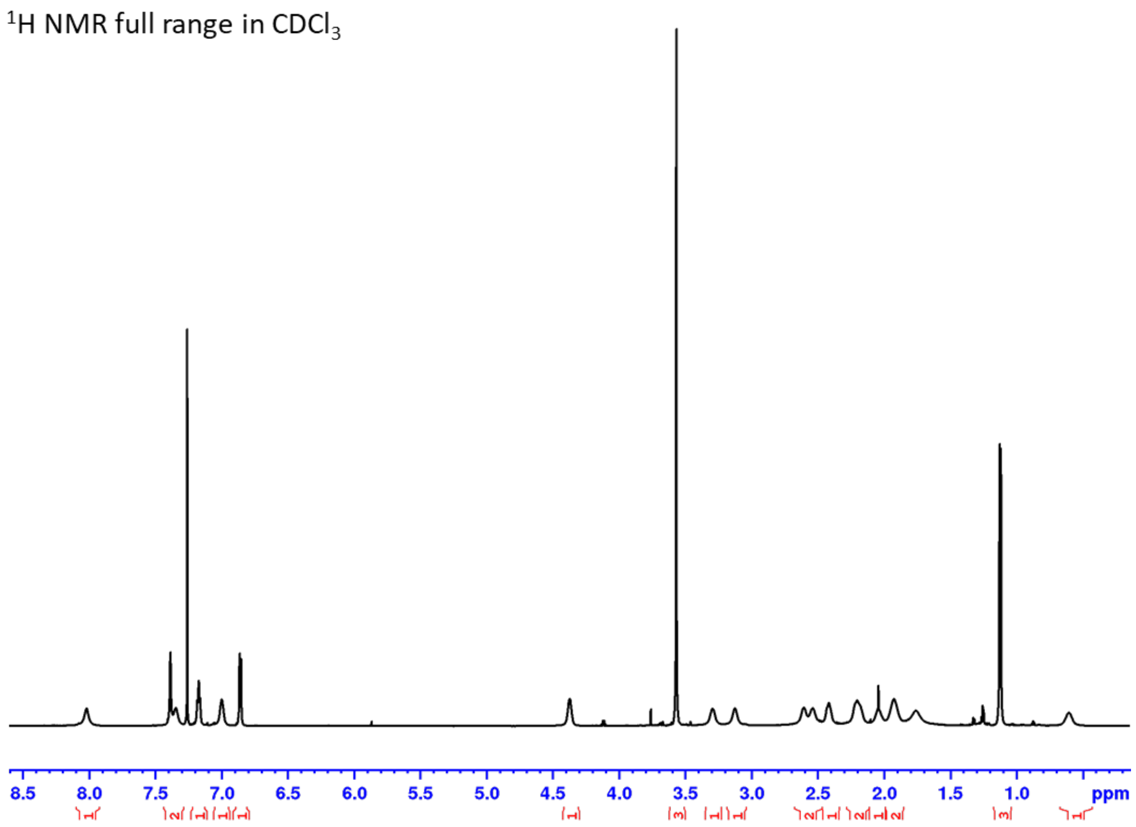


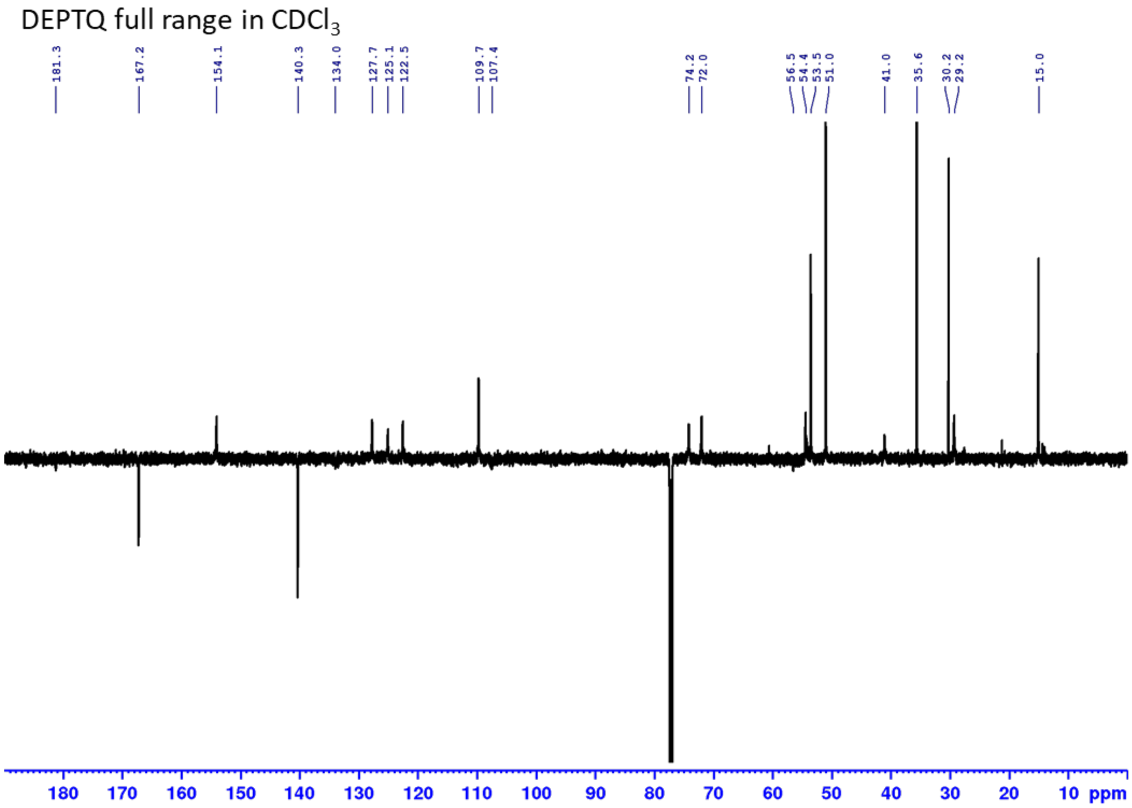


**Figure S63:** NMR spectra for **12b** (3*S*, 7*S*)**:** isomitraphylline.

<sup></sup>[1]H NMR full range in $CDCl_3$

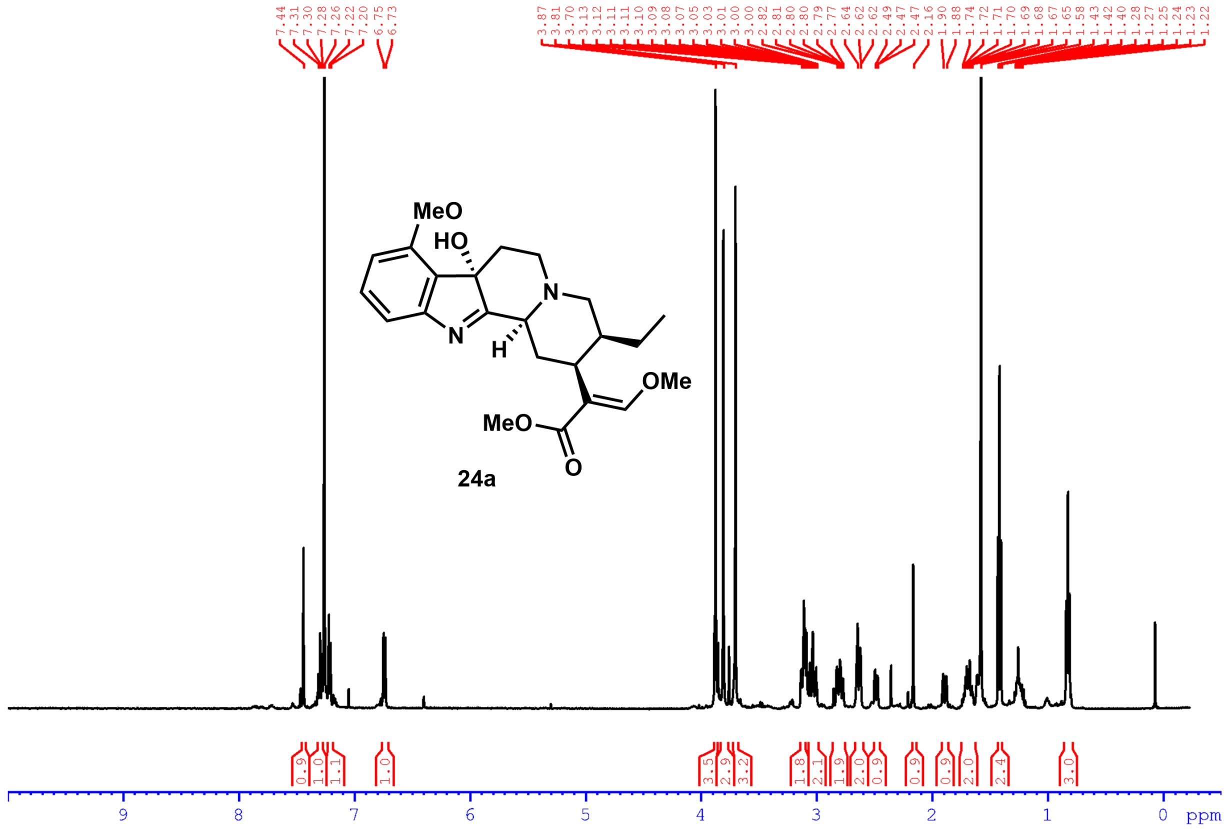

**Figure S64:** $^{1}$H NMR spectrum (500 MHz, $CDCl_3$) for 7-hydroxy mitragynine (**24a**).

$^1H$ NMR full range in $CDCl_3$

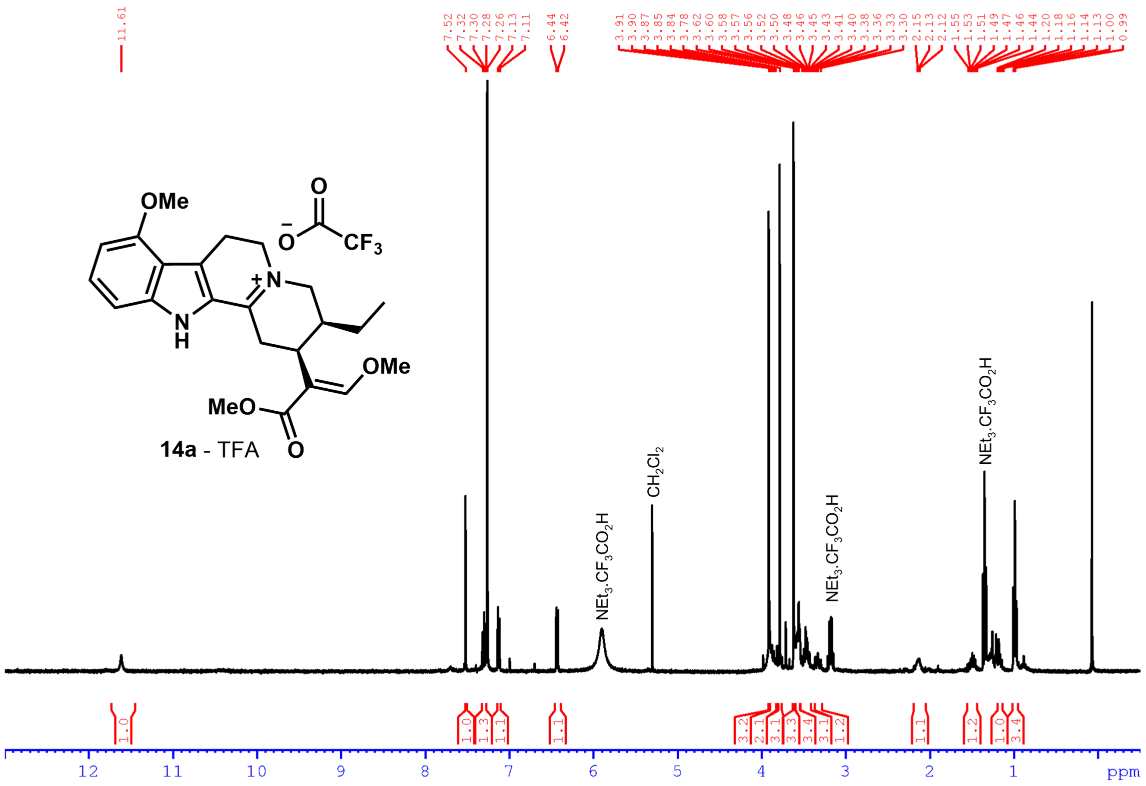

**Figure S65:** $^1H$ NMR spectrum (400 MHz, $CDCl_3$) for DHM-TFA (**14a**).

#### Supplementary References


(1) Nguyen, T. M.; Grzech, D.; Chung, K.; Xia, Z.; Nguyen, T.; Dang, T. T. Discovery of a Cytochrome P450 Enzyme Catalyzing the Formation of Spirooxindole Alkaloid Scaffold. *Front. Plant Sci.* **2023**, No. February, 1–7. https://doi.org/10.3389/fpls.2023.1125158.

(2) Laus, G.; Brössner, D.; Senn, G.; Wurst, K. Analysis of the Kinetics of Isomerization of Spiro Oxindole Alkaloids. *J. Chem. Soc. Perkin Trans. 2* **1996**, *9*, 1931–1936. https://doi.org/10.1039/P29960001931.

(3) Lombe, B. K.; Zhou, T.; Caputi, L.; Ploss, K.; Connor, S. E. O. Biosynthetic Origin of the Methoxy Group in Quinine and Related Alkaloids Angewandte. *Angew. Chemie* **2024**, *e202418306*, 1–8. https://doi.org/10.1002/anie.202418306.

(4) Haas, B. J.; Papanicolaou, A.; Yassour, M.; Grabherr, M.; Blood, P. D.; Bowden, J.; Couger, M. B.; Eccles, D.; Li, B.; Lieber, M.; Macmanes, M. D.; Ott, M.; Orvis, J.; Pochet, N.; Strozzi, F.; Weeks, N.; Westerman, R.; William, T.; Dewey, C. N.; Henschel, R.; Leduc, R. D.; Friedman, N.; Regev, A. De Novo Transcript Sequence Reconstruction from RNA-Seq Using the Trinity Platform for Reference Generation and Analysis. *Nat. Protoc.* **2013**, *8* (8), 1494–1512. https://doi.org/10.1038/nprot.2013.084.

(5) Martin, M. Cutadapt Removes Adapter Sequences from High-Throughput Sequencing Reads. *EMBnet J.* **2011**, 5–7.

(6) Gerlt, J. A.; Bouvier, J. T.; Davidson, D. B.; Imker, H. J.; Sadkhin, B.; Slater, D. R.; Whalen, K. L. Enzyme Function Initiative-Enzyme Similarity Tool ( EFI-EST ): A Web Tool for Generating Protein Sequence Similarity Networks. *BBA - Proteins Proteomics* **2015**, *1854* (8), 1019–1037. https://doi.org/10.1016/j.bbapap.2015.04.015.

(7) Cárdenas, P. D.; Sonawane, P. D.; Heinig, U.; Jozwiak, A.; Panda, S.; Abebie, B.; Kazachkova, Y.; Pliner, M.; Unger, T.; Wolf, D.; Ofner, I.; Vilaprinyo, E.; Meir, S.; Davydov, O.; Gal-on, A.; Burdman, S.; Giri, A.; Zamir, D. Pathways to Defense Metabolites and Evading Fruit Bitterness in Genus Solanum Evolved through 2-Oxoglutarate-Dependent Dioxygenases. *Nat. Commun.* **2019**, 1–13. https://doi.org/10.1038/s41467-019-13211-4.

(8) Sparkes, I. A.; Runions, J.; Kearns, A.; Hawes, C. Rapid, Transient Expression of Fluorescent Fusion Proteins in Tobacco Plants and Generation of Stably Transformed Plants. *Nat. Protoc.* **2006**, *1* (4), 2019–2025. https://doi.org/10.1038/nprot.2006.286.

(9) Caputi, L.; Franke, J.; Farrow, S. C.; Chung, K.; Payne, R. M. E.; Nguyen, T. D.; Dang, T. T. T.; Teto Carqueijeiro, I. S.; Koudounas, K.; De Bernonville, T. D.; Ameyaw, B.; Jones, D. M.; Curcino Vieira, I. J.; Courdavault, V.; O'Connor, S. E. Missing Enzymes in the Biosynthesis of the Anticancer Drug Vinblastine in Madagascar Periwinkle. *Science (80-. ).* **2018**, *360* (6394), 1235–1239. https://doi.org/10.1126/science.aat4100.

(10) Mcdonald, A. D.; Higgins, P. M.; Buller, A. R. Substrate Multiplexed Protein Engineering Facilitates Promiscuous Biocatalytic Synthesis. *Nat. Commun.* **2022**, *13*, 5242. https://doi.org/10.1038/s41467-022-32789-w.

(11) Kruegel, A. C.; Sames, D.; Javitch, J. A.; Majumdar, S. Deuterated Mitragynine Analogs as Safer Opioid Modulators in the Mitragynine Class. *Espacenet* **2020**, *WO20201602*.

(12) Chakraborty, S.; Uprety, R.; Slocum, S. T.; Irie, T.; Rouzic, V. Le; Li, X.; Wilson, L. L.; Scouller, B.; Alder, A. F.; Kruegel, A. C.; Ansono, M.; Varadi, A.; Eans, S. O.; Hunkele, A.; Allaoa, A.; Kalra, S.; Xu, J.; Pan, Y. X.; Pintar, J.; Kivell, B. M.; Pasternak, G. W.; Cameron, M. D.; Mclaughlin, J. P.; Sames, D. Oxidative Metabolism as a Modulator of Kratom's Biological Actions. *J. Med. Chem.* **2021**, *64*, 16553–16572. https://doi.org/10.1021/acs.jmedchem.1c01111.

(13) Misa, J.; Billingsley, J. M.; Niwa, K.; Yu, R. K.; Tang, Y. Engineered Production of Strictosidine and Analogues in Yeast. *ACS Synth. Biol.* **2022**, *11* (4), 1639–1649. https://doi.org/10.1021/acssynbio.2c00037.

(14) Wanner, M. J.; Claveau, E.; Van Maarseveen, J. H.; Hiemstra, H. Enantioselective Syntheses of

Corynanthe Alkaloids by Chiral Brønsted Acid and Palladium Catalysis. *Chem. - A Eur. J.* **2011**, *17* (49), 13680–13683. https://doi.org/10.1002/chem.201103150.

(15) Stærk, D.; Lemmich, E.; Christensen, J.; Kharazmi, A.; Olsen, C. E.; Jaroszewski, J. W. Leishmanicidal, Antiplasmodial and Cytotoxic Activity of Indole Alkaloids from Corynanthe Pachyceras. *Planta Med* **2000**, *66*, 531–536.

(16) Ren, J.; Ding, S. H.; Li, X. N.; Zhao, Q. S. Unified Strategy Enables the Collective Syntheses of Structurally Diverse Indole Alkaloids. *J. Am. Chem. Soc.* **2024**, *146* (11), 7616–7627. https://doi.org/10.1021/jacs.3c13869.

(17) Lounasmaa, M., Jokela, R., Laine, C., & Hanhinen, P. Preparation of (+-)-Hirsutine and (+-)-3-Isocorynantheidine. *Heterocycles* **1998**, *49*, 445–450.

(18) Flores-bocanegra, L.; Raja, H. A.; Graf, T. N.; Augustinovic, M.; Wallace, E. D.; Hematian, S.; Kellogg, J. J.; Todd, D. A.; Cech, N. B.; Oberlies, N. H. The Chemistry of Kratom [Mitragyna Speciosa]: Updated Characterization Data and Methods to Elucidate Indole and Oxindole Alkaloids. *J. Nat. Prod.* **2020**, *83*, 2165–2177. https://doi.org/10.1021/acs.jnatprod.0c00257.

(19) Paradowska, K.; Wolniak, M.; Pisklak, M.; Gliński, J. A.; Davey, M. H.; Wawer, I. 13C, 15N CPMAS NMR and GIAO DFT Calculations of Stereoisomeric Oxindole Alkaloids from Cat's Claw (Uncaria Tomentosa). *Solid State Nucl. Magn. Reson.* **2008**, *34* (4), 202–209. https://doi.org/10.1016/j.ssnmr.2008.10.002.

(20) Kruegel, A. C.; Gassaway, M. M.; Kapoor, A.; Váradi, A.; Majumdar, S.; Filizola, M.; Javitch, J. A.; Sames, D. Synthetic and Receptor Signaling Explorations of the Mitragyna Alkaloids: Mitragynine as an Atypical Molecular Framework for Opioid Receptor Modulators. *J. Am. Chem. Soc.* **2016**, *138* (21), 6754–6764. https://doi.org/10.1021/jacs.6b00360.

(21) Fulmer, G. R.; Miller, A. J. M.; Sherden, N. H.; Gottlieb, H. E.; Nudelman, A.; Stoltz, B. M.; Bercaw, J. E.; Goldberg, K. I. NMR Chemical Shifts of Trace Impurities: Common Laboratory Solvents, Organics, and Gases in Deuterated Solvents Relevant to the Organometallic Chemist. *Organometallics* **2010**, *29* (9), 2176–2179. https://doi.org/10.1021/om100106e.

(22) Andreev, I. A.; Boichenko, M. A.; Ratmanova, N. K.; Ivanova, O. A.; Levina, I. I.; Khrustalev, V. N.; Sedov, I. A.; Trushkov, I. V. 4-(Dimethylamino)Pyridinium Azide in Protic Ionic Liquid Media as a Stable Equivalent of Hydrazoic Acid. *Adv. Synth. Catal.* **2022**, *364* (14), 2403–2415. https://doi.org/10.1002/adsc.202200486.
